# Gene-expression profiling of single cells from archival tissue with laser-capture microdissection and Smart-3SEQ

**DOI:** 10.1101/207340

**Authors:** Joseph W. Foley, Chunfang Zhu, Philippe Jolivet, Shirley X. Zhu, Peipei Lu, Michael J. Meaney, Robert B. West

**Author notes:** Correponding author.

## Abstract

RNA sequencing (RNA-seq) is a sensitive and accurate method for quantifying gene expression. Small samples or those whose RNA is degraded, such as formalin-fixed, paraffin-embedded (FFPE) tissue, remain challenging to study with nonspecialized RNA-seq protocols. Here we present a new method, Smart-3SEQ, that accurately quantifies transcript abundance even with small amounts of total RNA and effectively characterizes small samples extracted by laser-capture microdissection (LCM) from FFPE tissue. We also obtain distinct biological profiles from FFPE single cells, which have been impossible to study with previous RNA-seq protocols, and we use these data to identify possible new macrophage phenotypes associated with the tumor microenvironment. We propose Smart-3SEQ as a highly cost-effective method to enable large gene-expression profiling experiments unconstrained by sample size and tissue availability. In particular, Smart-3SEQ’s compatibility with FFPE tissue unlocks an enormous number of archived clinical samples, and combined with LCM it allows unprecedented studies of small cell populations and single cells isolated by their *in situ* context.

## Introduction

*Omnis cellula e cellula* (every cell from a cell): this idea promoted by Rudolf Virchow, the grandfather of modern pathology, rings no less true today than 160 years ago [1]. Cell-centered concepts are emerging that suggest it is not just the genomic alterations of a cell but its developmental state that are critical to our understanding of disease. Technologies to study individual single cells, not complex aggregates of multiple cell types, will give us a greater understanding of the biology of many diseases. Multiple large-scale collaborative projects are underway to understand individual cell biology in both normal and pathological tissues [2, 3].

Single-cell gene-expression profiling is essential for these endeavors. Prominent new methods rely on fresh tissue to generate disaggregated intact single cells that can be profiled using microfluidic technologies [4, 5]. These approaches are generating extensive information on new cell types and cell-specific transcriptome profiles. Such information will have a significant impact on the clinical management of disease. However, this impact is diminished by the limitations of fresh clinical material. Fresh clinical material is difficult to obtain for research, cannot be banked, and the selection of material for analysis is based on *a priori* clinical knowledge and gross examination with no cellular context. Moreover, the isolation process often leads to selective degradation of specific cell types leading to a bias in the profiled cell populations. For these reasons, it is not suitable for many clinical study designs.

We have developed an RNA-seq method that addresses these problems. Our method, Smart-3SEQ, is robust in archival formalin-fixed, paraffin-embedded (FFPE) material and allows microscopic selection of specific cells. It combines the template-switching SMART method [6, 7] and protocol optimizations of Smart-seq2 [8] with the streamlined 3′ end-targeting approach of 3SEQ [9], yielding a simpler procedure than any of those. In addition, Smart-3SEQ incorporates unique molecular identifiers to increase the accuracy of transcript counting with low input amounts [10]. This method has conceptual similarities to the prototype RNA-seq method used to demonstrate UMIs, but with a streamlined protocol and optimizations for laser capture microdissection and FFPE material. Thus our protocol is cost-effective in reagent usage and working time, sensitive to single-cell amounts of RNA, and robust to degraded samples. Here we demonstrate that Smart-3SEQ quantifies transcript abundance accurately with a wide range of amounts of input; in particular it can create libraries from single cells dissected out of tissue with degraded RNA, which has previously only been possible with high-quality tissue samples [11], and this enables the study of cells in the tumor microenvironment with clinical samples.

## Results

### The Smart-3SEQ method

Previous single-cell RNA-seq protocols, Smart-seq [7] and Smart-seq2 [8], generate cDNA libraries by the SMART method [6]: they take advantage of Moloney murine leukemia virus–derived reverse transcriptase’s “template switching” ability to generate full-length, double-stranded cDNA molecules from RNA templates in a single incubation. However, when the original RNA is degraded, there may not be full-length molecules available to reverse-transcribe, so RNA integrity is very important with these methods. In contrast, the 3SEQ method [9] is optimal for degraded RNA such as that from FFPE tissue. In 3SEQ, the RNA is fragmented before reverse transcription, eliminating the difference between intact and degraded samples. Reverse transcription is still primed by an anchored oligo(dT) primer, so from each transcript only the fragment containing the beginning of the poly(A) tail is sequenced; this approach, also called digital gene expression, generates a short identifying “tag” sequence rather than read the full length of the transcript. Smart-3SEQ combines this 3′-targeted approach with template-switching in a streamlined and sensitive protocol (Figures 1, S1, S2; see Supplemental File 1 for a technical description and Table S1 for a step-by-step comparison with similar protocols).

**Figure 1:**
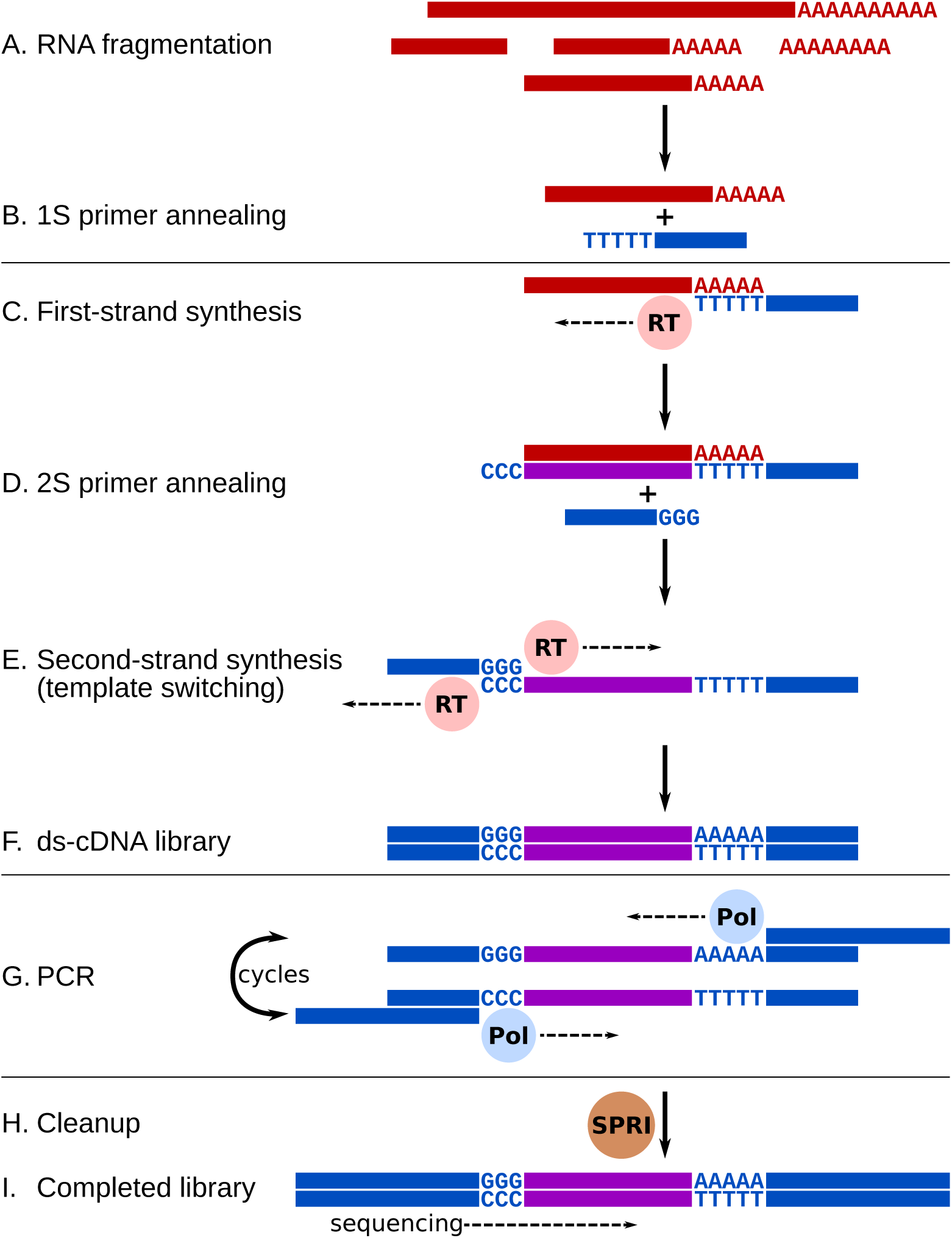
Conceptual diagram of the Smart-3SEQ library preparation method. Hands-on steps are separated by horizontal lines. A: Total RNA is denatured and fragmented by hydrolysis. B: The oligo(dT) primer, including a partial sequencing adapter, anneals at the beginning of the poly(A) tail. C: Reverse transcriptase synthesizes first-strand cDNA from the RNA template, and adds non-template dC at the end of the new strand. D: The second primer, which includes a second partial sequencing adapter, anneals to the new dC overhang. E: Reverse transcriptase synthesizes the second cDNA strand using the first as a template. F: After steps C–E, which occur consecutively in one incubation, the result is a double-stranded cDNA library with partial sequencing adapters at both ends. G: PCR with long primers amplifies the library and extends the adapters to full length, including multiplexing indexes. H: The only cleanup step in the protocol uses paramagnetic SPRI beads to purify the amplified library while excluding adapter dimers and short inserts. I: The final library contains the unknown cDNA sequence between the two sequencing adapters. The cDNA is sequenced in the orientation of the original RNA, yielding reads upstream of the end of the transcript (Figure S3). Note: See Supplemental File 1 for a detailed technical description, Figure S1 for a simplified diagram showing the practical workflow, and Figure S2 for a detailed technical diagram.

Smart-3SEQ has numerous practical advantages for many different applications of gene-expression profiling independent of its use with FFPE tissue (Table S2). Because of the method’s efficiency, with no inefficient dsDNA ligation and only one cleanup step, it is sensitive at very small amounts of starting RNA. The short number of steps and small reaction volumes allow a batch of libraries to be prepared in half a day with high throughput in a 96-well plate, at a reagent cost of about 5 USD per sample (Supplemental File 5). Library yields are predictable enough that the optimal number of PCR cycles can be reliably estimated from the amount of input material over five orders of magnitude. Furthermore, because Smart-3SEQ reads only a single fragment of each transcript, the one containing the polyadenylation site, it is not sensitive to transcript length, which is a confounder for whole-transcript RNA-seq [12]. Thus Smart-3SEQ allows simple and accurate quantification of transcript abundance; however, it cannot report information about splicing or genotypes, unless the splice junction or polymorphism occurs near the end of the gene, and it cannot detect non-polyadenylated transcripts. Like 3SEQ, because RNA fragmentation is the first step, it is also robust to pre-fragmented RNA, including damaged RNA from FFPE samples.

### Validation with reference samples

We validated the accuracy and sensitivity of the Smart-3SEQ method by using it to quantify the reference RNAs used in the SEQC Consortium’s benchmarking study [13, 14], though with lower amounts of RNA input. In the first experiment, we used ERCC Mixes 1 and 2, which contain 92 *in vitro*–transcribed, polyadenylated transcripts at known concentrations, which span 6 orders of magnitude and differ between the two mixes [15]. We prepared tenfold serial dilutions at 1/10–1/100,000 the stock concentration, then created a library from 1μL of each (10.4 fmol–1.04 amol, 6.23 billion–623,000 molecules); at each dilution level we tested two different numbers of PCR cycles (Figure S4).

The proportion of reads aligned to the reference sequences decreased mainly with RNA input, but the number of PCR cycles also made a large difference (Figure S5A). 99.99% of aligned reads were in the expected forward-strand orientation. The abundance of each transcript as measured by Smart-3SEQ corresponded linearly with its expected copy number in the sample (Figure 2A; average *r’* = 0.990 in the highest-input, lowest-PCR samples; see Methods for definition of *r’*). The lower information content of the lower-input samples was visible as an increase in the number of duplicate reads (Figure S5B): if we assume the lowest-input samples are sequenced to completion so that all unique sequences in the library have been detected at least once, our average observation of 189,322 non-duplicate reads among the low-PCR libraries implies a capture efficiency of 30%; for comparison, the theoretical maximum efficiency of GGG template-switching has been estimated as 46% [16]. As the amount of RNA input decreased, the accuracy of the measurements worsened for low-abundance transcripts (Figure S6; for 1 amol RNA, average *r’* = 0.926). Likewise the replicate correlations were very high for high inputs but decreased for low inputs (Figure S7), and the signal loss compared to the first dilution increased with subsequent dilutions (Figure S8). However, in the lowest-input samples the expected copy number of several transcripts was less than 1, so we expect considerable sampling error even in the true copy numbers of low-abundance transcripts, which is magnified by the recursive sampling of the serial-dilution process.

**Figure 2:**
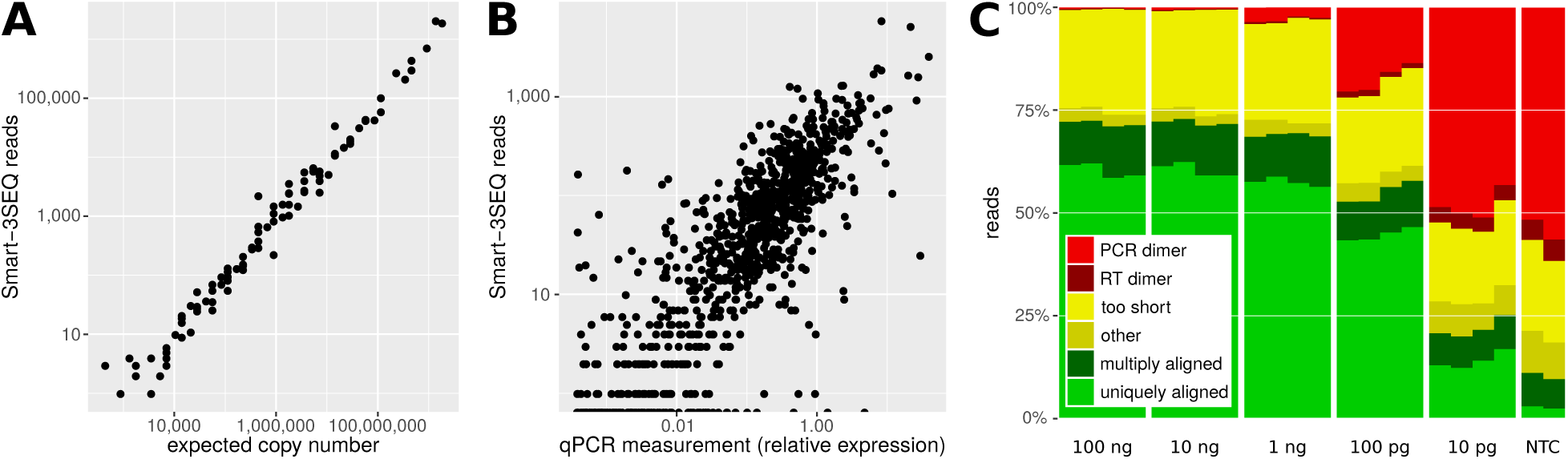
Technical validation of Smart-3SEQ with reference RNAs. A: Standard curve from ERCC transcripts, *r'* = 0.990. Each point represents one transcript sequence; “expected copy number” is the estimated number of copies of that transcript in the RNA sample (1μL at 1/10 dilution, 10fmol total), and “Smart-3SEQ reads” is the number of post-filter reads aligned uniquely to any part of that transcript’s sequence in the expected orientation (6.4M). Data shown are from the first replicate of the condition with 10fmol ERCC mix 2 and 7 PCR cycles, which had the highest read count for that condition; data from all samples are shown in Figure S6. B: Comparison with TaqMan qPCR quantification of human transcripts, *r'* = 0.854. qPCR measurements are normalized to the expression of RNA polymerase II. Data shown are from the first replicate of 100 ng UHRR, which had the highest read count for that amount of input RNA (2.7M sense-aligned to annotated genes); data from all samples are shown in Figure S13. C: Alignability of Smart-3SEQ reads from human reference RNA dilutions and no-template controls.

In a second validation experiment, we used the Human Brain Reference RNA (HBRR) and the Universal Human Reference RNA (UHRR), which were also used in the SEQC benchmarks. We mixed in the ERCC standards at the same relative concentration used by SEQC (1/500 ERCC stock concentration with 100ng/μL human RNA) to make our RNA samples comparable with theirs, then we serially diluted this mixture down to 10 pg total RNA (Figure S9), roughly the amount in a single mammalian cell [7, 8, 17, 18, 19, 20]. As in the previous experiment, we saw a high proportion of alignable reads in the high-input samples, which gradually declined as the number of PCR cycles approached 20 and then declined more steeply (Figure 2C); the lowest-input libraries accumulated some short sequence artifacts, but most of these did not align to the genome and a negligible share aligned to the transcriptome (Figure S10) so they did not affect measurements of gene expression. The diversity of the libraries also decreased with input amount, but the genome loci detected by the reads stayed fairly constant (Figure S11). Of the reads that aligned outside transcripts’ 3' ends, only a small fraction aligned upstream of genome-encoded poly(A) sites (Figure S12), suggesting off-target priming was not the major cause of these artifacts. The true abundances of the transcripts in these human samples are not known *a priori*, but it is still possible to validate Smart-3SEQ’s measurements by comparing them with those of an alternative method, qPCR, on the same reference RNA samples. Smart-3SEQ showed a strong correlation (average Spearman’s *ρ* = 0.845 for high-input samples; Figures 2B, S13) with TaqMan-based qPCR measurements of 999 genes [21] and with SYBR-based qPCR of 20,801 genes [13] (average high-input *ρ* = −0.827; Figure S14).

We compared the accuracy of Smart-3SEQ side-by-side with the SEQC data from standard RNA-seq performed on the same ERCC transcripts and human reference RNAs (250ng total RNA), as well as Smart-seq performed on only 10 pg of human reference RNAs with Smart-seq as implemented in the SMART-Seq v4 kit from Takara Bio USA (pers. comm.). With moderate amounts of input RNA, Smart-3SEQ showed equivalent accuracy to RNA-seq in the ERCC standard curves (Figures S15, S16). Likewise with ample RNA, Smart-3SEQ corresponded with both SYBR- and TaqMan-based qPCR roughly as well as high-input RNA-seq (Figures S17, S18), and with low input it performed slightly worse than Smart-seq (Figures S20, S21), though both methods performed worse with low input than Smart-3SEQ and RNA-seq did with high input. Although the direct correlation between Smart-3SEQ and the other methods was lower (Figures S19, S22), the qPCR platforms were equally correlated with each other as they were with the sequencing platforms (Figure S23). Thus we conclude that although Smart-3SEQ and whole-transcript RNA-seq may have different error profiles, they both quantify biological signal equally well.

To test Smart-3SEQ’s robustness to degraded RNA, we repeated the serial dilutions with pairs of fresh-frozen and FFPE samples. That is, from each fresh-frozen tissue sample, an FFPE block was also prepared, as previously described [9]. We used two cores each of solitary fibrous tumor and pigmented villonodular synovitis, all from different clinical subjects. Despite the large differences in RNA integrity, there were only small differences in the qualities of the final libraries (Figure S24), likely attributable to shorter cDNA inserts from FFPE (Figure S25). Although there was some bias in the gene-expression profiles according to the preparation method, the separation between biological categories was much stronger. The artifactual signal in the no-input controls, which may represent contamination of the library preparation reagents, cross-contamination of spuriously assigned sequence reads from the other libraries, or spuriously aligned sequencing artifacts, was distinct from the lowest-input samples (Figure S26). These results show that Smart-3SEQ maintains its sensitivity and accuracy even with low-quality RNA samples.

### Demonstration with microdissected cells

To take advantage of Smart-3SEQ’s compatibility with small amounts of degraded RNA, we used the method to study single-cell gene expression in the context of the tumor microenvironment, which is a mix of benign and malignant cells. We compared the gene expression measured in single cells to the gene expression obtained from bulk samples (100–500 cells) of homogeneous cell populations as identified by cytology and location within the tumor microenvironment. This is similar to the approach that others have used to assess the quality of single-cell RNA-seq libraries [23]. The previous 3SEQ method showed a strong correlation between gene-expression data obtained from matching fresh-frozen and FFPE material [9]. As such, we used bulk samples from FFPE as the reference point to compare with single-cell FFPE gene expression.

We reviewed FFPE tissue from a mastectomy specimen obtained within the prior year from a patient with ductal carcinoma *in situ* (DCIS), which had typically low-quality RNA (RIN 2.3, DV_200_ 30%–50%). We microdissected single-cell and bulk samples from ducts involved by DCIS and from an adjacent focus of stromal macrophages, 6 bulk tissue samples and 10 single cells of each type plus 6 no-cell controls (Figure 3; see Supplemental File 1 for validation of the dissection method). Unlike fresh-tissue single-cell RNA-seq, in which the tissue is processed without first visualizing the cells or their context within the tissue, the use of LCM allowed us to microscopically identify specific areas and cells to capture within the mastectomy specimens. This effectively allowed us to pick specific single cells of interest from the millions of cells present in the clinical archival specimen. Based on the above serial-dilution experiments with the human reference RNAs, a sample of 500 cells is well beyond enough input for high-quality data from a bulk library. This number of cells can also be obtained from a sections of a single breast duct involved by DCIS or a nearby collection of macrophages.

**Figure 3:**
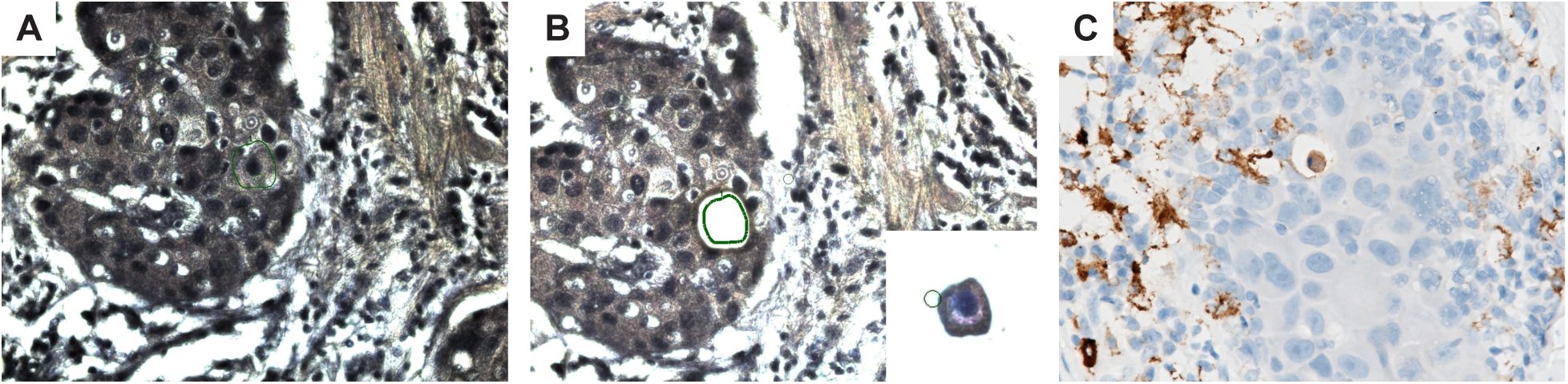
Laser-capture microdissection of ductal carcinoma *in situ*. A: Single cell within a duct involved by DCIS, targeted for dissection (green outline). B: Duct post-dissection and (inset) the captured cell on the LCM cap. C: Immunohistochemistry for macrophage marker CD163.

There are minor differences between the quality of the single-cell and bulk libraries. 27% of reads from the bulk libraries were uniquely alignable (Figure 4A); this is presumably lower than the yield from intact reference RNA because the RNA in FFPE tissue is fragmented below the desired length. The percentage of uniquely alignable reads was 17% in the single-cell libraries, likely lower because of the increased abundance of PCR primer dimers in samples with very low input. However, only 2% of reads from the negative-control libraries were uniquely alignable. As expected, we also saw a higher proportion of duplicate reads in the single-cell libraries, but the alignment loci stayed fairly consistent (Figure S27).

**Figure 4:**
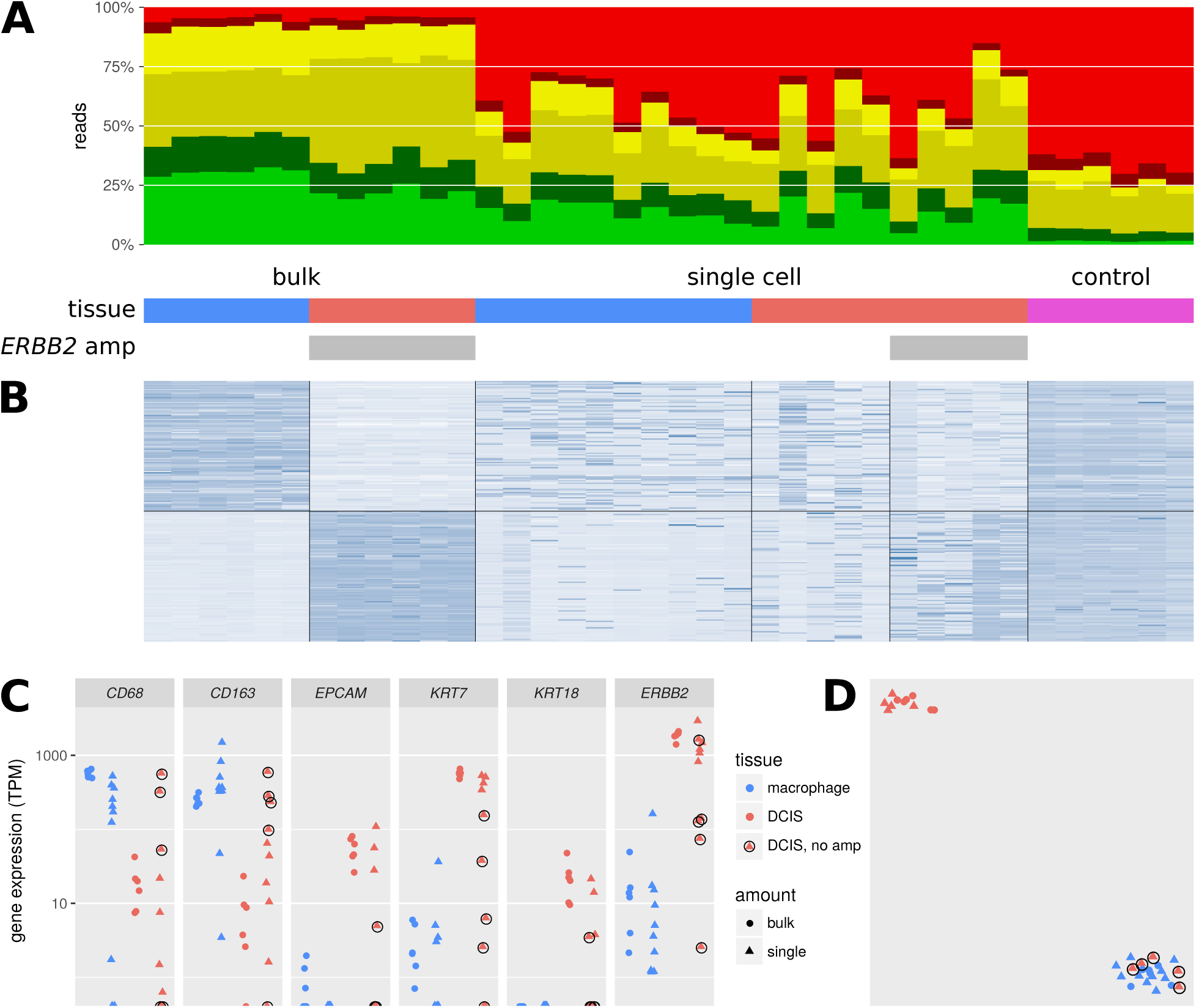
Gene-expression profiling on bulk and single-cell samples from FFPE tissue dissected by LCM. A: Alignability of Smart-3SEQ reads from FFPE bulk tissues, single cells, and negative controls. “ERBB2 amp” denotes the presence of the ERBB2 amplification in some samples (see Figure 5). B: Expression (regularized log read count, normalized by row) of the 100 genes with the greatest enrichment in bulk macrophage relative to bulk DCIS and the 100 genes with the opposite enrichment, all significant at P_adj_ < 0.01. C: Expression (transcripts per million) of known marker genes for macrophage (*CD68, CD163*) and DCIS (*EPCAM, KRT7, KRT18, ERBB2* (*HER2*)). Single cells from the DCIS tumor that lack the *ERBB2* amplification are circled. we infer that these cells are intraductal macrophages. D: t-SNE analysis of all genes; same plotting scheme as C.

To assess the degree to which Smart-3SEQ’s gene-expression profiles of dissected single cell can recapitulate the information from bulk cell populations, we first determined which genes were significantly differentially expressed (multiple testing–adjusted *p*_adj_ < 0.01) between bulk stromal macrophage and ductal tissue samples. As expected, the gene-expression profiles between ductal cells and stromal macrophages are considerably different and identify biomarkers indicative of DCIS and macrophage populations. The top 100 significantly differentially expressed genes, according to only the bulk samples, are shown as a heatmap to illustrate the differences (Figure 4B; these genes can be found by sorting and filtering Supplemental File 3). The significantly differentially expressed genes include common clinical biomarkers for macrophages (*CD68* and *CD163*) and DCIS cells (*EPCAM, KRT7, KRT18*, and *ERBB2 (HER2*)) (Figure 4C, Supplemental File 3). The single-cell samples generally have gene-expression profiles similar to the cell type–matched bulk samples. All the gene-expression patterns of single stromal macrophages match the bulk stromal macrophages. No stromal macrophages match ductal-cell signal. Remarkably, five ductal cells demonstrate a gene-expression pattern similar to the bulk stromal macrophages.

To explore whether we can use the single-cell data to identify distinct populations of cells, we performed t-SNE, a machine-learning technique that visualizes multidimensional data in two dimensions [24], on the entire dataset including negative controls (Figure 4D). t-SNE shows that the five ductal cells whose gene-expression profiles resemble those of macrophages also cluster with the stromal macrophage cells and the bulk stromal macrophage samples. The other five ductal cells cluster with the bulk ductal tissue samples while the control samples cluster distinctly from all others. These findings suggest that the bulk ductal tissue samples are heterogeneous with at least two different cell types, DCIS cells and intraductal macrophage cells, which are resolved with single-cell dissection but not bulk tissue homogenization. Immunohistochemistry for a macrophage marker on the duct used for dissection (*CD163*) confirms that there are macrophages alongside DCIS cells (Figure 3C).

To further clarify the cell status, we note that the DCIS cells from this mastectomy have an amplification of a region including the *ERBB2* (*HER2*) locus, which encodes an important therapeutic target. Examination of gene expression from the single and bulk DCIS cells demonstrates a coordinated increase in signal at a possible amplicon surrounding the *ERBB2* locus (Figure 5). This amplification appears in all of the bulk DCIS samples as well as the single ductal cells whose expression profiles match the bulk DCIS, but the amplification is absent in the bulk macrophage samples and in the single ductal cells that match the macrophage profile. We note that this is a qualitative observation to confirm a specific hypothesis; methods for rigorous unsupervised detection of genome amplifications from expression data are outside the scope of this report.

**Figure 5:**
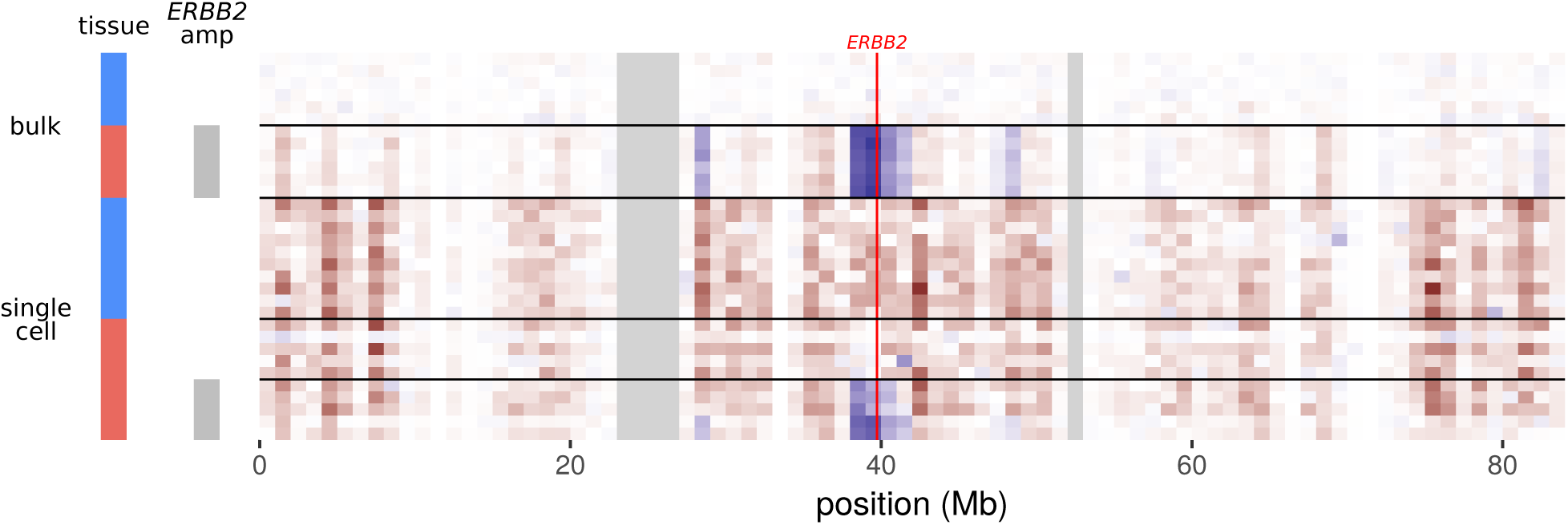
Inferred DNA copy number for Chromosome 17 in bulk and single cell samples. Genes are aggregated in blocks of 1 Mb by transcription termination site. Heatmap cells show expression normalized to the mean of the bulk macrophages. Red: lower expression than bulk macrophages; blue: higher expression than bulk macrophages; gray: no data. The bright red line shows the position of the *ERBB2* locus. Legend colors for cell types (left) match Figure 4.

There are 52 genes that were significantly differentially expressed (*p*_adj_ < 0.05) between single stromal macrophages and single ductal cells inferred to be macrophages (Supplemental File 3). Many of the most significant genes are related to extracellular matrix interactions and leukocyte signaling. These data raise the possibility that intraductal macrophages have a phenotype that is distinct from stromal macrophages, and further highlight the ability of single-cell LCM and Smart-3SEQ to dissect complex cell populations that would otherwise be missed in bulk analysis.

In a larger-scale replication of this study, we observed the same pattern of distinct single macrophages and heterogeneous single tumor cells (Figure S28). In particular, PCA (Figure S28C) showed several distinct trends. Among the bulk tissue samples, PC1 separated healthy macrophage from the tumors, as expected, while PC2 showed a continuum of the tumor samples with invasive ductal carcinoma (a more advanced tumor stage) at the extreme. On these same PCA axes, the single epithelium cells formed a new cluster intermediate between single IDC and single macrophage. However, the single DCIS cells were more heterogeneous, forming a cluster that spanned along the bulk DCIS–IDC continuum and subsumed the cluster of single IDC cells. This is consistent with the biology of the DCIS analyzed in this case: unlike the samples in the previous experiment, these samples were dissected from several different ducts in the same patient’s tissue, therefore these DCIS cells display more heterogeneity than the previous samples (Figure 4). Again, the single cells’ heterogeneity was concealed among the bulk-tissue samples, reinforcing the importance of genome-wide single-cell analysis, rather than bulk-tissue or gene-specific analysis, for understanding the tumor microenvironment.

## Discussion

Smart-3SEQ is a streamlined, sensitive, and robust method for gene-expression profiling that compares favorably with previous methods. The protocol has fewer steps and takes less time than any other RNA-seq method to date, and achieves a very low cost by using minimal volumes of common reagents (Table S2). Here we show that Smart-3SEQ quantifies transcript abundance accurately across at least five orders of magnitude of RNA input amounts, from common working concentrations down to single cells, including those with degraded RNA. This accuracy is maintained even with inexpensive sequencing conditions (Figures S30, S31); for example, at current list prices, an Illumina NextSeq run divided over a pool of 96 libraries (4M × 76 nt reads each) comes to about 25 USD per library, after about 5 USD for the library preparation. These attributes make Smart-3SEQ useful for a variety of biological research.

Combining Smart-3SEQ with laser-capture microdissection (LCM Smart-3SEQ) enables a new kind of single-cell genomics research on clinical FFPE samples. In most “single-cell” genomics experiments, a piece of tissue is homogenized and thousands of cells are used to make individual sequencing libraries, but the positional identities of the cells are lost in the context of the tissue and must be inferred from their genomic data—we propose to call this *reverse RNA-seq* (Figure 6), by loose analogy to reverse genetics. In contrast, LCM Smart-3SEQ allows for standard *forward RNA-seq* on specific single cells: the cells of interest can be identified under the microscope by markers such as morphology, location, or biomarker staining, and this prior classification is already known when the cells are profiled for gene expression. Our study is an illustration of the knowledge gained: we identified two sets of macrophages, one present in the stroma and one present within the duct associated with ductal carcinoma *in situ*. Conventional single-cell sequencing might fail to distinguish these two macrophage populations, which had only a very small number of differentially expressed genes, whereas LCM Smart-3SEQ began with the prior information that they were in different tissue compartments.

**Figure 6:**
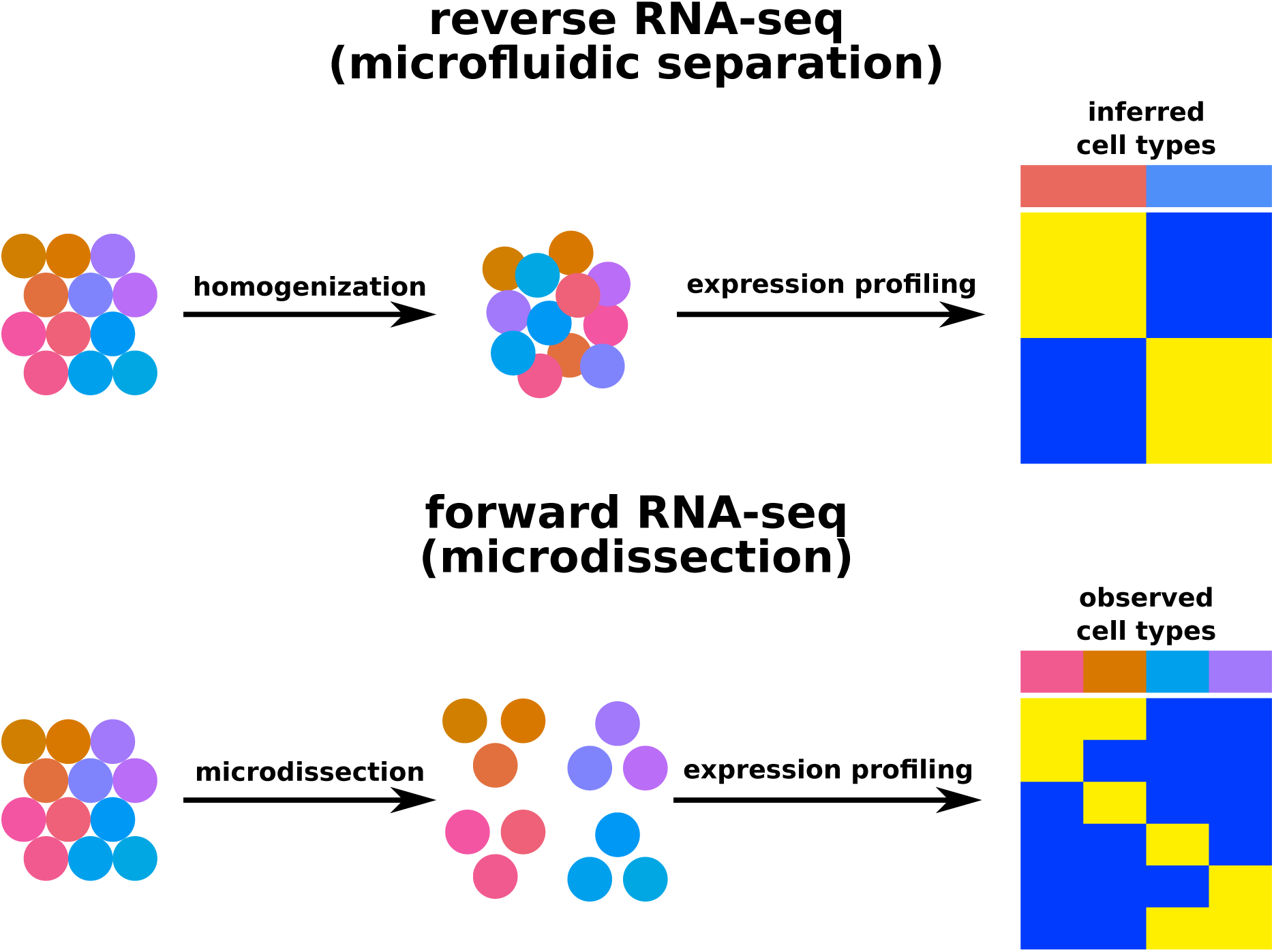
Conceptual diagram of different RNA-seq approaches. Reverse RNA-seq: Cells are disaggregated from tissue, destroying information about histological context and organization. After single-cell RNA-seq, expression profiles are used to retroactively infer categories of cells. Forward RNA-seq: Cells are dissected *in situ* according to their histology, so these *a priori* classes can inform differential gene-expression analysis.

We have demonstrated the success of our method for single-cell expression profiling on a number of levels: 1) our approach is effcient: we have obtained distinct and reproducible gene-expression profiles from our single-cell experiments; 2) the single-cell libraries recapitulate bulk gene-expression data; 3) we identified two distinct cell-type signatures, the DCIS profile and the macrophage profile, with clinically recognized biomarkers of each cell type; 4) the data are sufficiently quantitative to be used in conventional single-cell RNA-seq approaches; 5) the data are sufficiently quantitative that we can detect a DNA copy-number change; and 6) we can use the data to discover potentially new cell phenotypes, such as stromal macrophages vs. ductal macrophages, and describe gene-expression profile differences between the two cell subtypes. There are limitations to LCM Smart-3SEQ. LCM is labor-intensive and this limits the number of cells that can be collected and observed. The quality of RNA from FFPE material is poor. Recent FFPE clinical material that has been well archived can have an RNA integrity number (RIN) above 3. However, RNA quality from FFPE material decreases with time and decades-old FFPE material will yield degraded RNA that is challenging for the LCM Smart-3SEQ method. As single cells already have a limited amount of RNA, the age of the FFPE specimen may influence the cell-to-cell variation in absolute RNA expression and cell profile differences will be altered. The RNA obtained from a single cell may represent a fraction of the total possible RNA as the LCM may only capture part of the cell. For these reasons we do not propose LCM as a substitute for microfluidic single-cell methods. Rather, they are complementary approaches that solve different problems: microfluidic RNA-seq remains the best way to characterize thousands of cells from a single tissue sample and discover clusters of cell types, while LCM Smart-3SEQ allows the experimenter to choose the specific cells or tissue regions of interest, from many different tissue samples in a single experiment, including FFPE tissue. Likewise, LCM Smart-3SEQ is not a substitute for truly *in situ* hybridization or sequencing methods [25, 26, 27, 28], which completely preserve spatial information about the tissue sample but do not perform gene-expression profiling for the entire transcriptome of every cell. However, it is more sensitive to small and low-quality input material than previous methods based on microdissection, such as tomo-seq, which creates sequencing libraries from whole sections of tissue [29], and Geo-seq, which uses LCM but requires fresh-frozen tissue [30].

Our approach addresses a number of experimental-design problems that are difficult to overcome with conventional single-cell RNA-seq. FFPE archival clinical tissue cannot be examined by conventional single-cell RNA-seq as the cells cannot be physically dissociated. Non-archival tissue samples, both fresh and frozen, are difficult to collect for clinical studies, especially those that require large numbers of samples for assessing clinical outcomes. LCM Smart-3SEQ also enables studies on uncommon or microscopic lesions that are difficult to collect as fresh or fresh-frozen material. Thus, a much wider variety of cells and diseases can be assessed using archival material with Smart-3SEQ. There are also technical advantages of LCM Smart-3SEQ due to the ability to bank isolated single cells for future experiments. This includes the spiking in of known cells to control for batch effects with biological replicates.

The LCM Smart-3SEQ method creates a new histologically focused approach to studying small cell populations and individual cells. Cells of interest can be chosen by morphology, microenvironment location, or *in situ* biomarker status and then profiled to uncover previously unappreciated heterogeneity in gene expression, including subtle changes that might escape reverse RNA-seq analyses. This approach will allow transcriptome profiling to isolate and measure each cell individually, moving histology and molecular pathology to the level of cell-to-cell variation.

## Materials and methods

### Reference RNA preparation

ERCC standards and Human Brain Reference RNA were purchased from Thermo Fisher Scientific (catalog #4456739 and #AM6050 respectively). Universal Human Reference RNA was purchased from Agilent Technologies (#740000) and resuspended in RNA Storage Solution (Thermo Fisher Scientific #AM7001), which was also used for all RNA dilutions. HBRR and UHRR stocks were measured with a Quant-iT RNA Assay Kit (Thermo Fisher Scientific #Q10213) on a Qubit fluorometer and these concentrations were used to calculate dilutions. All RNA stocks were stored at −80 °C.

### Tissue preparation

Samples were collected with the approval of a HIPAA-compliant Stanford University Medical Center institutional review board. The FFPE tissue blocks were archived with the Stanford University Hospital Department of Pathology. RNA quality for the FFPE block was measured by extracting total RNA from a separate section with the RNeasy FFPE Kit (Qiagen #73504) and testing it on an Agilent Bioanalyzer with the RNA Pico kit.

### Slide preparation

Consecutive sections of the FFPE block were taken on a microtome at 7 μm thickness and mounted on glass slides with polyethylene naphthalate membranes (Thermo Fisher Scientific #LCM0522). Slides were stored overnight in a nitrogen chamber. The next day, slides were immersed 20 s each in xylenes (3 times), 100% ethanol (3 times), 95% ethanol (2 times), 70% ethanol (2 times), water, hematoxylin (Dako #S3309), water, bluing reagent (Thermo Fisher Scientific #7301), water, 70% ethanol (2 times), 95% ethanol (2 times), 100% ethanol (3 times), xylenes (3 times).

### Laser-capture microdissection

Slides were dissected immediately after staining. Histology was categorized by a board-certified pathologist (RBW) according to initial sections stained with hematoxylin and eosin. Cells were dissected on an Arcturus XT LCM System using both the ultraviolet (UV) laser to cut out each sample and the infrared laser to adhere it to a CapSure HS LCM Cap (Thermo Fisher Scientific #LCM0215). For bulk samples, roughly 500 cells were captured by area, according to density estimates by cell counting on small areas. For single cells, a cell was dissected from the same area as the corresponding bulk sample, then any additional cells adhering to the cap were ablated with the UV laser. For the ablation validation experiment, two regions of both types (still roughly 500 cells each) were captured on the same cap, and then one region or the other was ablated with the UV laser, except the no-ablation controls (therefore they had roughly 1000 cells). After LCM, the cap was sealed in an 0.5 mL tube (Thermo Fisher Scientific #N8010611) and stored at −80 °C until library preparation, which was performed within 3 days of dissection.

### Microscopy image preparation

LCM photographs were captured in JPEG format with the built-in camera and software of the Arcturus XT system. Photographs in Figures 3 and S29 were color-corrected to enhance contrast with the automatic “Stretch HSV” function in GNU Image Manipulation Program 2.8.20. All photographs were recompressed to reduce file size with the “jpeg-recompress” feature in JPEG Archive 2.1.1, using “veryhigh” quality, “accurate” mode, and the mean pixel error algorithm. All photographs are presented at the original resolution. A complete archive of microscopy photos from all dissections is included in Supplemental File 4.

### Library preparation

Sequencing libraries were prepared according to the Smart-3SEQ protocol (Supplemental File 2). Reference RNA libraries were prepared using the standard protocol for non-degraded RNA and the pre-SPRI pooling option, one batch for the ERCC experiment and one for the human reference RNAs, with the numbers of PCR conditions shown in Figures S4 and S9. The fresh-frozen vs. FFPE experiment used the corresponding versions of the protocol and only libraries of the same version were pooled; no-template controls were prepared with both versions. The fresh-frozen vs. FFPE experiment used the same numbers of PCR cycles for the same dilutions as the reference-RNA experiment. The LCM experiments used the special protocol for FFPE tissue on an Arcturus LCM HS cap and the pre-SPRI pooling option, a single batch for all LCM samples, using 22 PCR cycles for the bulk and ablated samples with roughly 500 cells, 25 cycles for the single cells, and 20 cycles for the no-ablation controls with roughly 1000 cells. No-template controls were prepared identically to single-cell samples in the same batch using empty LCM caps. Libraries were characterized immediately and stored at −20 °C until sequencing.

### Library characterization and sequencing

Libraries were profiled for size distribution on an Agilent 2200 TapeStation with High Sensitivity D1000 reagent kits and quantified by qPCR with a dual-labeled probe [31], and libraries were mixed to equimolarity according to the qPCR measurements. In the reference-RNA experiments, libraries were prepared according to the manufacturer’s instructions with a 1% spike-in of the *ϕ*X174 control library (Illumina #FC-110-3002) and sequenced on an Illumina NextSeq 500 instrument with a High Output v2 reagent kit (Illumina #FC-404-2005), reading 86 nt for read 1 and 6 nt for the P7 index read. The LCM experiments used 76 nt for read 1. In the experiment for testing the effect of read lengths, libraries were prepared according to the manufacturer’s instructions with a 1% *ϕ*X174 spike-in and sequenced on an Illumina MiSeq instrument with a Reagent Kit v3 (Illumina #MS-102-3001), reading 169 nt for read 1 and 6 nt for the P7 index read.

### Data preprocessing

In most experiments, base calls from the NextSeq were demultiplexed and converted to FASTQ format with bcl2fastq (Illumina Inc); adapter trimming was enabled and short trimmed sequences were retained for diagnostic purposes. For the MiSeq run, FASTQ generation, demultiplexing, and adapter trimming were performed by the MiSeq Reporter in “Generate FASTQ” mode. Only reads that passed the chastity filter were included in the total read count and further analysis. The first 5 nt of each read were removed from its sequence and appended to the read name as its UMI; the following 3 nt were assumed to derive from the G-overhang of the template-switch oligonucleotide and were discarded. Reads shorter than 8 nt after adapter trimming were assumed to be primer dimers and were excluded from other analysis, but still counted toward the total number of sequenced reads. Software versions and command-line arguments are listed in Table S3.

### Read alignment and counting

Reads from the experiment that used only ERCC RNAs were aligned to NIST’s empirical ERCC sequences [32] by NovoAlign (Novocraft Technologies Sdn Bhd), using A30 as the “adapter” sequence, and alignments with MAPQ < 10 (posterior probability < 0.9) were discarded. Reads from the experiment that mixed human reference RNAs with ERCC spike-ins were aligned by STAR [33] to a combination of the ERCC sequences and the hg38 reference genome sequence, masked for SNPs according to dbSNP version 150 (obtained from the UCSC Genome Browser) [34], and the GENCODE version 27 gene annotations [35] were provided as the reference transcriptome. Reads from the experiments with human tissue were aligned in the same way except their reference index contained no ERCC sequences. Alignments per human transcript were counted by featureCounts from the Subread suite [36], including only non-duplicate alignments in the correct (sense) orientation. In the replication of the FFPE LCM experiment, we discarded 19 failed libraries with fewer than 10,000 reads aligned to genes, including all 6 of the no-template controls. When a read aligned to more than one annotated gene, the category was decided by order of priority, with the most likely categories first: a read was considered “3′ end” if it aligned to the end of any annotated transcript, regardless of where it aligned on any other transcripts, or assigned to “exon” only if it did not align to any ends, or to “intron” only if it did not align to any exons, etc.

### Detection of duplicate reads

Smart-3SEQ reads only a small number of possible fragments from each transcript, so it is likely that some of them will break at the same base position by chance, and traditional deduplication according to genome position will incorrectly mark these as duplicates. The incorporation of UMIs before PCR allows distinguishing coincidental fragmentation duplicates from true PCR duplicates among reads that align to the same position. However, as reported previously [37], the simplistic approach of allowing only one hit per UMI per position also fails when the read density is high and the number of possible UMIs is low, such that even duplicates with the same genome position and same UMI may occur by chance.

We solved this problem with an extension of the algorithm proposed by Hatsugai et al. [38]. All reads that start at the same genome position are considered potential PCR duplicates. Among that set of reads, across the vector of read counts per UMI, the frequencies of observed counts are tallied. Then the number of non-duplicate reads is estimated as the weighted average of all counts’ predictions, where each observed count c predicts that all counts greater than *c* +1 per UMI are PCR duplicates, and the weight is the number of UMIs with that count. That is, the non-duplicate read count is estimated as

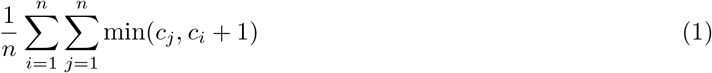

where *c_i_* is the number of reads with UMI *i* out of *n* possible UMIs (*n* = 1024 here as the UMIs are 5 nt long). Zero counts are included, so each UMI with zero hits votes for a limit of 1 non-duplicate read per UMI. Therefore when the total number of reads aligned to the genome position is small, this formula produces the same results as the traditional approach of allowing only one hit per UMI. However, unlike the traditional approach, it does not underestimate the non-duplicate read count when the total is large.

### Quantification of transcript abundance

Transcripts per million were computed differently between Smart-3SEQ and whole-transcript RNA-seq because the latter must be normalized by transcript length:

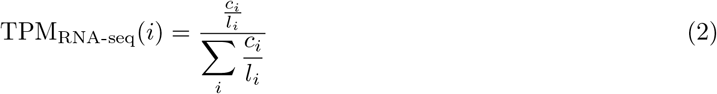

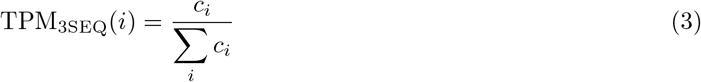

where *c_i_* is the number of reads aligned to transcript *i* and *l_i_* is the length of the annotated transcript.

Because transcript abundances and Smart-3SEQ read counts by transcript were log-distributed, but some counts were zero, we defined the linear correlation between read counts and other abundance measurements as

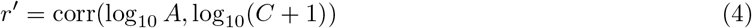

where *A* and *C* are the random variables of which the abundance measure *a* and read count *c*, respectively, are particular realizations, and corr(*X, Y*) is Pearson’s product-moment correlation.

### Analysis of gene expression

Differential gene expression was analyzed with DESeq2 1.6.1 [39]. All default options were used except dispersions were estimated with local fitting. The regularized log transformation was used to normalize read counts. PCA was performed on the normalized data with the prcomp function and t-SNE was performed with the Rtsne package [40] on the Euclidean distance matrix of the normalized data.

### Data access

The sequencing data generated in this study have been submitted to the NCBI BioProject database (https://www.ncbi.nlm.nih.gov/bioproject) under accession number PRJNA413176. Scripts used to perform the analyses in this study are collected in Supplemental Code.

## Supporting information

Supplemental File 2: Smart-3SEQ protocol

Supplemental File 3: Gene-expression table

Supplemental File 4: LCM photos

Supplemental File 5: Cost calculations

## Author contributions

JWF conceived the Smart-3SEQ method. JWF, PJ, CZ and SXZ developed the protocol. JWF and CZ performed the experiments shown here. JWF developed the data-processing pipeline. JWF and PL analyzed the data. JWF and RBW prepared the manuscript. MJM supervised JWF and PJ in the initial Smart-3SEQ phase of the project. RBW supervised JWF, CZ, SXZ, and PL in the later LCM phase of the project.

## Acknowledgments

We are grateful to the laboratory of Carlos D. Bustamante and especially to Shirley Sutton and Alexandra Sockell for use of the TapeStation and NextSeq instruments. We are also grateful to the McGill Group for Suicide Studies and especially to Gary G. Chen for use of the MiSeq instrument and guidance with sequencing. We thank Takara Bio USA for providing reference data on the performance of the SMART-Seq v4 kit. Some of the computations were performed on the Sherlock cluster of the Stanford Research Computing Center and the Guillimin cluster of McGill University’s High Performance Computing Centre. JWF was supported by a Foundation MIRI grant from Brain Canada. JWF and PJ were supported by the Ludmer Centre for Neuroinformatics and Mental Health. JWF, CZ, SXZ, PL, and RBW were supported by the National Cancer Institute of the National Institutes of Health under award number R01CA183904.

## Competing interests

The authors declare no competing financial interests.

