## Supplemental File 1: Supplemental material for "Gene-expression profiling of single cells from archival tissue with laser-capture microdissection and Smart-3SEQ"

### Detailed explanation of the Smart-3SEQ method

First, the input RNA is fragmented to a desirable size by a divalent cation in high heat. Whole cells or tissue can be lysed with the addition of a detergent in the same step; lysing fixed tissue may require adding proteinase K at this step and an inhibitor afterward. There is no need to purify the RNA after fragmentation as it performed in the presence of some components of the next reaction mix, nor is mRNA enrichment or rRNA depletion required. Instead, immediately after RNA fragmentation, reverse transcription is primed with an oligonucleotide comprising a $3'$ anchored oligo(dT) primer, which hybridizes to the beginning of the RNA template's poly(A) tail (so only the single fragment that contains this site is reverse-transcribed), and a non-complementary $5'$ sequence matching the innermost portion of the downstream sequencing adapter (P7 on the Illumina platform). This incorporates the partial adapter into the first cDNA strand and eliminates the need to add that adapter by ligation later.

After extending the first cDNA strand, MMLV-derived reverse transcriptase tends to extend several non-template bases at the $3'$ end, which are primarily dC. This provides a target for hybridization with a second oligonucleotide, which comprises a short $3'$ oligo(G) primer and the innermost portion of the upstream sequencing adapter (Illumina P5). Reverse transcriptase then performs a "template switch", extending a second cDNA strand from this new primer. Thus the reverse transcription produces both the first and second cDNA strands in a single incubation, and this ds-cDNA has partial sequencing adapters at both ends. Note that the template-switch oligonucleotide consists mainly of DNA but the $3'$ guanine residues are RNA to reduce off-target strand invasion. Each oligonucleotide in the template-switching reverse transcription also includes a blocking group (biotin) at its $5'$ end to discourage concatenation of additional adapters.

All that remains to produce a sequencing-ready library is to extend the adapters to full length, including the multiplexing barcodes, and to amplify the library to sufficient concentration for quality control and pooling. Both purposes are served by PCR with primers matching the sequences of the entire adapters, which anneal to the partial adapters on the cDNA and extend them to full length. Finally, the only cleanup step in the protocol is to purify the amplified dsDNA library by a single SPRI procedure, using stringent conditions to avoid retaining molecules that are too short to be useful. Optionally, because they are now labeled with separate barcodes, the amplified libraries can be pooled before cleanup to combine them into a single small volume, reducing the amount of downstream handling and yielding acceptable concentrations from lower numbers of PCR cycles.

When the library is sequenced, each read contains up to five sections (Figure S3A): 1) the UMI, a set of random bases included in the second-strand primer to discriminate PCR duplicates from fragmentation duplicates; 2) a short stretch of Gs; 3) cDNA sequence matching the source transcript; 4) a long stretch of As, if the read length is longer than the cDNA insert; and 5) potentially the downstream adapter sequence, if the read length is much longer than the cDNA insert, though in practice bases downstream of a homopolymer tend to be poorly read. When the cDNA sequences are aligned to the reference transcriptome, they align in the sense orientation slightly upstream of the polyadenylation site (Figure S3B) and the read count is directly proportional to the abundance of the source transcript, regardless of the transcript's length.

During the development of the protocol, we extensively tested RNA fragmentation conditions and settled on a relatively gentle protocol (5 min at 80 °C with concentrated magnesium; typically this is done at 94 °C, often for a longer duration), which retains fairly large fragments from intact RNA that are easy to discriminate from adapter dimers in size selection. In our further optimizations for FFPE RNA there was no improvement, compared with these already gentle conditions, from further reducing the fragmentation or eliminating it altogether by rearranging the order of reagent addition. FFPE RNA always yields short fragments so the FFPE version of the protocol has a less stringent size selection, and that is the only difference between FFPE and frozen samples in the protocol for isolated RNA.

There is, however, a substantial difference in the modified protocols for working directly on LCM tissue. We found it necessary to incubate FFPE tissue at 60 °C for a full hour, in the presence of proteinase K, to fully lyse the tissue. To avoid further fragmenting the RNA during this incubation, we changed the compositions of the reaction mixes to remove any magnesium from the initial lysis mix, and in that condition we did see a large improvement in the quality of the libraries.

### Validation of the laser ablation method

Dissecting single cells with LCM is difficult and sometimes more than one cell is recovered on the cap. We ensured that our libraries came from true single cells by destroying the extraneous cells with the UV laser (Figures S29A, S29B). To verify that this eliminates the signal from the ablated cells, we performed an experiment on a larger scale. On each cap we collected a roughly equal number of macrophages and DCIS cells, then on some caps we ablated all cells of one type, while on other caps we performed no ablation as a control (3 bulk ablations for each tissue type plus 2 no-ablation controls). Compared with the bulk samples (6 per tissue type), the gene-expression data from the no-ablation controls resembled a mix of both cell types, while the profiles from the ablated samples resembled the bulk data from their unablated cell types (Figures S29C, S29D).

### Sections in the supplemental table of gene expression analysis

The "sheets" in the Excel spreadsheet of gene expression data from the FFPE LCM experiment contain the output of read counting, normalization, and DESeq2 analyses of differential gene expression.

- `count`: number of reads uniquely aligned to each gene
- `TPM`: read counts normalized to transcripts per million
- `rlog`: read counts normalized to regularized logarithm by DESeq2
- `bulk comparison`: bulk DCIS vs. bulk macrophage
- `single comparison`: single DCIS vs. single macrophage, excluding DCIS cells that lack the *ERBB2* amplification
- `amp comparison`: single DCIS with the amplification vs. single DCIS without the amplification
- `site comparison`: single DCIS without the amplification (inferred ductal macrophage) vs. single macrophage (stromal)
- `single vs. bulk`: single cells vs. bulk tissue, controlling for tissue type, excluding DCIS cells without the amplification
- `DCIS vs. macrophage`: DCIS vs. macrophage, single cells and bulk combined and controlled for, excluding DCIS cells without the amplification

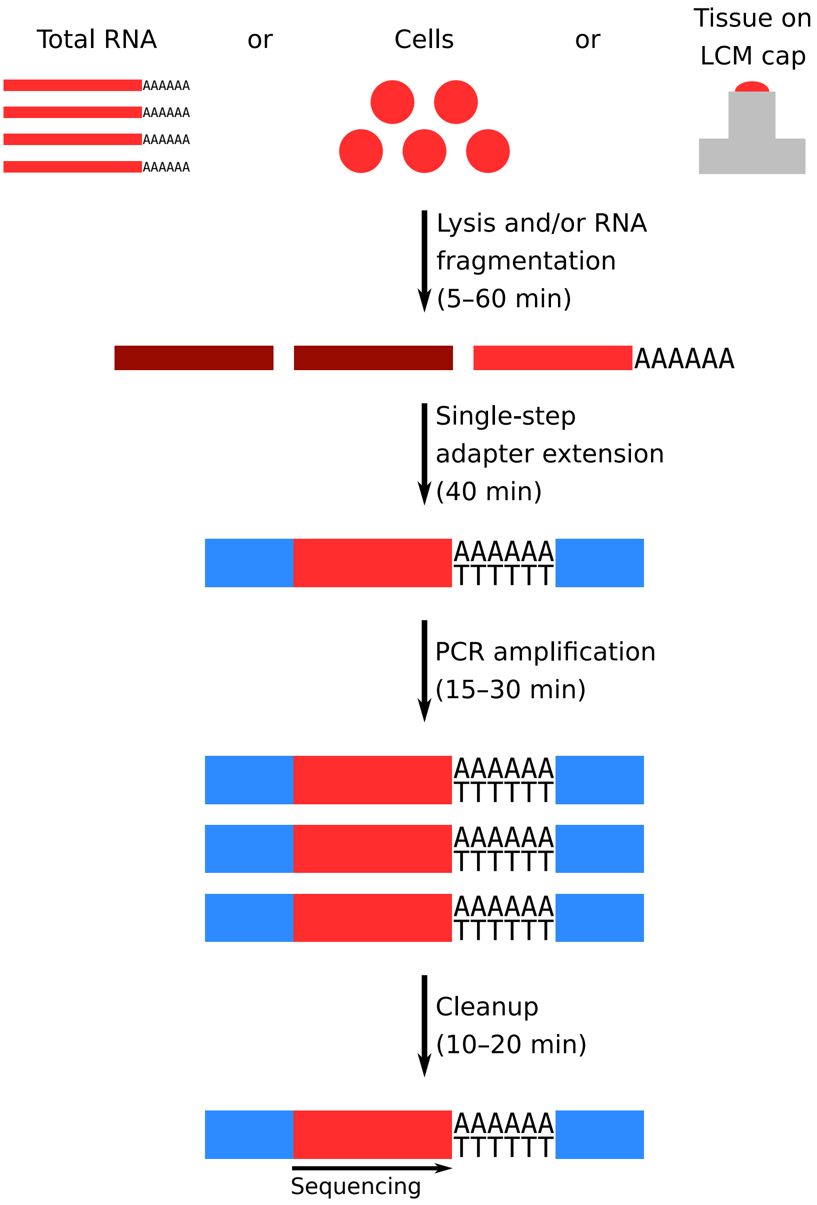


Figure S1: Schematic of the Smart-3SEQ workflow.

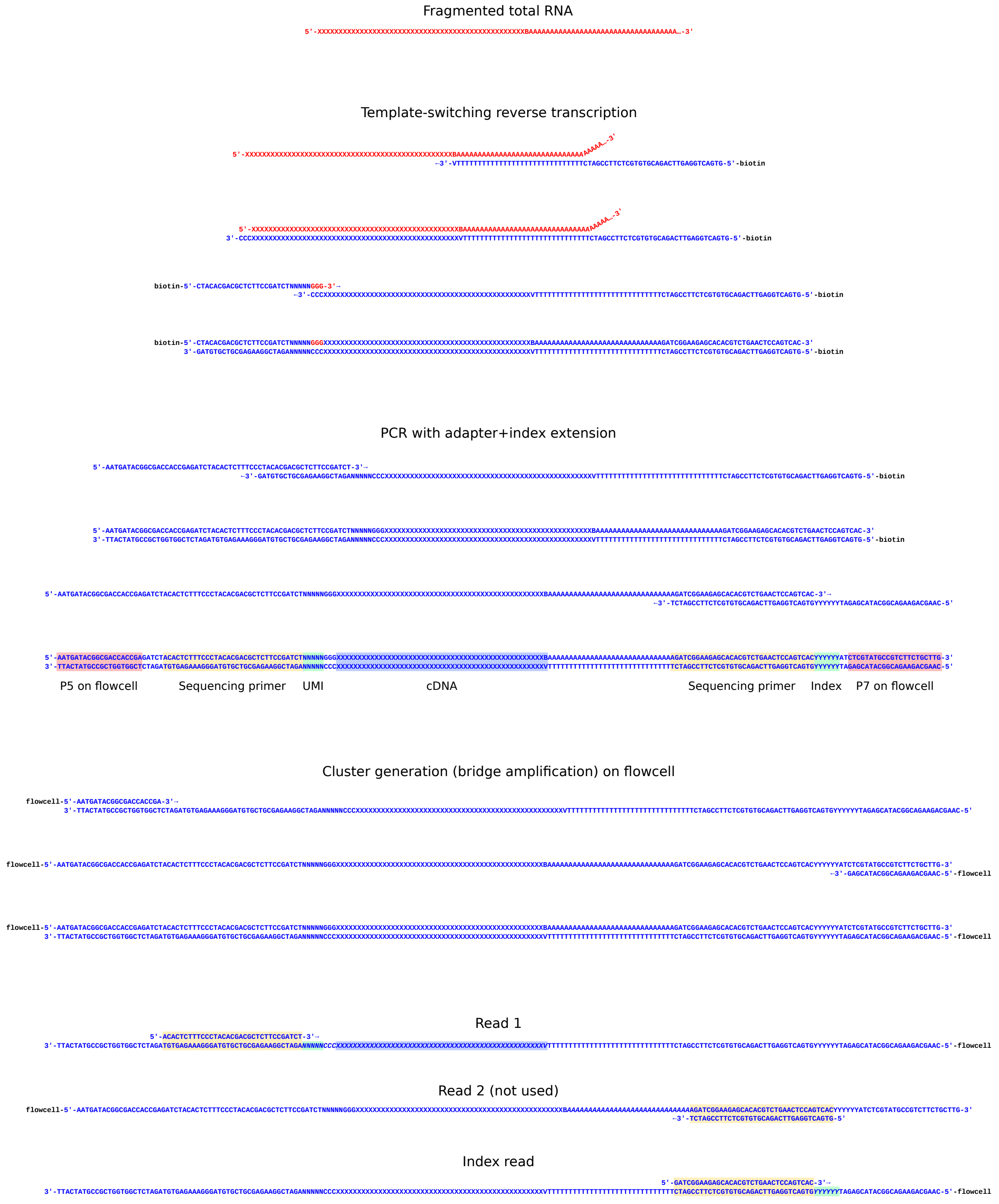


Figure S2: Schematic of oligonucleotide sequences used in Smart-3SEQ library preparation and sequencing on the Illumina platform with single indexing. Oligonucleotide sequences © 2017 Illumina, Inc. All rights reserved. Derivative works created by Illumina customers are authorized for use with Illumina instruments and products only. All other uses are strictly prohibited.

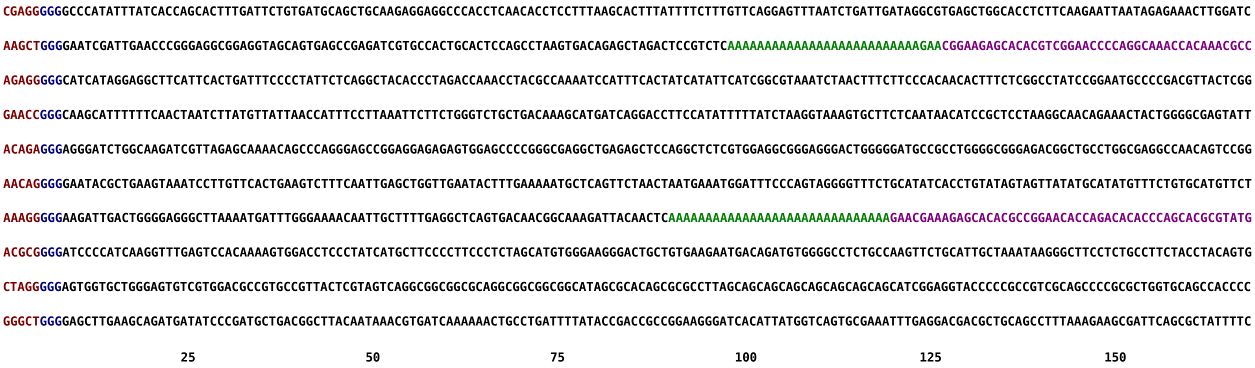


(A)

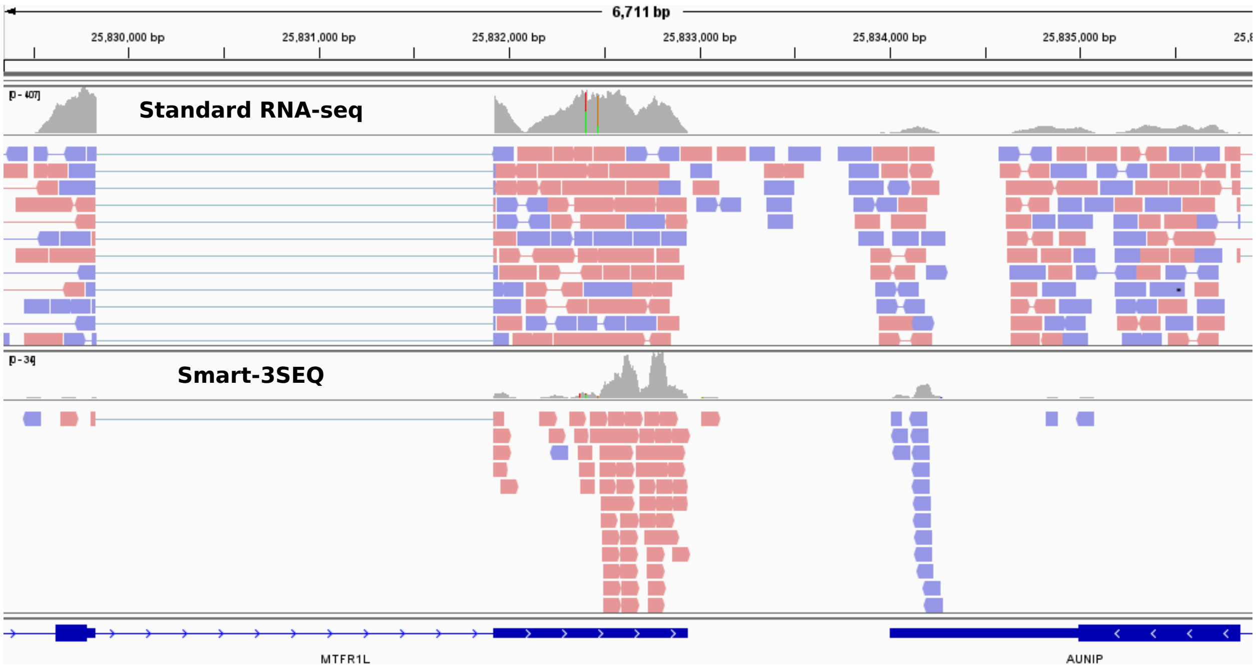


(B)

Figure S3: Example data from Smart-3SEQ. A: Read sequences. Before the cDNA sequence, each read begins with a 5 nt UMI (red), which is tracked for deduplication, then a $G_3$ stretch (blue) derived from the overhang required for template-switching, which is discarded. The length of the remaining cDNA insert (black) depends on the integrity of the input RNA: damaged RNA will yield shorter fragments. If the reads are longer than the inserts, they may continue into the poly(A) sequence (green), whose length is expected to match that of the oligo(dT) reverse-transcription primer, 30 nt. Any further base calls (purple) are unreliable because of the difficulty of sequencing through the homopolymer. B: IGV screenshot of example alignments, alongside RNA-seq data (top) from the same sample. RNA-seq reads span all exons, while Smart-3SEQ yields sense-oriented reads that only align directly upstream of the transcription termination site. They may still straddle an exon–exon junction if the final exon is short.

Table S1: Smart-3SEQ workflow compared with selected RNA-seq methods. Analogous steps are aligned in the same row. For brevity, tagmentation with a third-party kit is shown for Takara SMART-Seq v4 rather than sonication and the manufacturer's ligation-based library preparation. RT: reverse transcription. TS-RT: template-switching reverse transcription.

| TruSeq Stranded Total RNA (Illumina) | SMART-Seq v4 (Takara) | SMARTer Pico (Takara) | 3SEQ (Beck et al. 2010) | Smart-3SEQ |
|---|---|---|---|---|
| RNA isolation | RNA isolation or cell lysis | RNA isolation | RNA isolation | RNA isolation or cell/tissue lysis |
| Ribosomal RNA depletion & cleanup | | | Poly(A) selection & cleanup | |
| | | | mRNA QC | |
| RNA fragmentation | | RNA fragmentation | RNA fragmentation | RNA fragmentation |
| First-strand RT, random-primed | TS-RT, oligo(dT)-primed | TS-RT, random-primed (adds adapters) | First-strand RT, oligo(dT)-primed (adds adapter) | TS-RT, oligo(dT)-primed (adds adapters and UMI) |
| Second-strand synthesis | | | Second-strand synthesis | |
| | cDNA LD-PCR | | | |
| Cleanup | Cleanup | | Cleanup | |
| | cDNA QC | | | |
| A-tailing | | | A-tailing | |
| | | | Cleanup | |
| Dual adapter ligation | cDNA tagmentation | | Second adapter ligation | |
| Cleanup & size-selection | | | Cleanup & size-selection | |
| Library PCR | Library PCR | Library PCR | Library PCR | Library PCR |
| Cleanup & size-selection | Cleanup & size-selection | Cleanup & size-selection | Cleanup & size-selection | Cleanup & size-selection |
| | | Ribosomal cDNA depletion | | |
| | | Library PCR 2 | | |
| | | Cleanup & size-selection | | |

Table S2: Comparison of Smart-3SEQ with selected RNA-seq methods. Cost per library includes all reagents (kits, SPRI beads, enzymes) but not consumables (tubes, pipet tips) and is rounded to the nearest 5 USD.

| | TruSeq Stranded Total RNA (Illumina) | SMART-Seq v4 (Takara) | Smart-seq2 (Picelli et al. 2014) | SMARTer Pico (Takara) | 3SEQ (Beck et al. 2010) | Smart-3SEQ |
|---|---|---|---|---|---|---|
| Cost per library (USD) | $165 | $115 | $60 | $45 | $120 | $5 |
| Required total RNA | 100 ng | 10 pg | 10 pg | 250 pg | 10 µg | 10 pg |
| Protocol time | 2 working days | 2.5 working days | 2.5 working days | 5 hours | 2 working days | 3 hours |
| Strand-specific? | Yes | No | No | Yes | Yes | Yes |
| Supports damaged RNA? | Yes | No | No | Yes | Yes | Yes |
| Works directly on cells? | No | Yes | Yes | No | No | Yes |
| Works directly on FFPE tissue? | No | No | No | No | No | Yes |

Table S3: Software and configurations used.

| Program (suite) | Version | Important arguments |
|---|---|---|
| bcl2fastq | 2.17.1.1 | --minimum-trimmed-read-length 0 --mask-short-adapter-reads 0 |
| NovoAlign | 3.08.00 | -H --trim3HP -a AAAAAAAAAAAAAAAAAAAAAAAAAAAAAA |
| STAR | 2.5.3a | --outFilterMultimapNmax 1 --outFilterMismatchNmax 999 --clip3pAdapterMMp 0.2 --clip3pAdapterSeq AAAAAAAAAAAAAAAAAAAAAAAAAAAAAA |
| featureCounts (subread) | 1.5.0-p2 | -s 1 --read2pos 5 |

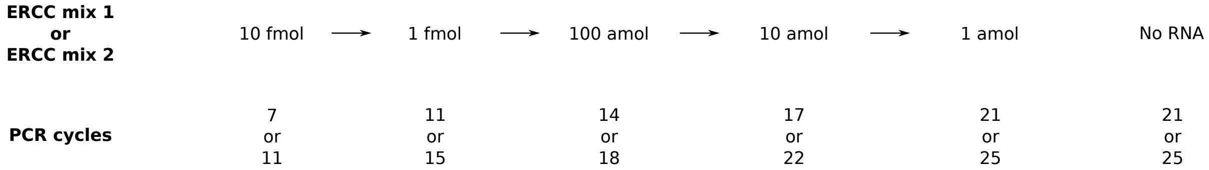


Figure S4: Experimental design of validation with ERCC standards.

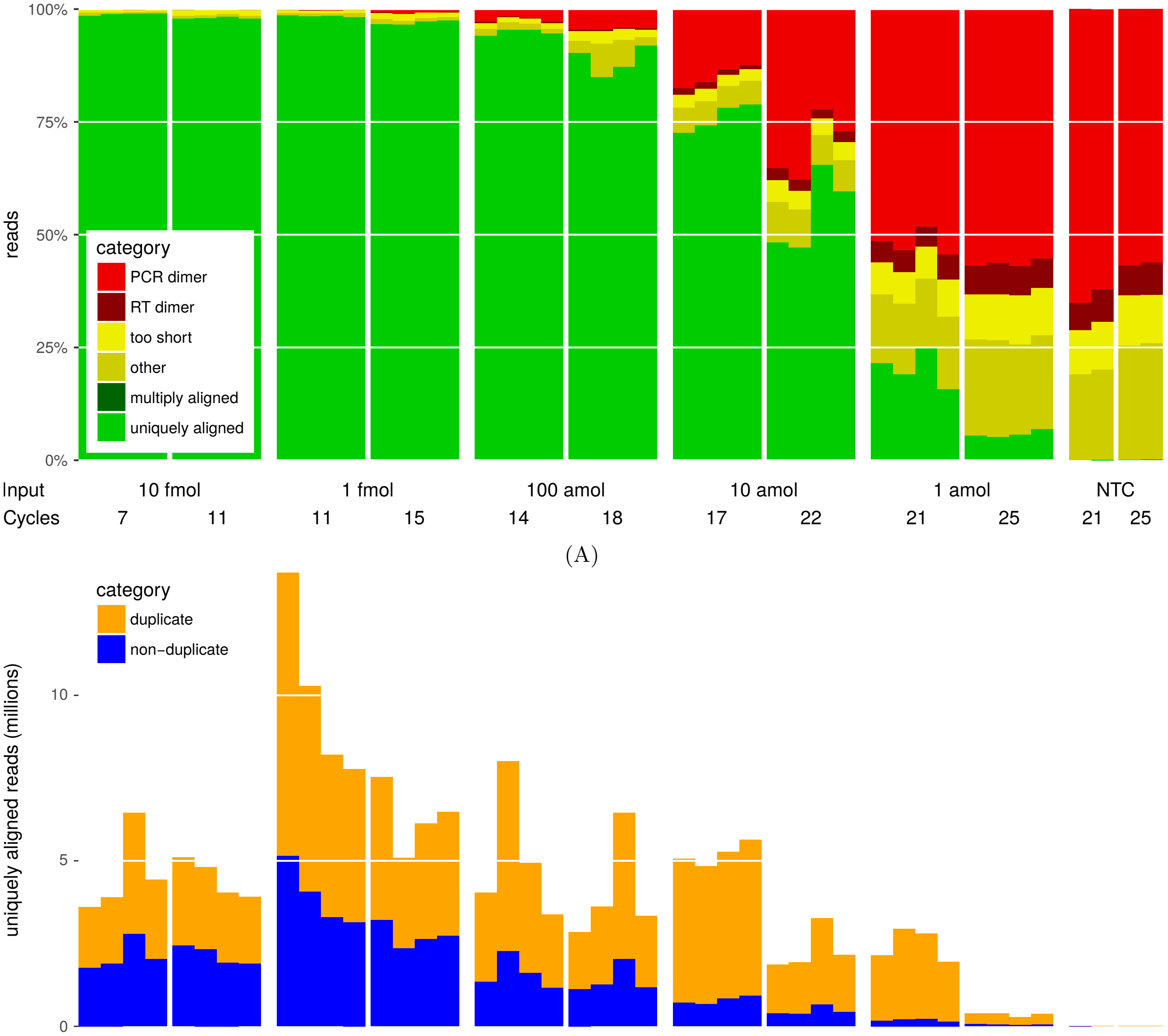


(B)

Figure S5: Quality statistics of Smart-3SEQ reads from ERCC dilutions and no-template controls. A: Read alignability. B: PCR duplication, as measured by UMIs.

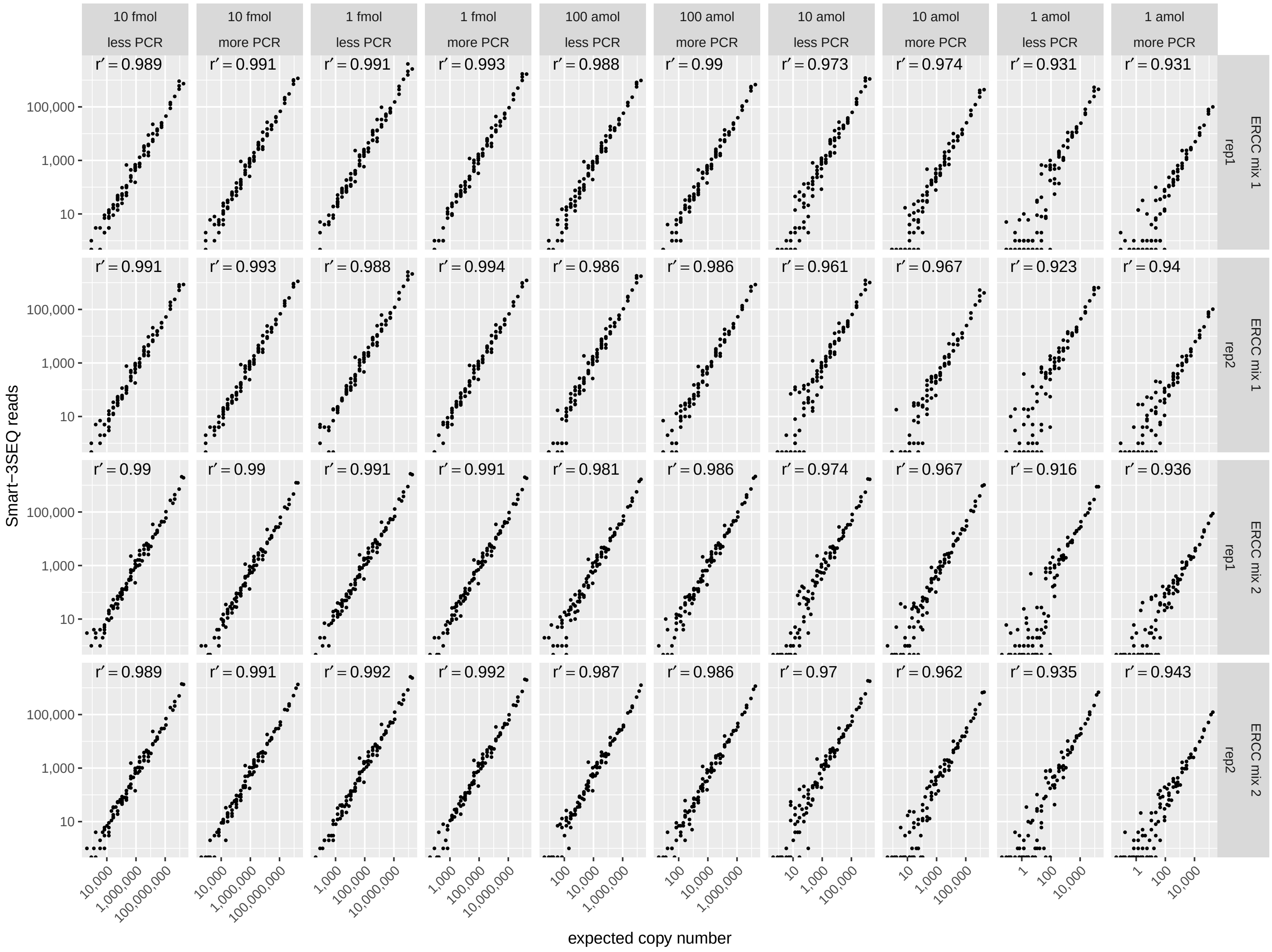


Figure S6: Accuracy of Smart-3SEQ. Smart-3SEQ standard curves from ERCC experiment. Each point is a single transcript.

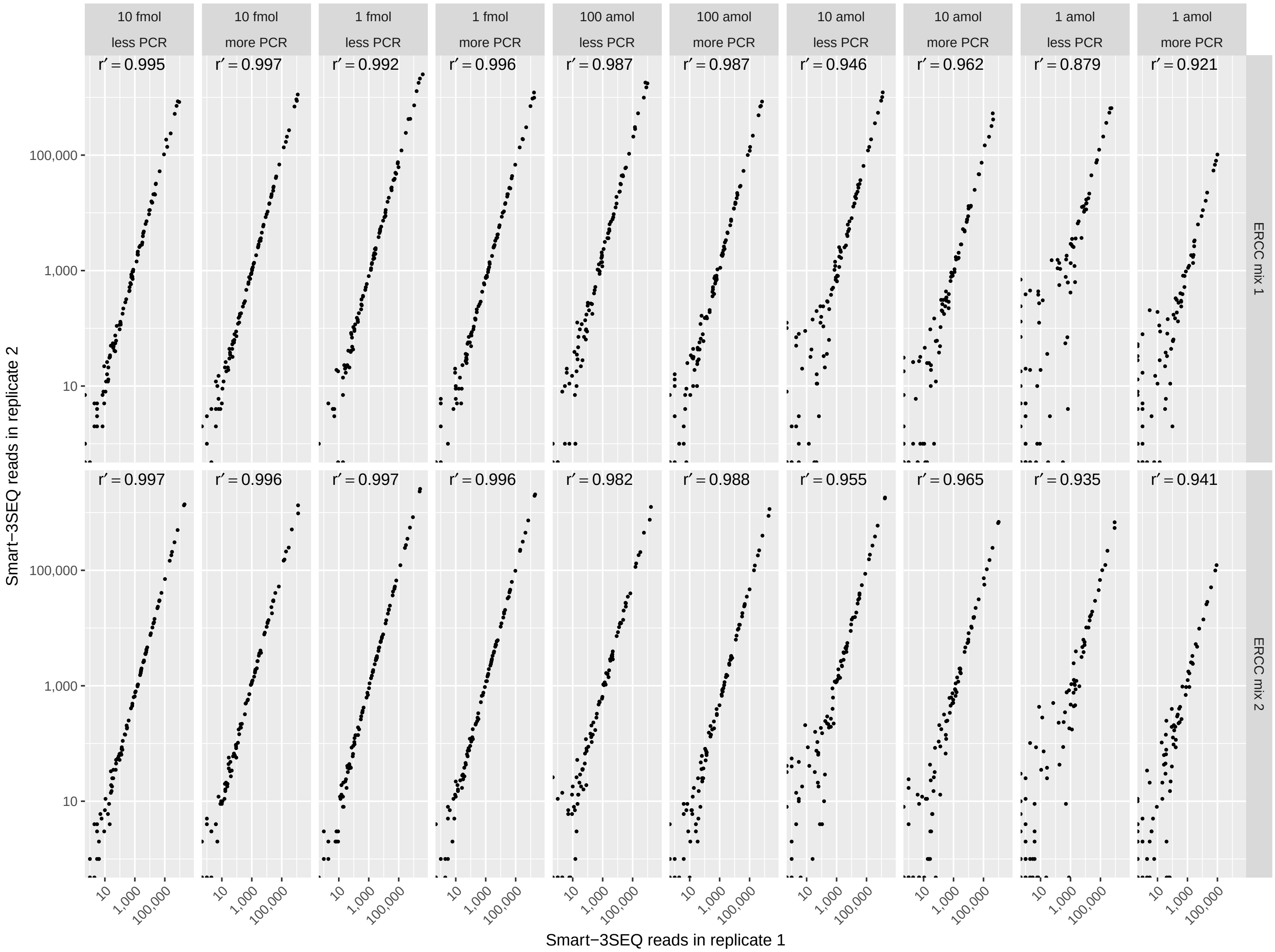


Figure S7: Precision of Smart-3SEQ. Correlation of technical replicates in ERCC experiment.

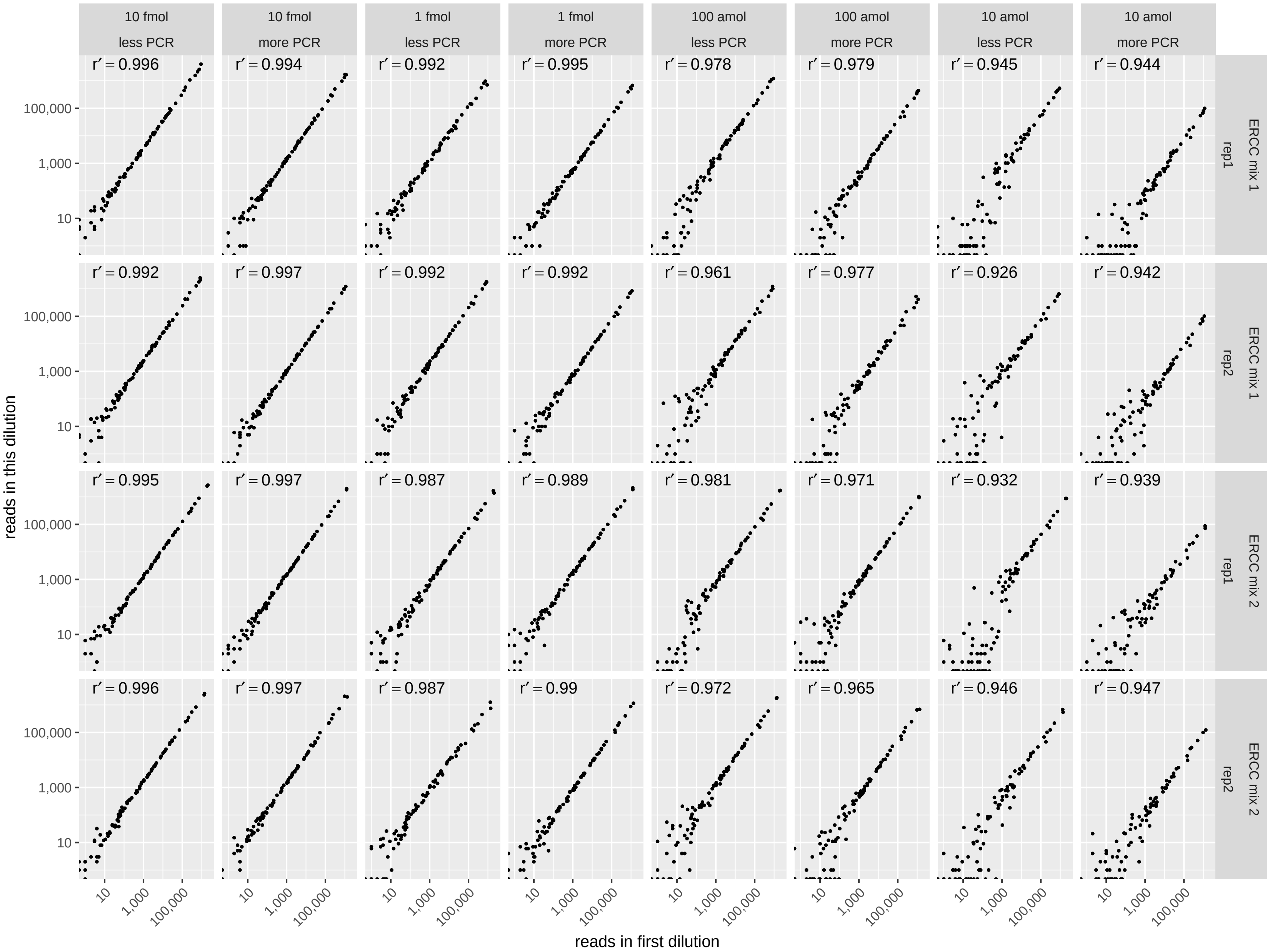


Figure S8: Sensitivity of Smart-3SEQ. Correlation of read counts in each subsequent dilution with those in the first dilution.

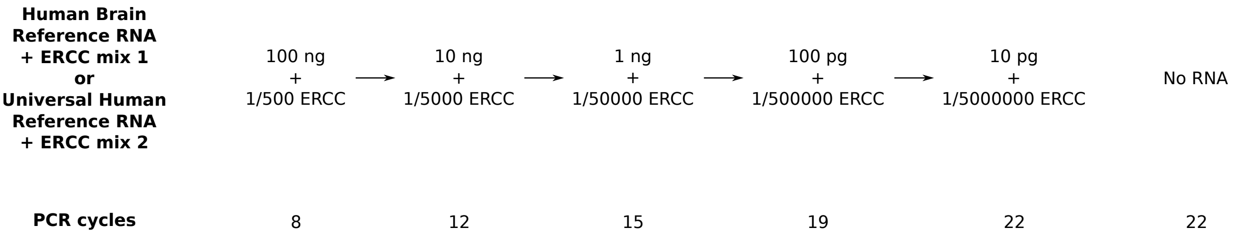


Figure S9: Experimental design of validation with human reference RNAs.

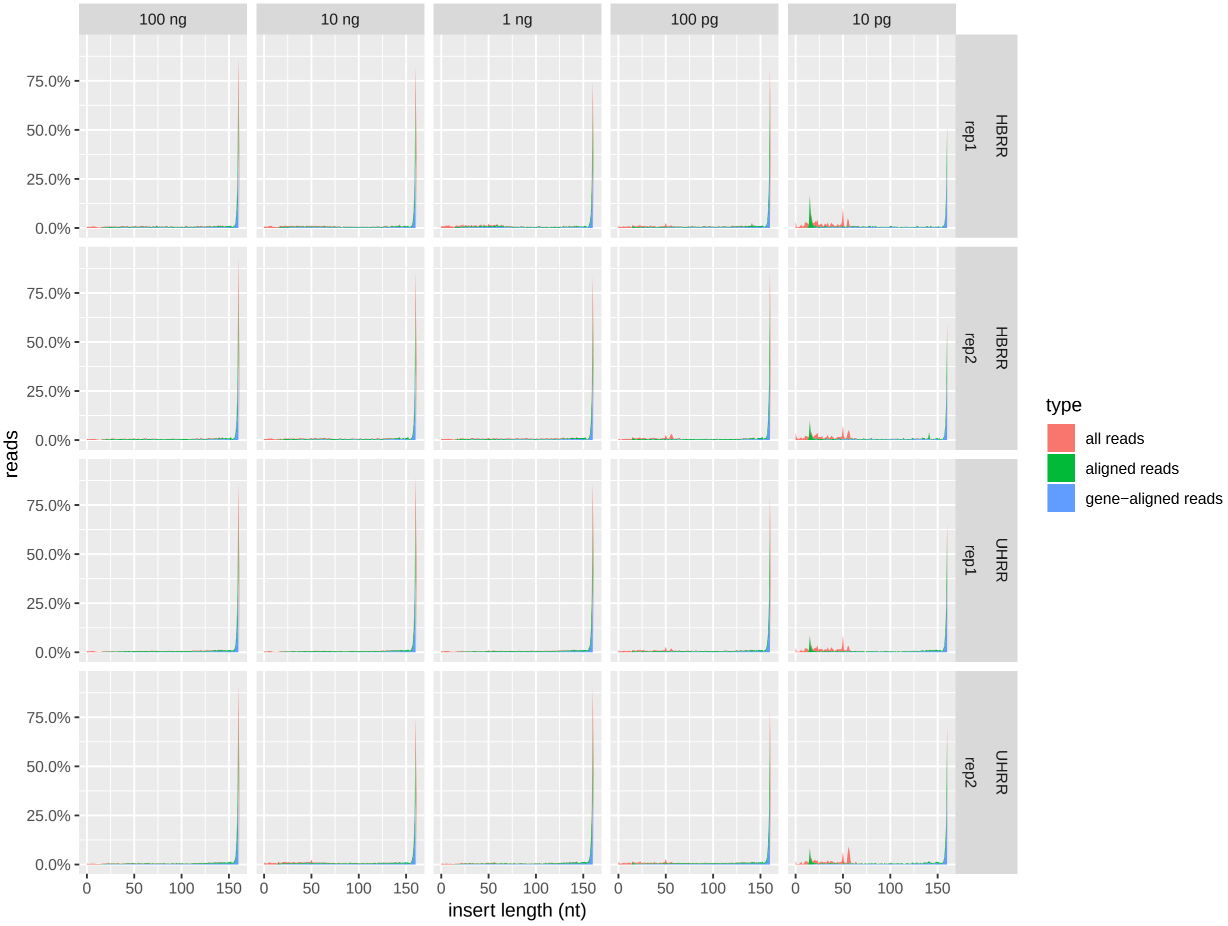


Figure S10: Insert lengths of Smart-3SEQ reads from human reference RNAs (MiSeq data). Insert length is inferred as the length of the read between the NNNNNGGG header and the beginning of the poly(A) tail or sequencing adapter; the read length is 169 nt and the header is 8 nt so all inserts longer than 161 nt are rounded down to 161. “All reads”: all post-filter reads reported by the MiSeq. “Aligned reads”: only the subset of reads that were aligned uniquely by STAR. “Gene-aligned reads”: only the further subset of reads that were aligned uniquely to an annotated transcript by featureCounts.

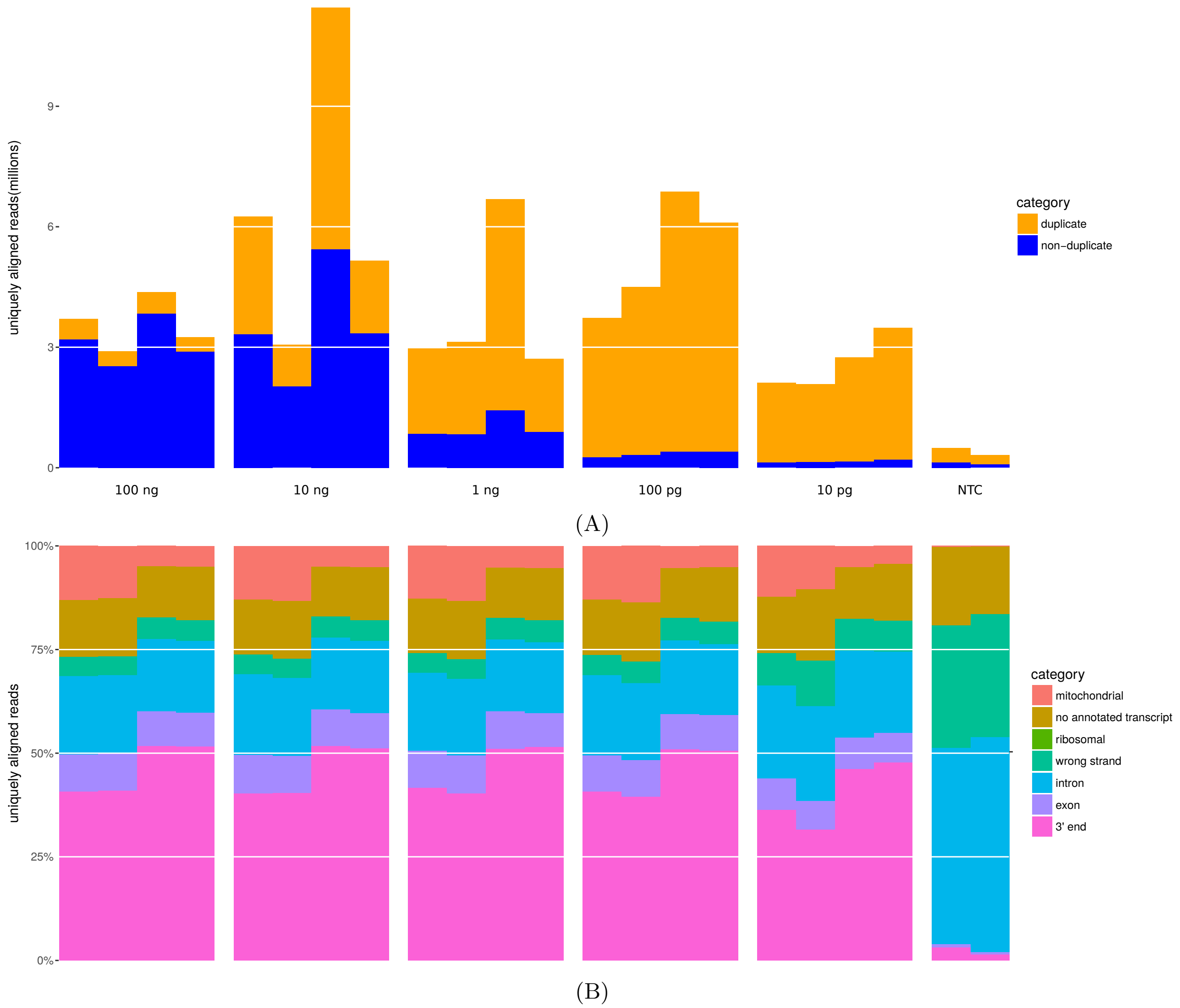


Figure S11: Quality statistics of Smart-3SEQ reads from human reference RNAs and no-template controls. A: PCR duplication, as measured by UMIs. B: Alignment position types. The "ribosomal" category is invisible because only 11 or fewer reads from any sample aligned to ribosomal genes.

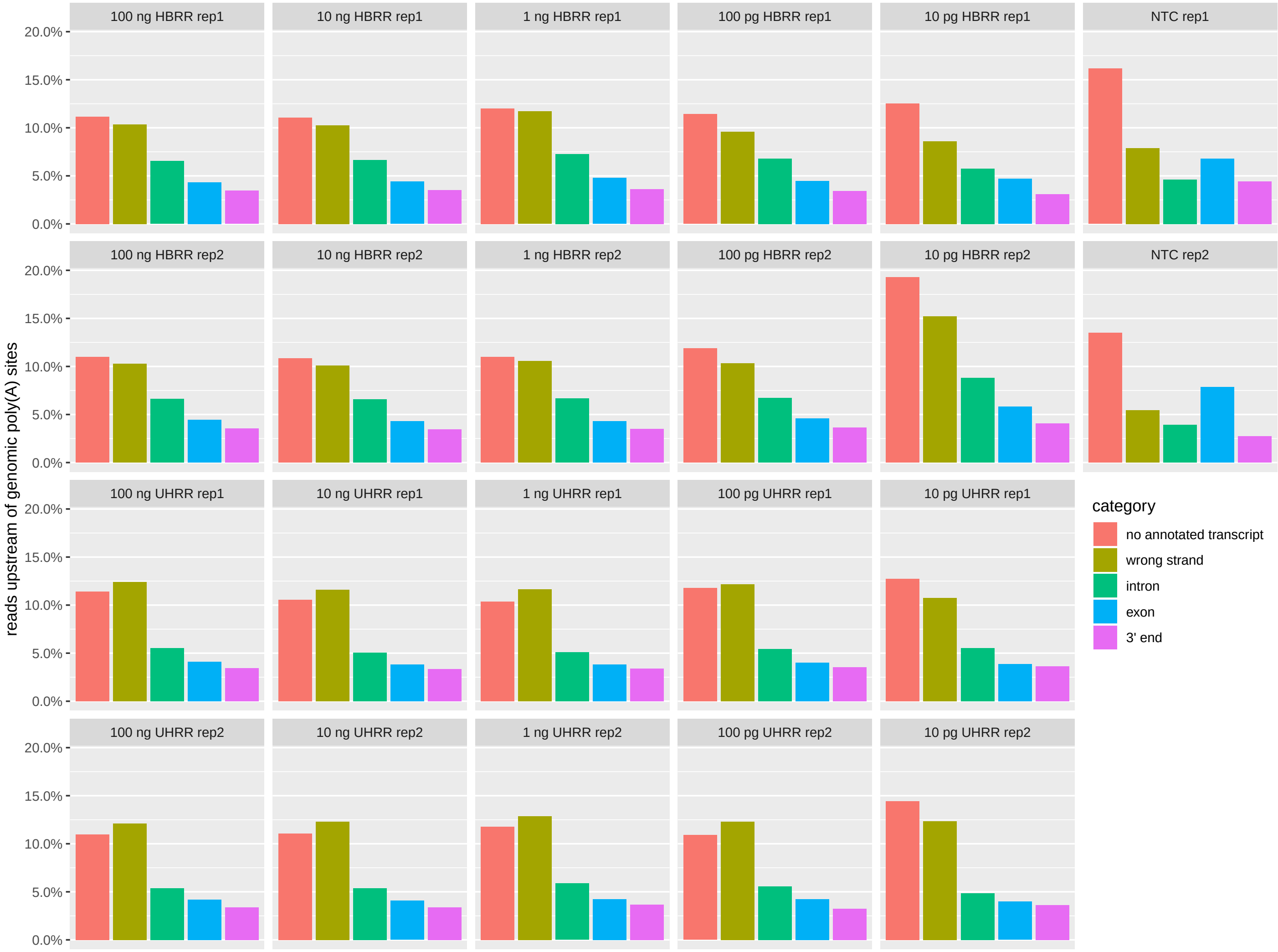


Figure S12: Smart-3SEQ reads from human reference RNA aligned upstream of genomic poly(A) sites, categorized as in Figure S11B. Poly(A) sites were identified as runs of 15 or more A's in the reference genome sequence, with non-A bases allowed as long as there were never two or more consecutive non-A bases and the first and last base of the site were both A's. A read was considered to be upstream of a poly(A) site if it was on the same strand and the entire length of the read aligned between the last base of the site and a range 100 nt upstream of the first base of the site. By this definition about 11% of bases in the reference genome are upstream of poly(A) sites. Mitochondrial and ribosomal hits were negligible because of the low number of poly(A) sites in the mitochondrial genome and the low number of reads aligned to ribosomal genes.

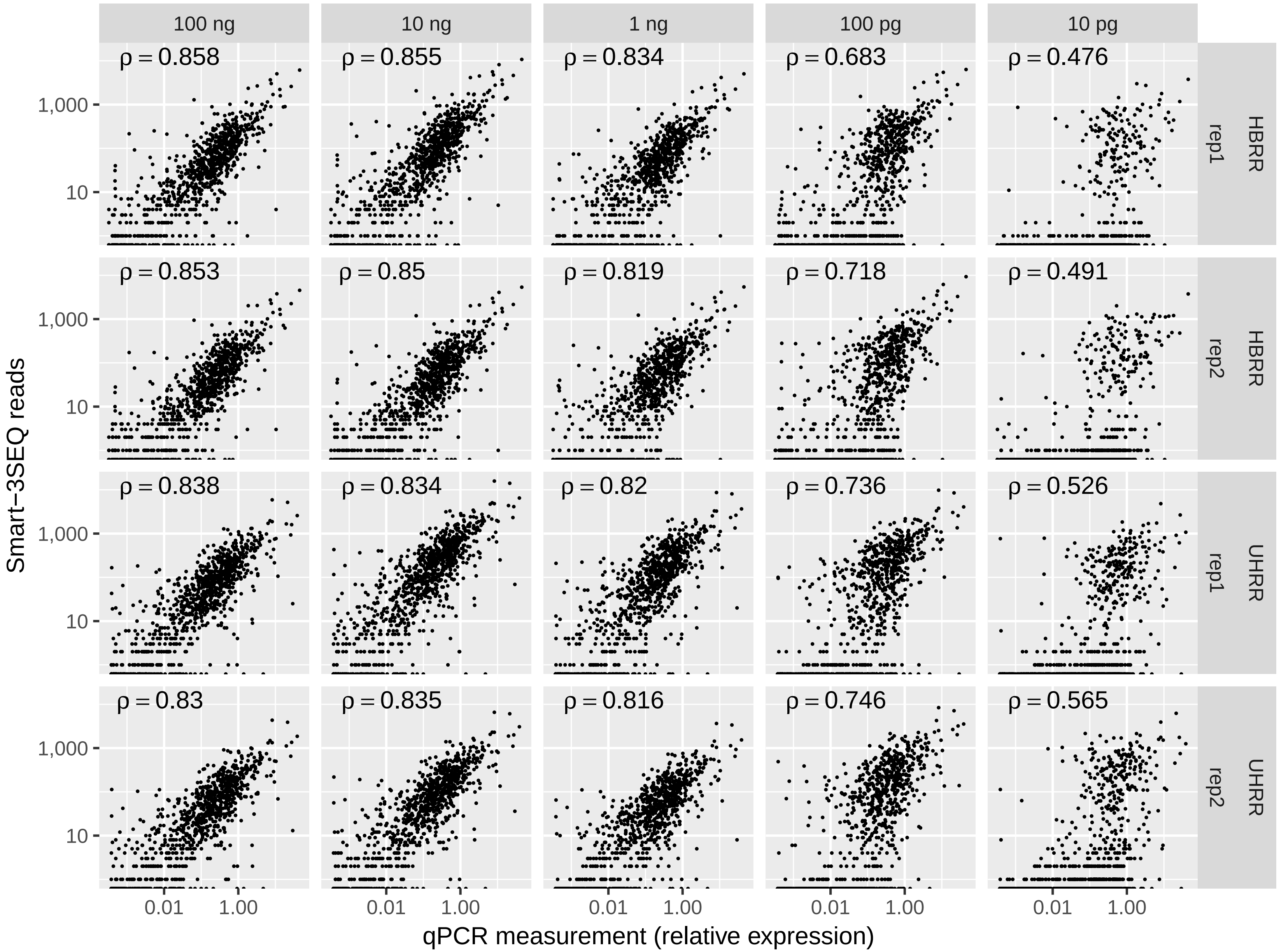


Figure S13: Correspondence of Smart-3SEQ results with TaqMan qPCR measurements. Each point is a single gene with available TaqMan qPCR data. $\rho$: Spearman's rank correlation.

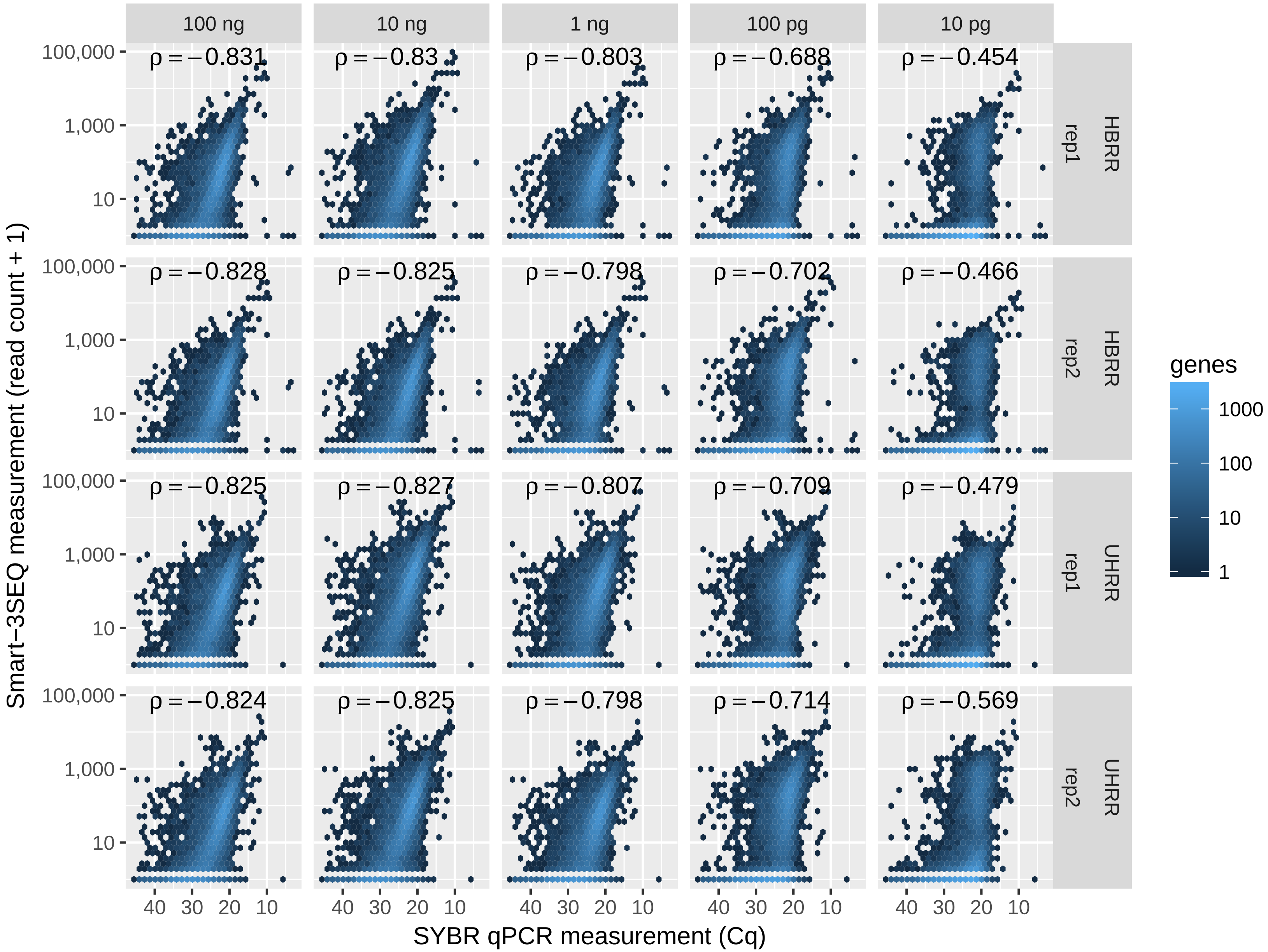


Figure S14: Correspondence of Smart-3SEQ results with SYBR qPCR measurements. Hexagonal bins are colored, on a logarithmic scale, by the number of genes among those with available SYBR qPCR data.

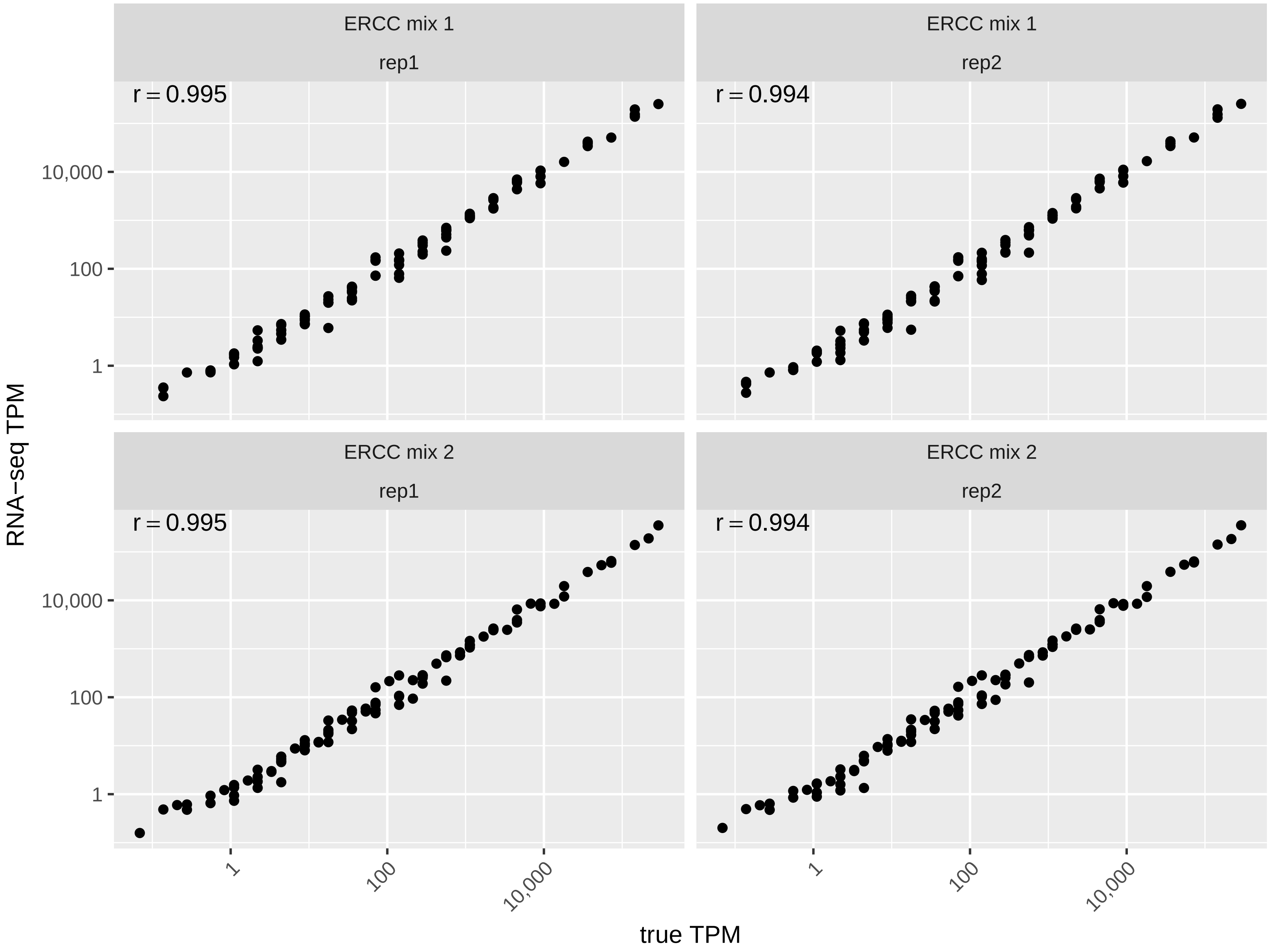


Figure S15: Accuracy of SEQC RNA-seq on ERCC standards.

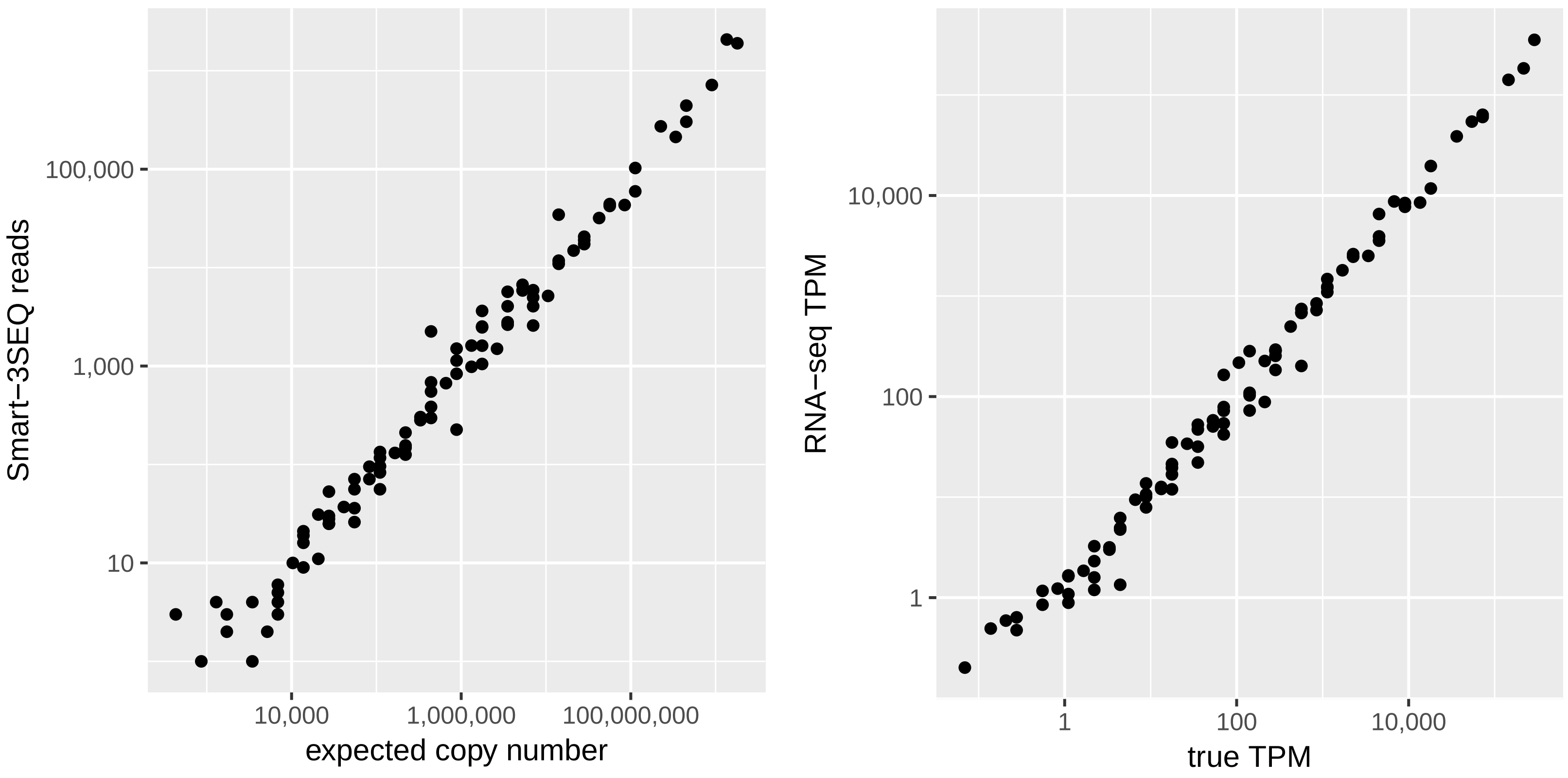


Figure S16: Comparative accuracy of Smart-3SEQ and SEQC RNA-seq on ERCC standards using the libraries with the greatest sequencing depth. Smart-3SEQ $r' = 0.990$, RNA-seq $r = 0.994$.

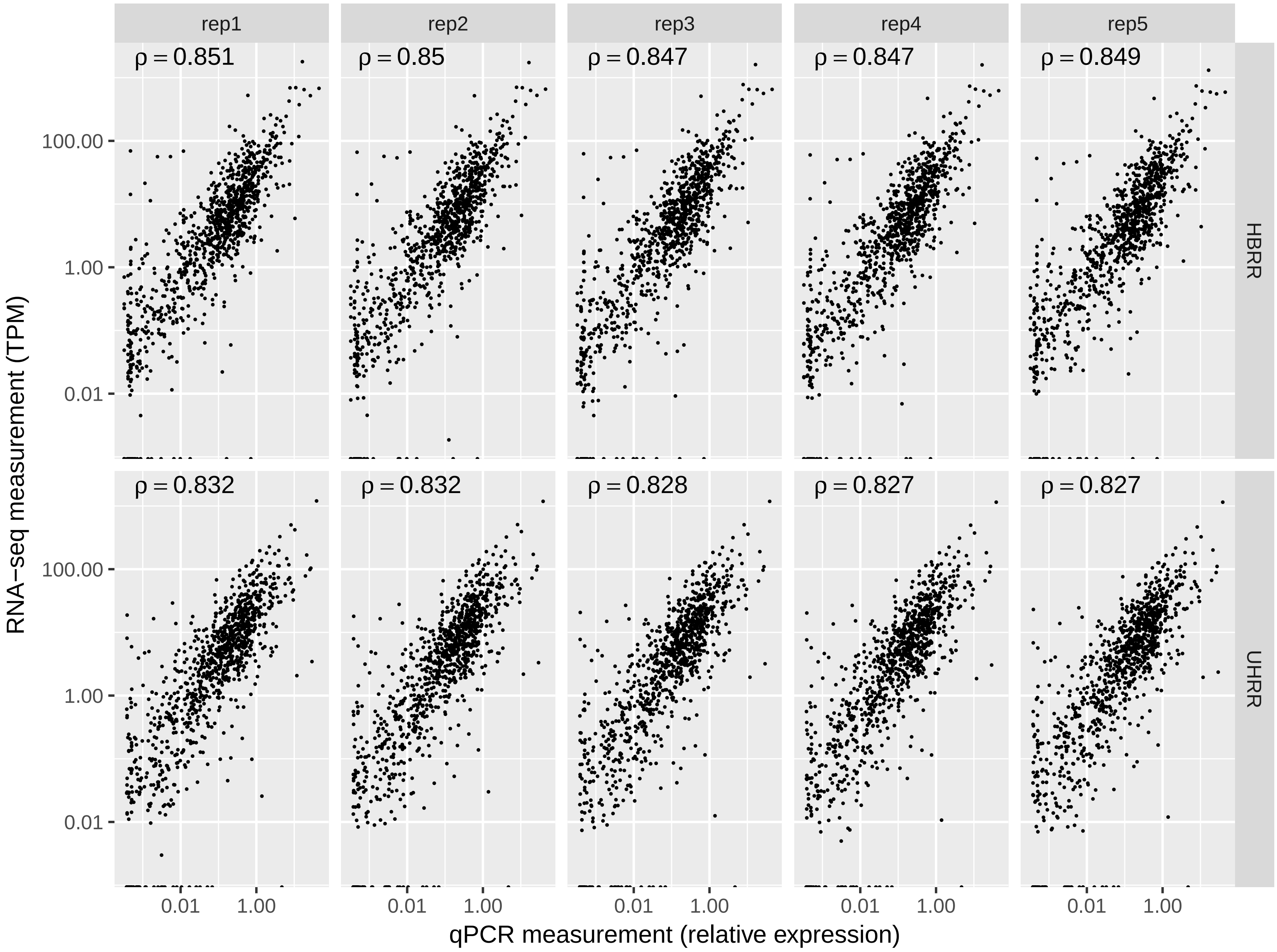


Figure S17: Correspondence of SEQC RNA-seq results with TaqMan qPCR measurements.

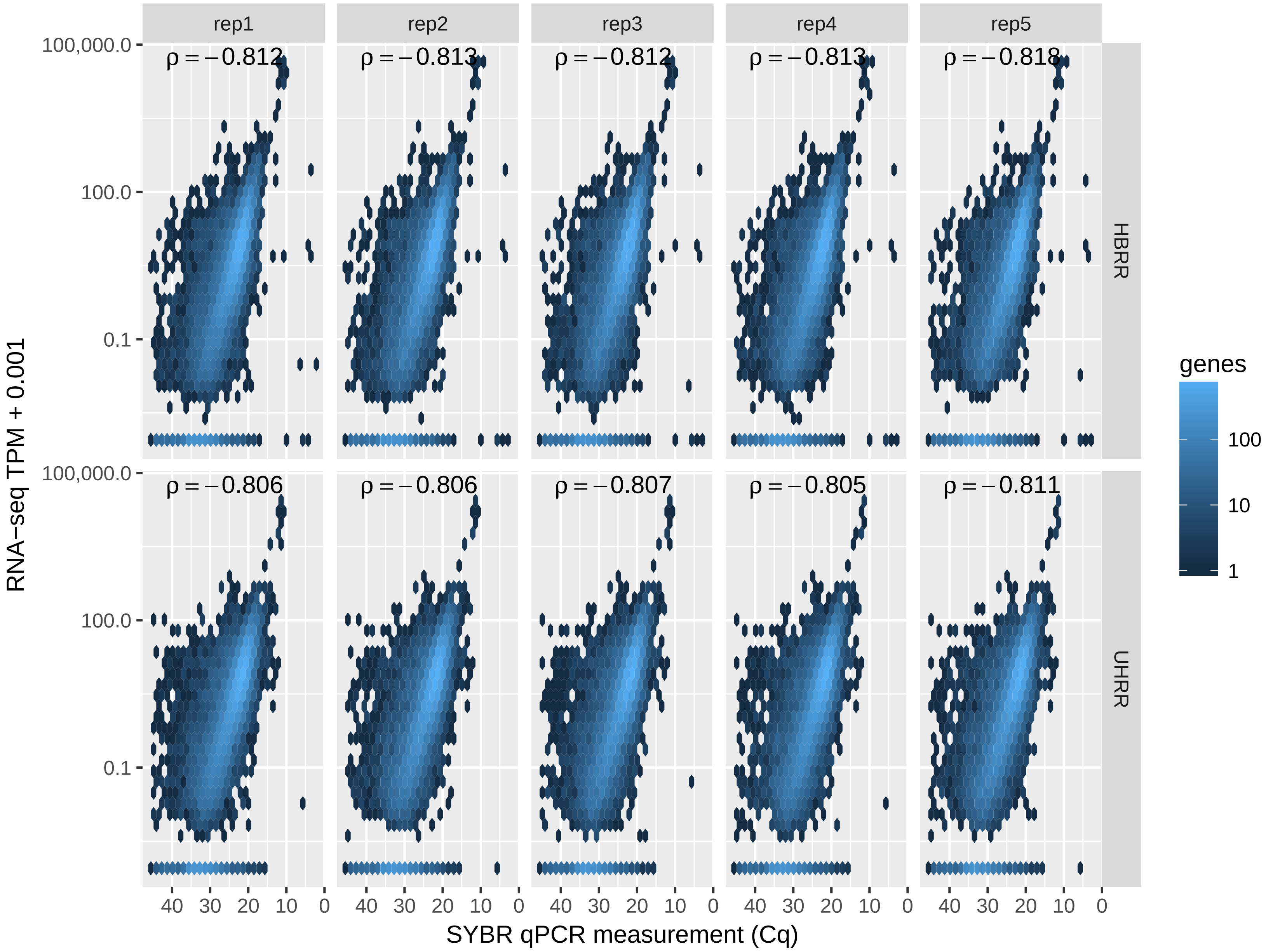


Figure S18: Correspondence of SEQC RNA-seq results with SYBR qPCR measurements.

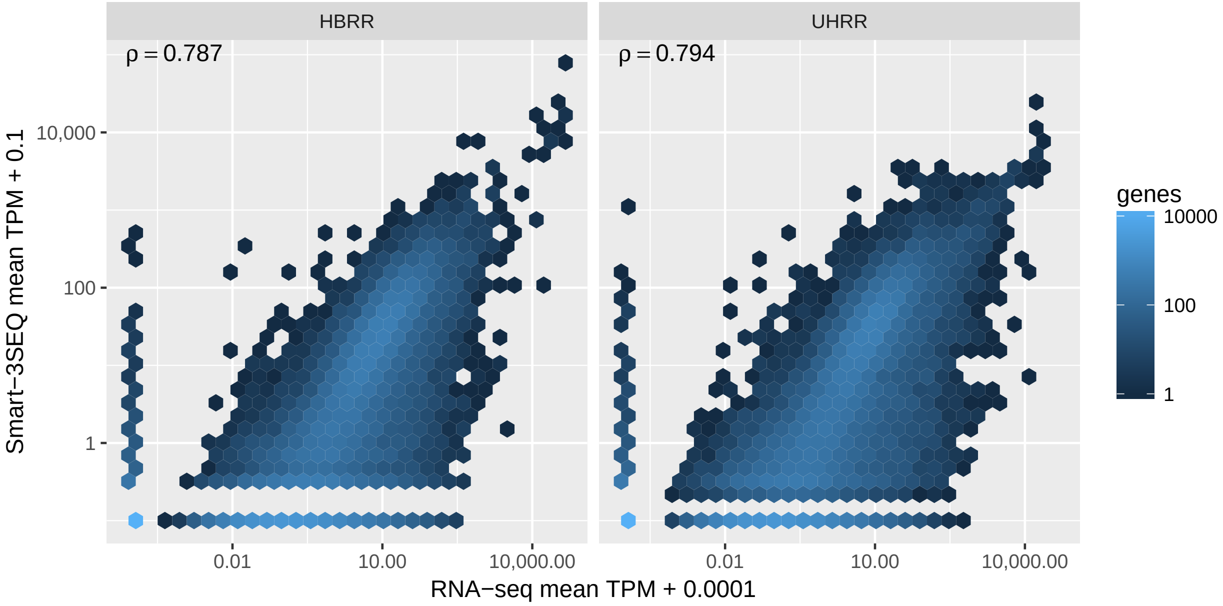


Figure S19: Correspondence of Smart-3SEQ results (means of all 100 ng replicates) with SEQC RNA-seq results (means of all replicates). All annotated genes are shown.

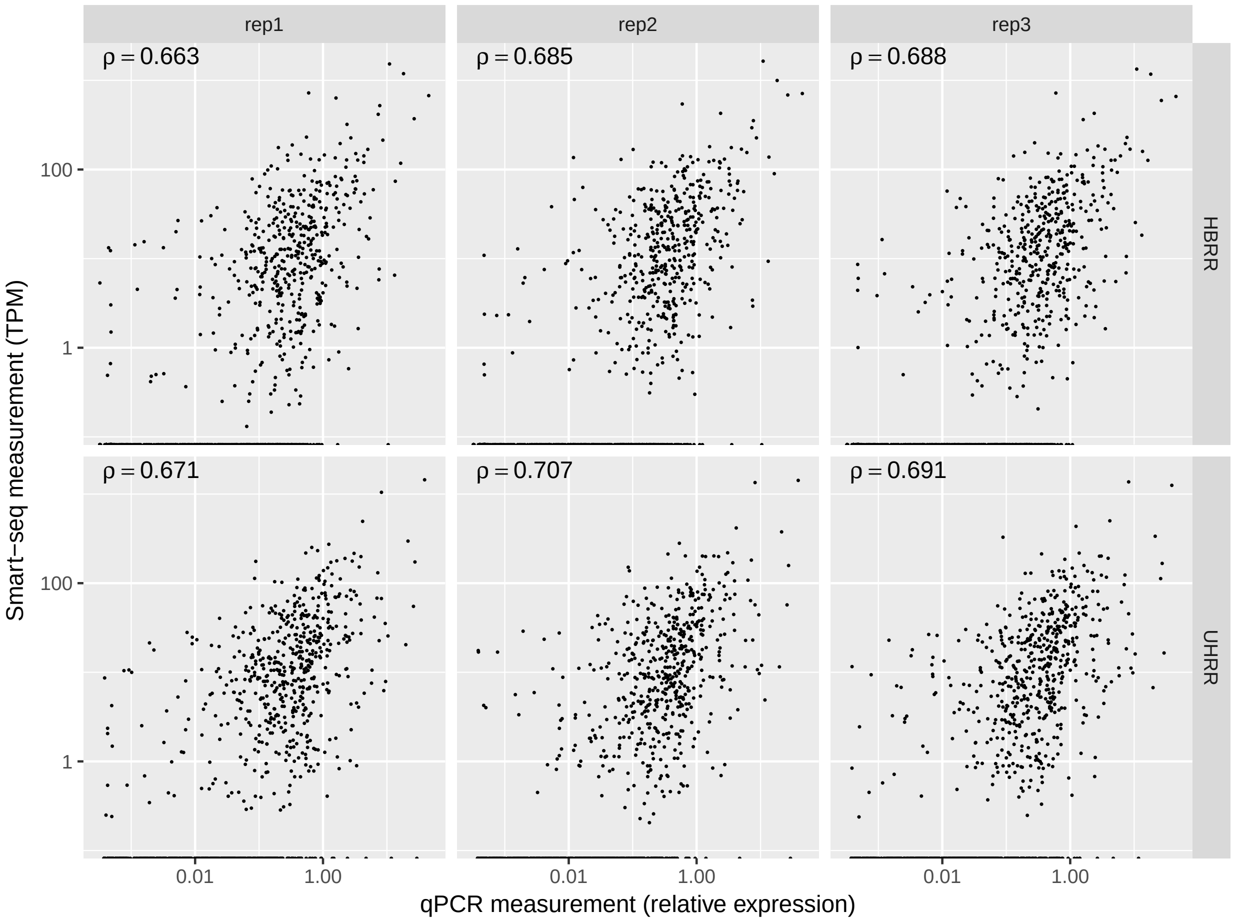


Figure S20: Correspondence of Takara SMART-Seq v4 results with TaqMan qPCR measurements.

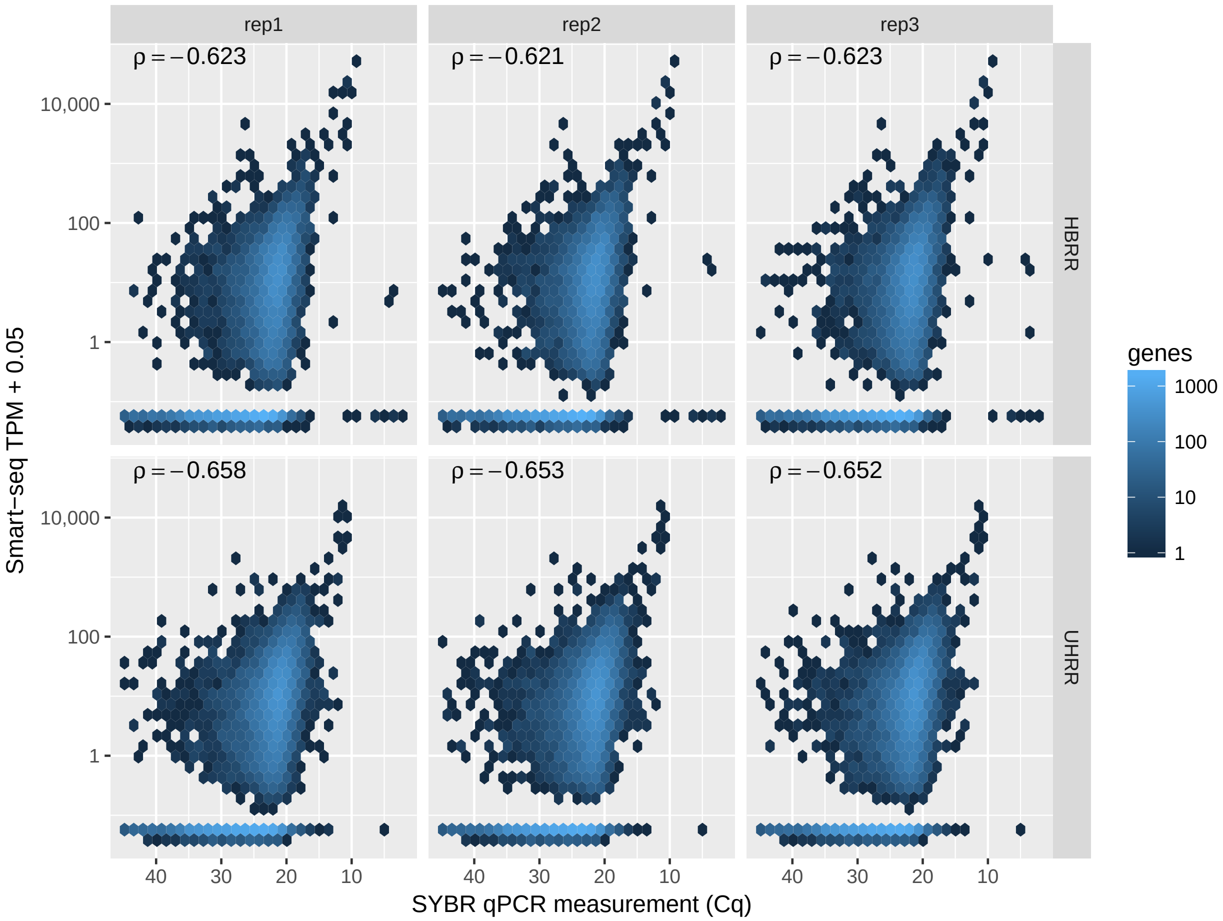


Figure S21: Correspondence of Takara SMART-Seq v4 results with SYBR qPCR measurements.

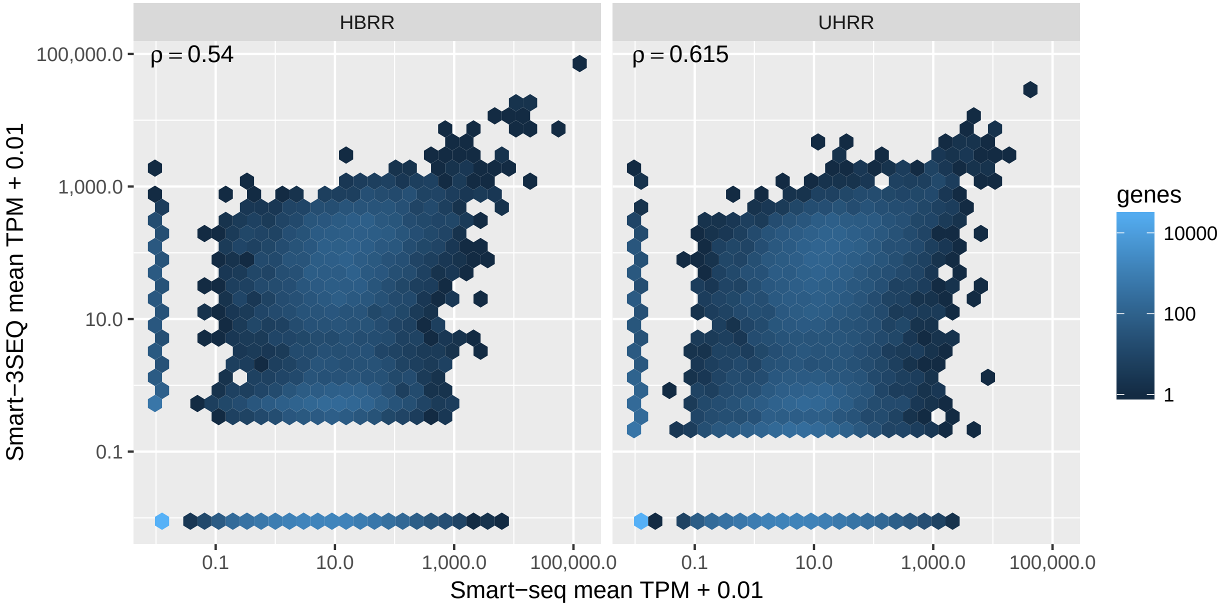


Figure S22: Correspondence of Smart-3SEQ results (means of all 10 pg replicates) with Takara SMART-Seq v4 results (means of all replicates). All annotated genes are shown.

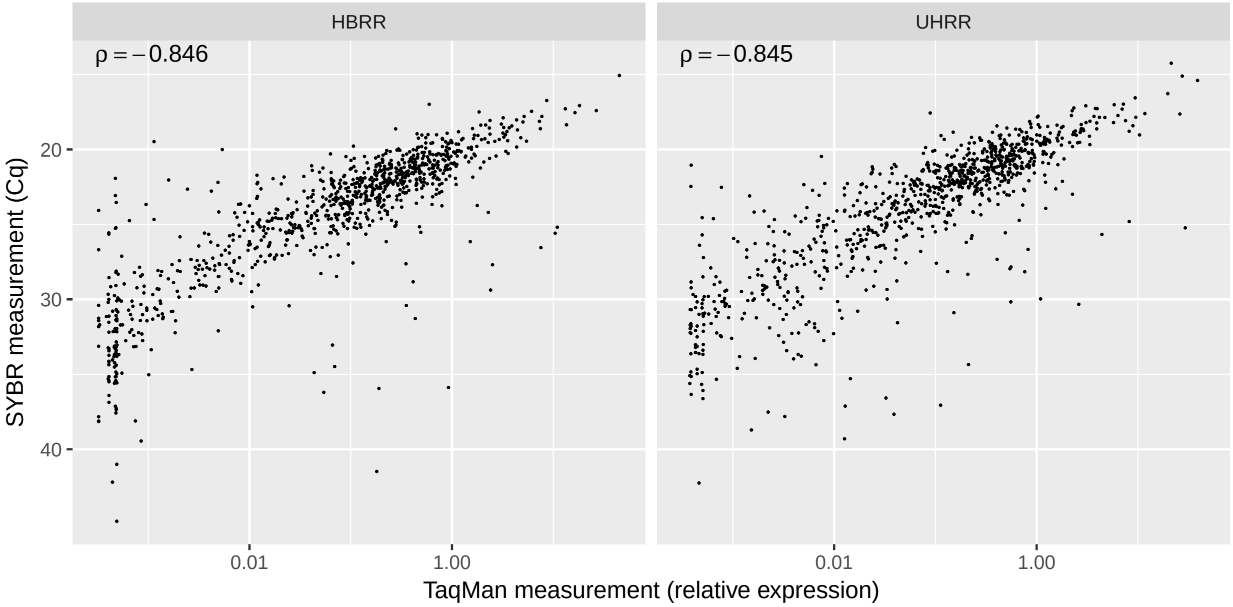


Figure S23: Correspondence of the two different qPCR measurements. Each point is a single gene with available data from both qPCR platforms.

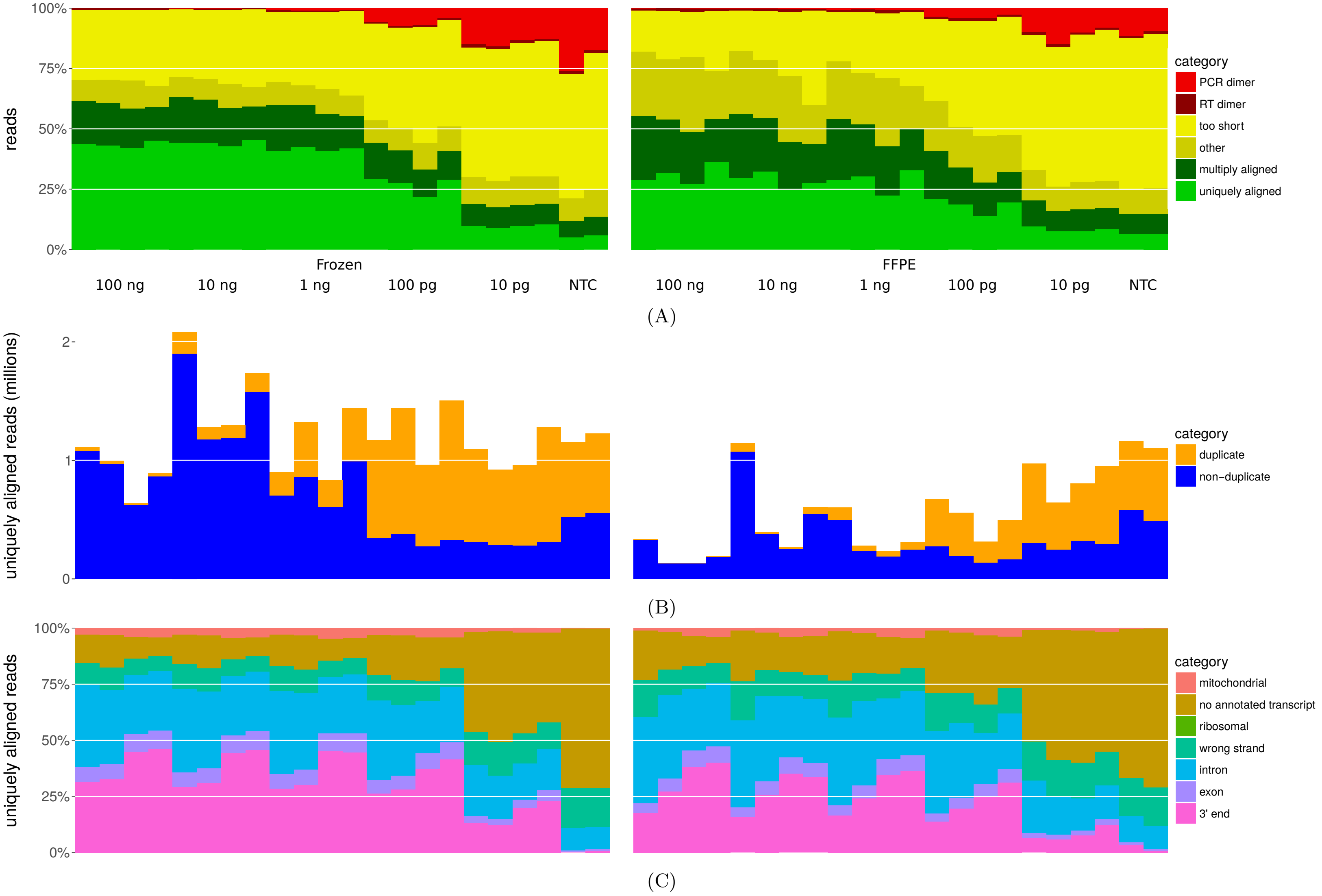


Figure S24: Quality statistics of Smart-3SEQ reads from RNA isolated from frozen vs. FFPE tissue. In order from left to right, each dilution level's libraries are SFT1, SFT2, PVNS1, PVNS2. A: Read alignability. B: PCR duplication, as measured by UMIs. C: Alignment position types. The "ribosomal" category is invisible because only 10 or fewer reads from any sample aligned to ribosomal genes. SFT1: frozen RIN 8.4, FFPE RIN 2.3. SFT2: 6.8, 2.4. PVNS1: 6.2, 2.4. PVNS2: 8.9, 4.3.

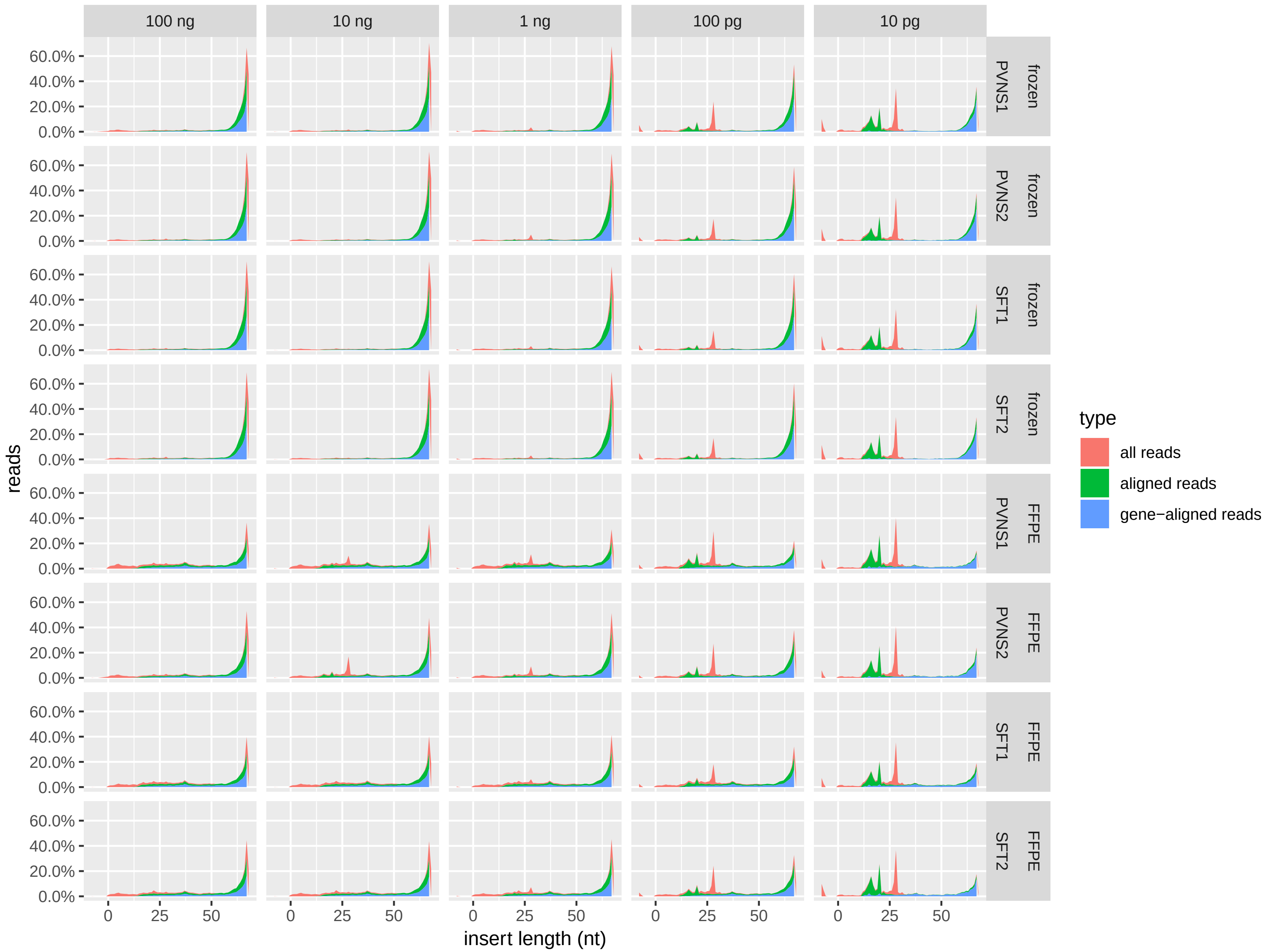


Figure S25: Insert lengths of Smart-3SEQ reads from RNA isolated from frozen vs. FFPE tissue. Insert length is inferred as in Figure S10; the read length is 76 nt so long inserts are rounded down to 68. An insert length of 0 implies an RT primer dimer (adapters and header but no cDNA), while −8 implies a PCR primer dimer (no header either).

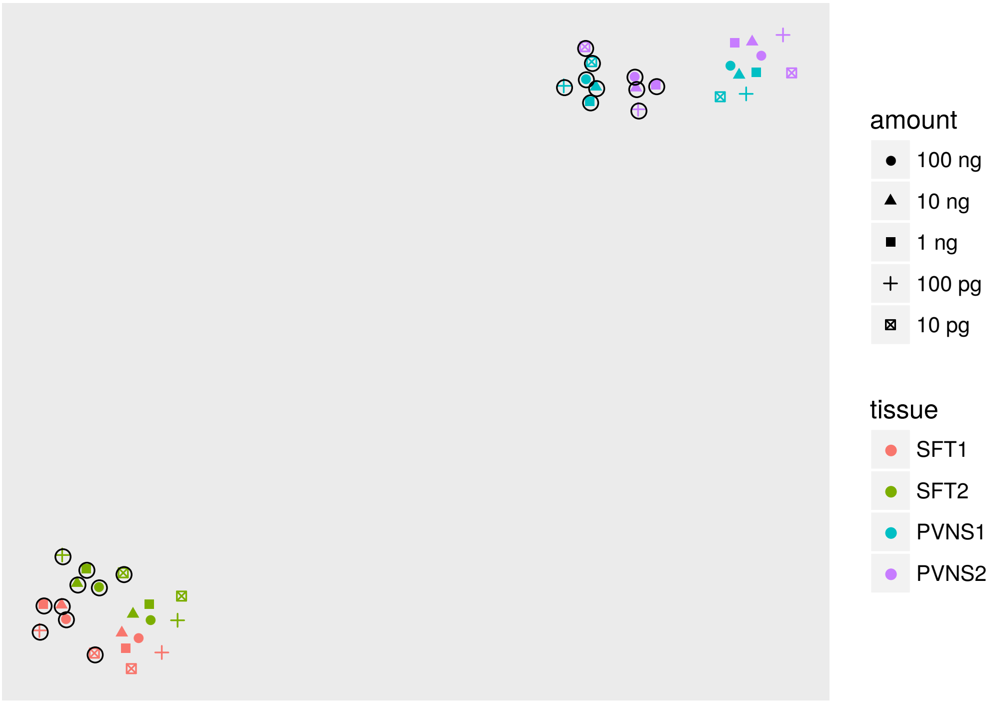


(A)

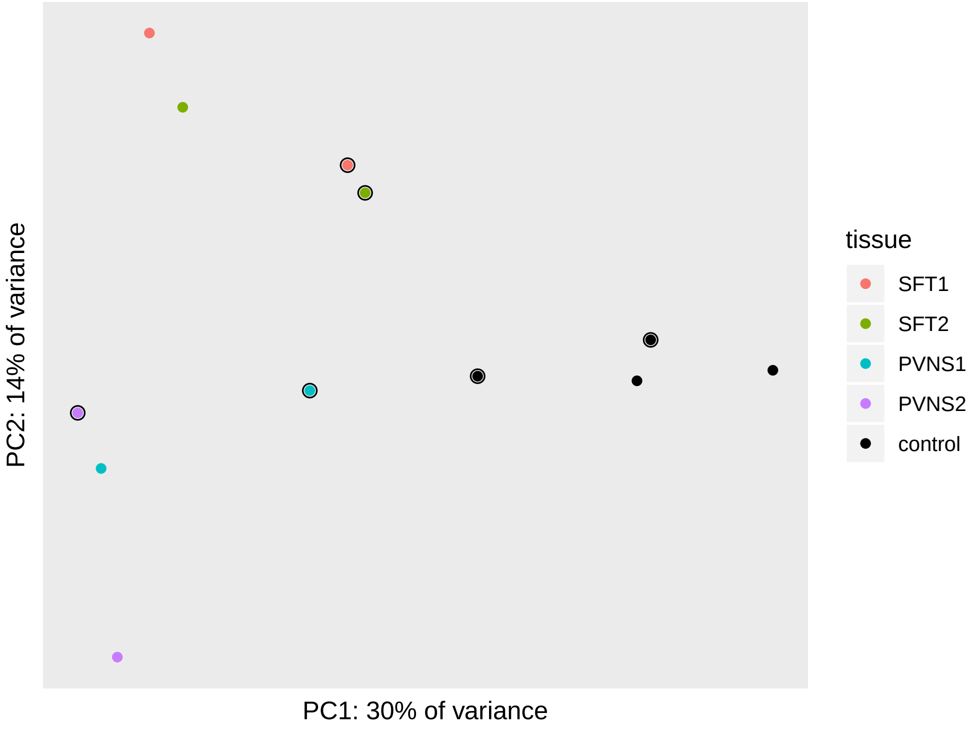


(B)

Figure S26: Cluster analysis of all genes in libraries from the comparison of fresh-frozen vs. FFPE tissue. FFPE samples (or controls from the protocol for damaged RNA) are circled and frozen samples are uncircled. SFT1: frozen RIN 8.4, FFPE RIN 2.3. SFT2: 6.8, 2.4. PVNS1: 6.2, 2.4. PVNS2: 8.9, 4.3. A: *t*-SNE of all non-control libraries. B: PCA of only the 10 pg libraries and the no-template controls.

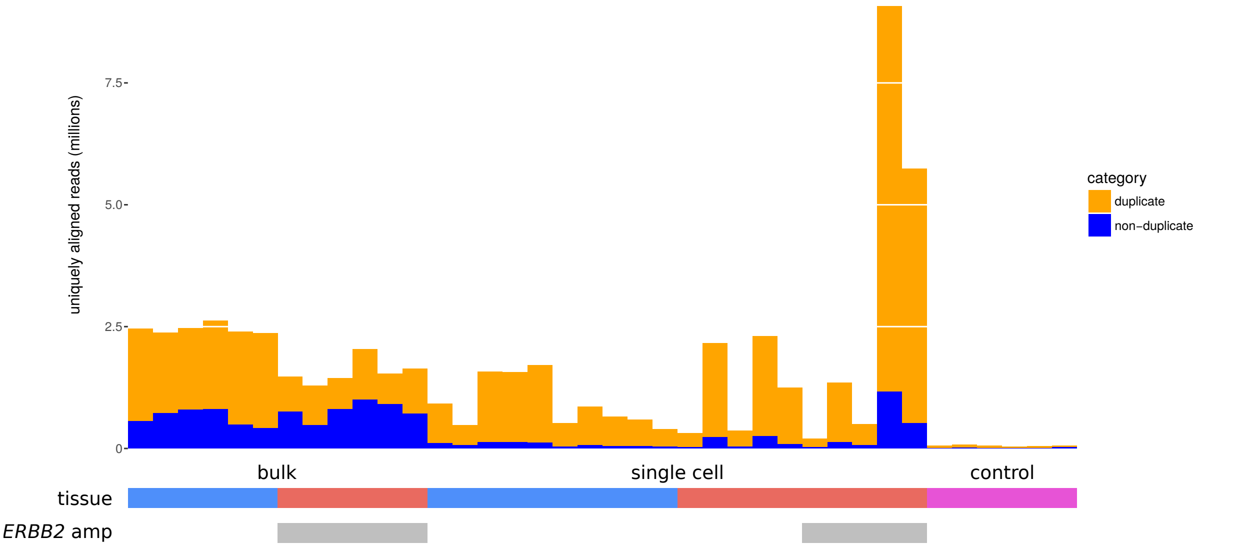


(A)

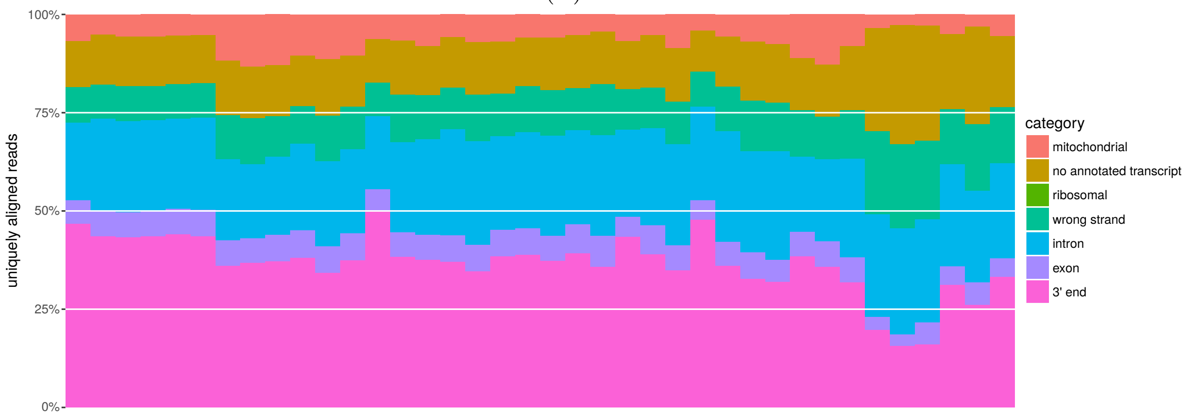


(B)

Figure S27: Quality statistics of Smart-3SEQ reads from LCM FFPE samples and no-template controls. A: PCR duplication, as measured by UMIs. B: Alignment position types. The “ribosomal” category is invisible because only 50 or fewer reads from any sample aligned to ribosomal genes.

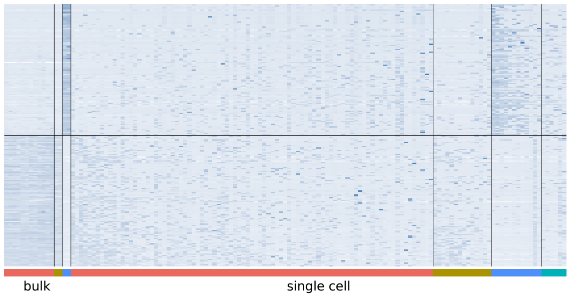


(A)

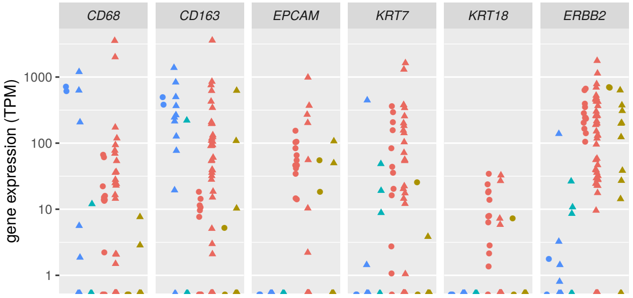


(B)

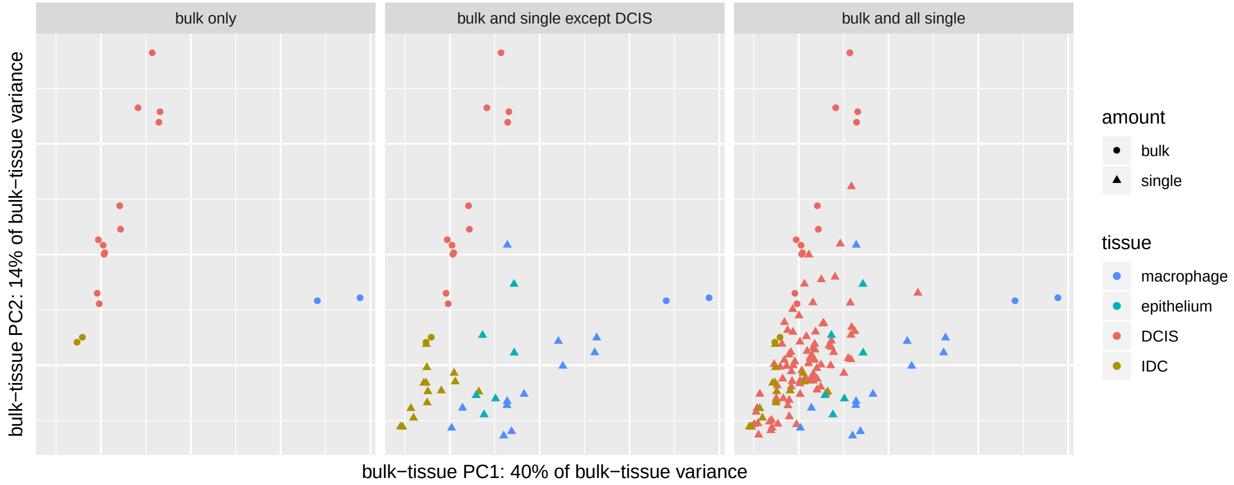


(C)

Figure S28: Replication of the FFPE LCM experiment on a larger scale: 12 bulk DCIS, 2 bulk macrophage, 2 bulk invasive ductal carcinoma (IDC, a more advanced tumor), 96 single DCIS, 16 single macrophage, 16 single IDC, and 6 single normal epithelium cells as positive controls in addition to 6 negative controls, all from a second case of breast cancer. Color scheme is the same in all panels. A: Expression (regularized log read count, normalized by row) of the 100 genes with the greatest enrichment in bulk macrophage relative to bulk tumor (both DCIS and IDC) and the 100 genes with the opposite enrichment, all significant at $p_{\text{adj}} < 0.01$. Single-cell libraries are displayed in order, from left to right, of decreasing similarity (Pearson correlation) to the corresponding bulk profile (mean of bulk libraries). B: Expression (transcripts per million) of known marker genes for macrophage (*CD68*, *CD163*) and tumor (*EPCAM*, *KRT7*, *KRT18*, *ERBB2* (*HER2*)). C: Principal components analysis of all genes (regularized log read counts), with both bulk and single-cell libraries rotated onto PCs that were trained on only the bulk data. The first subpanel shows only the bulk libraries, the second subpanel adds the single cells of all types except DCIS, and the third subpanel adds the single DCIS cells as well, thus showing all libraries from this experiment.

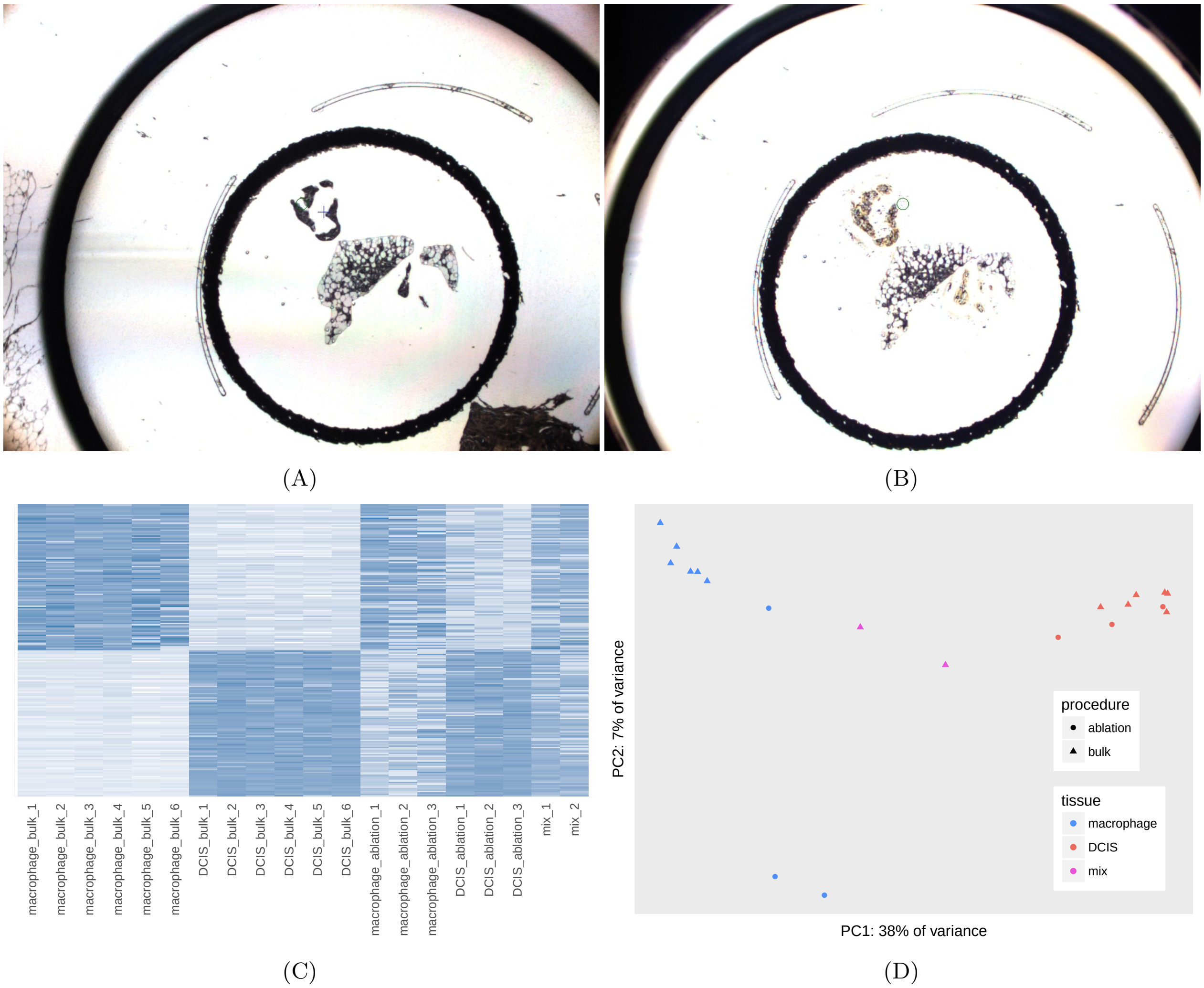


Figure S29: Validation of the laser ablation method. A: Example of mixed DCIS regions and macrophage regions on the same LCM cap. B: DCIS tissue ablated by destruction with the UV laser. C: Expression (regularized log read count, normalized by row) of the 100 genes with the greatest enrichment in bulk macrophage relative to bulk DCIS and the 100 genes with the opposite enrichment, all significant at $p_{adj} < 0.05$. Because the genes are chosen by differential expression in the bulk samples, those samples necessarily show the strongest distinction in this figure, but that distinction is recapitulated independently in the ablated samples and not in the unablated mixes. D: Principal components analysis of all genes. Note that the first PC, which discriminates DCIS from macrophage, captures substantially more variance than the second and all subsequent PCs; lower-ranked PCs commonly isolate a small number of samples by detecting common outliers after higher-ranked PCs have captured the strongest variables in the experimental design.

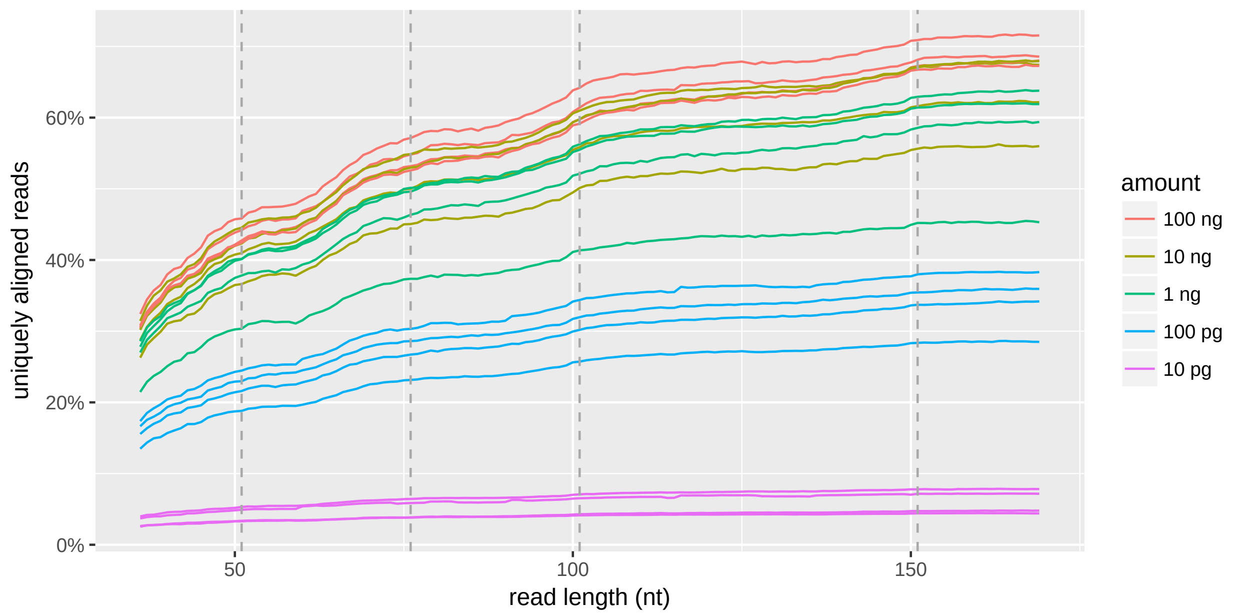


(A)

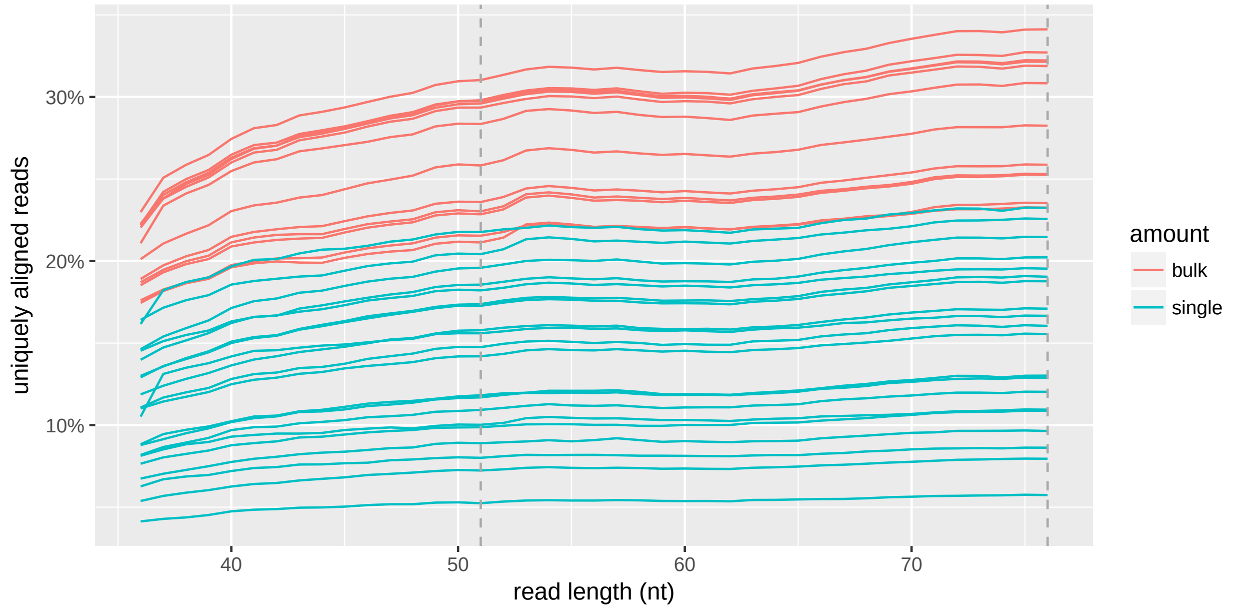


(B)

Figure S30: Diminishing returns from increased read lengths. Results are simulated by truncating reads to the specified length and rerunning the alignment pipeline. Each line traces results from a single library. Commonly used lengths are noted with dashed lines. A: Dilution series of human reference RNAs (high-quality RNA; long fragments), sequenced with long reads on an Illumina MiSeq. B: Bulk tissue and single cells from LCM on FFPE tissue (degraded RNA; short fragments).

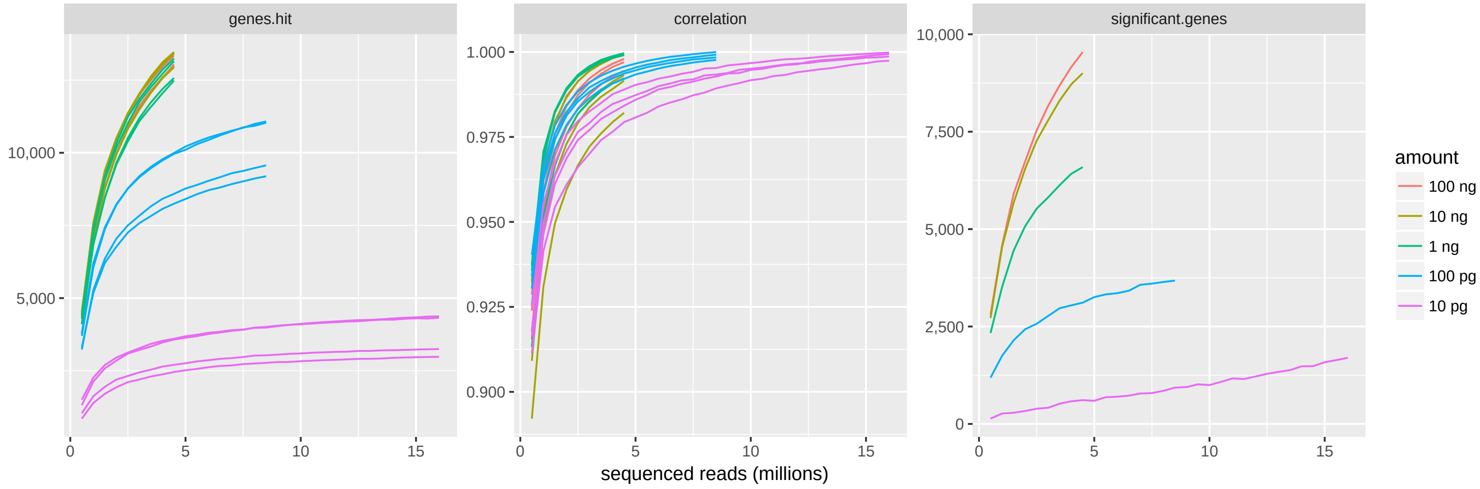


(A)

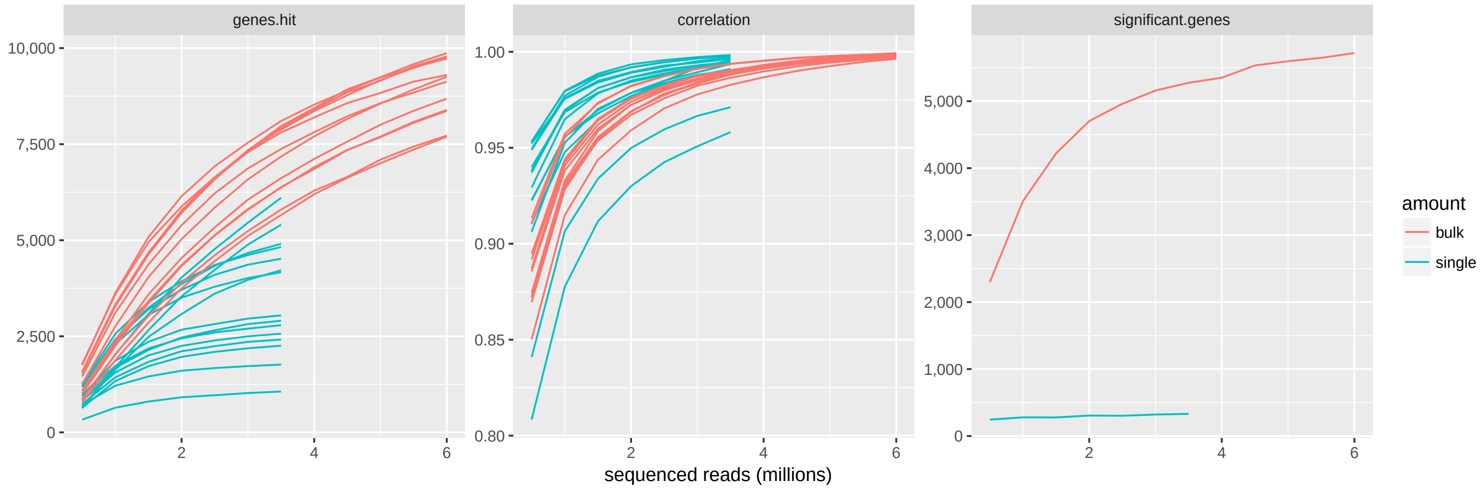


(B)

Figure S31: Diminishing returns from increased sequencing depth. Results are simulated by binomial subsampling of the gene-aligned read counts. `genes.hit`: number of genes with at least 10 reads aligned. `correlation`: Pearson correlation of gene-expression values (as $\log_{10}(c+1)$ for read count $c$) between subsampled data and original. `significant.genes`: number of genes with $p_{\text{adjusted}} < 0.05$ for significant differential expression between biological categories (HBRR vs. UHRR or DCIS vs. macrophage), using only the data from this amount of RNA/tissue. A: Dilution series of human reference RNAs (HBRR vs. UHRR); 2 library replicates per condition at each dilution. B: Bulk tissue and single cells from LCM on FFPE tissue (macrophage vs. DCIS; single DCIS-labeled cells lacking *ERBB2* amplification not included); 6 dissection replicates per condition for bulk and 10 vs. 5 single cells.
