## Supplemental File 2: Smart-3SEQ protocol for "Gene-expression profiling of single cells from archival tissue with laser-capture microdissection and Smart-3SEQ"

### Gene-expression profiling by Smart-3SEQ

Joseph W. Foley
Department of Pathology
Stanford University School of Medicine
January 8, 2019

#### Abstract

This protocol combines aspects of 3SEQ (Beck et al., PLOS ONE 2010), Smart-seq2 (Picelli et al., Nat Methods 2013), and UMIs (Kivioja et al., Nat Methods 2011) into a streamlined method for making a sequencing library to measure the expression of polyadenylated RNAs. It is fast (3 hr), inexpensive (< $10 / library), sensitive (tested successfully with 10 pg human total RNA, equivalent to one cell), and scalable (all steps performed in PCR tubes or microplates). It generates digital gene-expression data equivalent to 3SEQ's: one read per RNA molecule in the starting material, at the 3′ end of the gene, which allows simpler quantification of transcript abundance at the expense of most information about splicing isoforms.

#### Experimental design considerations

Gene-expression profiling requires large sample sizes for powerful statistical analysis, like any other biological experiment. Because this protocol is rapid and inexpensive, it is not necessary to scale down your experiment to fit time or budget constraints; it is better to sequence 10 biological replicates at 50% depth than 5 replicates at 100% depth (see e.g. Liu et al., Bioinformatics 2014). As such, some volumes in this protocol may be impractical to pipet accurately for a single tube, and are intended to be prepared in a master mix with excess volume.

Plan the allocation of your index sequences before beginning. If possible, every library in your entire experiment should have a unique index, allowing you to sequence them all together in every lane/flowcell and avoid confounding by any technical variation on the sequencer, though this is not likely to be large. On the other hand, if you are pooling a very small number of libraries, you must take care to balance the color usage in the sequencer. See Illumina's “TruSeq Library Prep Pooling Guide” for details.

This method is sensitive over at least five orders of magnitude of input amount; the only variable that needs to be changed is the number of PCR cycles. It is ideal to dilute your more concentrated samples so that all samples in the experiment start from the same amount of material. However, an accurate measurement of that amount is crucial; fluorometry (Qubit, RiboGreen, Bioanalyzer) is strongly recommended instead of spectrophotometry.

#### Pre-SPRI pooling option

The normal protocol makes a separate library from each RNA sample (Option 1: Individual libraries). However, if you are working in large batches, you can combine the libraries before the purification step and elute the pool into the same volume, concentrating the yield (Option 2: Pre-SPRI pooling). This saves money and time, and also reduces the number of PCR cycles required to generate enough material for QC and sequencing. It is crucial to pool low-input libraries that would require too many PCR cycles if processed separately. Every library in the pool must have a different index sequence, and this option may result in less balanced read counts from library to library than if you quantify them separately.

#### Choosing the number of PCR cycles

Compute the number of required PCR cycles based on the amount of total RNA input, with the formula

$$\text{cycles} = 19 - \frac{\log(N \times \text{ng total RNA per sample})}{\log(1.9)}$$

where $N$ is the number of samples you will combine with the pre-SPRI pooling option, or $N = 1$ if you use the individual library option.

The nomogram on the final page of this document shows the results of the formula. For example, if you prepare individual libraries from 100 ng total RNA each, use 12 PCR cycles; or if you pool 25 libraries made from 1 ng total RNA each, you can use as few as 14 PCR cycles. If you use 20 or more PCR cycles, you will get a lower proportion of usable reads, due to the abundance of adapter dimers. Avoid using 24 or more PCR cycles.

This was calibrated on intact human RNA and may need adjustment for different kinds of samples. Recalibrate by testing a fixed amount of RNA with different numbers of cycles. For convenience, you can run all the tubes in the same program, then quickly open the thermal cycler and transfer a tube to 72 °C for 1 min (in a second thermal cycler holding constant temperature) at the end of each cycle. Do not omit this final 72 °C incubation. Aim for the highest number of cycles that does not show signs of overamplification (see "Quantification and validation of the libraries").

#### Required materials

##### *Equipment*

- Adjustable-volume micropipets
- Programmable thermal cycler(s)
- Mini-centrifuge for PCR tubes or microplates, e.g. Ohaus #FC5306, Fisher #14-100-143
- *Required only for individual library option:* Magnetic separation block for 0.2 mL PCR tubes or microplates, e.g. V&P #772F4-1, #771LD-4CS,
- *Required only for pre-SPRI pooling option:* Magnetic separation block for 1.5/2.0 mL microcentrifuge tubes, e.g. Thermo Fisher #12321D
- *Recommended:* separate workstations for pre-PCR and post-PCR steps, to avoid contamination

##### *Consumables*

- Low-retention 0.2 mL PCR tubes or microplates, e.g. Axygen Maxymum Recovery, Eppendorf LoBind
- *Required only for pre-SPRI pooling option:* Low-retention 1.5 or 2.0 mL microcentrifuge tubes, e.g. Eppendorf LoBind
- *Required only for Arcturus LCM with HS caps:* GeneAmp 0.5 mL PCR tubes, Thermo Fisher #N8010611
- *Recommended:* low-retention disposable pipet tips, e.g. Eppendorf Dualfilter T.I.P.S. LoRetention

##### *Reagents*

- SMARTScribe reverse transcriptase (includes 5X reaction buffer and 20 mM DTT), Clontech #639536
- Nuclease-free dNTP mix, 10 mM each, e.g. Thermo Fisher #R0192
- Trimethylglycine (TMG; also called glycine betaine), 5 M solution, e.g. Sigma-Aldrich #B0300
- RNase inhibitor, 20X, e.g. Thermo Fisher #AM2694
- Magnesium chloride ($MgCl_2$), 80 mM solution (200 mM for FFPE LCM), e.g. Sigma-Aldrich #63069
- HiFi HotStart ReadyMix, 2X, Kapa #KK2601
- DNA SPRI bead mix, e.g. Beckman Coulter #B23317 or homemade
- 80% ethanol in molecular biology–grade water, freshly prepared on the day of the procedure
- DNA storage buffer, e.g. TE, TE+Tween, Qiagen EB (pure water is not a buffer and not recommended)
- Oligonucleotide primers (see "Oligonucleotide primer designs (Illumina-specific)")
- *Required only if using cells or tissue:* Triton X-100, 0.5% (v/v) in molecular biology–grade water
- *Required only for LCM FFPE tissue:* Proteinase K, e.g. NEB #P8107S
- *Required only for LCM FFPE tissue:* Proteinase K inhibitor, EMD Millipore #539470 (resuspend at 5 mM in DMSO)
- *Optional to reduce cost for pre-SPRI pooling option:* DNA SPRI bead binding buffer

Equivalent reagents can probably be substituted for most of these recommendations with no problems, but do not change the enzyme mixes (Clontech SMARTScribe and Kapa HiFi). DTT, dNTPs, and proteinase K inhibitor are unstable and/or susceptible to degradation by repeated freezes and thaws; upon receipt, separate these into small aliquots and keep them frozen.

#### Thermal cycler programs

##### *Program 1: Frag + TS-RT*

| | | |
|---|---|---|
| 80 °C | hold | (add tube containing RNA and Fragmentation Mix) |
| 80 °C | 5 min | |
| 42 °C | hold | (add TS-RT Mix to tube) |
| 42 °C | 30 min | |
| 70 °C | 10 min | |
| 4 °C | hold | |

##### *Program 1a: TS-RT Only*

This program is only used with LCM tissue on an Arcturus cap.

| | | |
|---|---|---|
| 42 °C | hold | (add LCM TS-RT Mix to lysate in tube) |
| 42 °C | 30 min | |
| 70 °C | 10 min | |
| 4 °C | hold | |

##### *Program 2: PCR*

| | | |
|---|---|---|
| 98 °C | hold | (add tube containing TS-RT product and PCR reagents) |
| 98 °C | 45 s | |
| cycles: | | |
| 98 °C | 15 s | |
| 60 °C | 30 s | |
| 72 °C | 10 s | |
| 72 °C | 60 s | |
| 4 °C | hold | |

#### Procedure

1. Place the RNase inhibitor and SMARTScribe reverse transcriptase on ice and thaw the frozen TS-RT reagents to room temperature in your pre-PCR workstation.
2. Start Program 1: Frag + TS-RT in a designated amplification-free thermal cycler to pre-warm it.
3. To 1 µL total RNA sample in a low-retention PCR tube or microplate well, add 4.0 µL Fragmentation Mix and mix by pipetting:

| | |
|---|---|
| 2.0 µL | SMARTScribe first-strand reaction buffer, 5X |
| 1.0 µL | nuclease-free dNTP mix, 10 mM each |
| 0.5 µL | 1S primer, 20 µM |
| 0.5 µL | $MgCl_2$, 80 mM |

4. When Program 1 has reached the 80 °C hold, place the tube in the thermal cycler and end the hold.
5. When the thermal cycler reaches the 42 °C hold, remove and briefly centrifuge the tube, then add 5.0 µL TS-RT Mix and mix by pipetting:

| | |
|---|---|
| 2.0 µL | TMG, 5 M |
| 1.0 µL | DTT, 20 mM |
| 0.5 µL | RNase inhibitor, 20X |
| 0.5 µL | 2S primer, 20 µM |
| 1.0 µL | SMARTScribe reverse transcriptase, 100 U/µL |

6. Immediately return the tube to the thermal cycler and end the 42 °C hold.
7. Thaw the frozen PCR reagents to room temperature.
8. Just before needed, start Program 2: PCR in a thermal cycler designated for PCR, using the appropriate number of cycles for your library (see "Choosing the number of PCR cycles").
9. Once Program 1 reaches the 4 °C hold, remove and briefly centrifuge the tube, then, still in your pre-PCR workstation, add 2.5 µL PCR primer mix with this library's unique combination of indexes, 2 µM each.
10. Add 12.5 µL HiFi HotStart ReadyMix, 2X, and mix by pipetting.
11. When Program 2 has reached the 98 °C hold, place the tube in the thermal cycler and end the hold.
12. Continue with the cleanup protocol for either Option 1 or Option 2.

##### *Option 1: Individual libraries*

1. When Program 2 reaches the 4 °C hold and the sample has cooled, briefly centrifuge the tube, add 55 µL diluted SPRI bead mix and mix very well:

| | |
|---|---|
| 25 µL | molecular biology–grade water |
| 30 µL | SPRI bead mix |

2. Incubate the tube 1 min at room temperature.
3. Place the tube on the magnet and wait for the beads to separate completely (~ 2 min on V&P 771LD).

4. Without disturbing the pellet, remove and discard all the supernatant. *Note:* low-retention pipet tips are not necessary for steps 4–6.
5. Still on the magnet, add 200 µL freshly prepared 80% ethanol to the tube and wait 30 s.
6. Remove and discard the supernatant. Use a smaller pipet to collect residual droplets.
7. Leave the tube open to air-dry 1 min. Do not overdry the pellet or it will be difficult to resuspend.
8. Remove the tube from the magnet, thoroughly resuspend the pellet in 10 µL DNA Storage Buffer and wait 30 s.
9. Return the tube to the magnet and allow the beads to separate completely.
10. Transfer the supernatant to a new tube. This is your sequencing-ready library.

##### *Option 2: Pre-SPRI pooling*

1. When Program 2 reaches the 4 °C hold and the samples have cooled, briefly centrifuge the tubes, then combine the samples into a single 1.5 mL ($N \leq 37$ samples) or 2.0 mL ($N \leq 50$) low-retention tube.
2. Add 15 × *N* µL SPRI bead mix and mix very well.
3. Incubate the tube 5 min at room temperature.
4. Place the tube on the magnet and wait for the beads to separate completely.
5. Without disturbing the pellet, remove and discard all the supernatant. *Note:* low-retention pipet tips are not necessary for steps 5 and 9–12.
6. Remove the tube from the magnet and resuspend the pellet in 96 µL Re-SPRI Mix (briefly centrifuge the tube if necessary to collect the pellet):

| | |
|---|---|
| 60 µL | molecular biology–grade water |
| 36 µL | SPRI bead mix or bead binding buffer |

7. Incubate the tube 5 min at room temperature.
8. Place the tube on the magnet and wait for the beads to separate completely.
9. Without disturbing the pellet, remove and discard all the supernatant.
10. Still on the magnet, add 1 mL freshly prepared 80% ethanol to the tube and wait 30 s.
11. Remove and discard the supernatant, then repeat the wash (step 10).
12. Remove and discard all the remaining supernatant. Use a smaller pipet to collect residual droplets.
13. Leave the tube open to air-dry 2 min. Do not overdry the pellet or it will be difficult to resuspend.
14. Remove the tube from the magnet, thoroughly resuspend the pellet in 10 µL DNA Storage Buffer (briefly centrifuge the tube if necessary to collect the pellet) and wait 30 s.
15. Return the tube to the magnet and allow the beads to separate completely.
16. Transfer the supernatant to a new tube. This is your sequencing-ready library pool.

#### Quick reference

"Quick reference: Smart-3SEQ from isolated RNA or cells" and the subsequent versions for Arcturus LCM are concise reminders for users who are familiar with the protocol.

#### Safe stopping points

The reaction mixture can be stored at 4 °C overnight after Program 1 or Program 2.

#### Using damaged RNA

If your RNA sample is already very fragmented (e.g. from FFPE tissue), use an 0.7X SPRI ratio instead of 0.6X, i.e. use 35 µL of SPRI bead mix to prepare the dilute SPRI bead mix and then use 60 µL of that (individual library option), or 17.5 × *N* µL followed by 60 µL water + 42 µL SPRI mix (pre-SPRI pooling option). Expect a narrow distribution of smaller library molecules, and a lower percentage of alignable reads, which align very close to the end of each transcript.

#### Starting from cells instead of isolated RNA

For step 3, suspend cell(s) in a 5 µL solution comprising the normal 4 µL Fragmentation Mix + 1 µL 0.25% Triton X-100 and proceed normally. The ideal number of PCR cycles may match the recommendations for the equivalent amount of RNA (1 mammalian cell ≈ 10 pg RNA) but should probably be recalibrated.

#### Using Arcturus laser-capture microdissection

We prefer membrane slides for easier LCM, using the UV laser to cut the slide membrane. Glass membrane slides adhere to frozen tissue sections better than framed membrane slides in our experience. There does not seem to be much advantage to the framed membrane slides even for FFPE tissue.

The protocol is easier and much less time-consuming with MicroCaps since they can go directly into the same 0.2 mL PCR tubes used in the rest of the protocol, but they require doubling the reaction volumes to cover the surface of the cap. This may increase the relative amount of PCR primer dimers with very small tissue samples, so it is also possible to use HS caps with the normal volumes. Use MicroCaps if possible or HS Caps if necessary. To use an HS cap, reduce all volumes in the special protocol below by half, and use the original volumes after returning to the main protocol. Carefully pipet 5 µL lysis mix directly onto the center of the HS cap and avoid letting the droplet touch the 0.5 mL tube when you put it on. If it does, collect the entire droplet in the tube by brief centrifugation and try to put it back on the center of the cap. After lysis, briefly centrifuge the tube + cap to collect the lysate, then transfer it to a low-retention 0.2 mL PCR tube and continue.

#### Starting from fresh-frozen tissue on an Arcturus LCM MicroCap

Even with good handling we still observe some degradation of RNA from fresh-frozen tissue, probably during the LCM procedures. Minimize the time the sample spends at room temperature.

1. Pre-warm an incubator at 60 °C with the metal CapSure incubation block inside. *Note:* It may help to place an open container of water in the incubator for humidity.
2. Aliquot 10 µL LCM Lysis Mix into an 0.2 mL low-retention PCR tube:

| | |
|---|---|
| 4 µL | SMARTScribe first-strand reaction buffer, 5X |
| 2 µL | nuclease-free dNTP mix, 10 mM each |
| 1 µL | 1S primer, 20 µM |
| 1 µL | $MgCl_2$, 80 mM |
| 1 µL | Triton X-100, 0.5% (v/v) |
| 1 µL | RNase inhibitor, 20X |

3. Tightly seal the MicroCap into the tube, then flick it upside-down to cover the cap with the liquid.
4. Place the upside-down cap and tube in the pre-warmed incubation block.
5. Incubate 20 min at 60 °C.
6. Pre-warm a designated amplification-free thermal cycler with Program 1a: TS-RT Only.
7. When the incubation is complete, briefly centrifuge the tube + cap to collect the lysate, then remove the cap. You can inspect it under a microscope later to verify complete lysis.
8. Add 10 µL LCM TS-RT Mix to the tube and mix by pipetting:

| | |
|---|---|
| 4 µL | TMG, 5 M |
| 2 µL | DTT, 20 mM |
| 1 µL | molecular biology–grade water |
| 1 µL | 2S primer, 20 µM |
| 2 µL | SMARTScribe reverse transcriptase, 100 U/µL |

9. Place the tube in the thermal cycler and end the 42 °C hold. The lid of the tube might not seal as well after the MicroCap has stretched it out, but the lid of the thermal cycler will hold it closed.
10. Continue with the normal protocol from step 7 (thaw the PCR reagents). Follow the special instructions for “Using damaged RNA”. Double the volumes of the PCR primers and PCR master mix. In the SPRI cleanup, simply use 35 µL SPRI bead mix with no water (Option 1) or 35 × *N* µL (Option 2, first cleanup; keep the second cleanup the same). See “Quick reference: Smart-3SEQ starting from fresh-frozen tissue on Arcturus LCM” for a concise but complete protocol with all the steps and volumes.

#### Starting from FFPE tissue on an Arcturus LCM MicroCap

This version of the protocol is optimized for formalin-fixed, paraffin-embedded tissue sections, which require longer lysis time and have degraded RNA even before you start the protocol.

1. Pre-warm an incubator at 60 °C with the metal CapSure incubation block inside. *Note:* It may help to place an open container of water in the incubator for humidity.
2. Aliquot <u>10 µL FFPE LCM Lysis Mix</u> into an 0.2 mL low-retention PCR tube:

| | |
|---|---|
| 4 µL | TMG, 5 M |
| 2 µL | nuclease-free dNTP mix, 10 mM each |
| 1 µL | 1S primer, 20 µM |
| 1 µL | Triton X-100, 0.5% (v/v) |
| 2 µL | Proteinase K, diluted to 0.125 µg/µL in molecular biology–grade water |

3. Tightly seal the MicroCap into the tube, then flick it upside-down to cover the cap with the liquid.
4. Place the upside-down cap and tube in the pre-warmed incubation block.
5. Incubate <u>60 min at 60 °C</u>.
6. Pre-warm a designated amplification-free thermal cycler with <u>Program 1a: TS-RT Only</u>.
7. When the incubation is complete, briefly centrifuge the tube + cap to collect the lysate, then remove the cap. You can inspect it under a microscope later to verify complete lysis.
8. Add <u>10 µL FFPE LCM TS-RT Mix</u> to the lysate and mix by pipetting:

| | |
|---|---|
| 4.0 µL | SMARTScribe first-strand reaction buffer, 5X |
| 2.0 µL | DTT, 20 mM |
| 1.0 µL | RNase inhibitor, 20X |
| 0.4 µL | 2S primer, 50 µM |
| 0.4 µL | $MgCl_2$, 200 mM |
| 0.2 µL | Proteinase K inhibitor, 5 mM |
| 2.0 µL | SMARTScribe reverse transcriptase, 100 U/µL |

9. Place the tube in the thermal cycler and end the 42 °C hold. The lid of the tube might not seal as well after the MicroCap has stretched it out, but the lid of the thermal cycler will hold it closed.
10. Continue with the normal protocol from step 7 (thaw the PCR reagents). Follow the special instructions for “Using damaged RNA”. Double the volumes of the PCR primers and PCR master mix. In the SPRI cleanup, simply use 35 µL SPRI bead mix with no water (Option 1) or 35 × *N* µL (Option 2, first cleanup; keep the second cleanup the same). See “Quick reference: Smart-3SEQ starting from fresh-frozen tissue on Arcturus LCM” for a concise but complete protocol with all the steps and volumes.

#### Quantification and validation of the libraries

The final yield should be 10 µL of amplified library at 5 to 50 nM, with most of the fragments between 200 and 600 bp. The size distribution can be verified by running the library undiluted on an Agilent Bioanalyzer 2100 DNA High Sensitivity chip or equivalent, which will also give a rough reading of the concentration. The electropherogram may show evidence of overamplification: a secondary bump or especially wide smear of molecules that migrate more slowly than the rest, because they comprise complementary annealed adapters and noncomplementary, unannealable inserts. These libraries can still be sequenced, but the Bioanalyzer will report inaccurate molarities, and it is ideal to recalibrate the PCR cycles to the maximum number that does not produce this artifact. A small bump at about 85 bp is a normal byproduct of the TS-RT reaction and should not affect sequencing; a spike at about 160 bp indicates adapter dimers that will reduce the number of usable sequence reads, which are caused by too many PCR cycles.

To determine pooling volumes (if using Option 1: Individual libraries) and optimize cluster density, it is helpful to measure the concentration more precisely by qPCR, which measures sequenceable molecules rather than total DNA content. Kapa's Library Quantification Kits are well designed for this purpose, although they are based on SYBR Green I, which measures total DNA mass and therefore requires precise knowledge of the average fragment size to calculate molarity. Alternatively, use a dual-labeled hydrolysis probe to directly measure molarity instead of mass.

Current Illumina sequencing protocols require only 5 µL pooled library mix at 4 nM combined concentration and only sequence a fraction of this, but your service providers may request more; ask for their requirements.

#### Sequencing

A good target depth for the human transcriptome is 1–10 million reads. Long reads (> 50 nt) are not required as this method is for counting reads, not bases. Slightly longer reads may reduce alignment ambiguity, but there are diminishing returns; you can quantify the effect by truncating and reprocessing long reads. No custom sequencing primers are required, and standard Illumina indexing is used. Paired-end reads are not recommended, as read 2 will start with a long homopolymer and may become unreadable afterward. Likely due to the enrichment of poly(A) inserts, libraries from this method may produce many clusters that fail Illumina's chastity filter. ΦX174 spike-in is not required, but a 1% spike-in is useful for troubleshooting in case of instrument failure. Average base quality will be poor in bases 6–8 and will decrease more rapidly than usual across the length of the read, but poly(A) trimming corrects the latter issue. The three all-G cycles may cause a loss of focus on the HiSeq 4000 and NovaSeq; using the latest instrument software and/or spiking in more ΦX174 may resolve this problem but we have not tested it thoroughly.

#### Data processing

All reads will begin with a five-base unique molecular identifier (UMI) (Kivioja et al., Nat Methods 2011) followed by three G's, then the sense-oriented cDNA sequence starting 50–500 nt upstream of the source RNA's poly(A) tail, followed by a long stretch of A's. Special processing is required before alignment to extract the cDNA inserts and annotate the reads with their UMI sequences.

The processed reads can be aligned to a reference genome/transcriptome with any standard aligner. They will be sense-oriented (“`fr-secondstrand`” in the Tuxedo suite, “`-s 1`” in the Subread suite) and fall slightly upstream of each gene's transcription termination site, though a short, highly expressed gene may also have a spike precisely at its transcription start site from reading full-length transcripts. There may be artifacts from internal priming at A-rich regions within transcript bodies, but in our data we find that removing these reduces the quantitative accuracy.

Because this method only generates one sequenceable fragment per RNA molecule, normalization by transcript length is not required; transcript abundance is directly proportional to read count.

Suggested pipeline:

1. Demultiplex pooled libraries and trim adapter sequence with Illumina software.
2. Extract UMI sequence and G-overhang from FASTQ data, and remove A-tails, with `umi_homopolymer.py`.
3. Align reads to genome + transcriptome, and sort alignments by position, with STAR.
4. Count reads from multiple libraries with "`featureCounts -s 1`" (Subread suite).
5. Analyze read counts with DESeq2.

We are currently testing an optimized UMI-aware deduplication method. Do not use UMI-unaware deduplication; it is likely that different transcripts will sometimes produce identical 3′-end fragments by chance.

#### Oligonucleotide primer designs (Illumina-specific)

Oligonucleotide sequences © 2017 Illumina, Inc. All rights reserved. Derivative works created by Illumina customers are authorized for use with Illumina instruments and products only. All other uses are strictly prohibited.

All sequences are in the order 5′→3′. Resuspend lyophilized oligonucleotides in DNA storage buffer, and verify by UV spectrophotometry that the yields are within 10% of expectations. According to the measured concentrations, dilute 1S and 2S primers to 20 µM small aliquots. (The FFPE LCM protocol requires 50 µM 2S primer instead of 20 µM.) Dilute each combination of P5 and P7 PCR primers together into small aliquots containing a mix of both P5 and P7 at 2 µM each (e.g. combine equal volumes of P5 at 4 µM and P7 at 4 µM).

##### *Abbreviations (IDT codes)*

`/5Biosg/`: 5′ biotin
`V`: equimolar mix of A, C, G
`N`: equimolar mix of A, C, G, T
`rG`: riboguanosine (all other nucleosides are deoxy)
`*`: phosphorothioate backbone

##### *Template-switching reverse transcription primers*

Use RNase-free HPLC purification for all. This reduces the complexity of the random bases, but that is less important than the purity of full-length molecules. The 2S primer may need to be ordered as an "RNA oligo" (e.g. IDT requires this) since it contains some ribonucleotides.

First strand (1S) primer: `/5Biosg/GT GAC TGG AGT TCA GAC GTG TGC TCT TCC GAT CTT TTT TTT TTT TTT TTT TTT TTT TTT TTT TV`

Second strand (2S) primer: `/5Biosg/CT ACA CGA CGC TCT TCC GAT CTN NNN NrGrG rG`

##### *Single-indexing PCR primers*

These primers add a single i7 index (6 nt) to each library. Use HPLC purification for all. Up to 47 libraries may be combined in one pool for sequencing (46 with Illumina's two-color chemistry on the NextSeq, MiniSeq, and NovaSeq).

P5 universal: `AAT GAT ACG GCG ACC ACC GAG ATC TAC ACT CTT TCC CTA CAC GAC GCT CTT CCG ATC* T`

P7 indexed: `CAA GCA GAA GAC GGC ATA CGA GAT [i7] GTG ACT GGA GTT CAG ACG TGT GCT CTT CCG ATC* T` where [i7] is one of the following index sequences:

| | | | | | | | |
|---|---|---|---|---|---|---|---|
| 1 | CGT GAT | 13 | TTG ACT | 25 | ATC AGT | 37* | ATT CCG |
| 2 | ACA TCG | 14 | GGA ACT | 26* | GCT CAT | 38* | AGC TAG |
| 3 | GCC TAA | 15 | TGA CAT | 27 | AGG AAT | 39* | GTA TAG |
| 4 | TGG TCA | 16 | GGA CGG | 28* | CTT TTG | 40* | TCT GAG |
| 5 | CAC TGT | 17* | CTC TAC | 29* | TAG TTG | 41*‡ | GTC GTC |
| 6 | ATT GGC | 18 | GCG GAC | 30* | CCG GTG | 42* | CGA TTA |
| 7 | GAT CTG | 19 | TTT CAC | 31*‡ | ATC GTG | 43* | GCT GTA |
| 8 | TCA AGT | 20 | GGC CAC | 32* | TGA GTG | 44* | ATT ATA |
| 9 | CTG ATC | 21 | CGA AAC | 33* | CGC CTG | 45* | GAA TGA |
| 10 | AAG CTA | 22 | CGT ACG | 34* | GCC ATG | 46* | TCG GGA |
| 11‡ | GTA GCC | 23† | CCA CTC | 35* | AAA ATG | 47* | CTT CGA |
| 12 | TAC AAG | 24* | GCT ACC | 36* | TGT TGG | 48* | TGC CGA |

* Indices 1-16, 18-23, 25, and 27 match the TruSeq LT system so you can use that setting for a sample sheet in the Illumina Experiment Manager or BaseSpace. The other indices require loading special files.
† Index 23 is G-rich in sequencing orientation and therefore produces low signal on the two-color chemistry of the NextSeq, MiniSeq, and NovaSeq; avoid using index 23 on these platforms.
‡ Index 41 differs from indices 11 and 31 by only 2 bases; avoid pooling 11+41 or 31+41.

*Important:* **These are the reverse complements of the index sequences that are read by the instrument.**

#### PCR primers for higher-plexity pooling

Using longer index sequences, it is possible to pool larger numbers of libraries in the same sequencing run. In particular, IDT and Illumina have created new index designs for 96- or 384-plex unique dual indexing (UDI), where each library has a unique P7 index as well as a unique P5 index. Smart-3SEQ is compatible with this scheme but the UDI adapters are normally sold in double-stranded form for ligation-based protocols; contact IDT to get a quote for ordering the UDI PCR primers.

The original index-hopping problem solved by UDIs does not appear to affect Smart-3SEQ as much as other methods. A less expensive alternative for high plexity is to use only single-indexing with one side of the UDI pairs, e.g. order 96 or 384 of only the P7 UDI PCR primers and combine them with the universal P5 primer as above.

However, in our data from low-quality libraries (FFPE LCM), we sometimes observe sequencing artifacts in the i7 index read but not in the i5 index read, so we get better results from an inverted scheme using P5 indexes and a universal P7 primer (as above but with no i7 sequence). The problem is that i5-only indexing requires unusual sequencing conditions. The NextSeq 500 requires a minimum of 2 cycles to be read from the i7 index (index read 1) before it will read the i5 index (index read 2), and the unwanted i7 cycles should be masked in `bcl2fastq`. This works well on the NextSeq but Illumina has advised us that this i5-only scheme will probably fail on the MiSeq; we have no information about other sequencers.

### Quick reference: Smart-3SEQ from isolated RNA or cells

#### *Fragmentation Mix, 4 µL + 1 µL RNA (or suspend cells in 5 µL)*

| | **RNA** | **Cells** |
|---|---|---|
| SMARTScribe buffer, 5X | 2.0 µL | 2.0 µL |
| dNTP mix, 10 mM each | 1.0 µL | 1.0 µL |
| 1S primer, 20 µM | 0.5 µL | 0.5 µL |
| $MgCl_2$, 80 mM | 0.5 µL | 0.5 µL |
| Triton X-100, 0.25% | | 1.0 µL |

#### *TS-RT Mix, 5 µL*

| | |
|---|---|
| TMG, 5 M | 2.0 µL |
| DTT, 20 mM | 1.0 µL |
| RNase inhibitor, 20X | 0.5 µL |
| 2S primer, 20 µM | 0.5 µL |
| SMARTScribe, 100 U/µL | 1.0 µL |

#### *PCR*

| | |
|---|---|
| PCR primer pair, 2 µM each | 2.5 µL |
| HiFi ReadyMix, 2X | 12.5 µL |

#### *Cleanup*

| | **Individual libraries** | | **Pre-SPRI pooling** | |
|---|---|---|---|---|
| | **Intact RNA (55 µL)** | **Degraded RNA (60 µL)** | **Intact RNA** | **Degraded RNA** |
| Water | 25 µL | 25 µL | | |
| SPRI bead mix | 30 µL | 35 µL | 15 × *N* µL | 17.5 × *N* µL |

Mix well and incubate 1 min.

Separate beads on magnet and discard supernatant.

*Pre-SPRI pooling option only:*

Resuspend in 96 µL Re-SPRI Mix (102 µL if RNA was degraded):

| | **Intact RNA** | **Degraded RNA** |
|---|---|---|
| Water | 60 µL | 60 µL |
| SPRI bead mix | 36 µL | 42 µL |

Incubate 1 min away from magnet.

Separate beads on magnet and discard supernatant.

Wash with fresh 80% ethanol 30 s.

Discard supernatant and air-dry 1 min.

Elute in 10 µL DNA Storage Buffer 30 s away from magnet.

Separate beads on magnet and transfer supernatant for storage.

### Quick reference: Smart-3SEQ starting from fresh-frozen tissue on Arcturus LCM

#### *LCM Lysis Mix (20 min @ 60 °C)*

| | MicroCap (10 µL) | HS Cap (5 µL) |
|---|---|---|
| SMARTScribe buffer, 5X | 4.0 µL | 2.0 µL |
| dNTP mix, 10 mM each | 2.0 µL | 1.0 µL |
| 1S primer, 20 µM | 1.0 µL | 0.5 µL |
| $MgCl_2$, 80 mM | 1.0 µL | 0.5 µL |
| Triton X-100, 0.5% | 1.0 µL | 0.5 µL |
| RNase inhibitor, 20X | 1.0 µL | 0.5 µL |

#### *LCM TS-RT Mix (Program 1a, TS-RT Only)*

| | MicroCap (10 µL) | HS Cap (5 µL) |
|---|---|---|
| TMG, 5 M | 4.0 µL | 2.0 µL |
| DTT, 20 mM | 2.0 µL | 1.0 µL |
| water | 1.0 µL | 0.5 µL |
| 2S primer, 20 µM | 1.0 µL | 0.5 µL |
| SMARTScribe, 100 U/µL | 2.0 µL | 1.0 µL |

#### *PCR*

| | MicroCap | HS Cap |
|---|---|---|
| PCR primer pair, 2 µM each | 5.0 µL | 2.5 µL |
| HiFi ReadyMix, 2X | 25.0 µL | 12.5 µL |

#### *Cleanup*

| | Individual libraries | | Pre-SPRI pooling | |
|---|---|---|---|---|
| | **MicroCap** | **HS Cap (60 µL)** | **MicroCap** | **HS Cap** |
| Water | | 25 µL | | |
| SPRI bead mix | 35 µL | 35 µL | 35 × *N* µL | 17.5 × *N* µL |

Mix well and incubate 1 min.

Separate beads on magnet and discard supernatant.

*Pre-SPRI pooling option only:*

Resuspend in 102 µL Re-SPRI Mix:

| | |
|---|---|
| Water | 60 µL |
| SPRI bead mix | 42 µL |

Incubate 1 min away from magnet.

Separate beads on magnet and discard supernatant.

Wash with fresh 80% ethanol 30 s.

Discard supernatant and air-dry 1 min.

Elute in 10 µL DNA Storage Buffer 30 s away from magnet.

Separate beads on magnet and transfer supernatant for storage.

### Quick reference: Smart-3SEQ starting from FFPE tissue on Arcturus LCM

#### *FFPE LCM Lysis Mix (60 min @ 60 °C)*

| | MicroCap (10 µL) | HS Cap (5 µL) |
|---|---|---|
| TMG, 5 M | 4.0 µL | 2.0 µL |
| dNTP mix, 10 mM each | 2.0 µL | 1.0 µL |
| 1S primer, 20 µM | 1.0 µL | 0.5 µL |
| Triton X-100, 0.5% | 1.0 µL | 0.5 µL |
| Proteinase K, 0.125 µg/µL | 2.0 µL | 1.0 µL |

#### *FFPE LCM TS-RT Mix (Program 1a, TS-RT Only)*

| | MicroCap (10 µL) | HS Cap (5 µL) |
|---|---|---|
| SMARTScribe buffer, 2X | 4.0 µL | 2.0 µL |
| DTT, 20 mM | 2.0 µL | 1.0 µL |
| RNase inhibitor, 20X | 1.0 µL | 0.5 µL |
| 2S primer, 50 µM | 0.4 µL | 0.2 µL |
| $MgCl_2$, 200 mM | 0.4 µL | 0.2 µL |
| Proteinase K inhibitor, 5 mM | 0.2 µL | 0.1 µL |
| SMARTScribe, 100 U/µL | 2.0 µL | 1.0 µL |

#### *PCR*

| | MicroCap | HS Cap |
|---|---|---|
| PCR primer pair, 2 µM each | 5.0 µL | 2.5 µL |
| HiFi ReadyMix, 2X | 25.0 µL | 12.5 µL |

#### *Cleanup*

| | Individual libraries | | Pre-SPRI pooling | |
|---|---|---|---|---|
| | MicroCap | HS Cap (60 µL) | MicroCap | HS Cap |
| Water | | 25 µL | | |
| SPRI bead mix | 35 µL | 35 µL | 35 × *N* µL | 17.5 × *N* µL |

Mix well and incubate 1 min.

Separate beads on magnet and discard supernatant.

*Pre-SPRI pooling option only:*

Resuspend in 102 µL Re-SPRI Mix:

| | |
|---|---|
| Water | 60 µL |
| SPRI bead mix | 42 µL |

Incubate 1 min away from magnet.

Separate beads on magnet and discard supernatant.

Wash with fresh 80% ethanol 30 s.

Discard supernatant and air-dry 1 min.

Elute in 10 µL DNA Storage Buffer 30 s away from magnet.

Separate beads on magnet and transfer supernatant for storage.

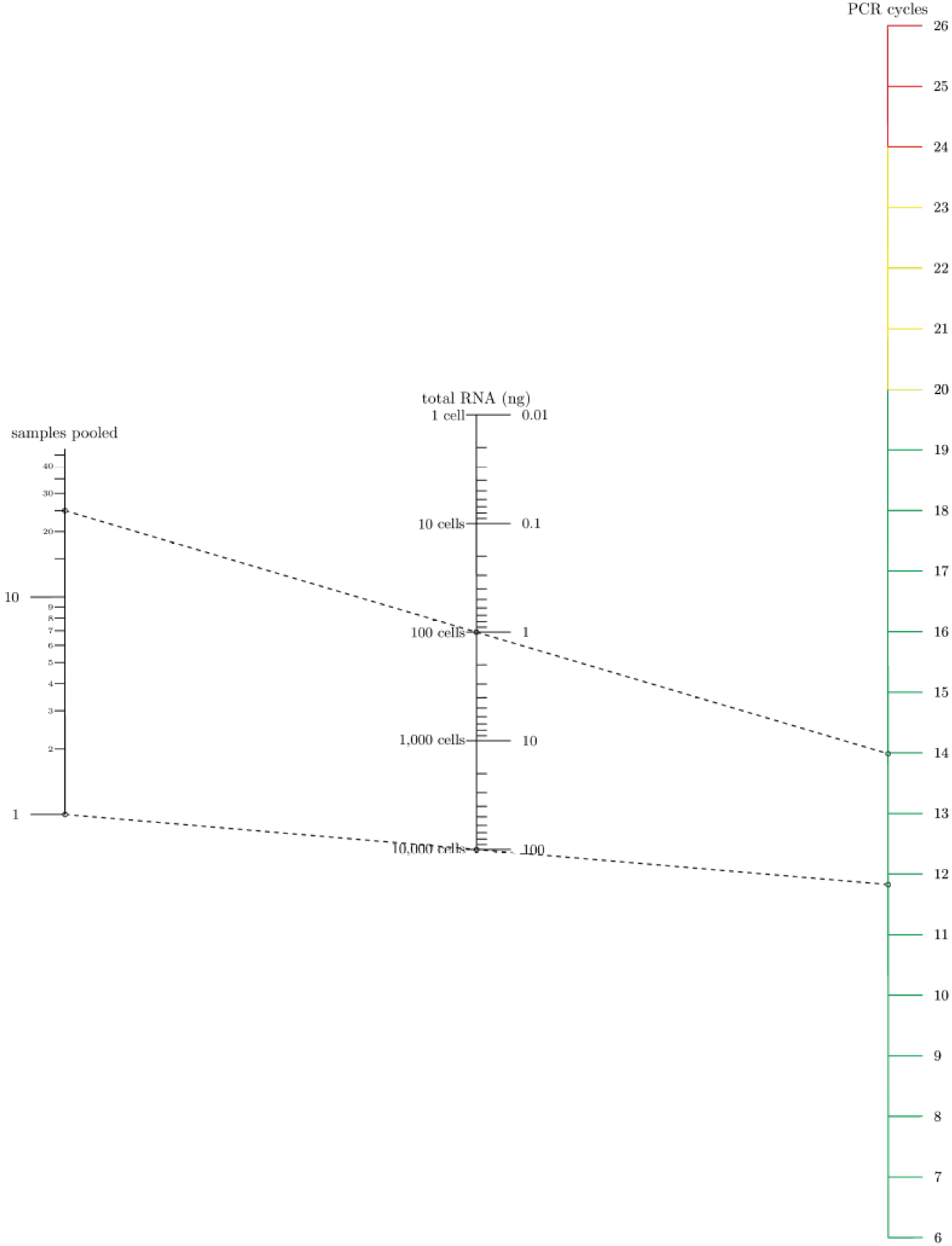

samples pooled
total RNA (ng)
PCR cycles
1
2
3
4
5
6
7
8
9
10
20
30
40
1 cell
10 cells
100 cells
1,000 cells
10,000 cells
0.01
0.1
1
10
100
