## Supplementary figures and images for "Gene-expression profiling of single cells from archival tissue with laser-capture microdissection and Smart-3SEQ"

### DCIS-1 40x.jpeg

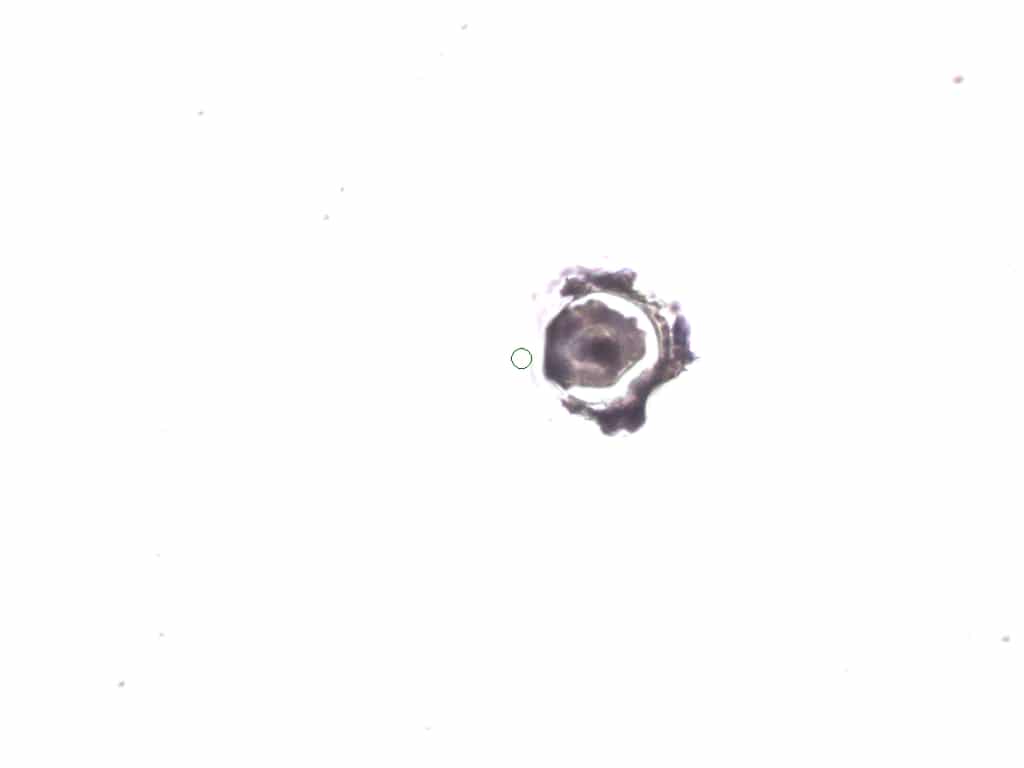

### DCIS-1 BEFORE.jpeg

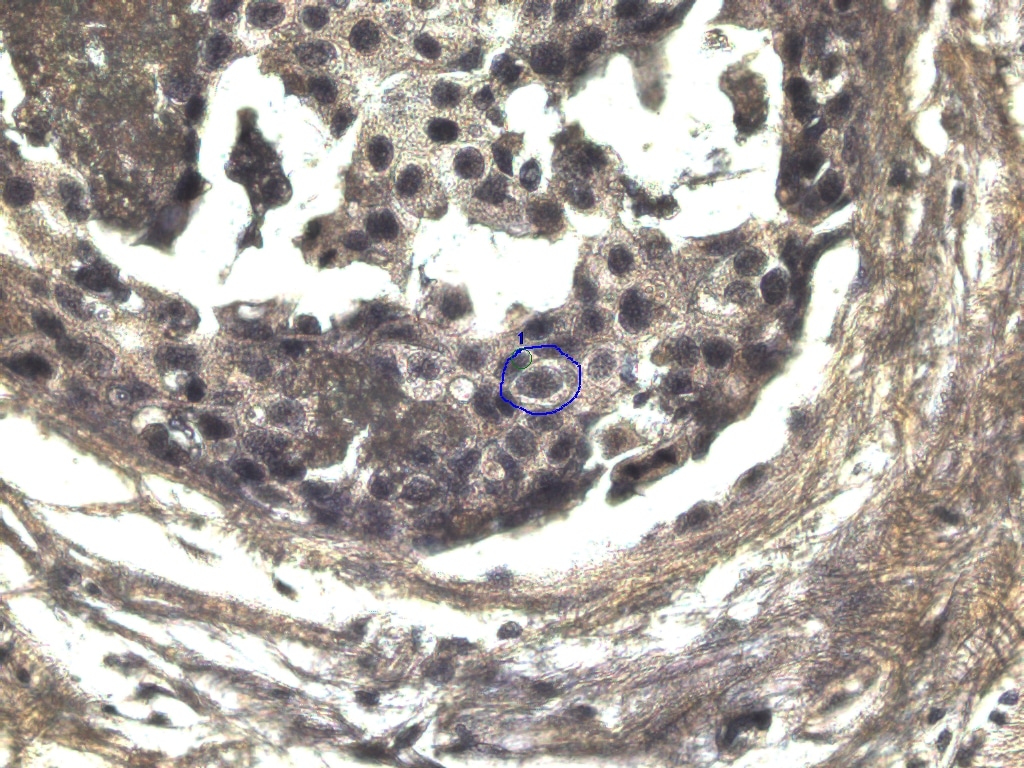

### DCIS-1AFTER40x.jpeg

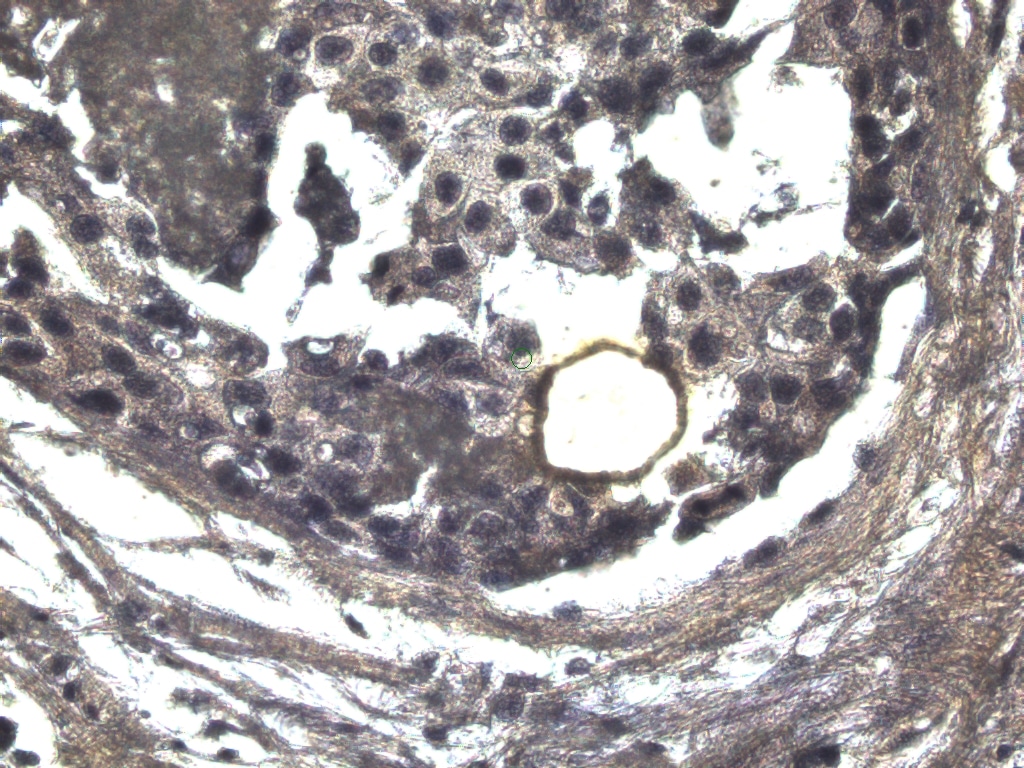

### DCIS-2 BEFORE.jpeg

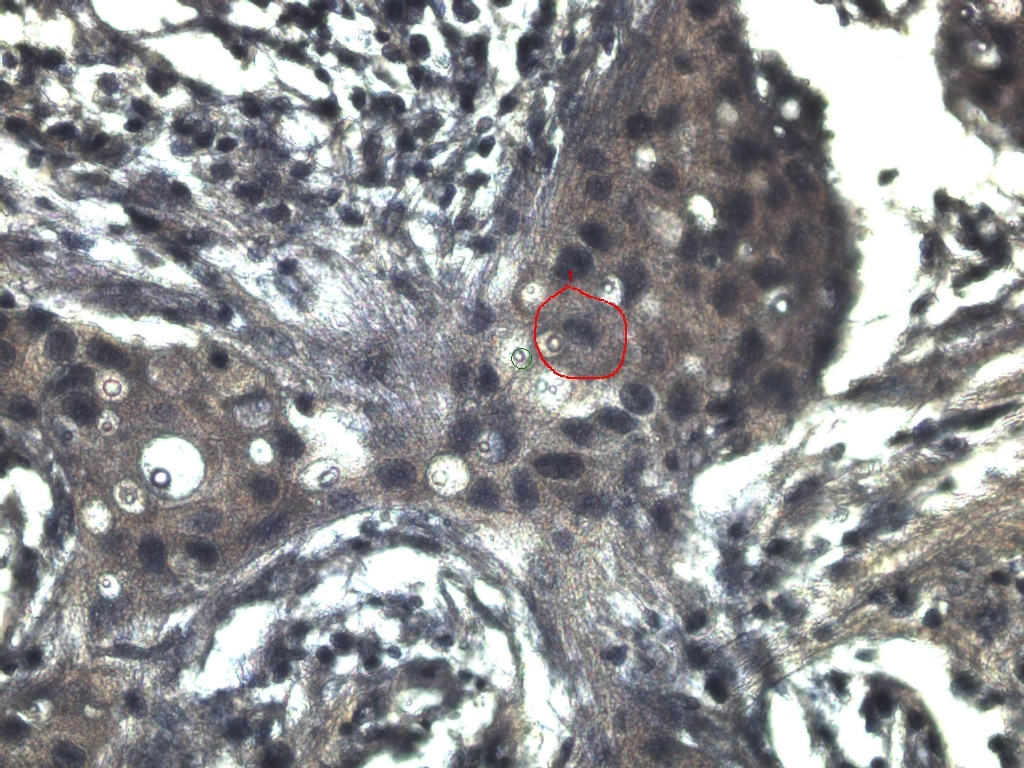

### DCIS-3 40x.jpeg

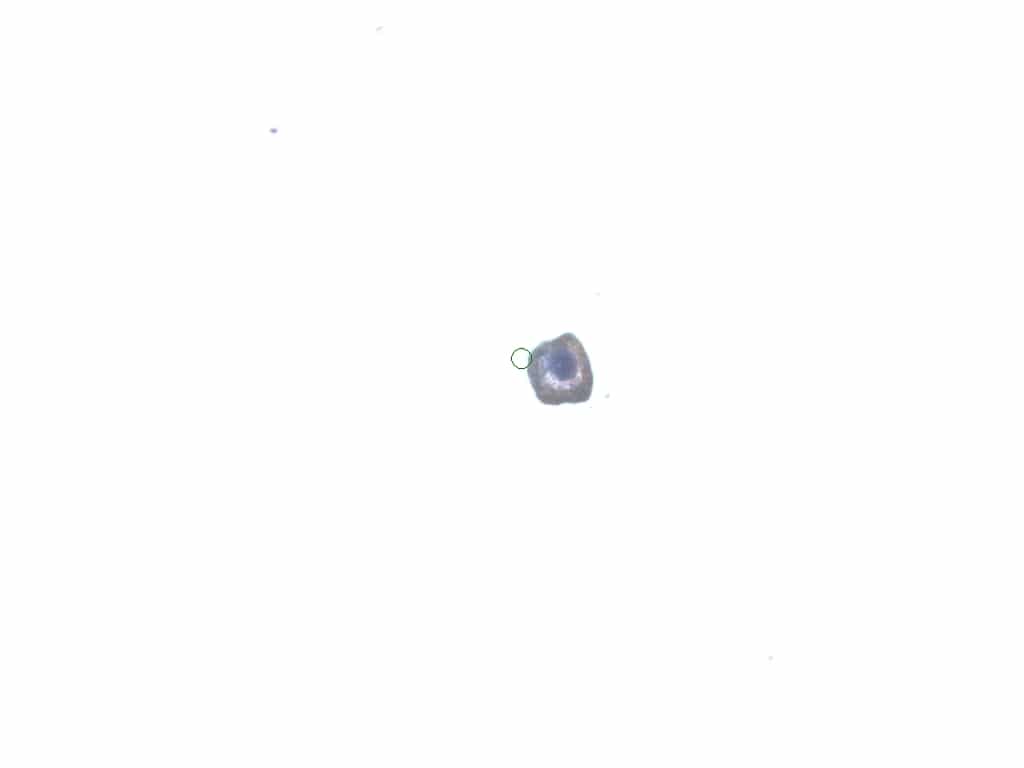

### DCIS-4 40x.jpeg

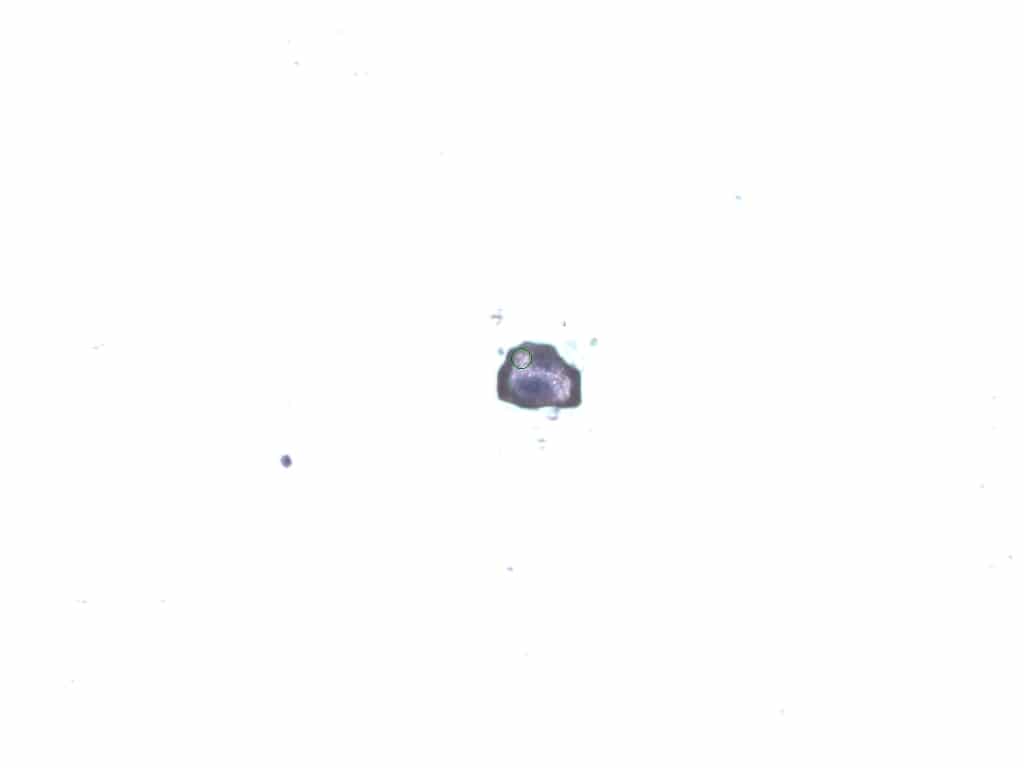

### DCIS-4 AFTER.jpeg

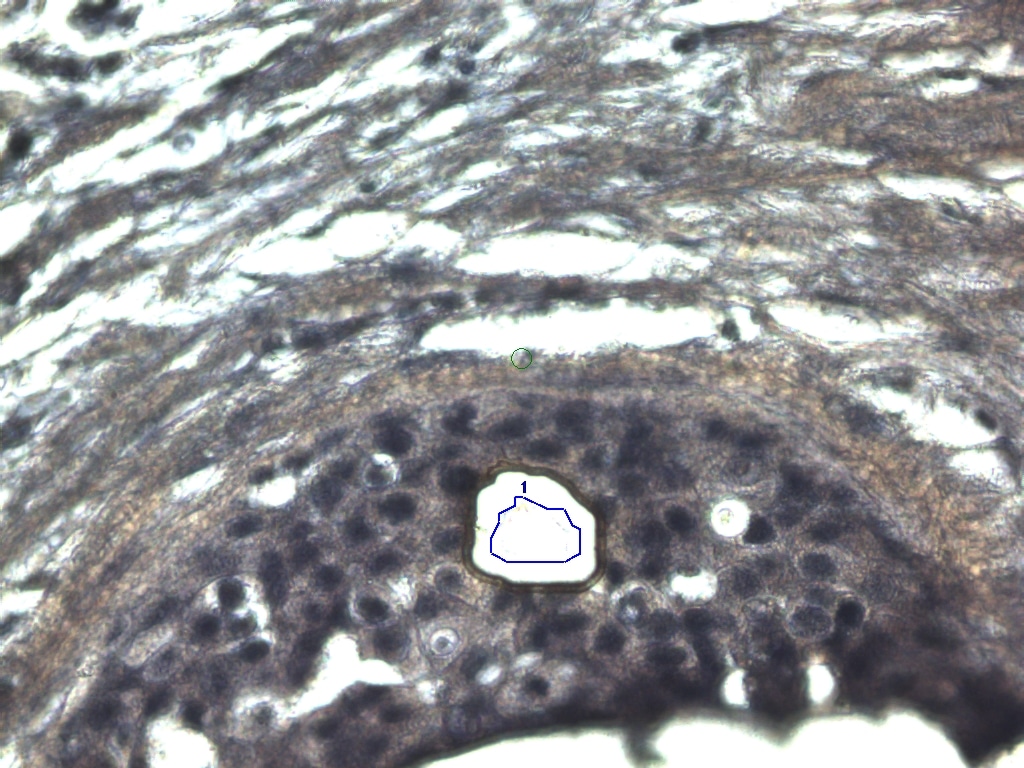

### DCIS-5 AFTER.jpeg

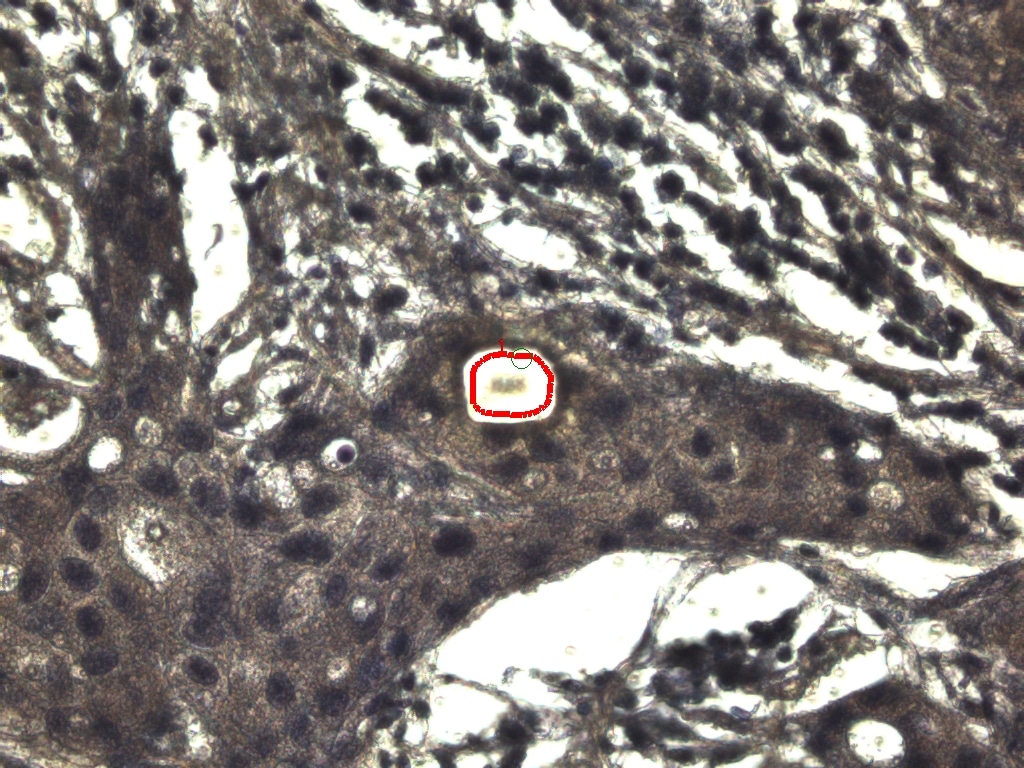

### DCIS-5 BEFORE.jpeg

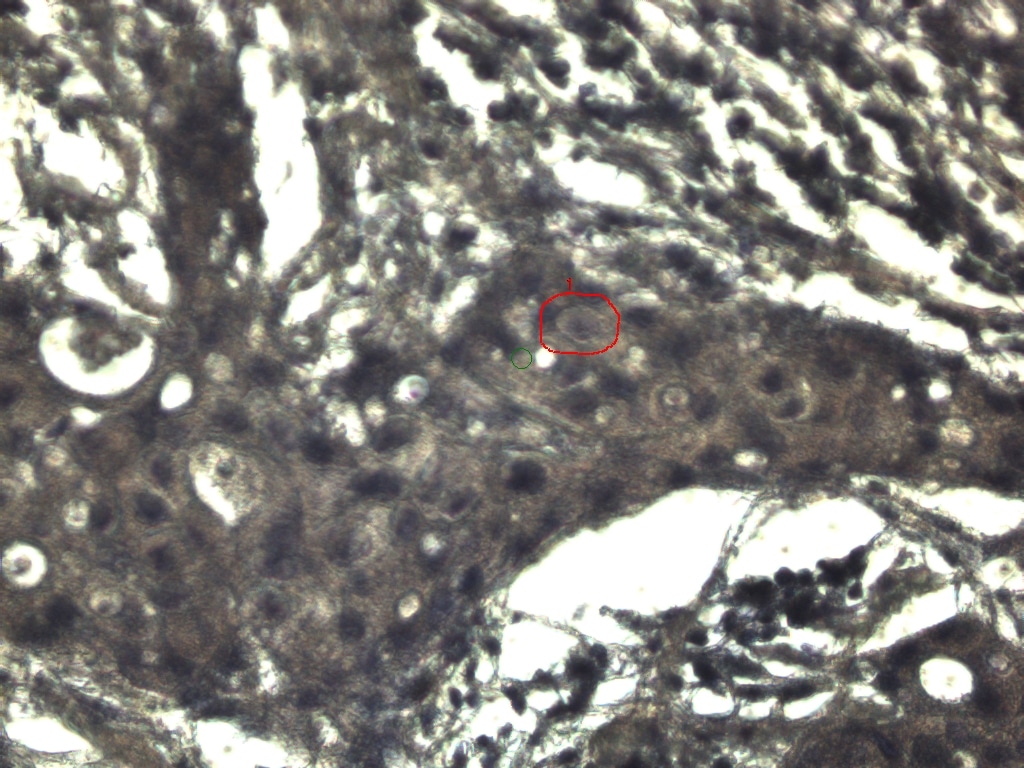

### DCIS-5.jpeg

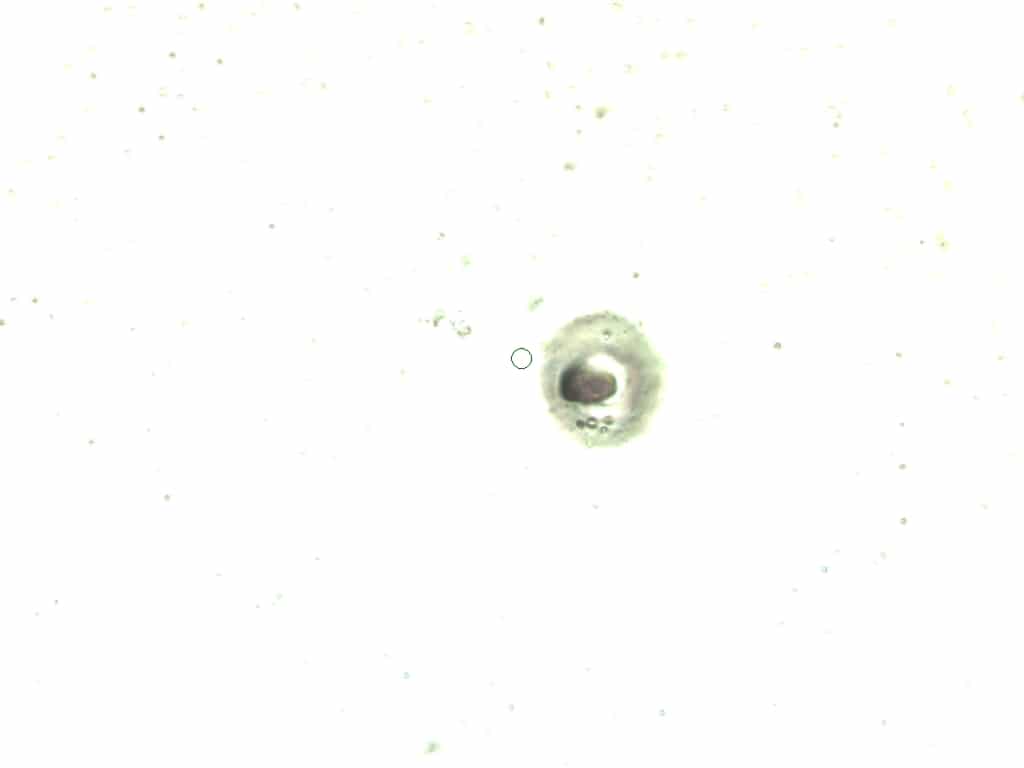

### DCIS-6 AFTER.jpeg

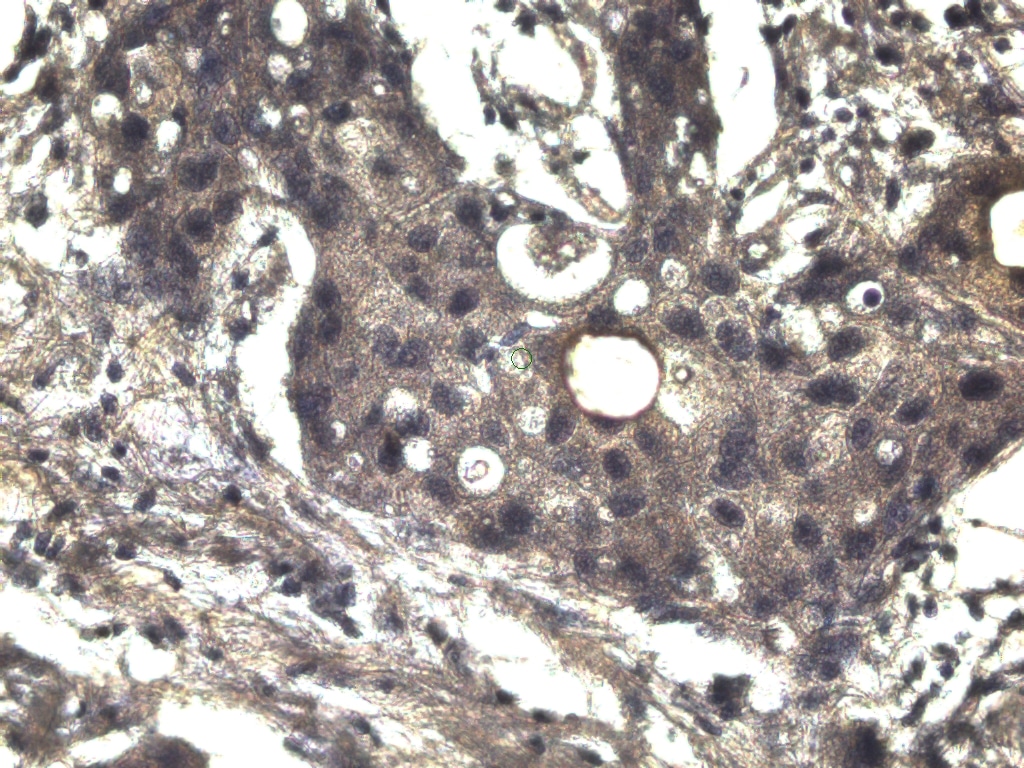

### DCIS-6 BEFORE.jpeg

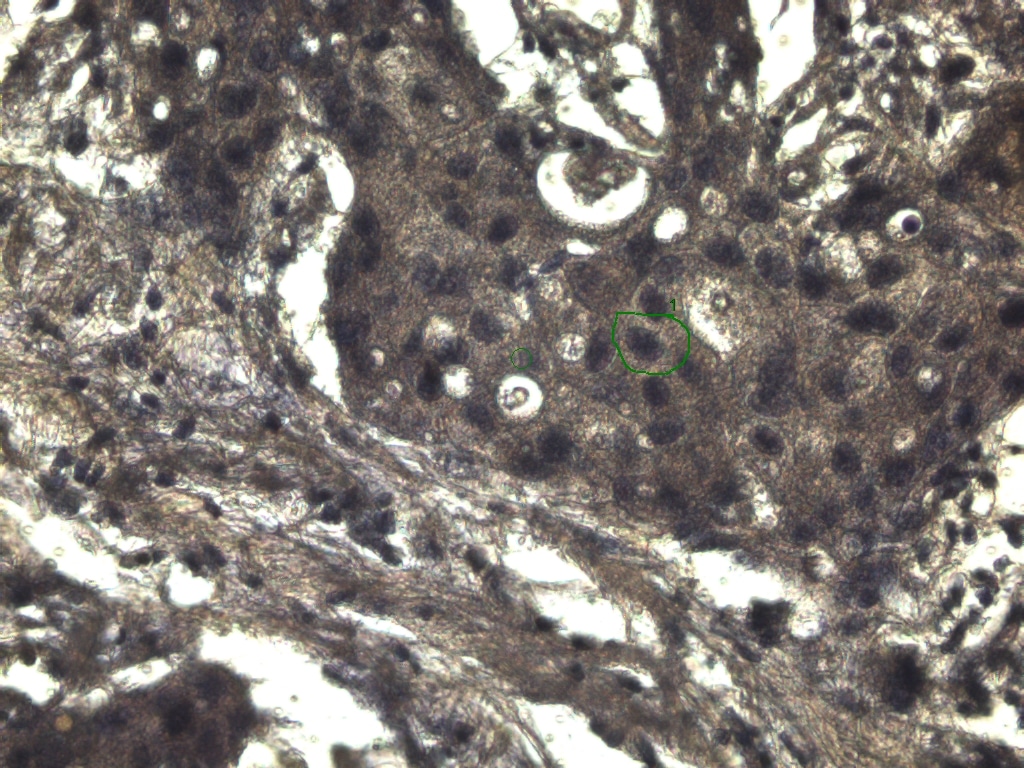

### DCIS-7 AFTER.jpeg

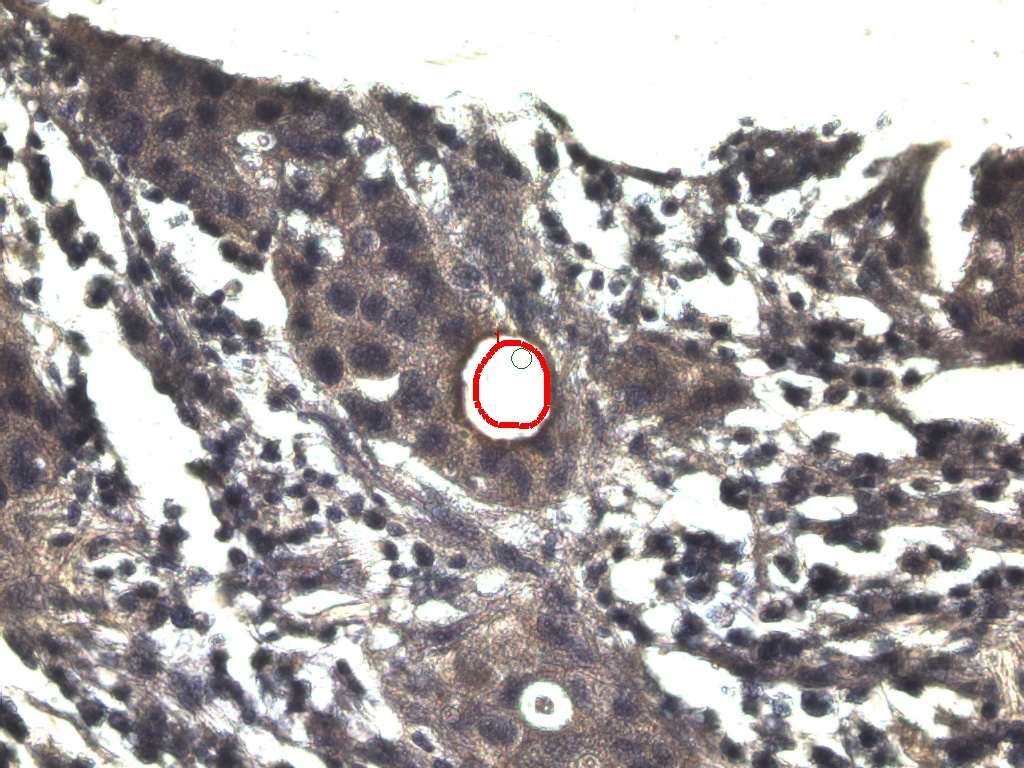

### DCIS-7 BEFORE.jpeg

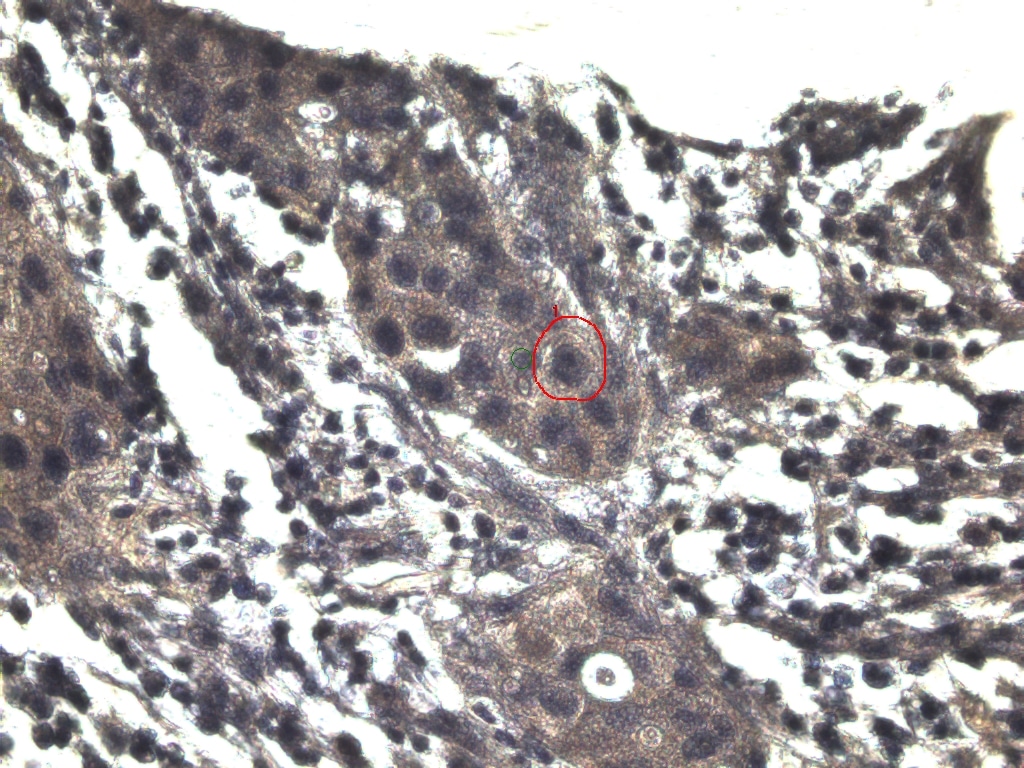

### DCIS-7.jpeg

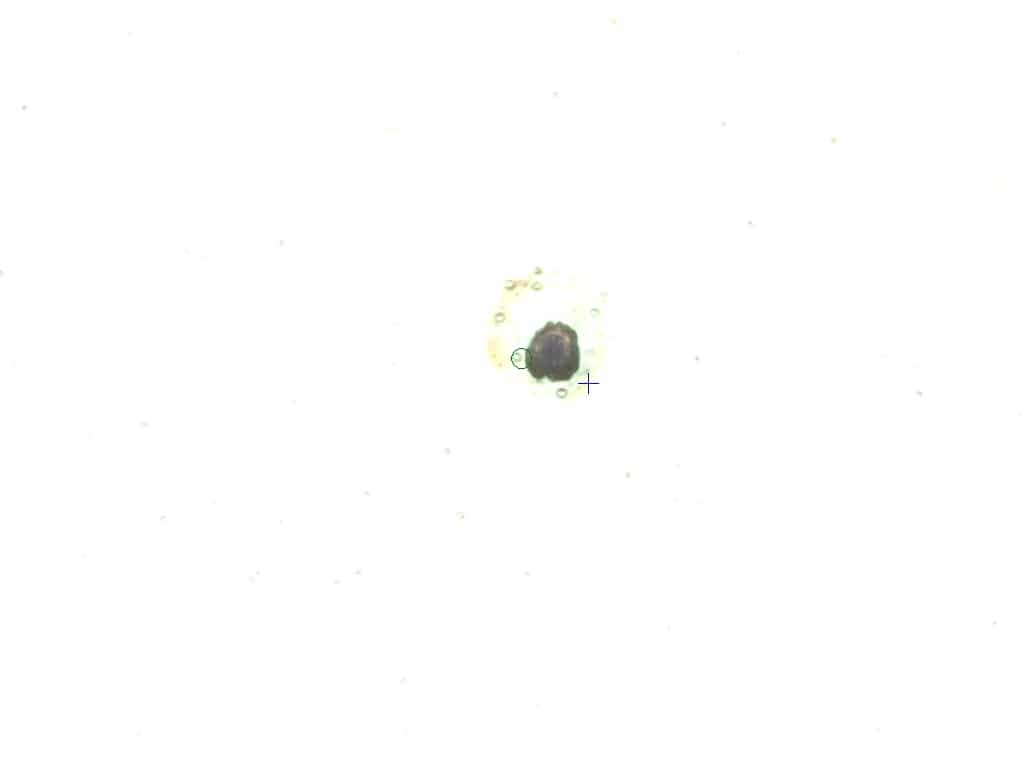

### DCIS-8 AFTER.jpeg

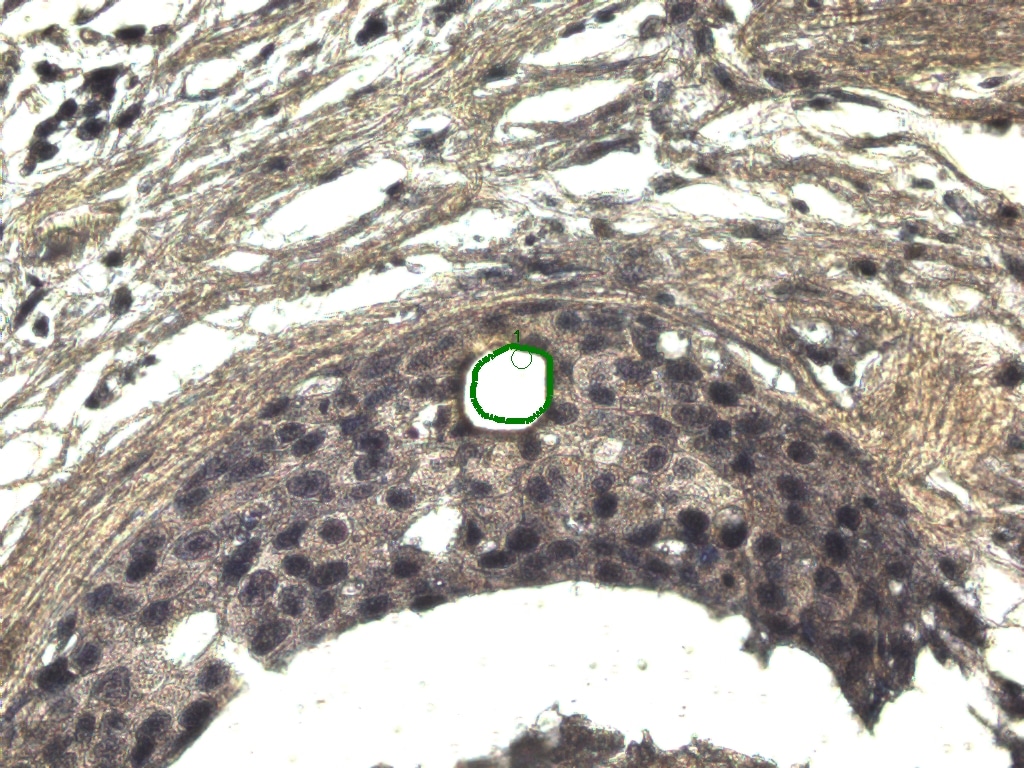

### DCIS-8 BEFORE.jpeg

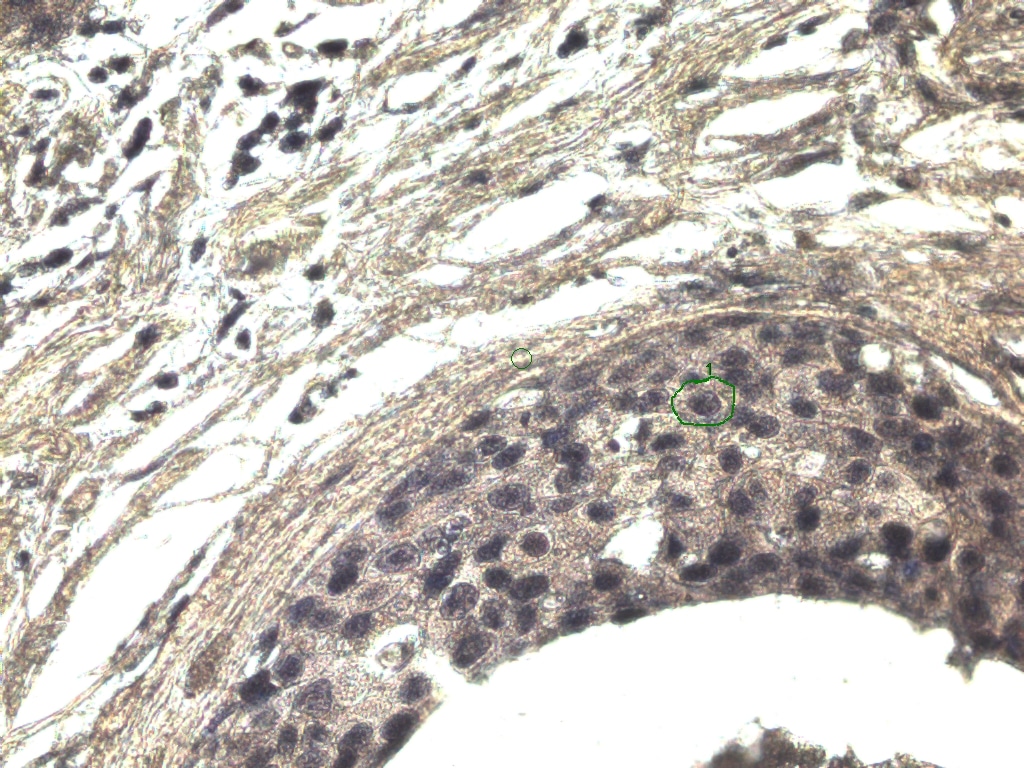

### DCIS-8.jpeg

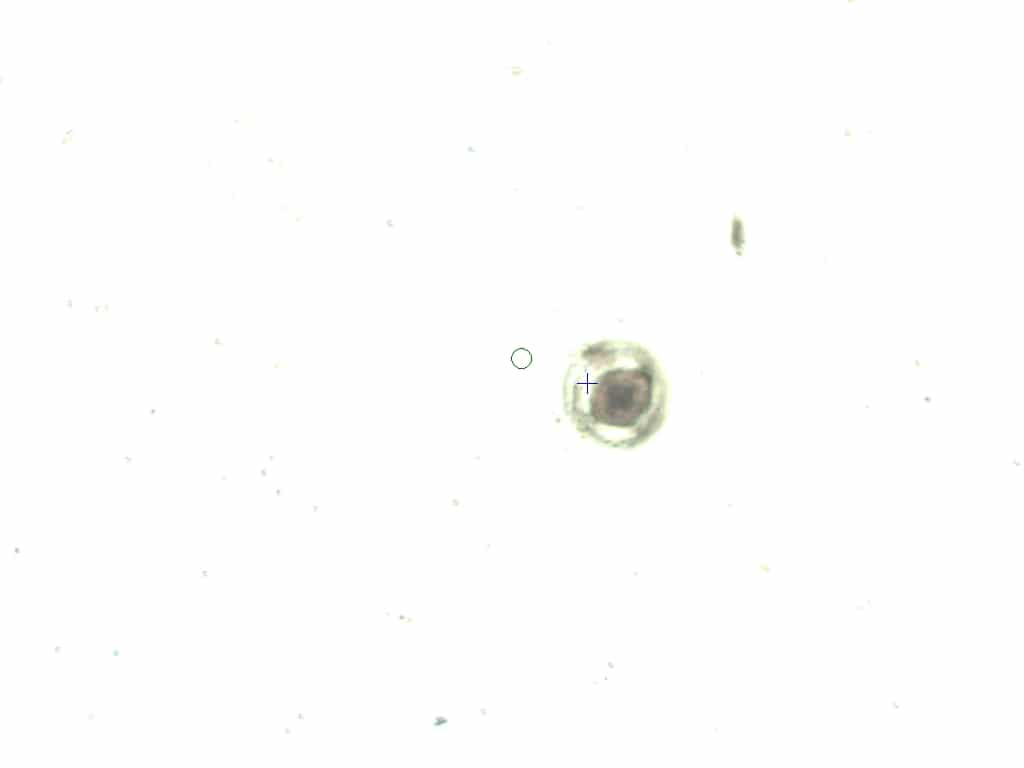

### DCIS-9 AFTER ABLATION.jpeg

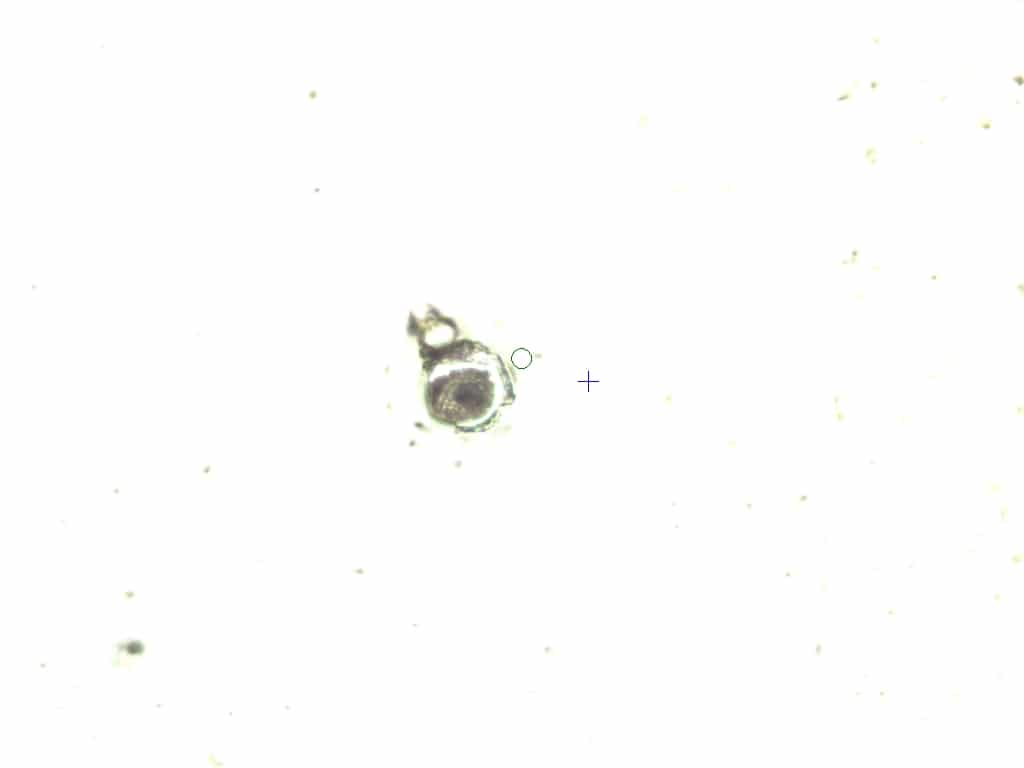

### DCIS-9 AFTER.jpeg

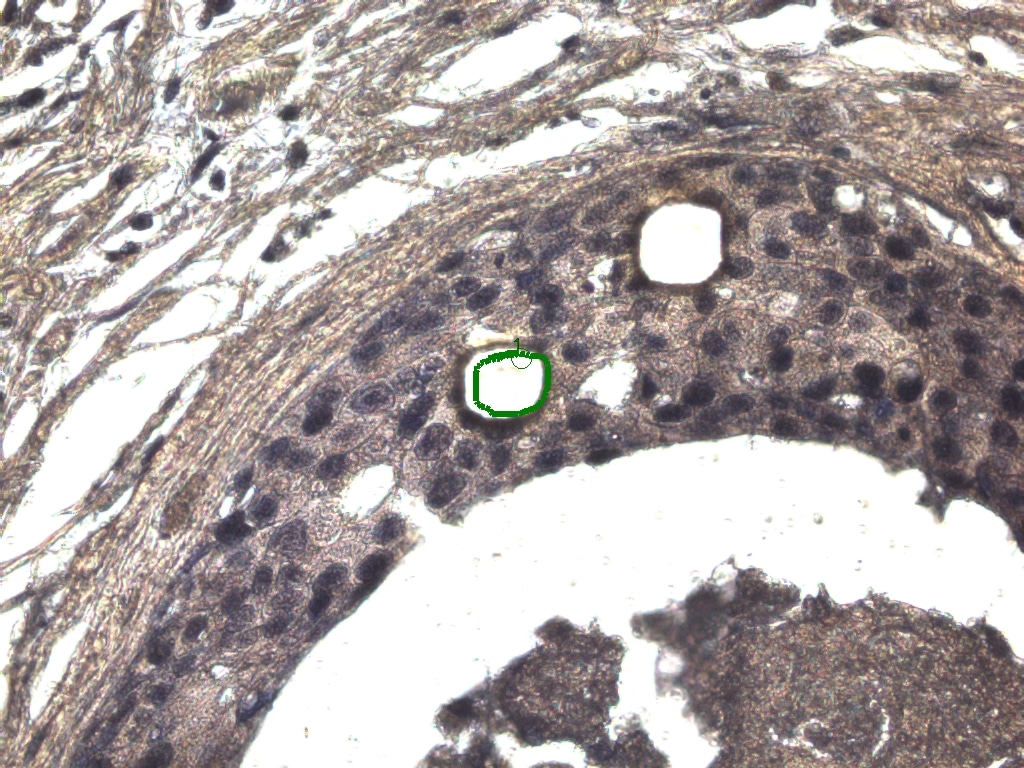

### DCIS-9.jpeg

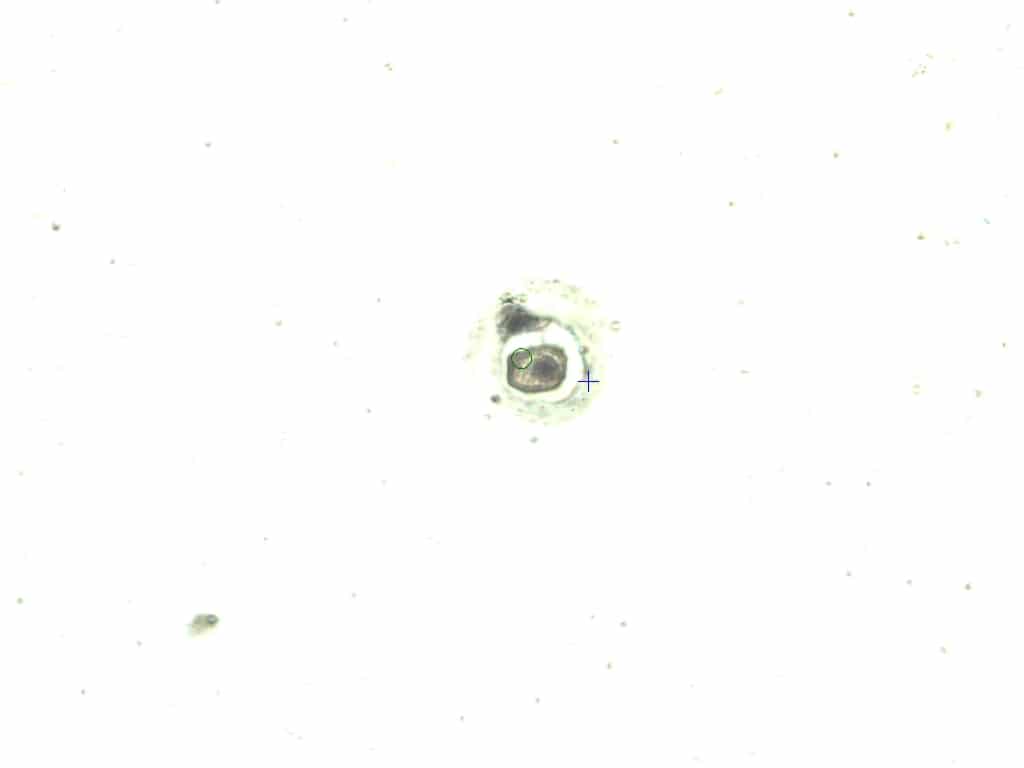

### DCIS-10 AFTER.jpeg

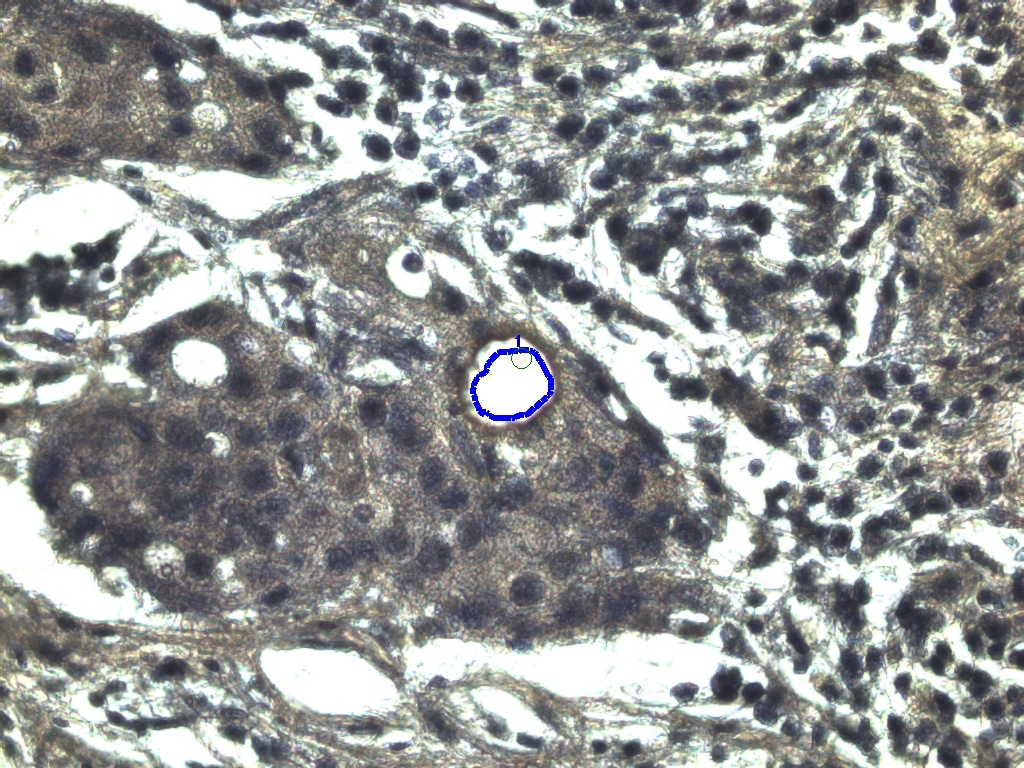

### DCIS-10 BEFORE.jpeg

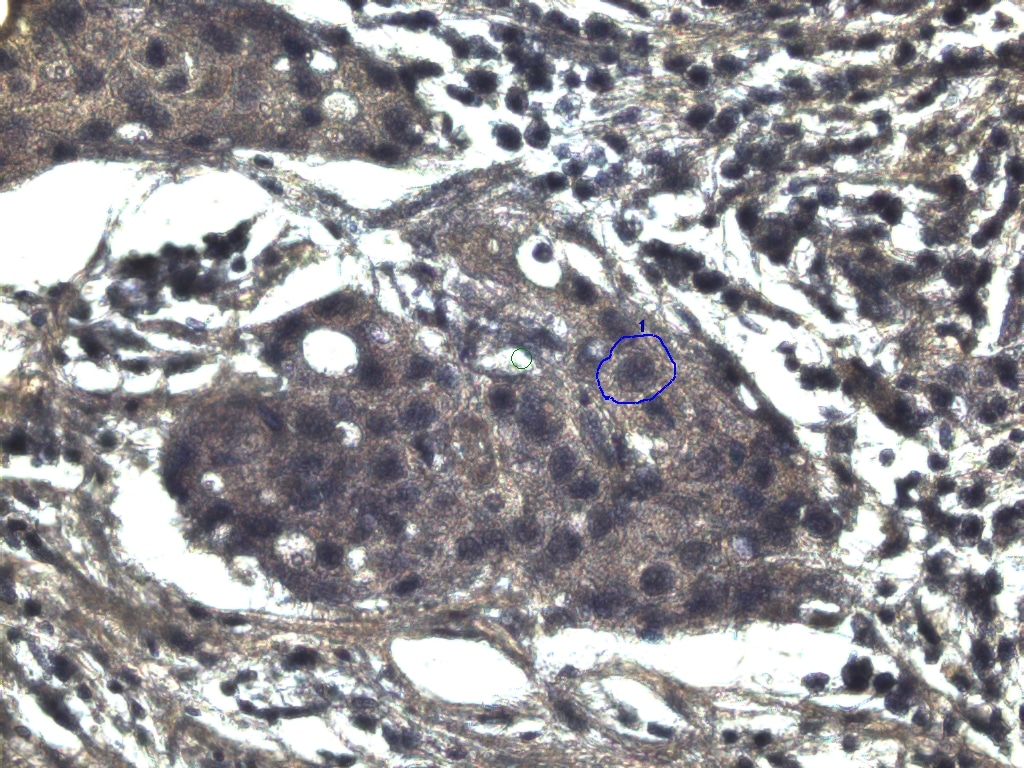

### DCIS-10.jpeg

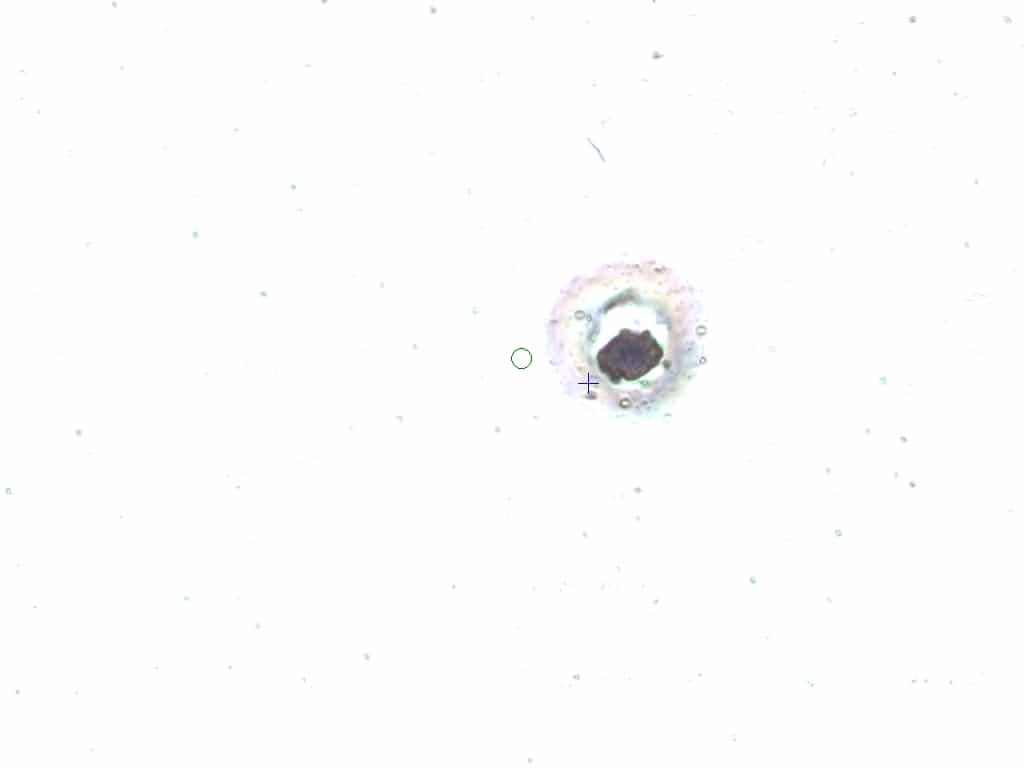

### MAC1 AFTER.jpeg

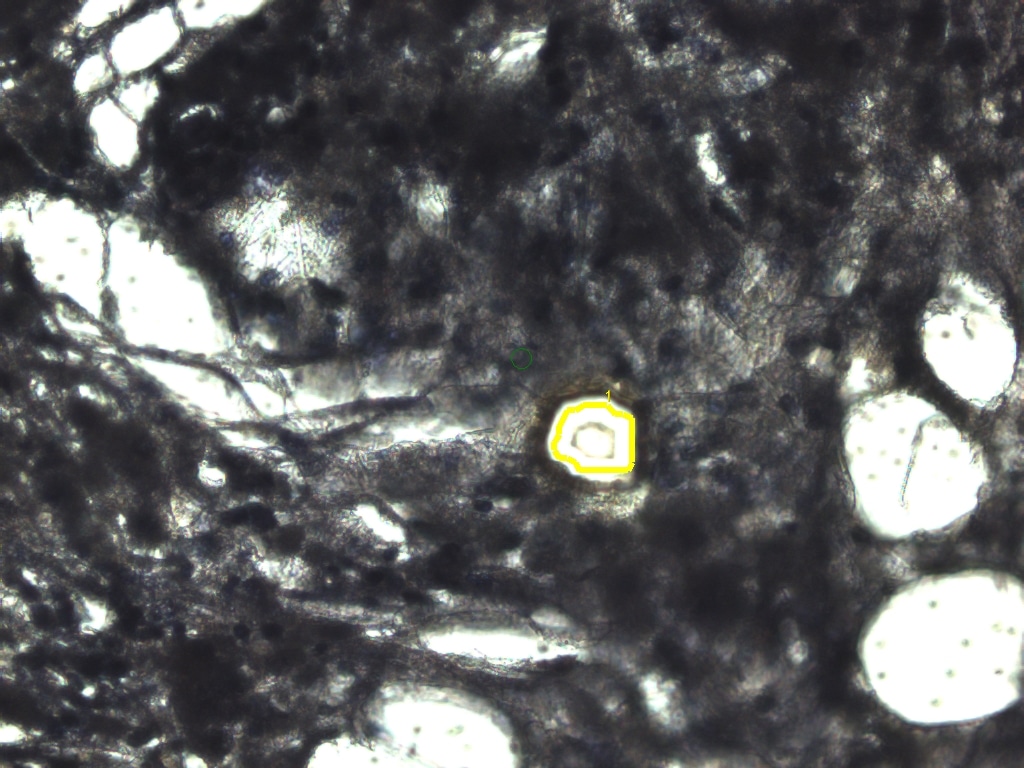

### MAC1 BEFORE.jpeg

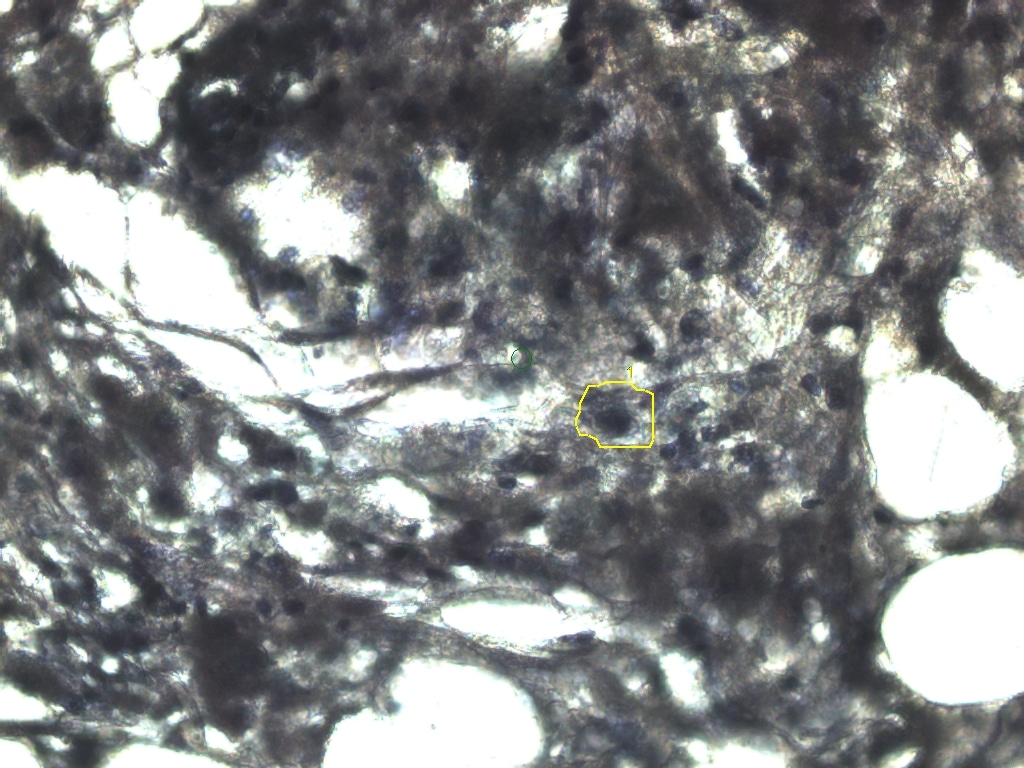

### MAC1.jpeg

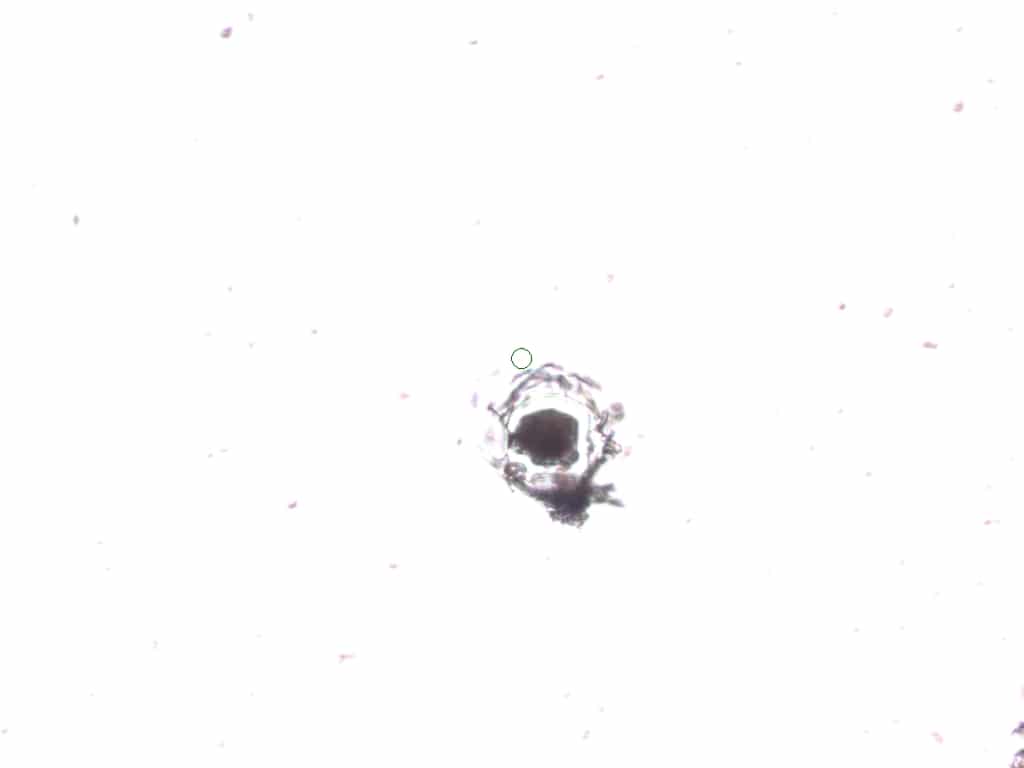

### MAC2 AFTER ABLATION.jpeg

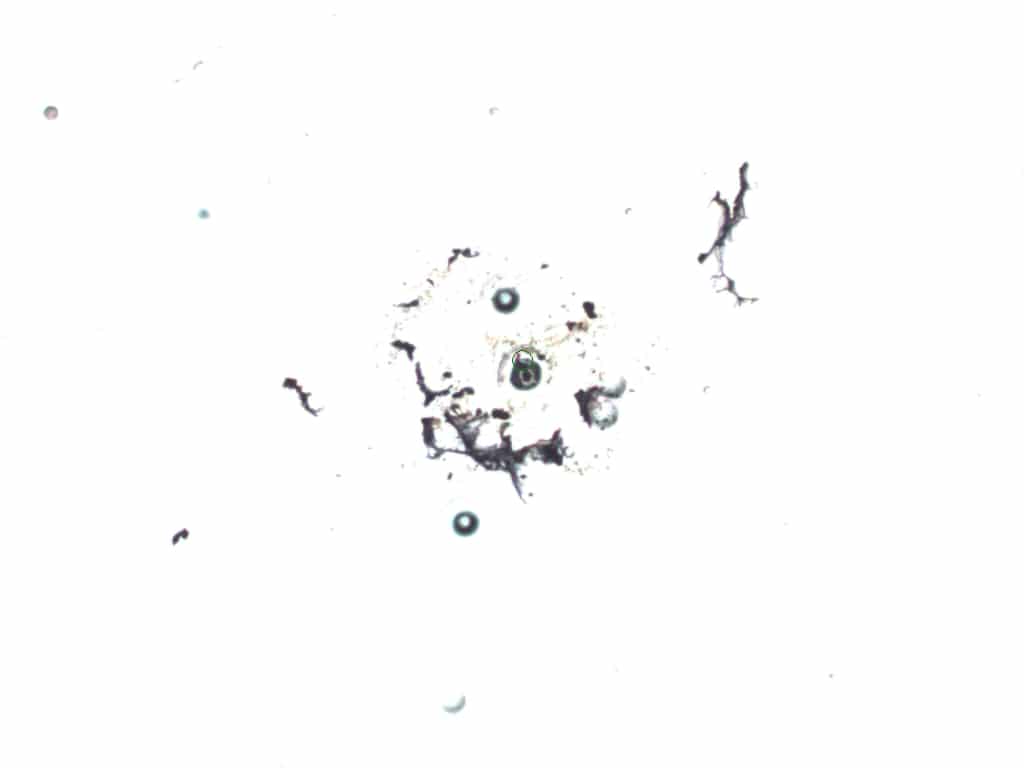

### MAC2 BEFORE.jpeg

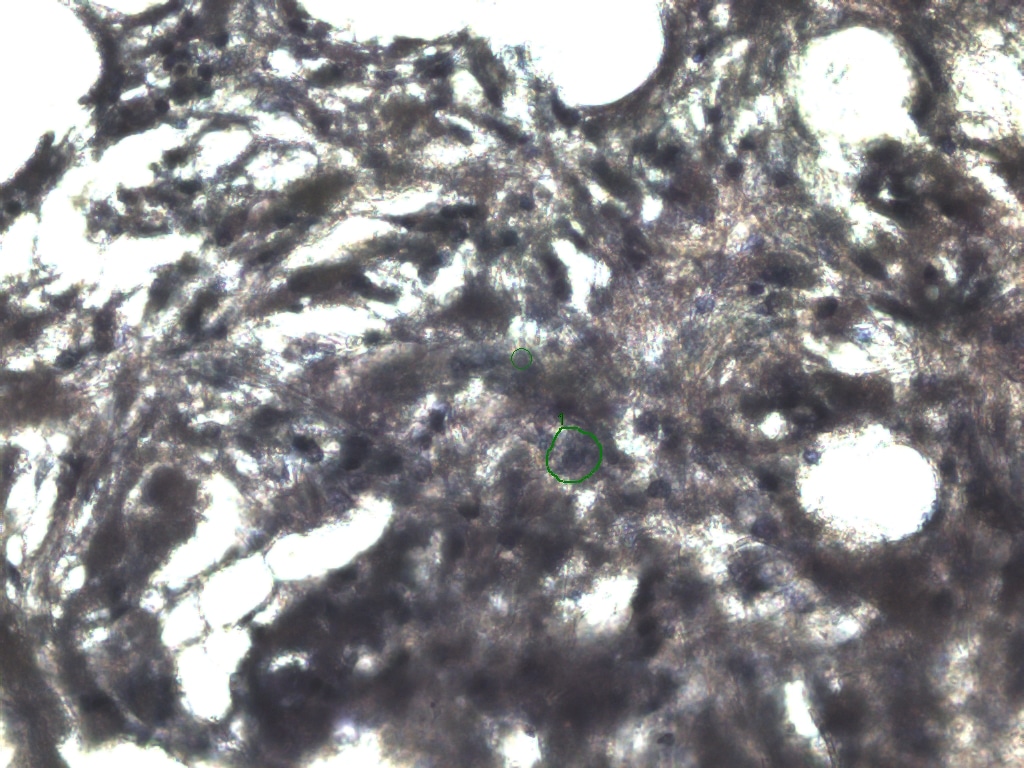

### MAC2.jpeg

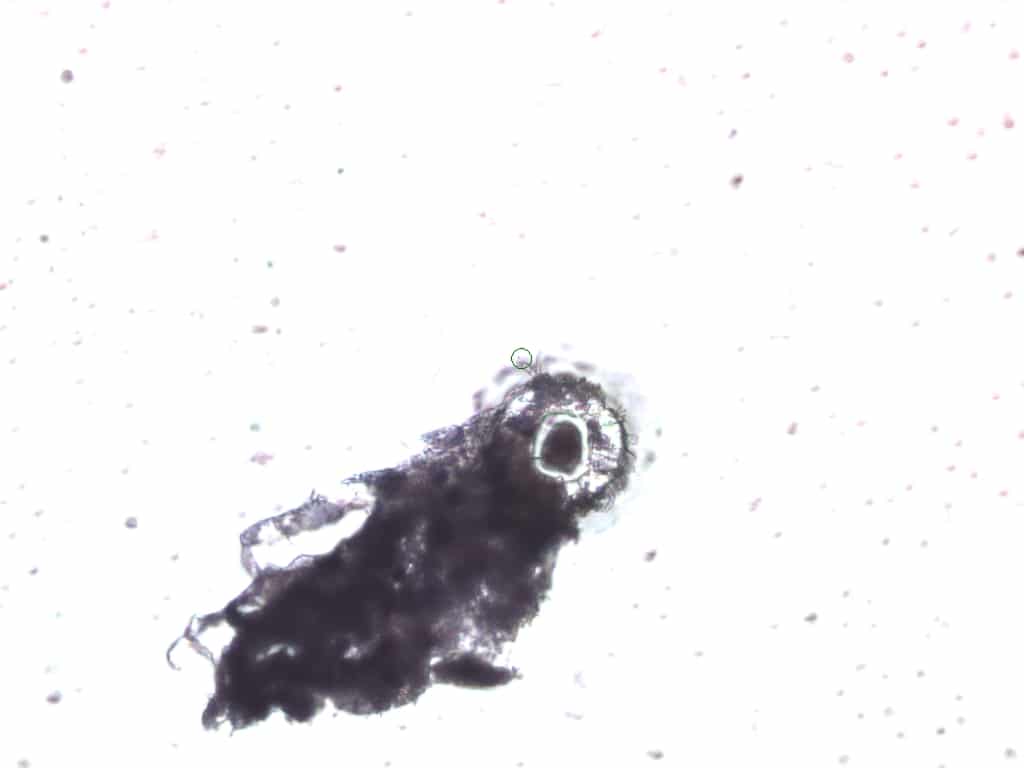

### MAC3 AFTER.jpeg

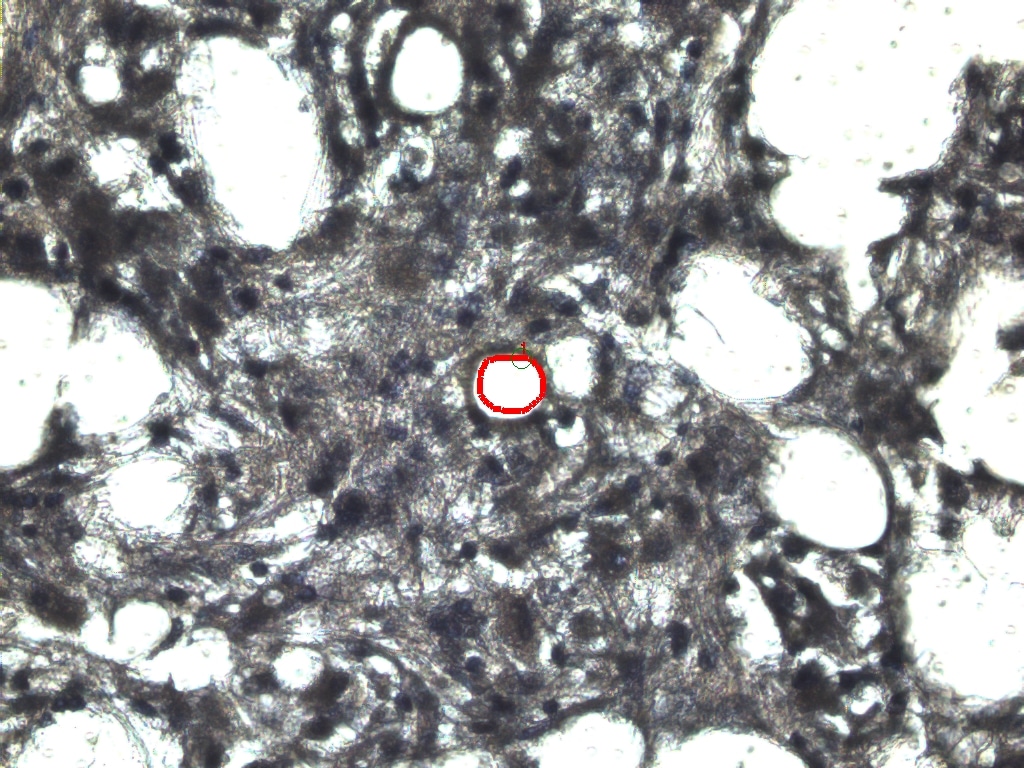

### MAC3 BEFORE.jpeg

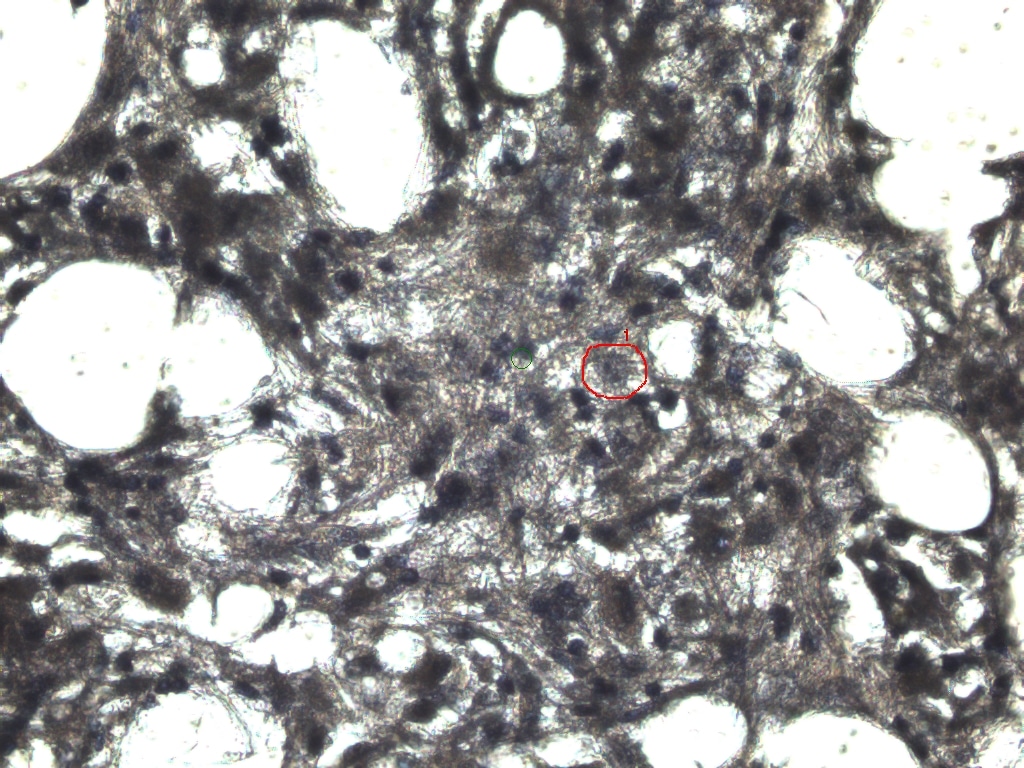

### MAC3.jpeg

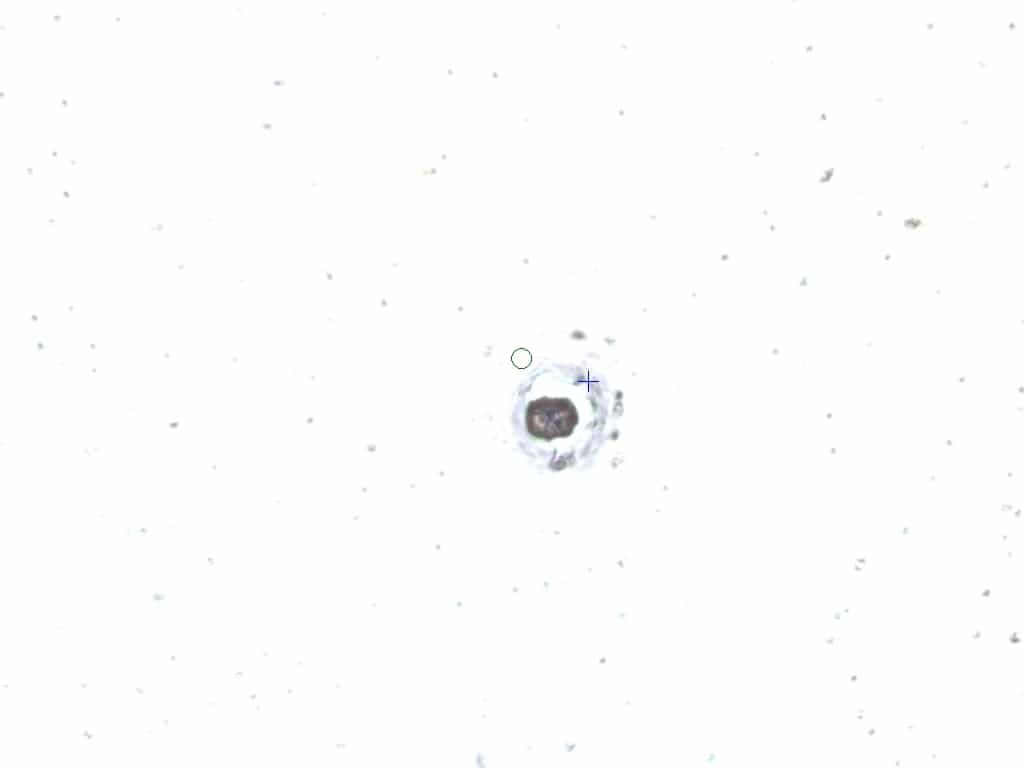

### MAC4 AFTER.jpeg

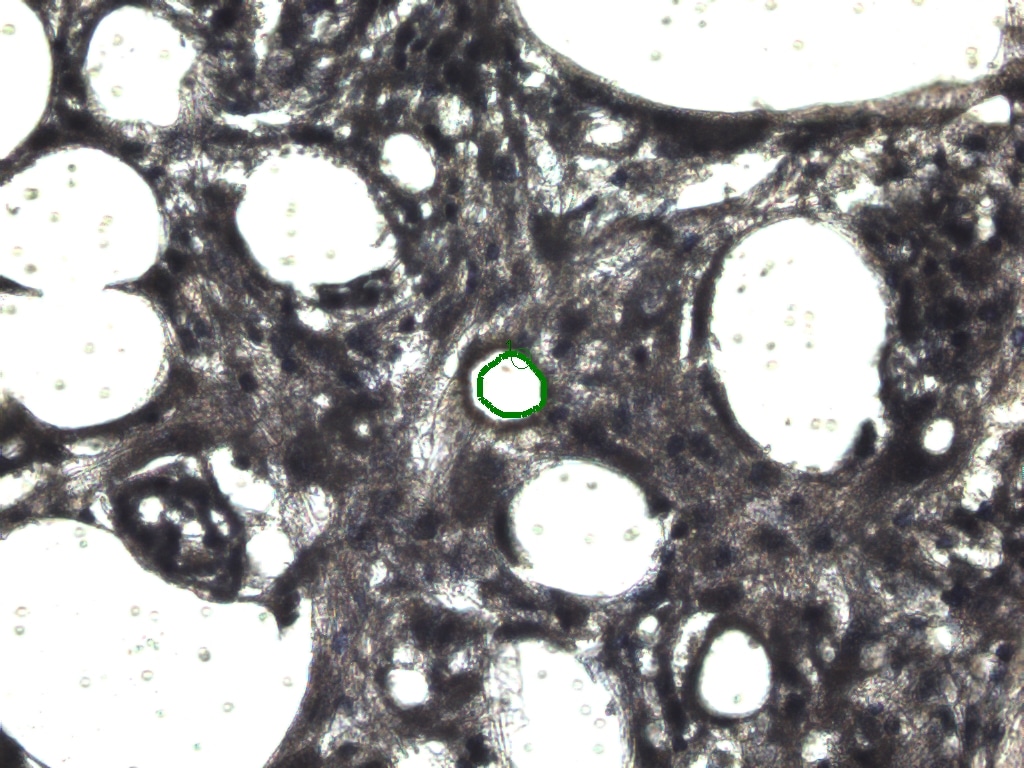

### MAC4 BEFORE.jpeg

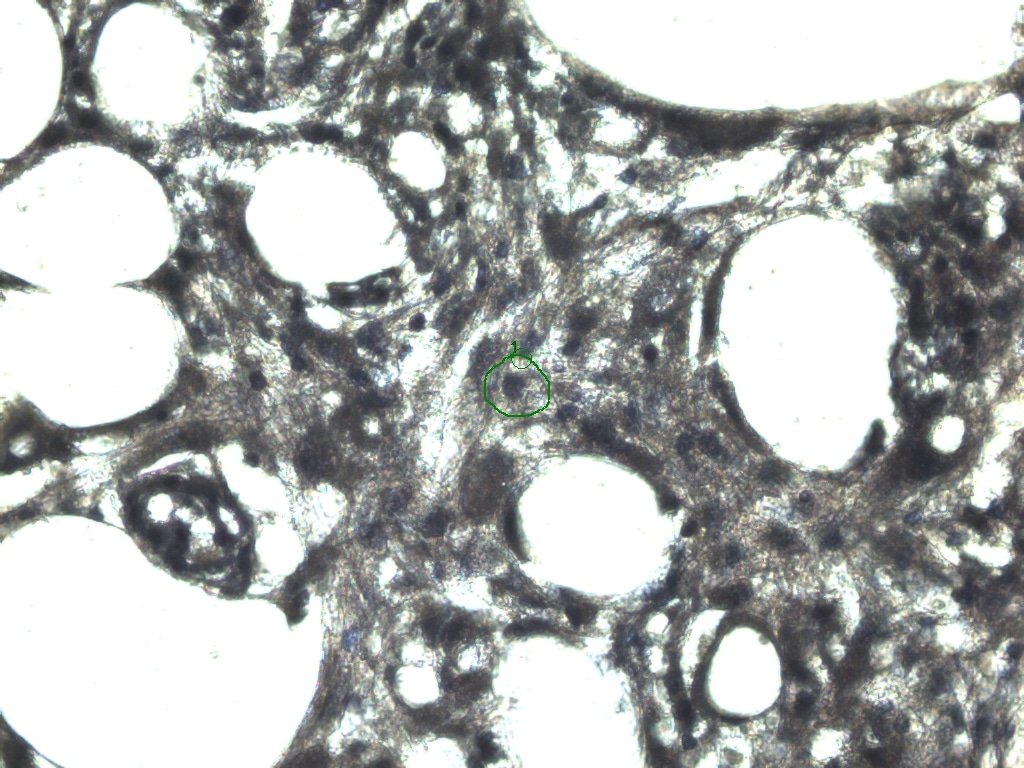

### MAC4.jpeg

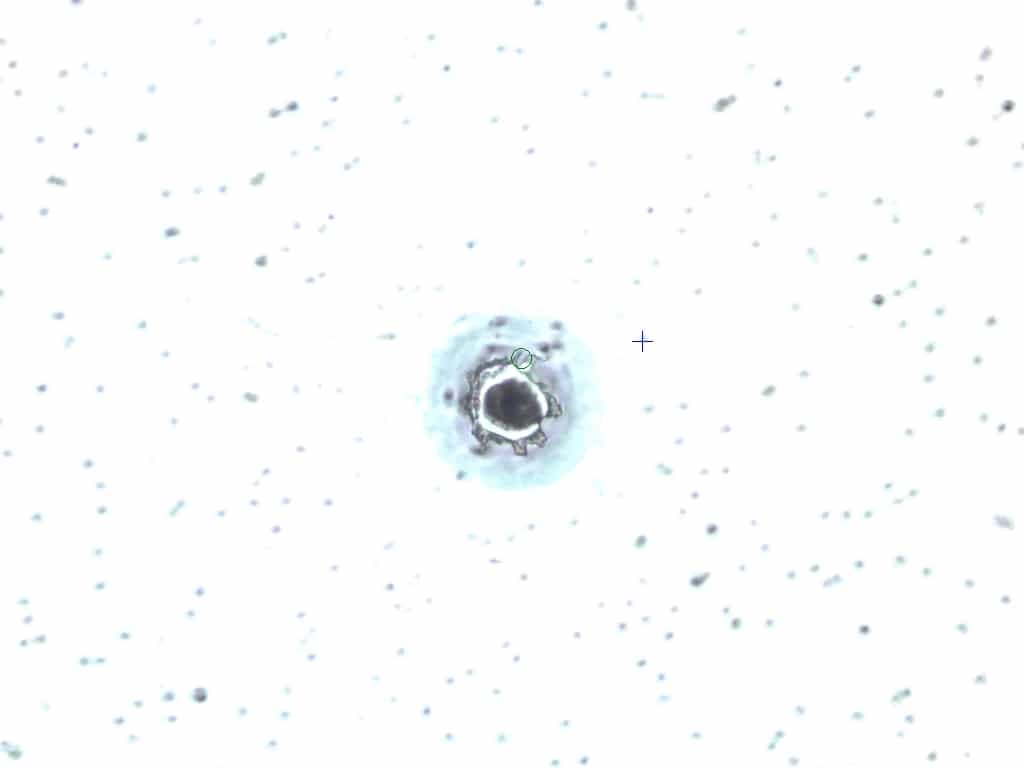

### MAC5 AFTER ABLATION FINAL.jpeg

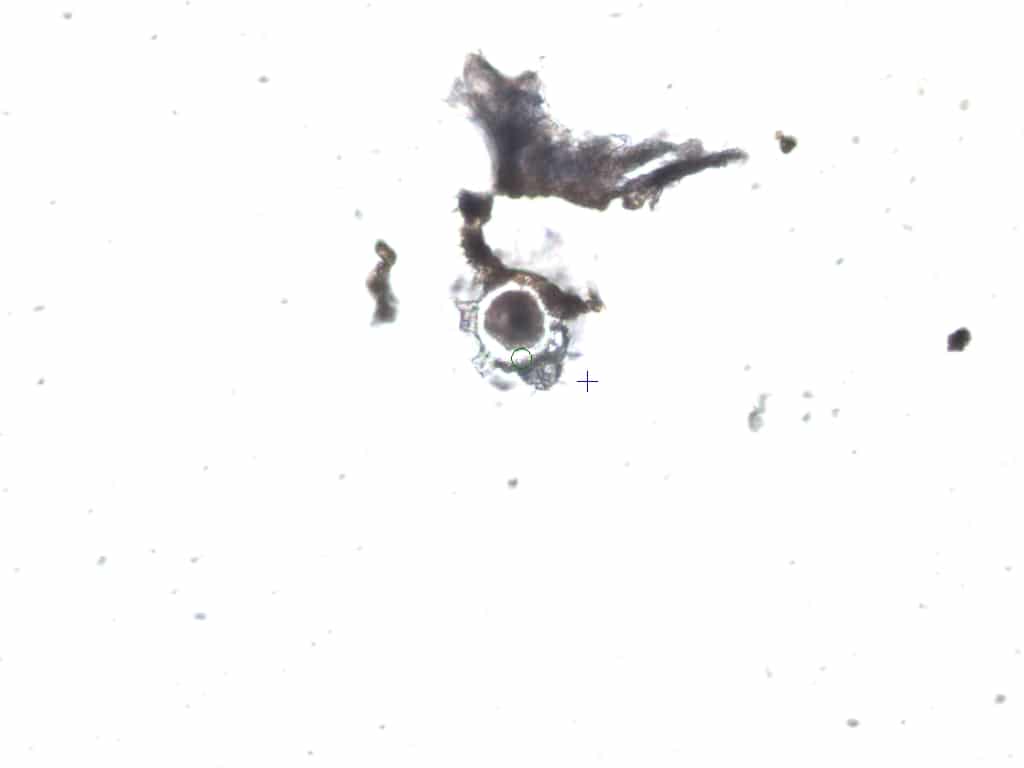

### MAC5 AFTER ABLATION.jpeg

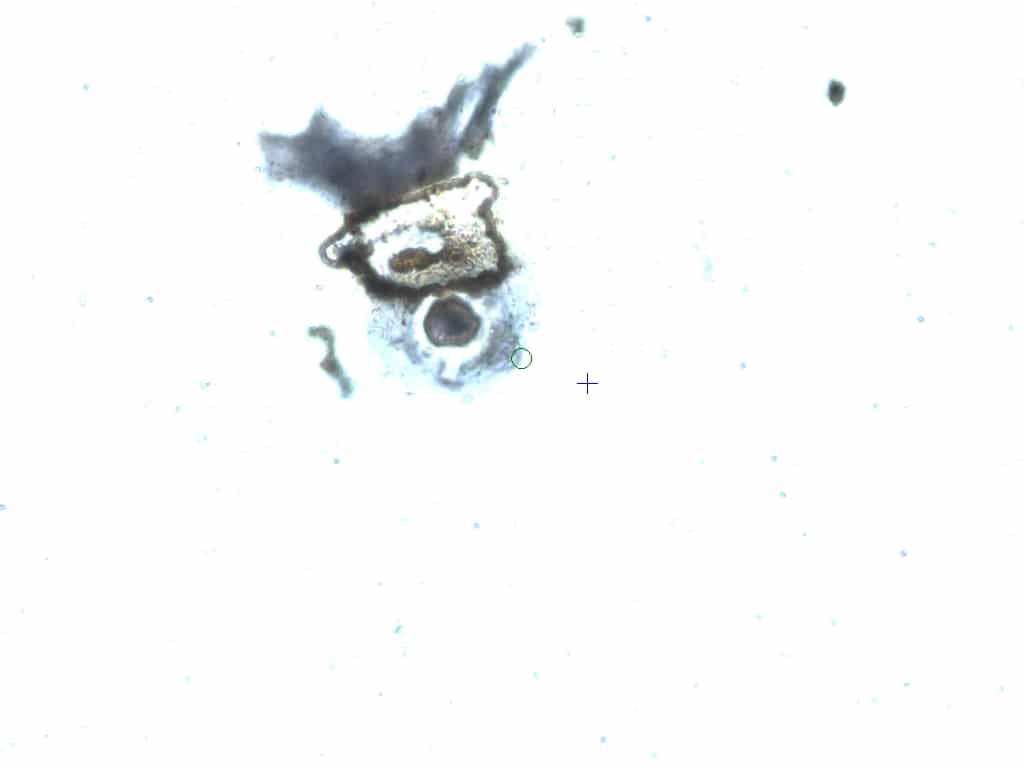

### MAC5 AFTER.jpeg

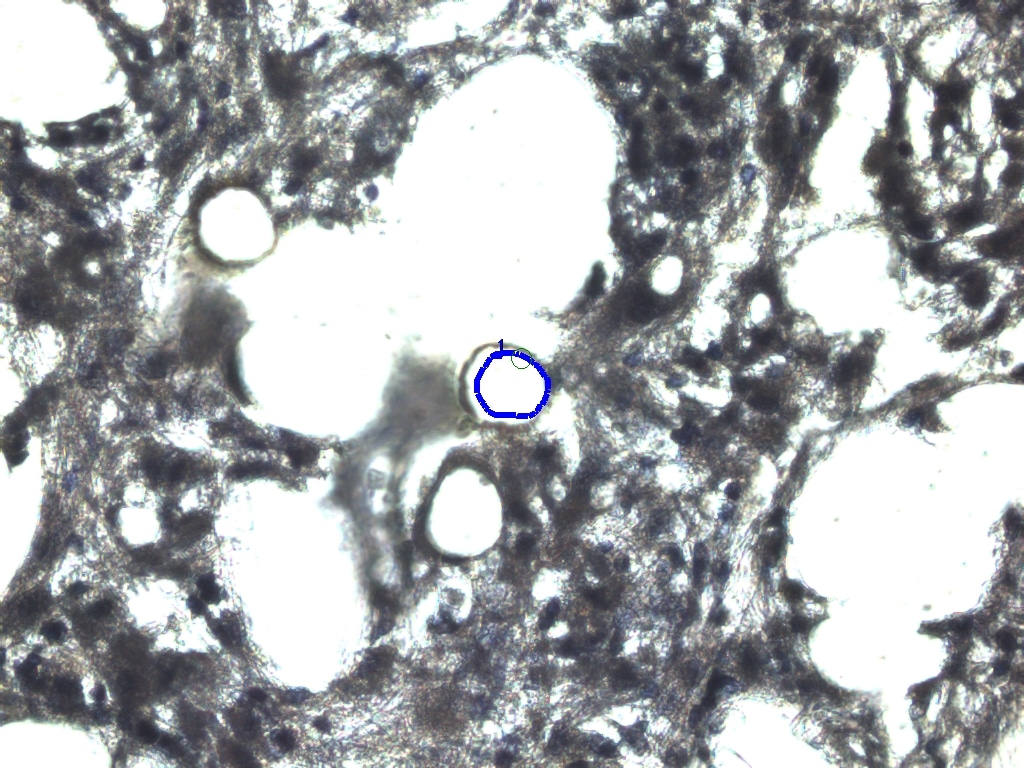

### MAC5 BEFORE.jpeg

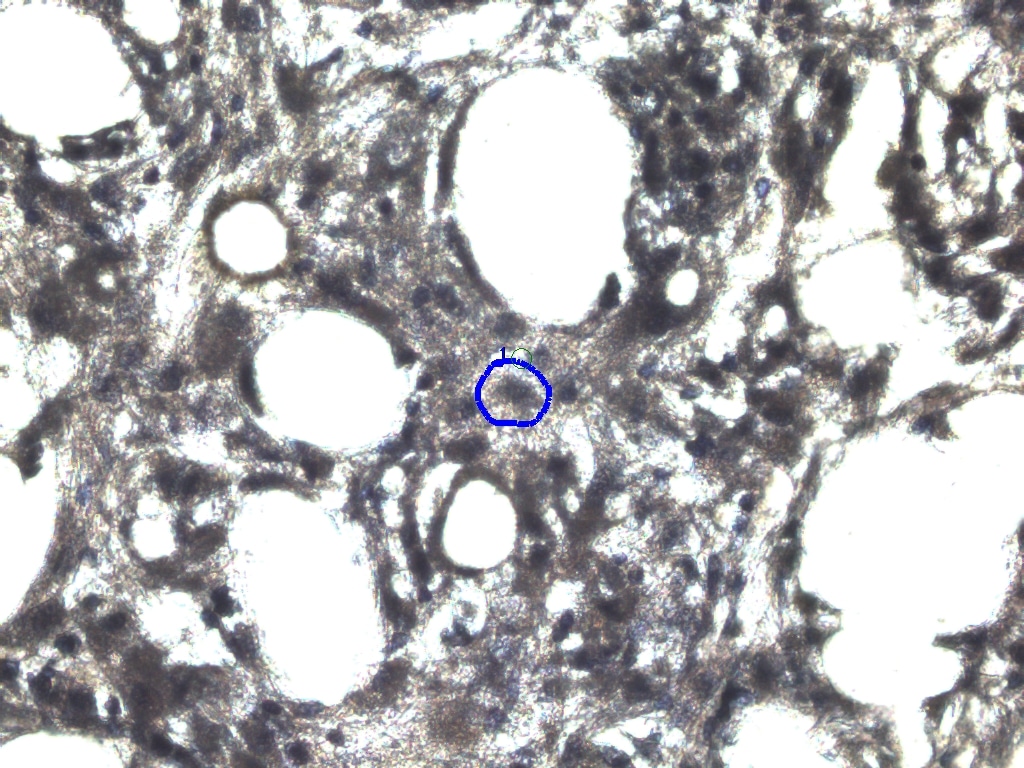

### MAC5.jpeg

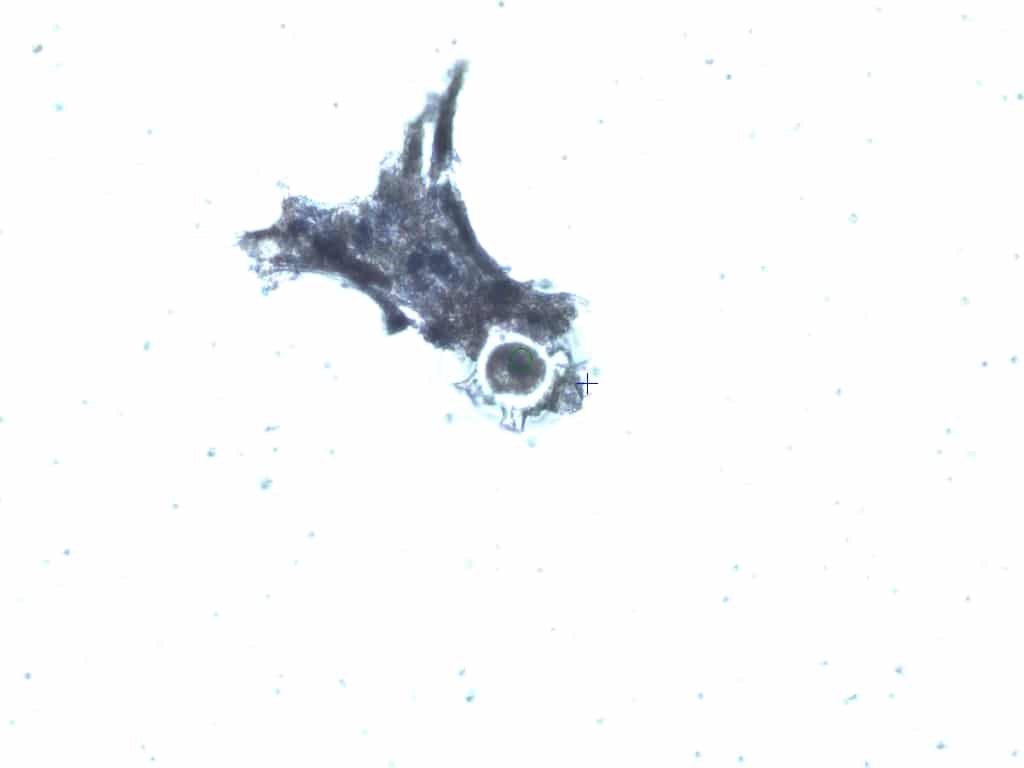

### MAC6 AFTER.jpeg

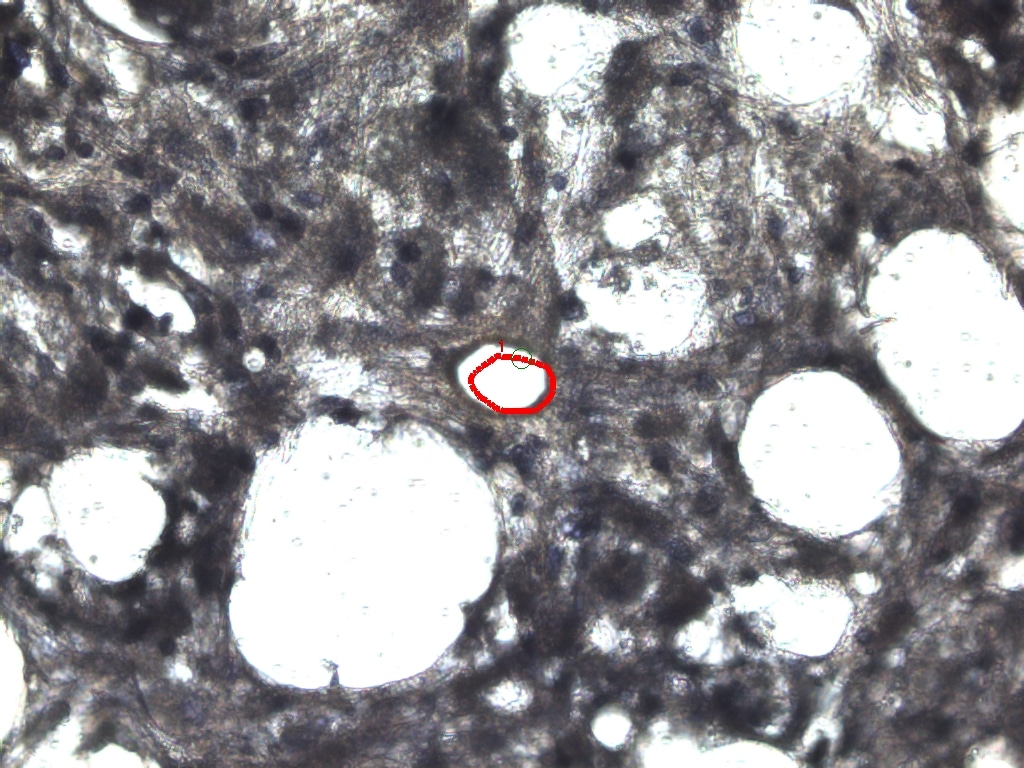

### MAC6 BEFORE.jpeg

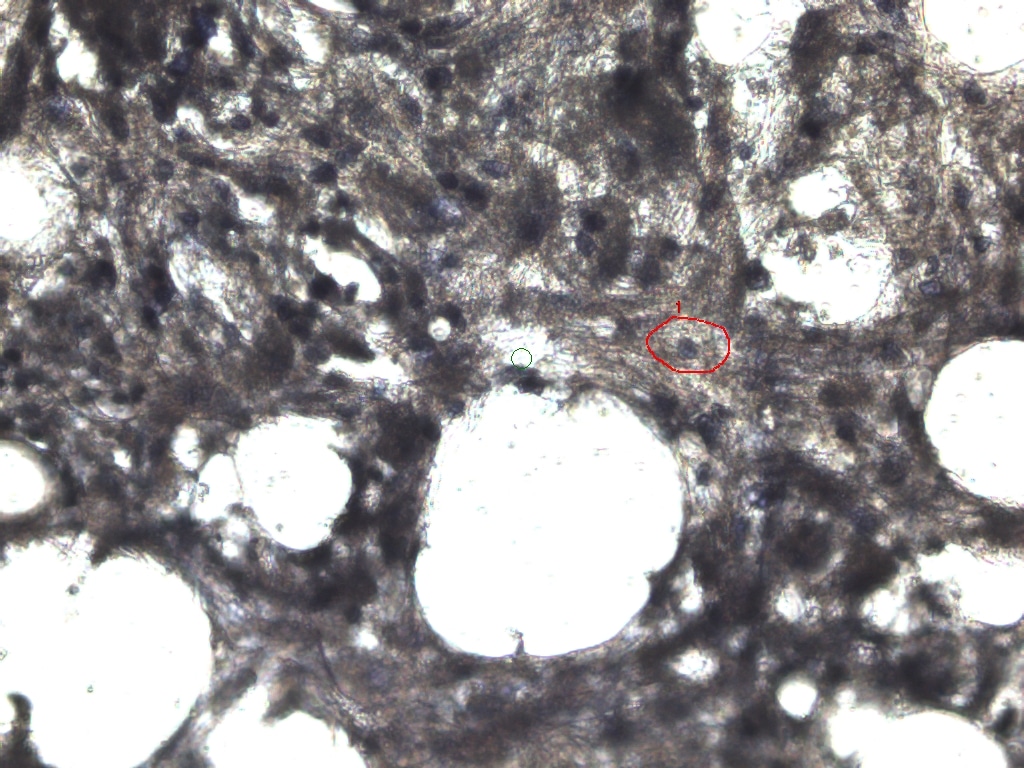

### MAC6.jpeg

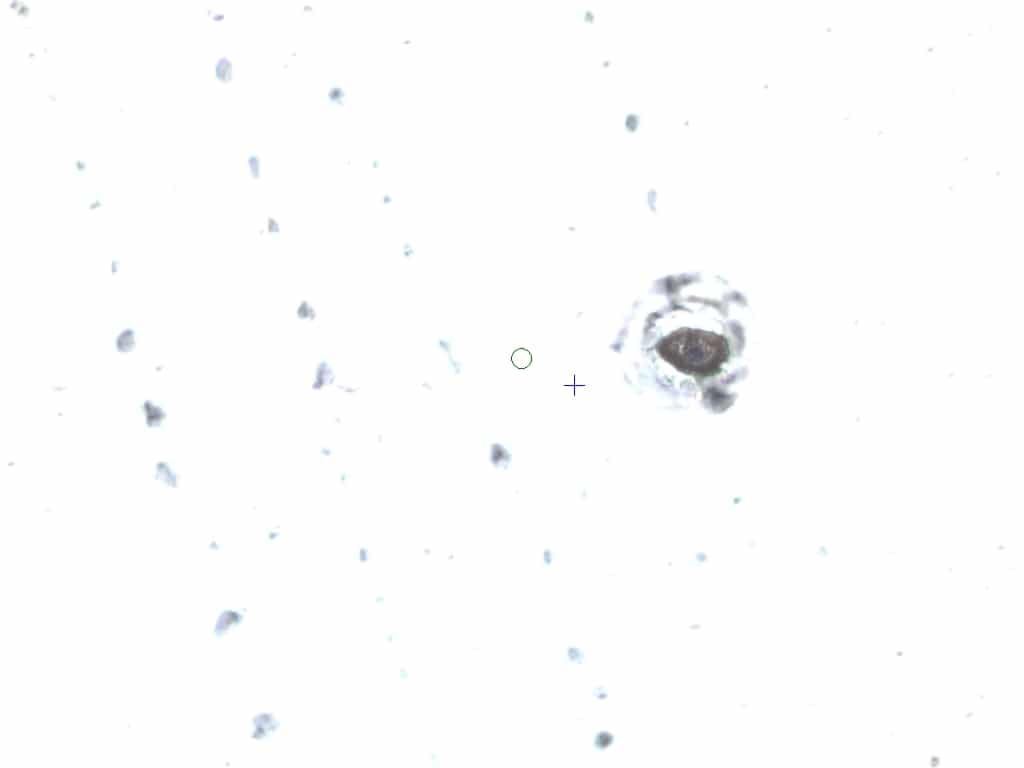

### MAC7 AFTER ABLATION.jpeg

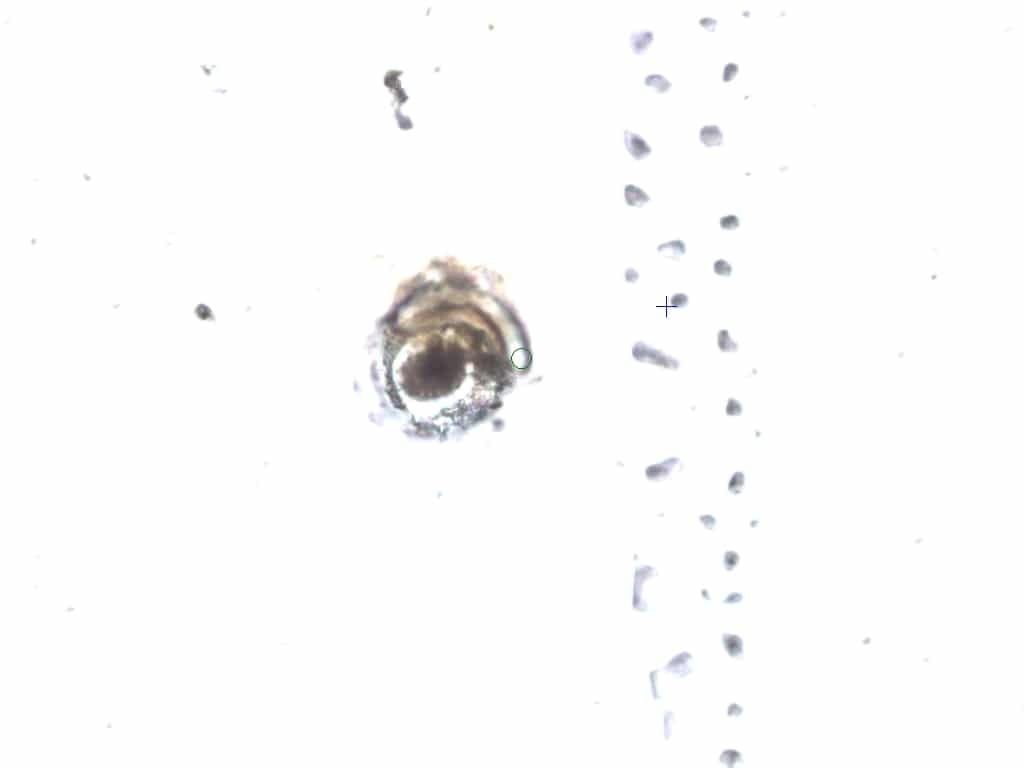

### MAC7 AFTER.jpeg

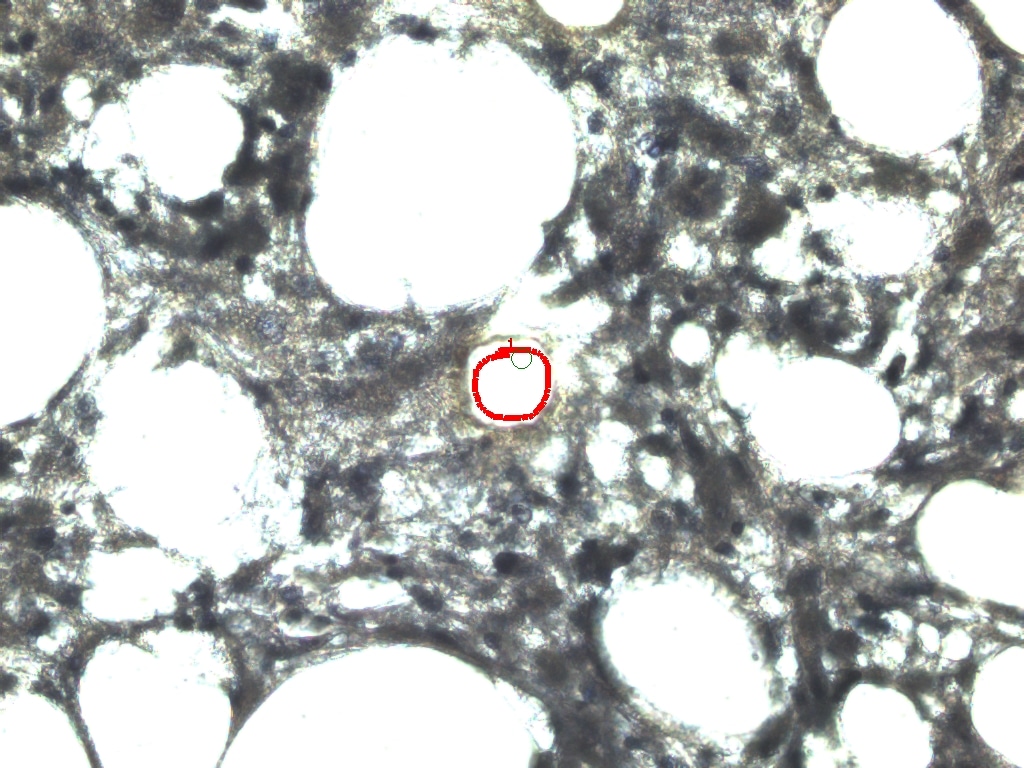

### MAC7 BEFORE.jpeg

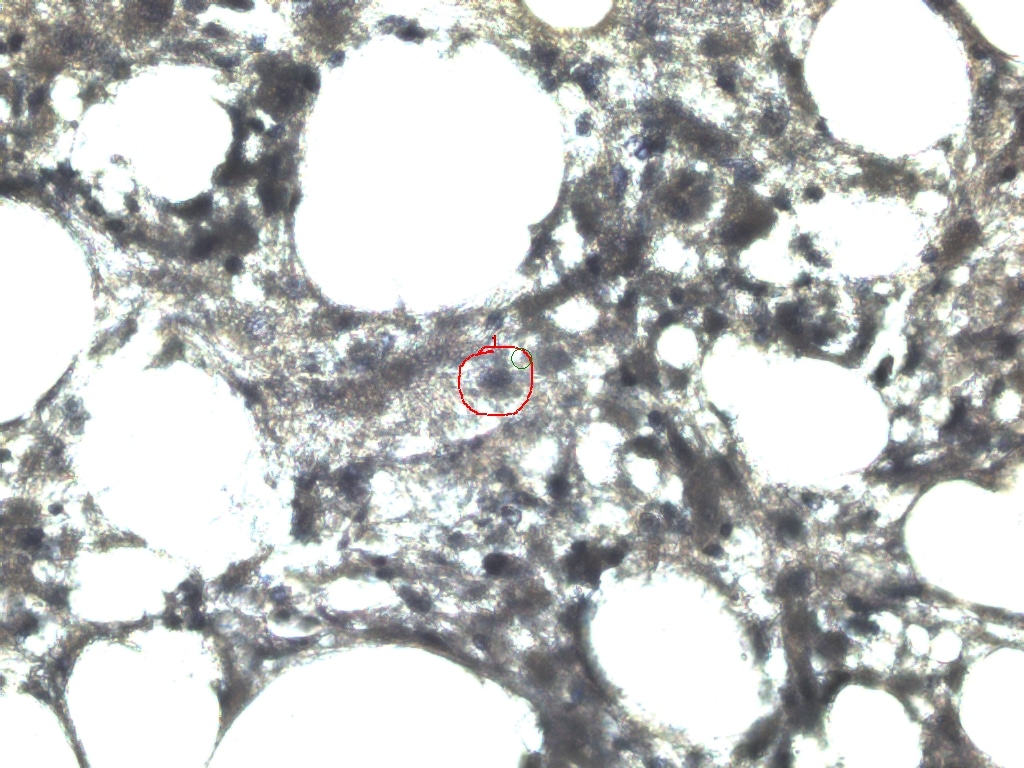

### MAC7.jpeg

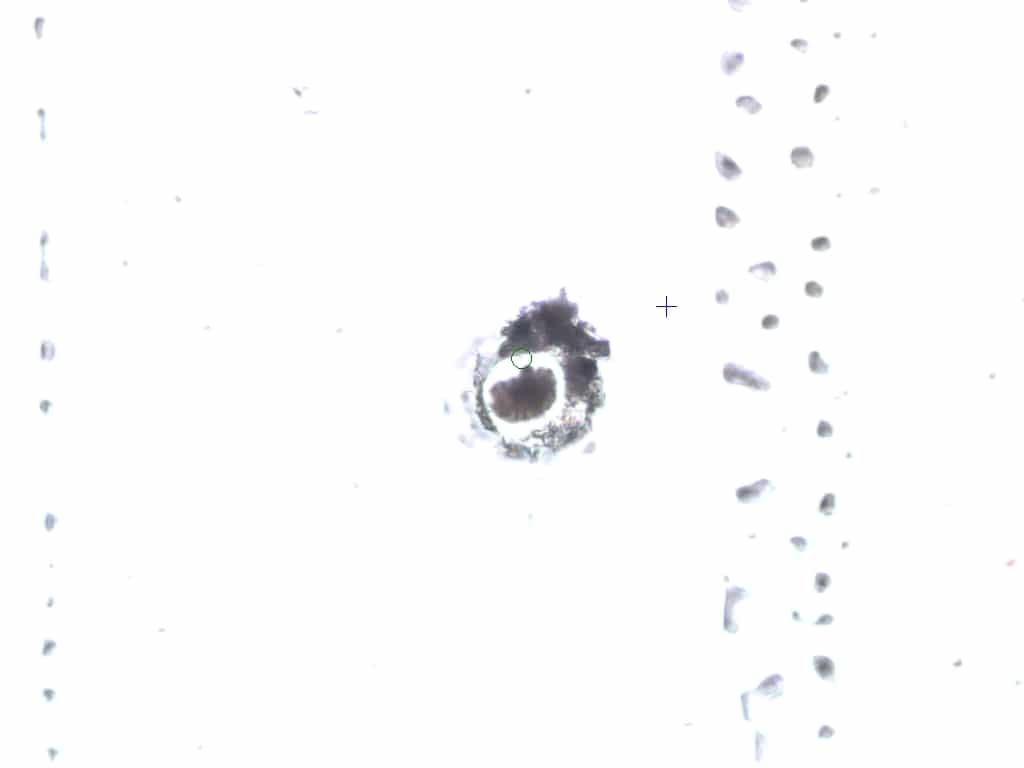

### MAC8 AFTER.jpeg

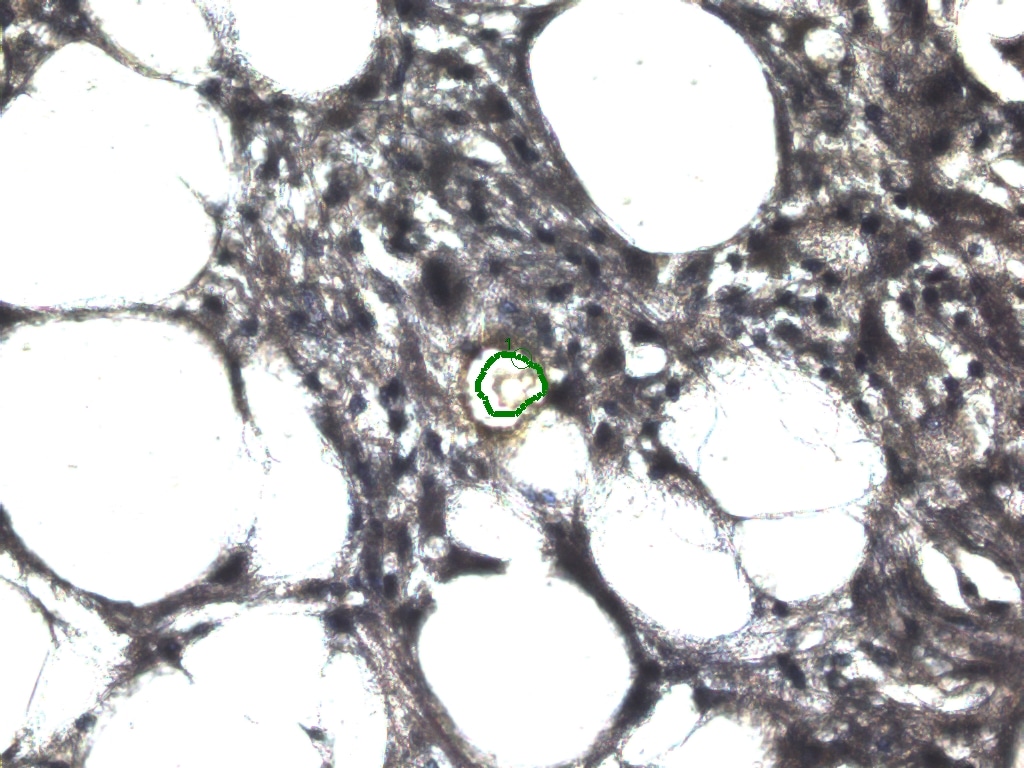

### MAC8 BEFORE.jpeg

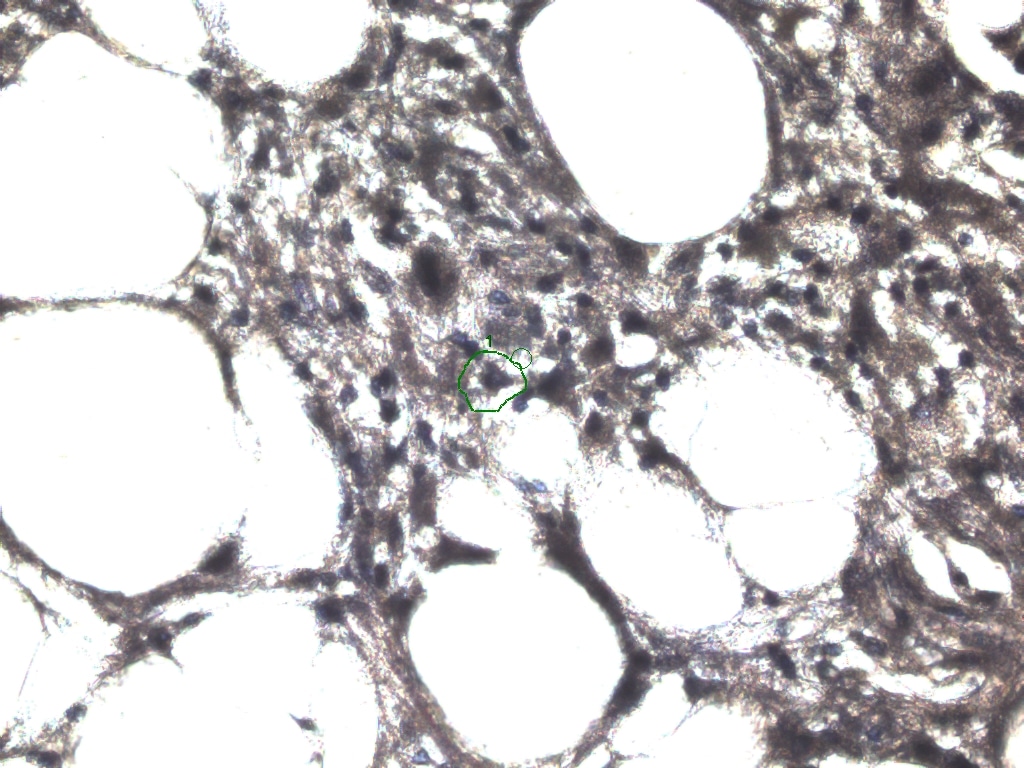

### MAC8.jpeg

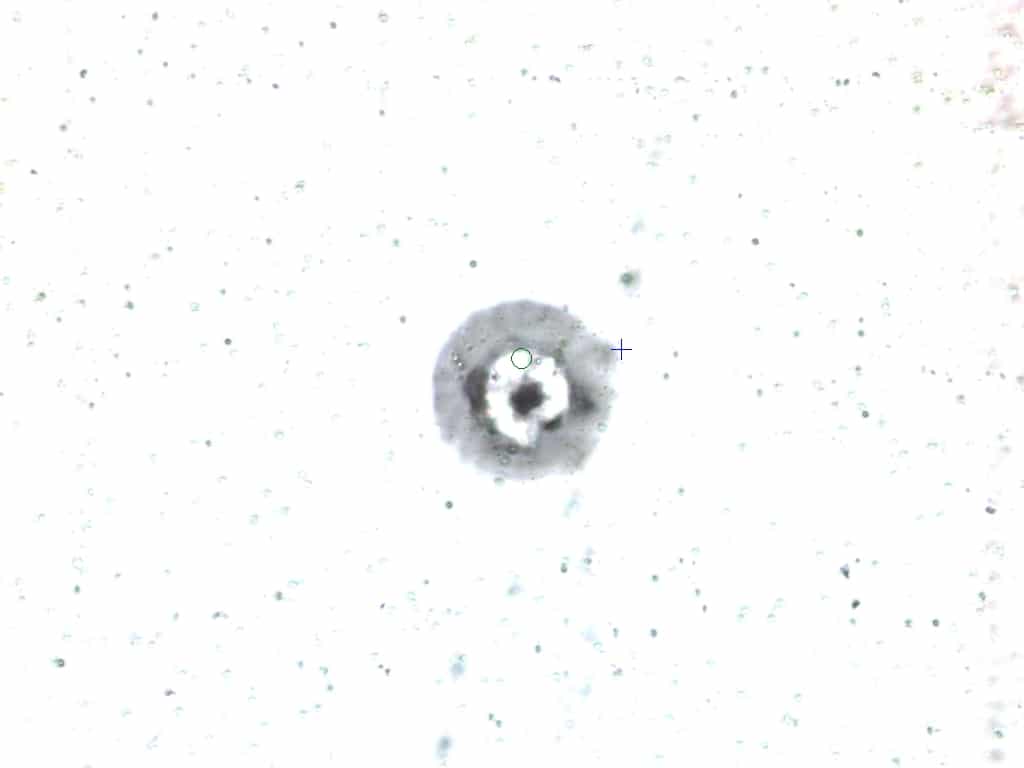

### MAC9 AFTER ABLATION.jpeg

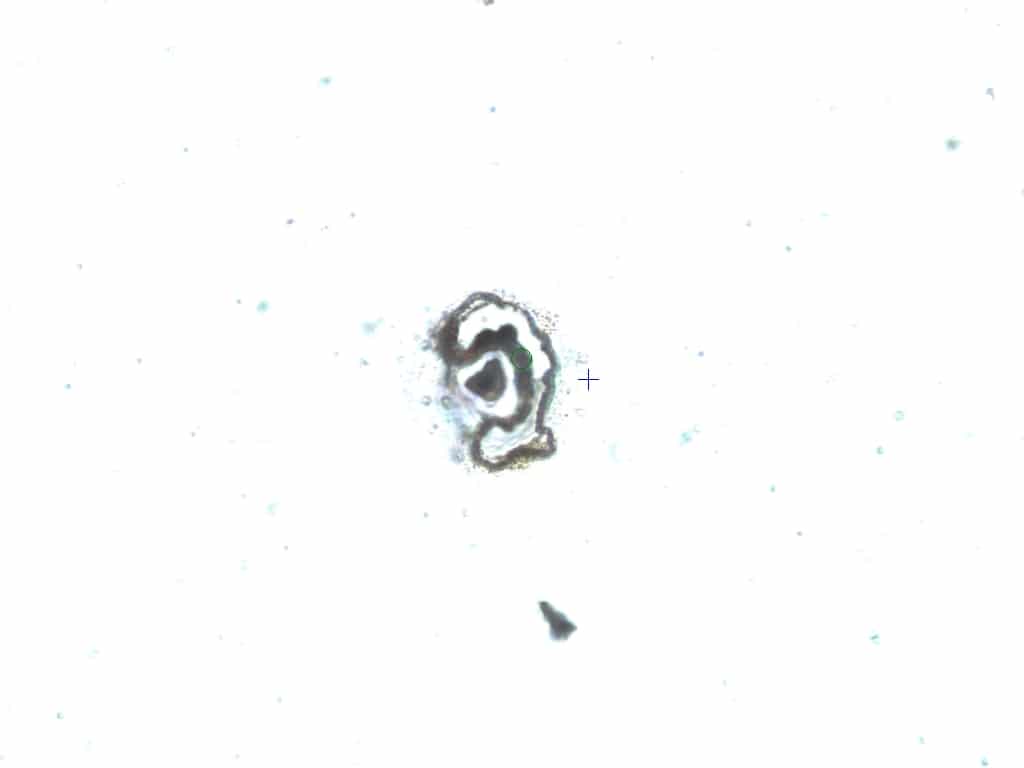

### MAC9 AFTER.jpeg

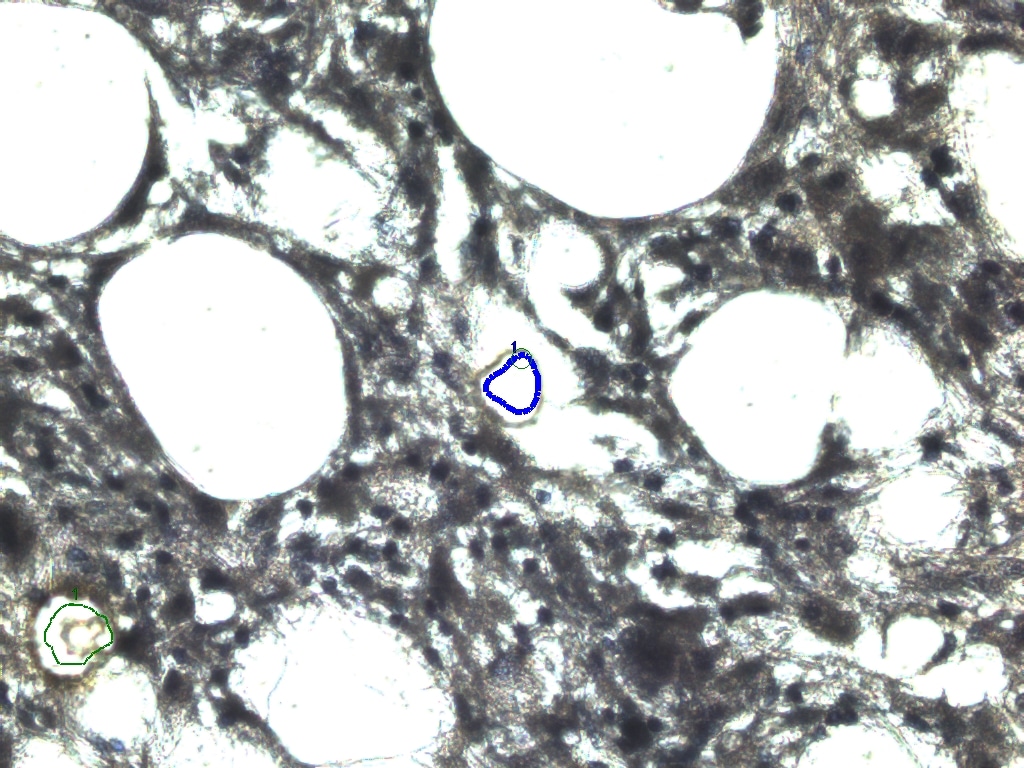

### MAC9 BEFORE.jpeg

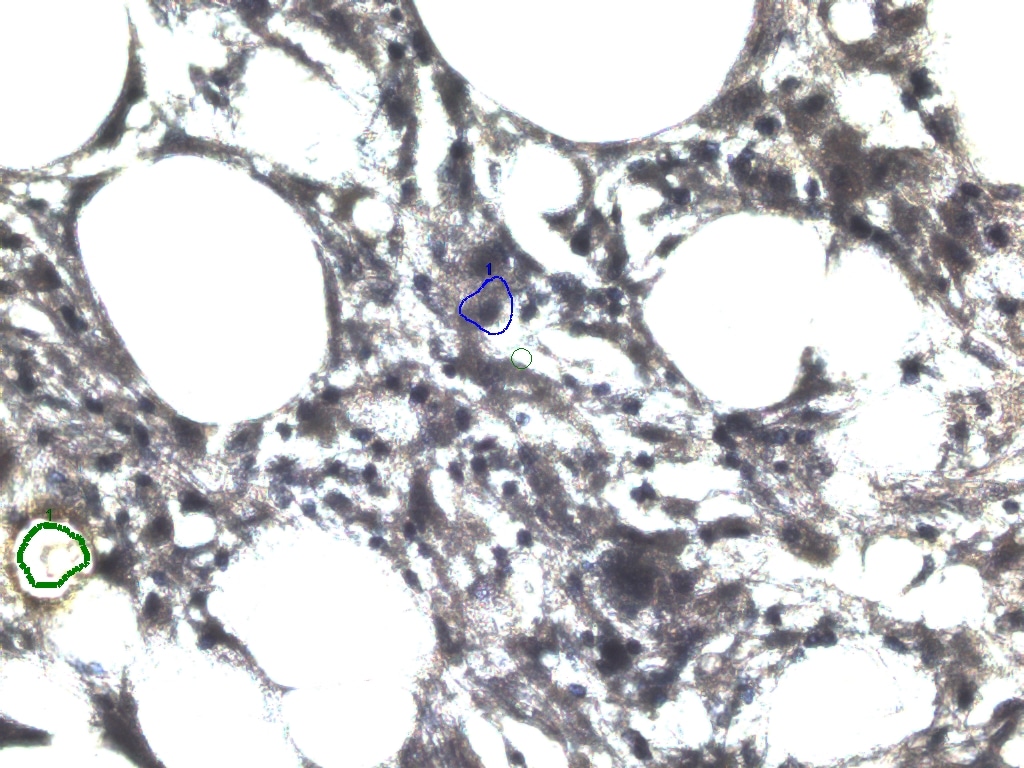

### MAC9.jpeg

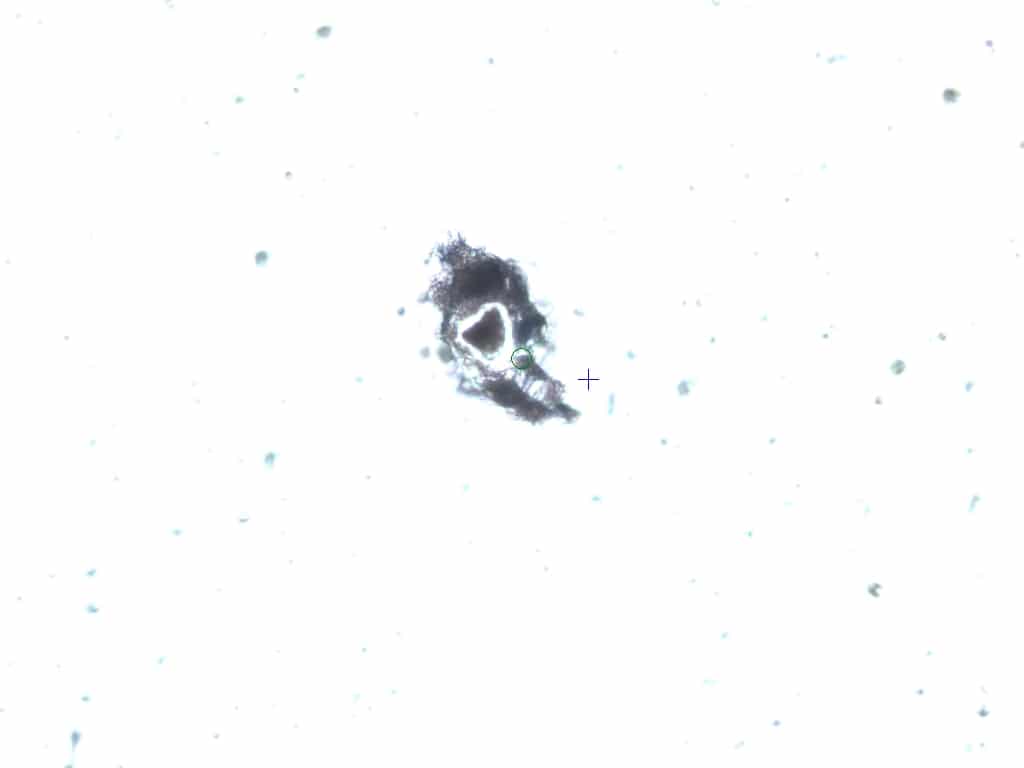

### MAC10 AFTER.jpeg

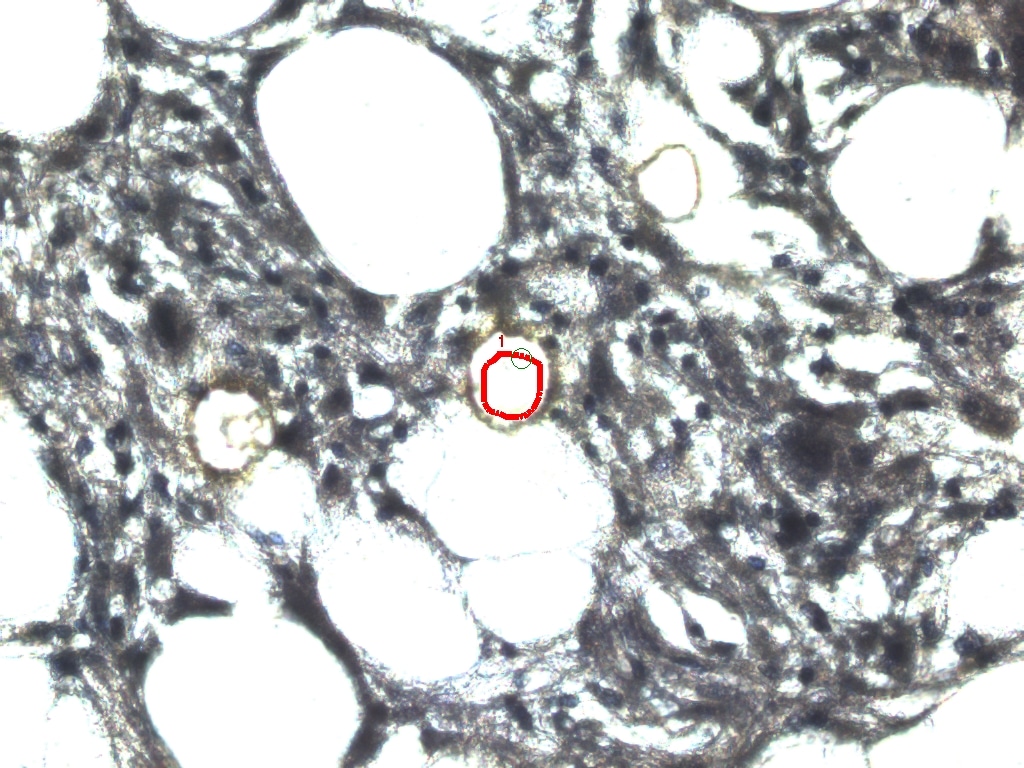

### MAC10 BEFORE.jpeg

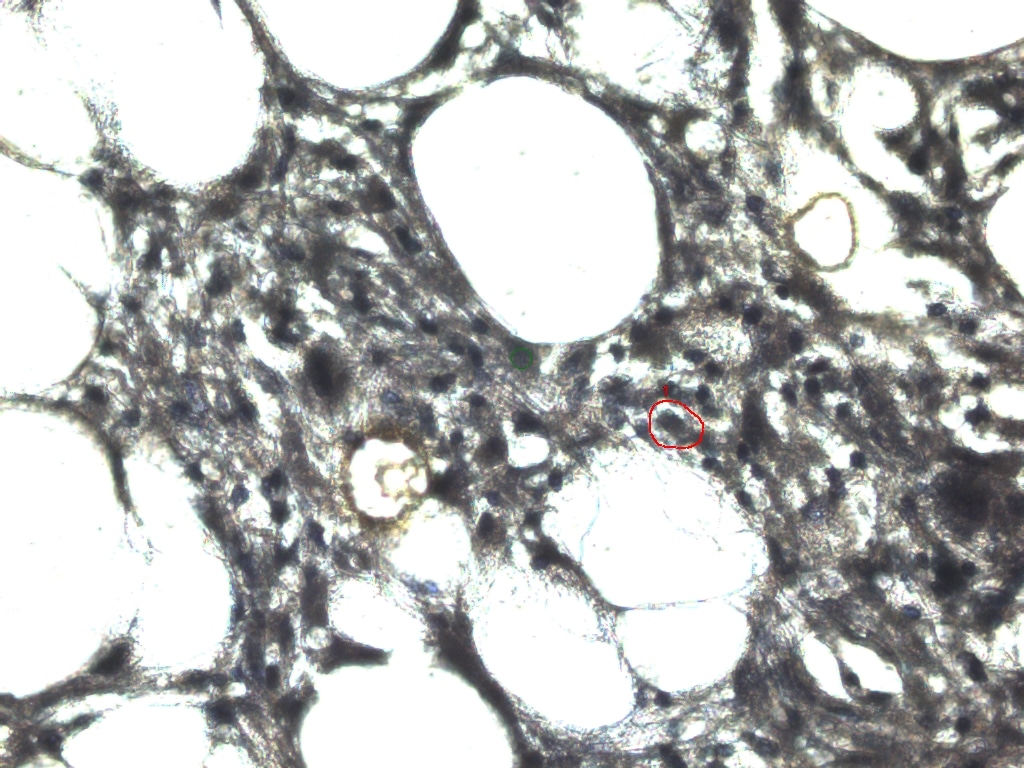

### MAC10.jpeg

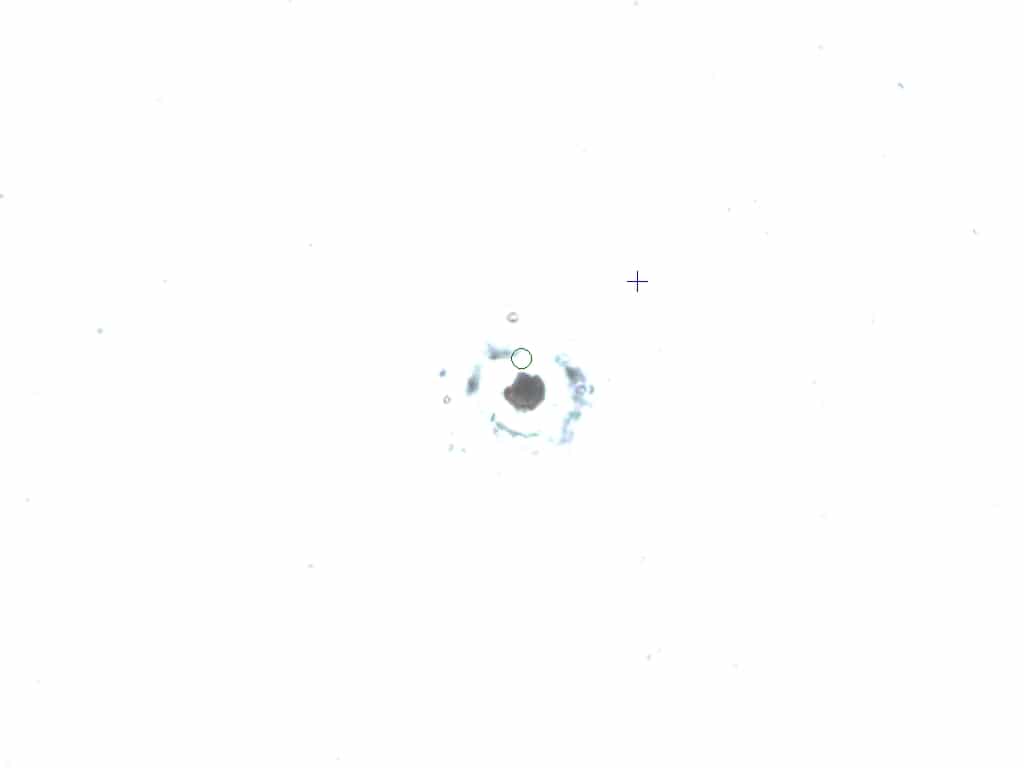
