## Supplementary figures and images for "Gene-expression profiling of single cells from archival tissue with laser-capture microdissection and Smart-3SEQ"

### CAP1 BEFORE ABLATION.jpeg

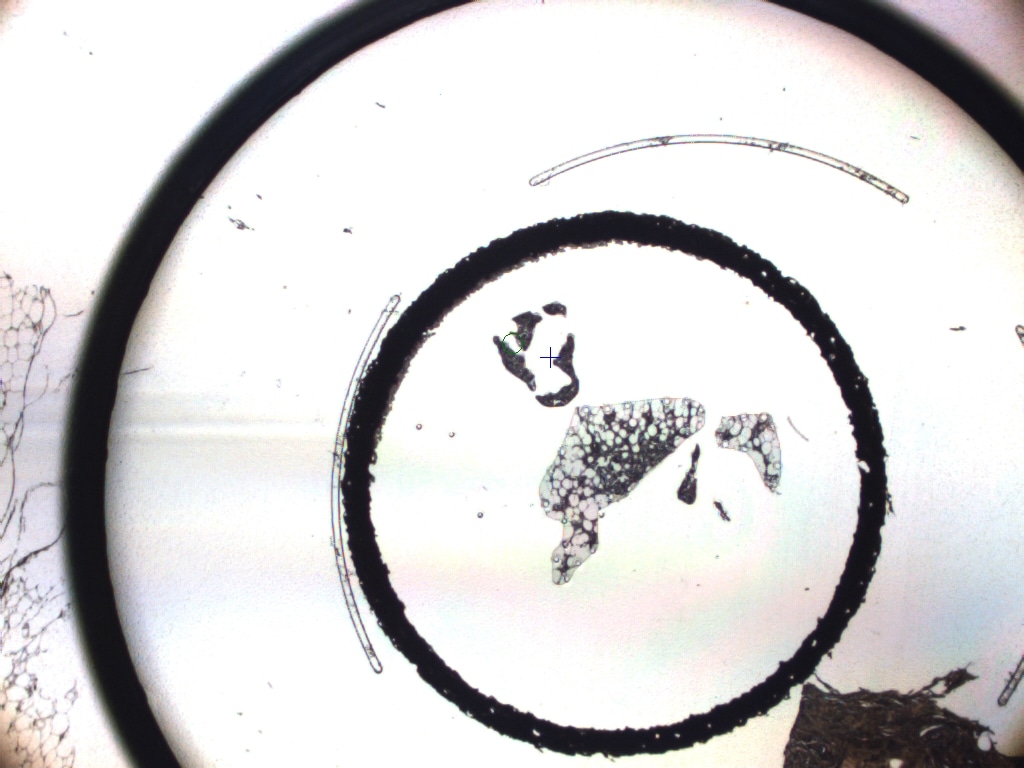

### CAP1AFTERABLATION 10x.jpeg

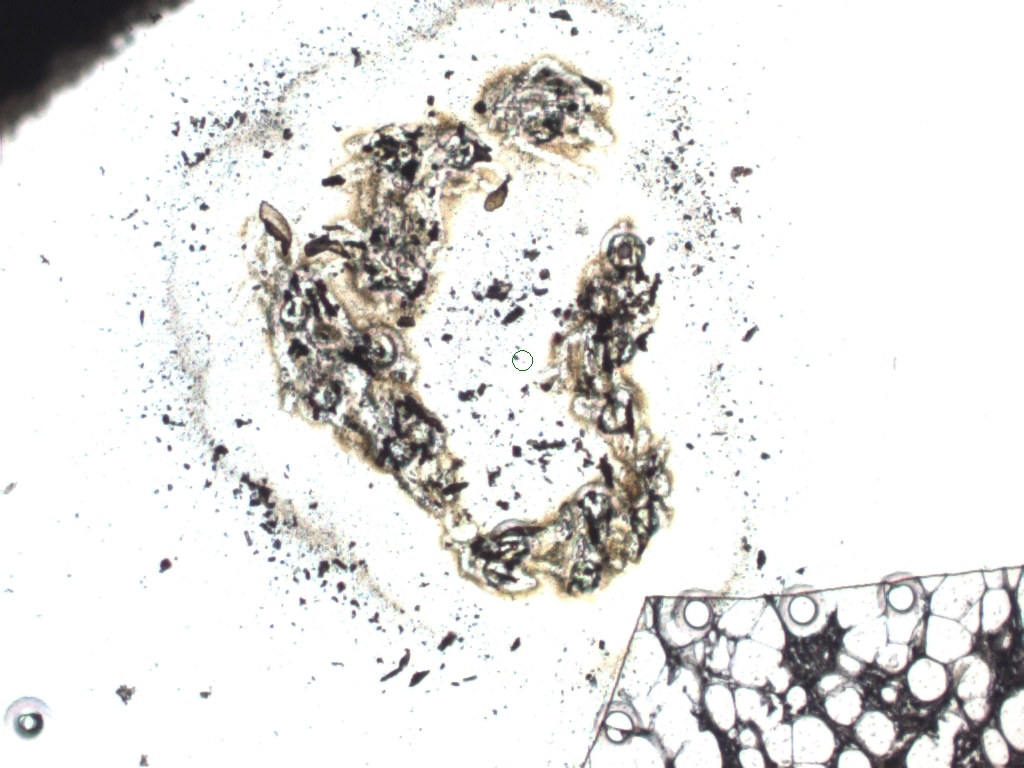

### CAP1AFTERABLATION.jpeg

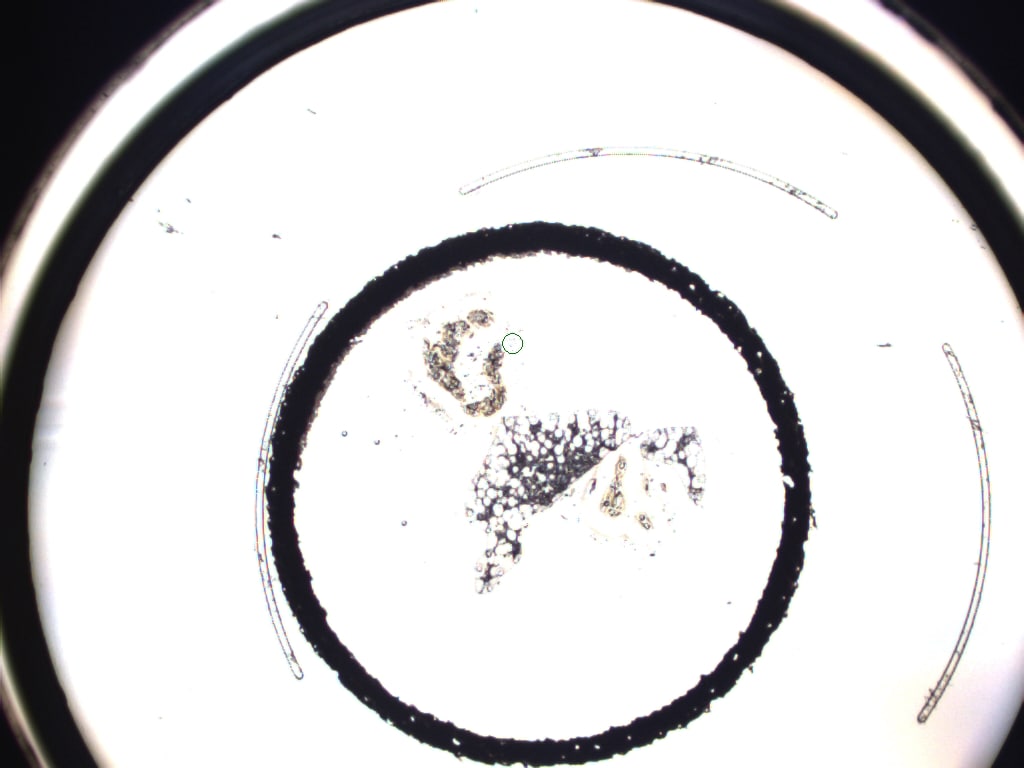

### DCIS1 .jpeg

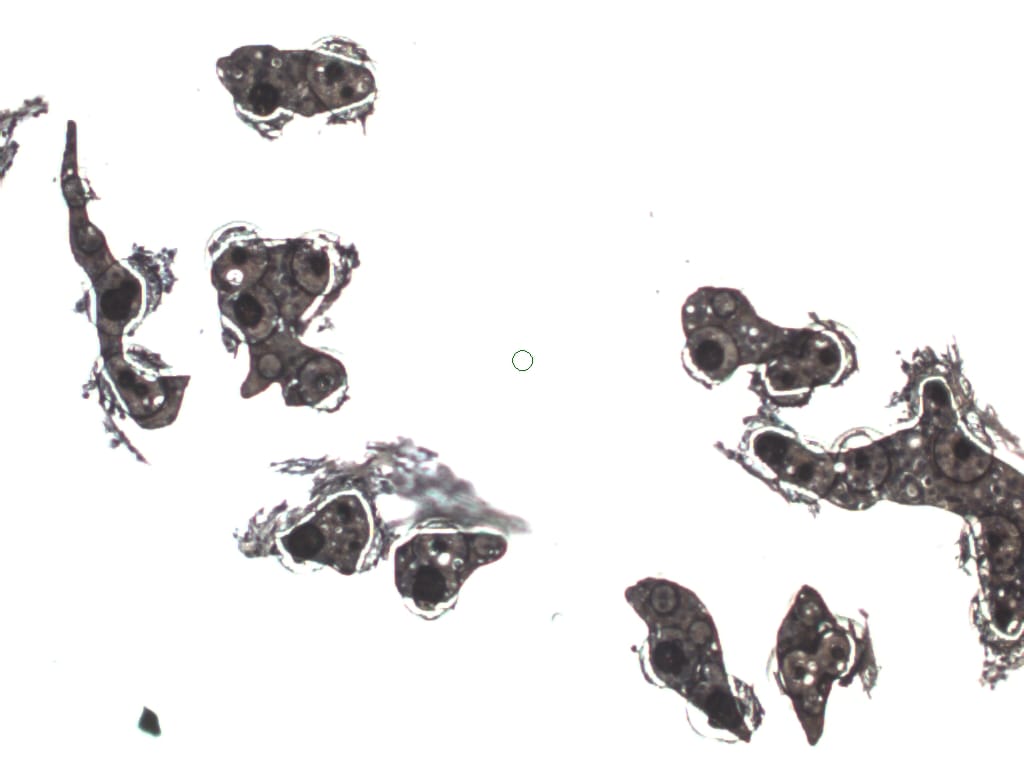

### DCIS1 BEFORE.jpeg

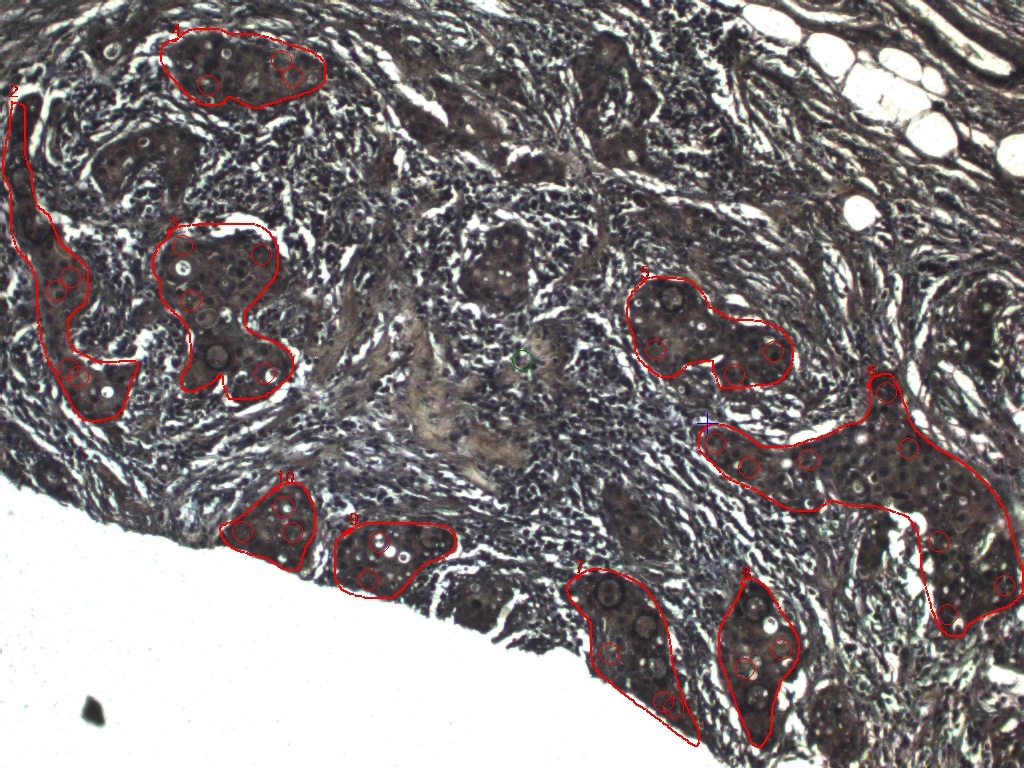

### DCIS1-2 BEFORE.jpeg

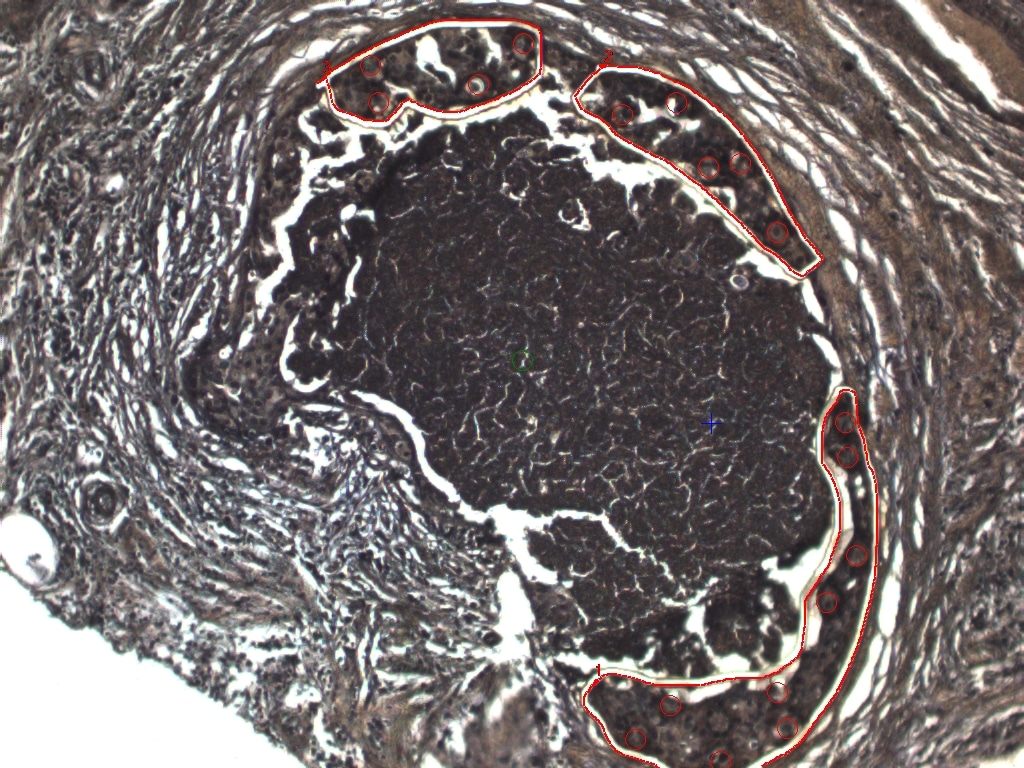

### DCIS1MAC1 ABLATION .jpeg

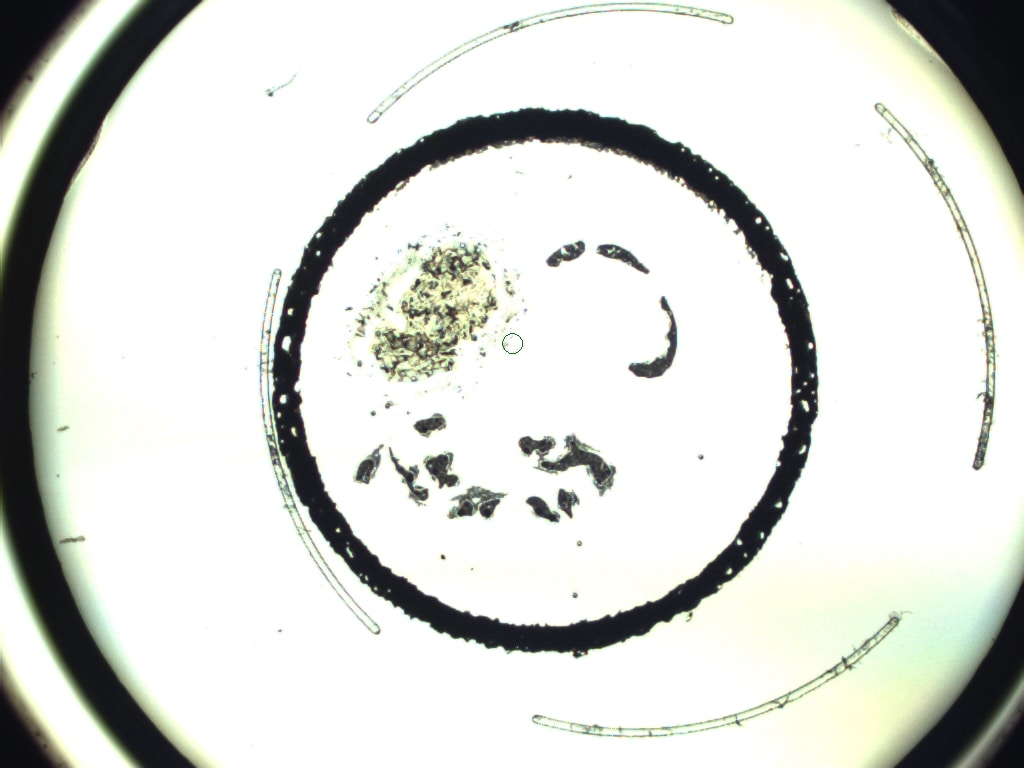

### DCIS1MAC1 BEFORE .jpeg

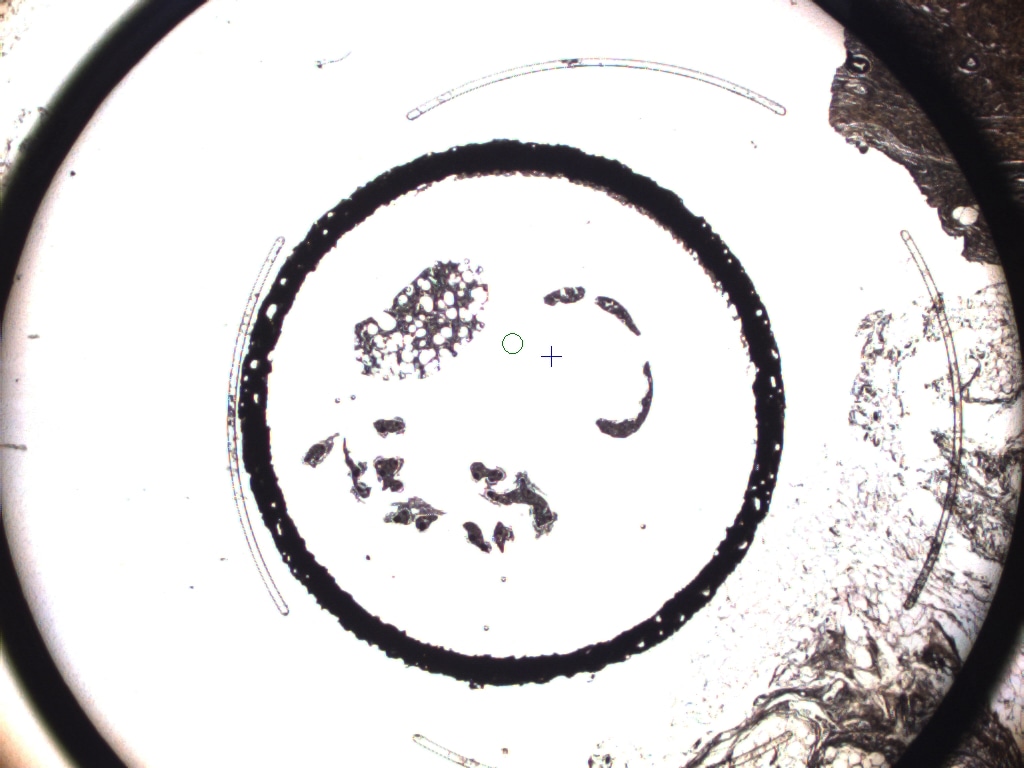

### DCIS2 BEFORE.jpeg

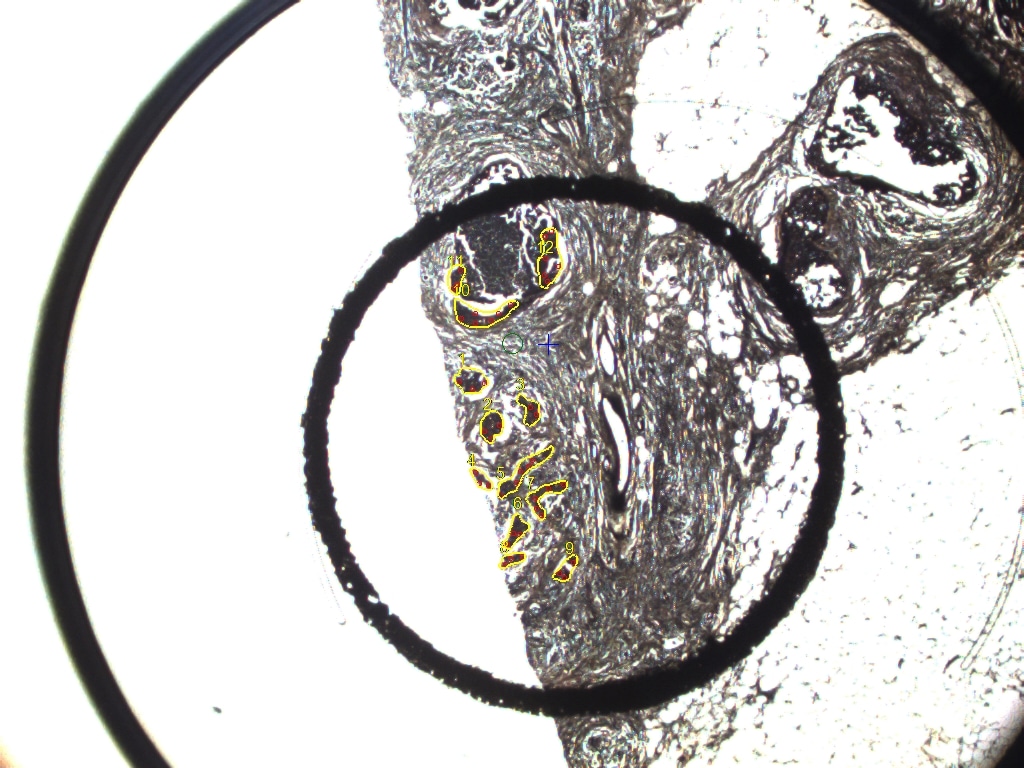

### DCIS2MAC2 ABLATION 10x .jpeg

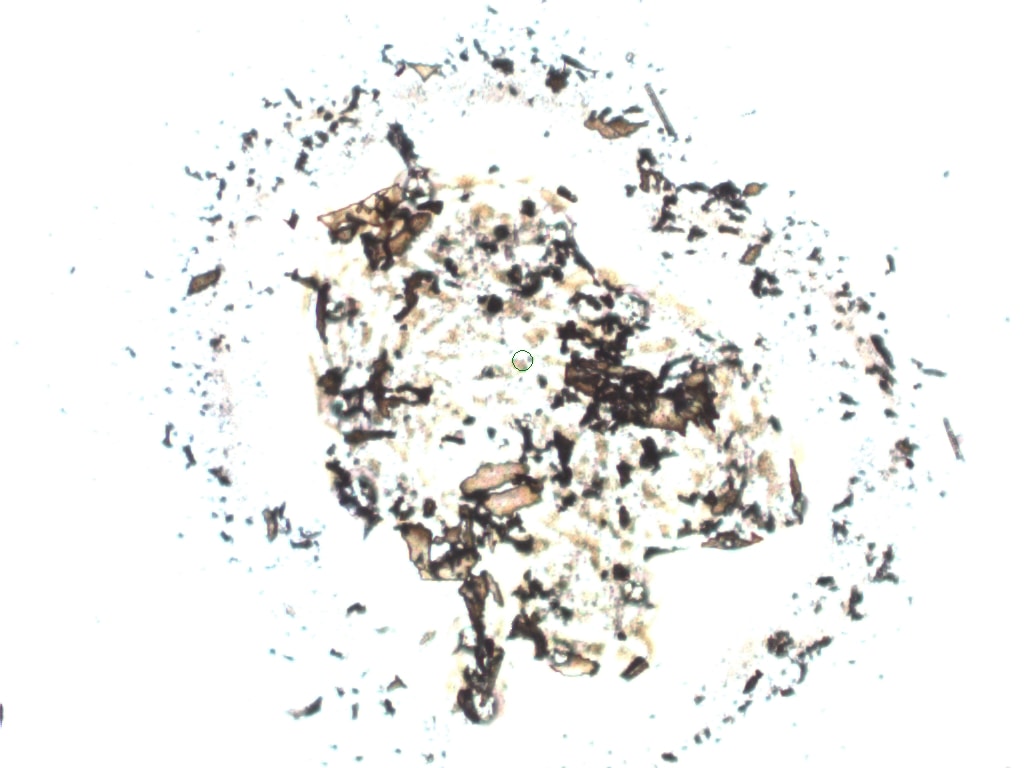

### DCIS2MAC2 ABLATION .jpeg

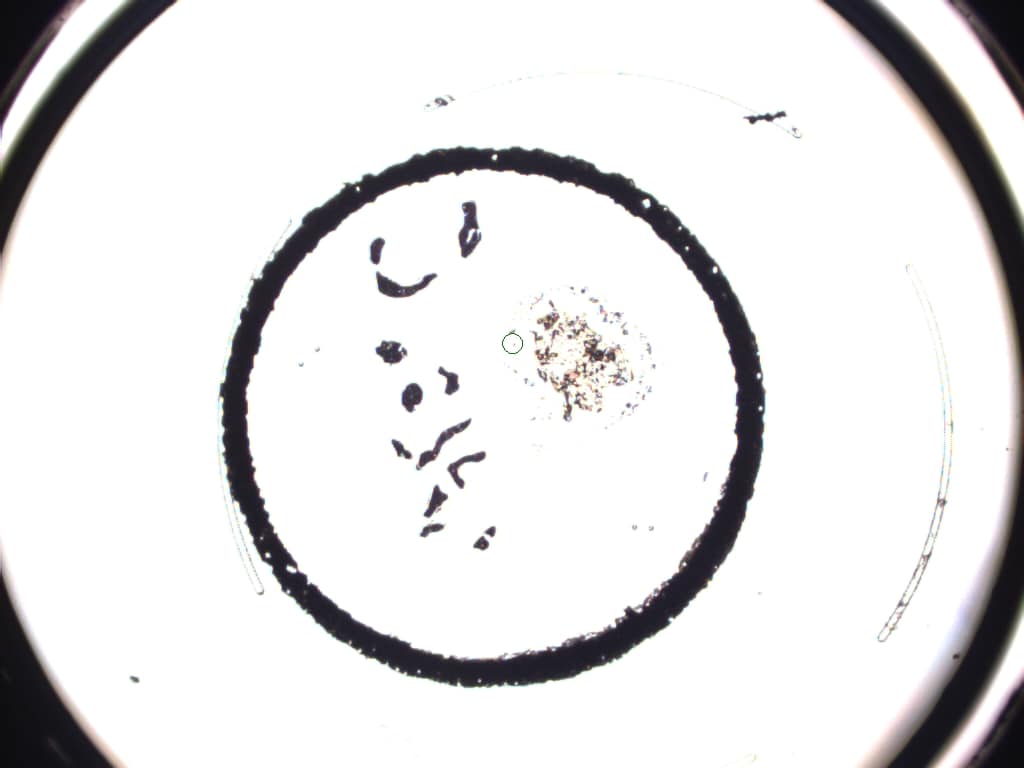

### DCIS2MAC2 BEFORE .jpeg

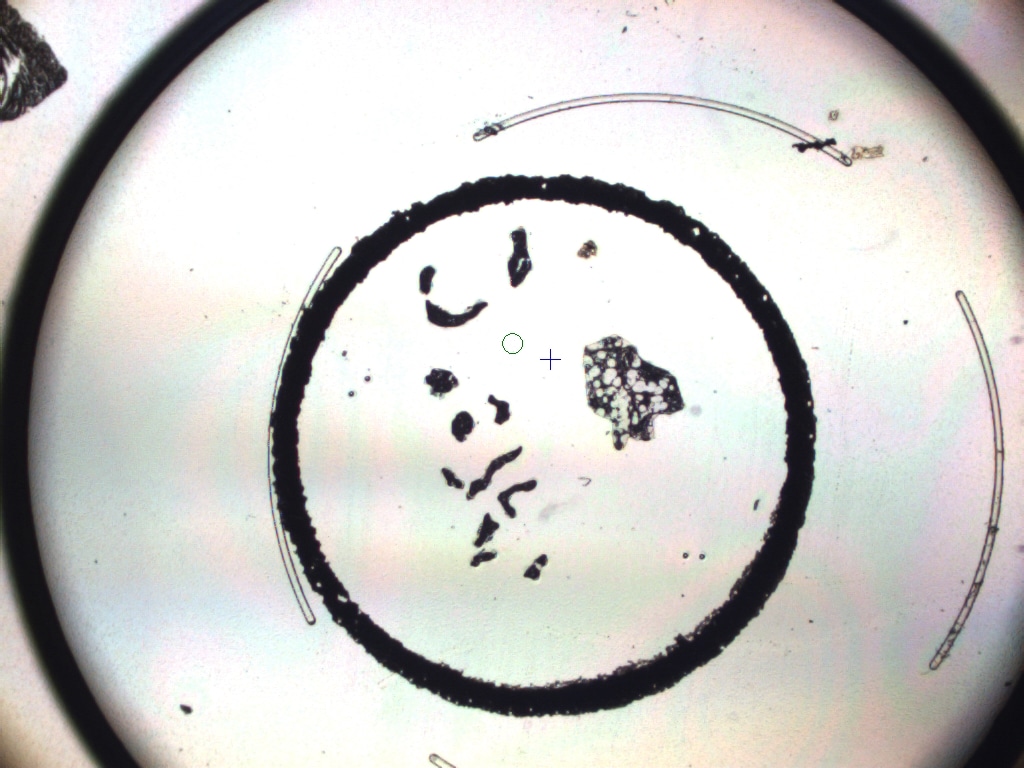

### DCIS3 BEFORE2X.jpeg

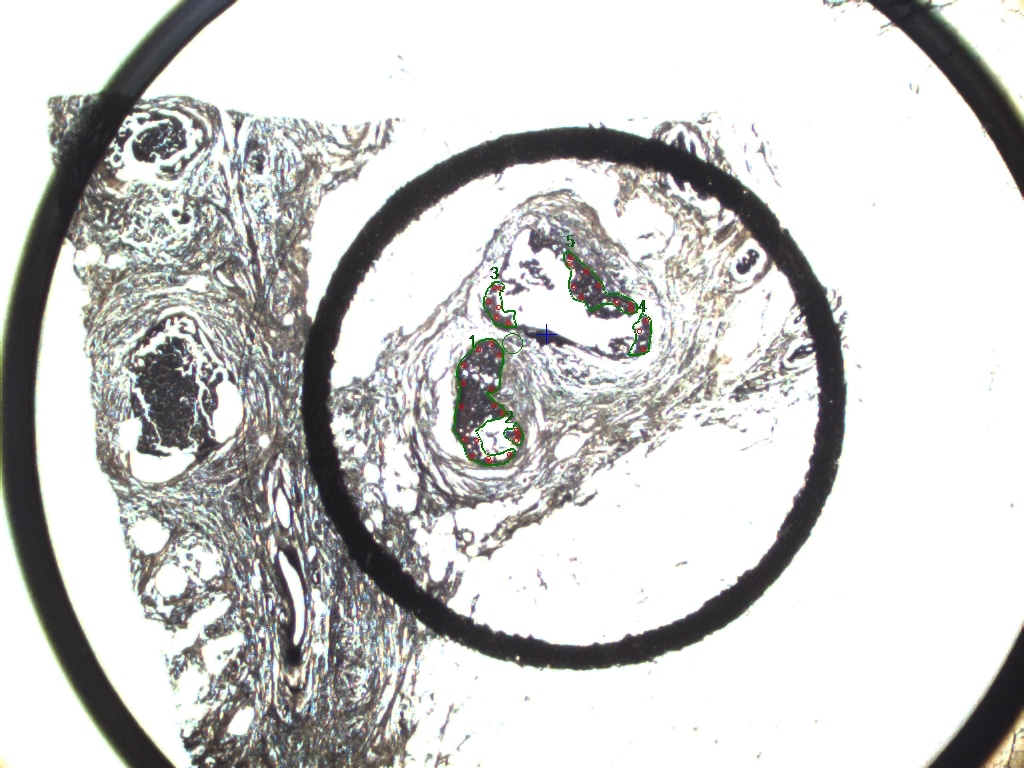

### DCIS3 BEFORE.jpeg

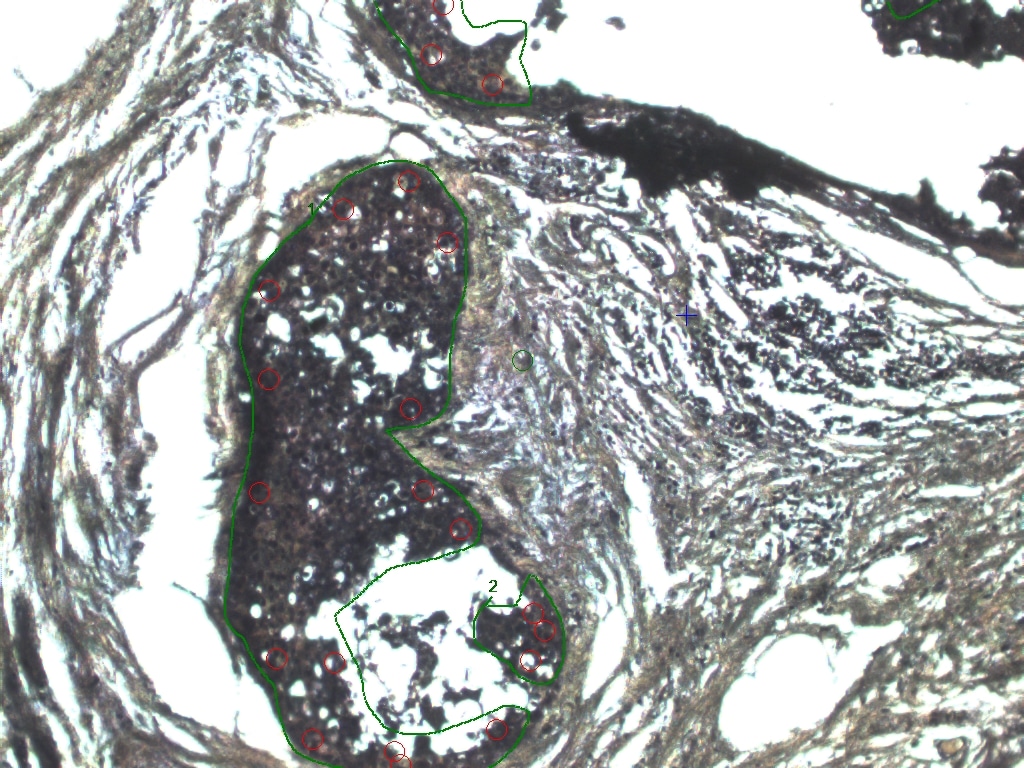

### DCIS3-2 BEFORE 10x.jpeg

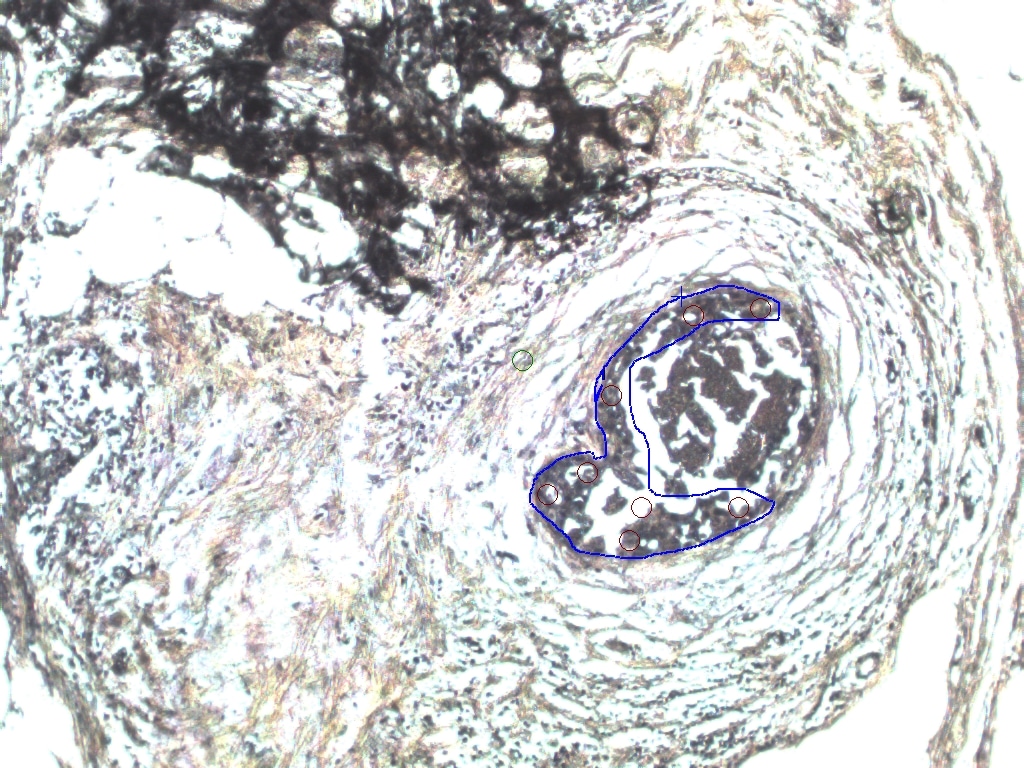

### DCIS3MAC3 ABLATION 10x .jpeg

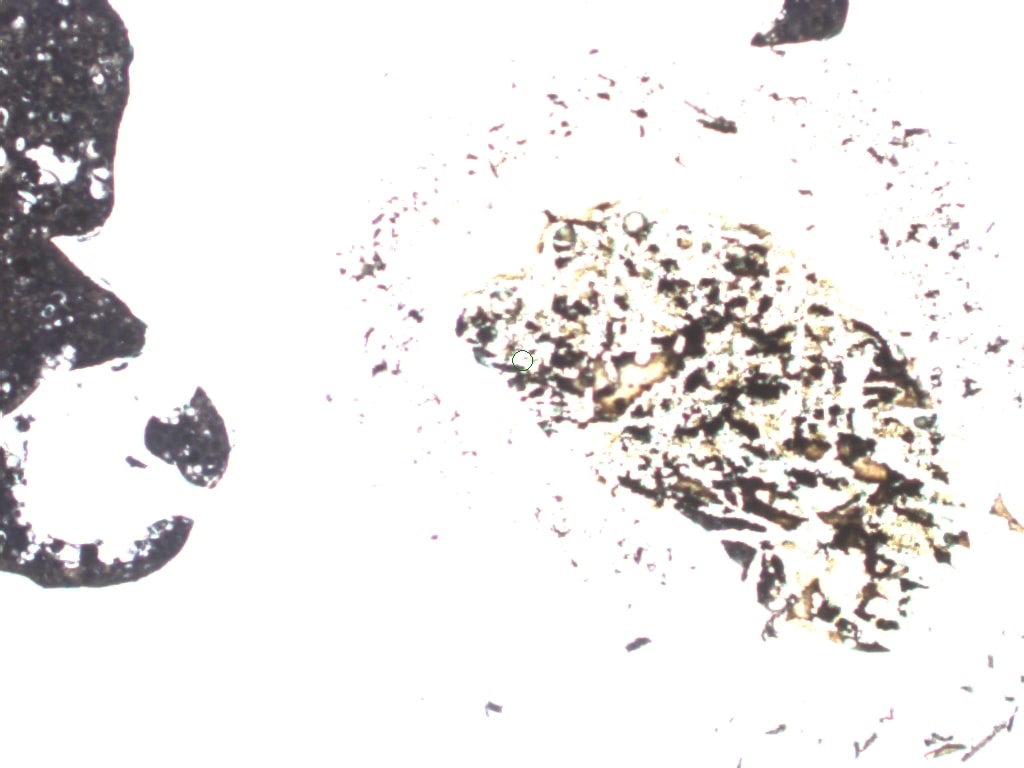

### DCIS3MAC3 ABLATION .jpeg

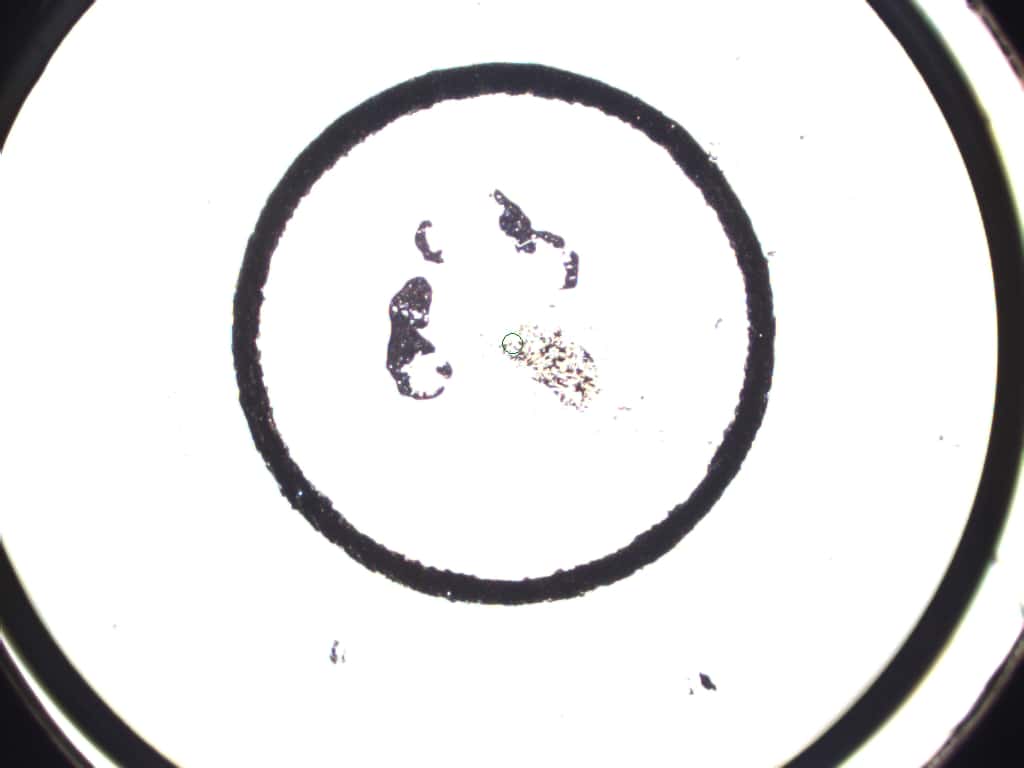

### DCIS3MAC3 BEFORE .jpeg

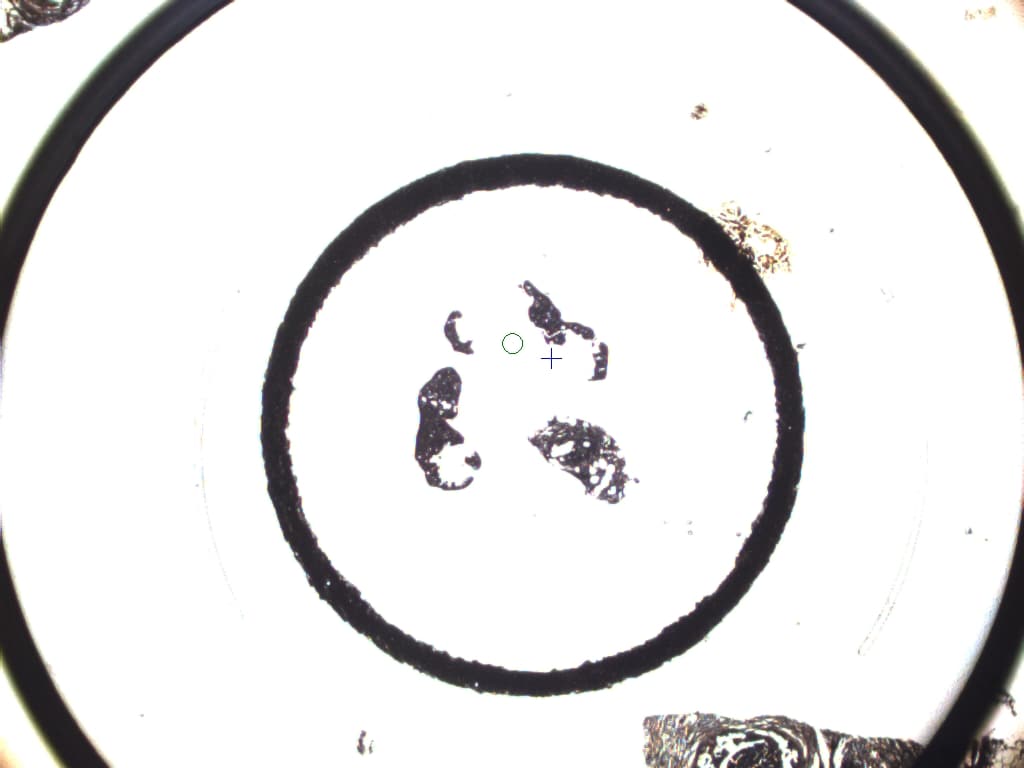

### DCIS 1 BEFORE2.jpeg

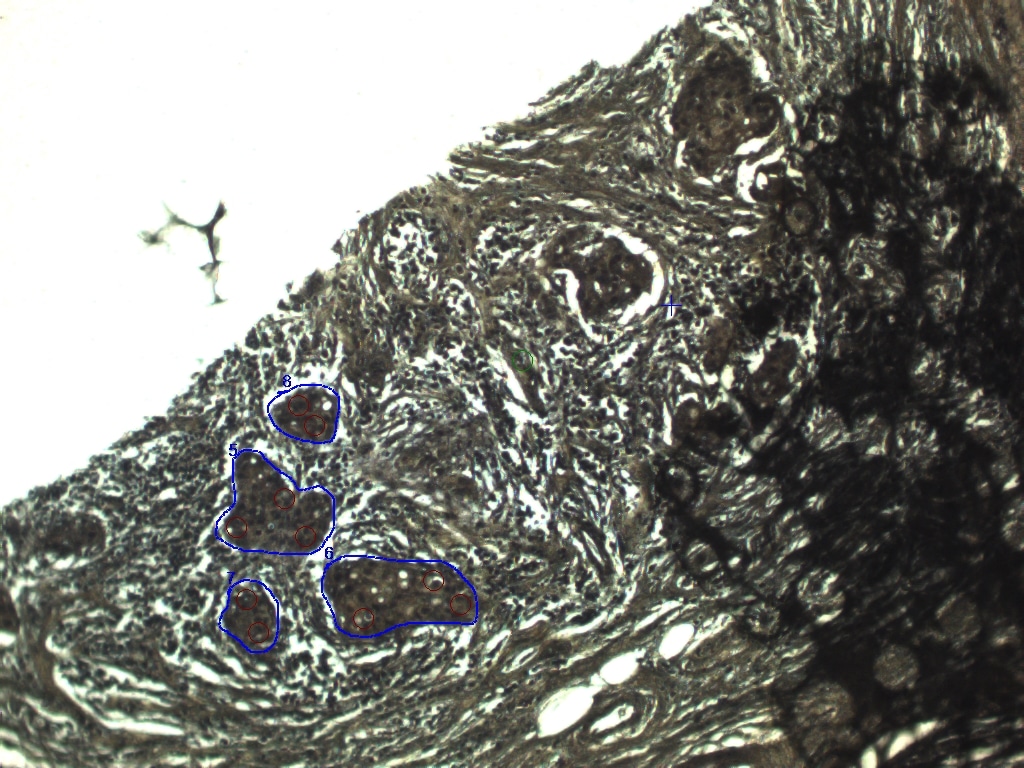

### DCIS 1 BEFORE.jpeg

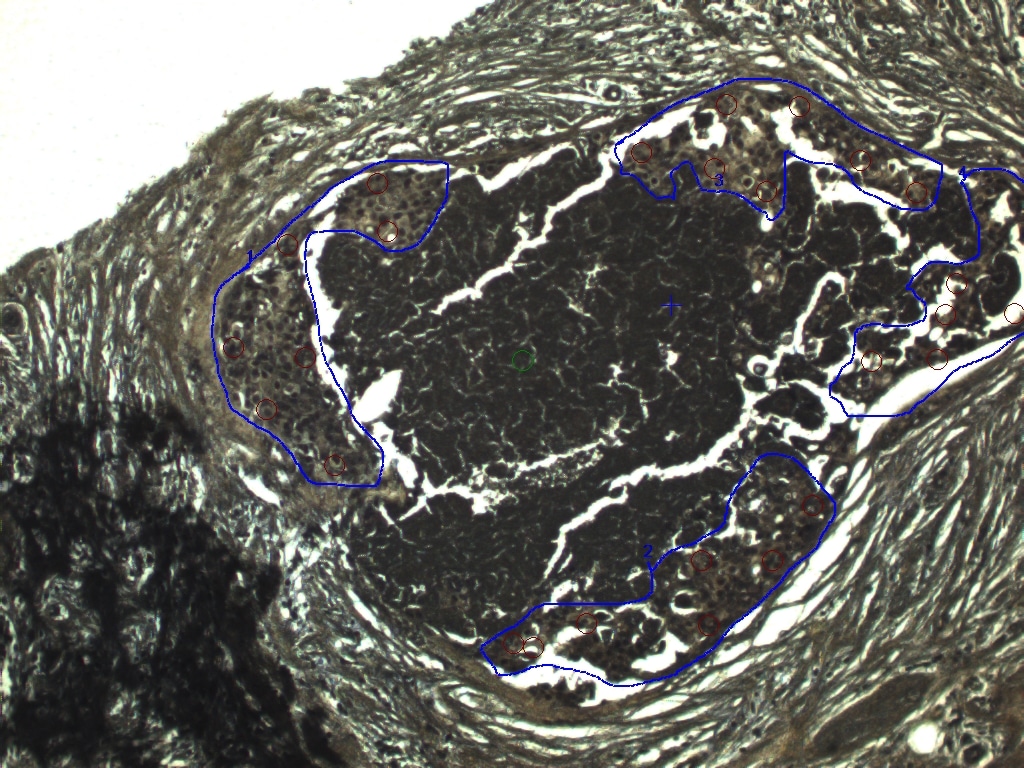

### DCIS 1AFTER.jpeg

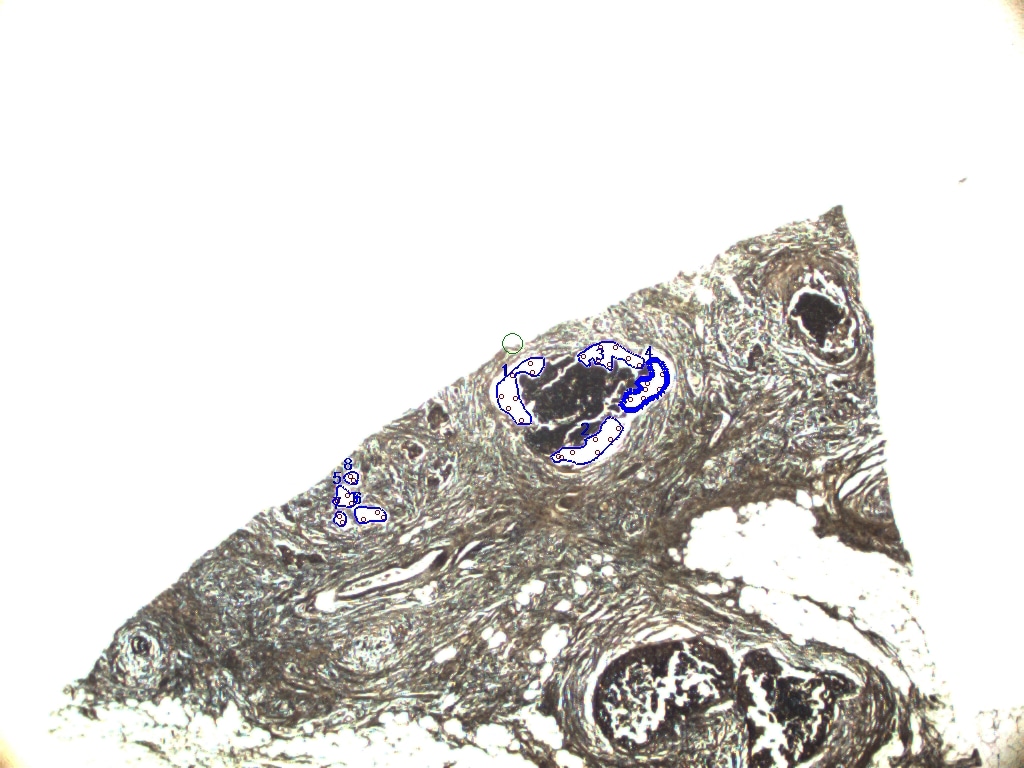

### DCIS-1 BEFORE.jpeg

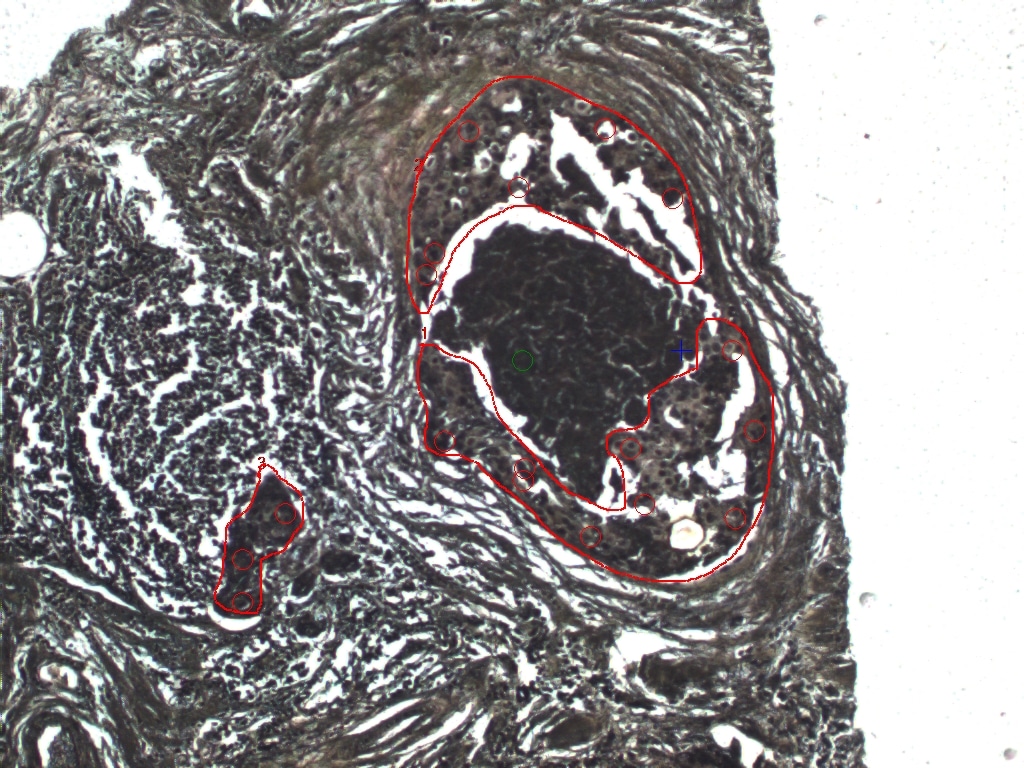

### DCIS-1-2 BEFORE.jpeg

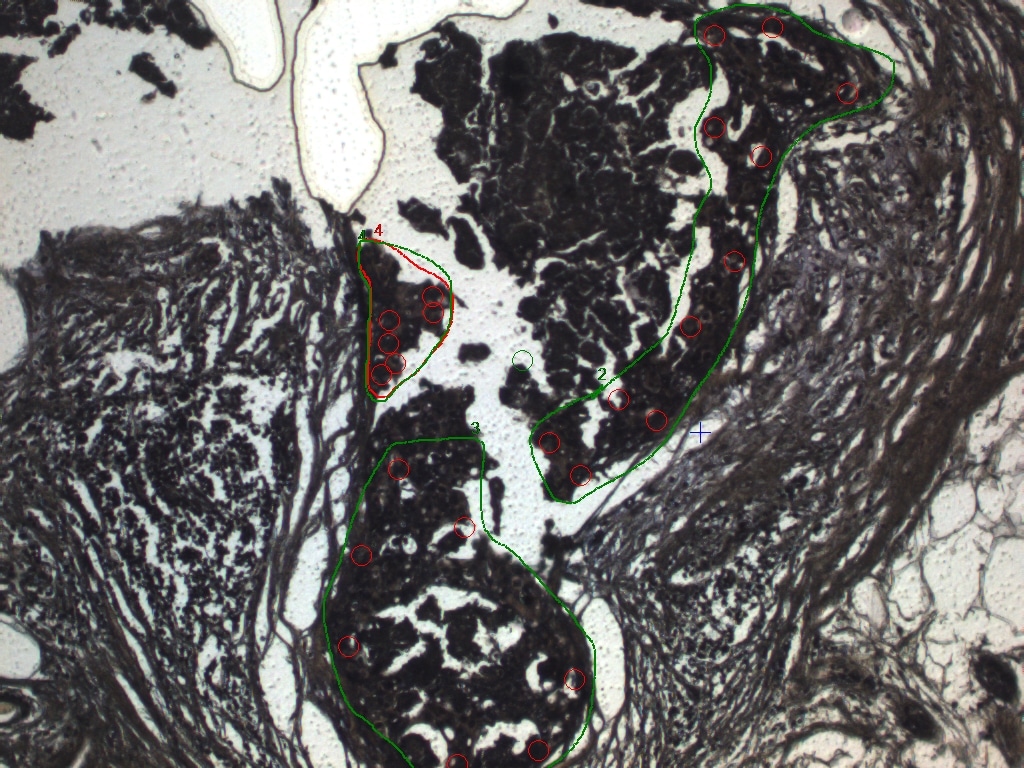

### DCIS-1-2AFTER.jpeg

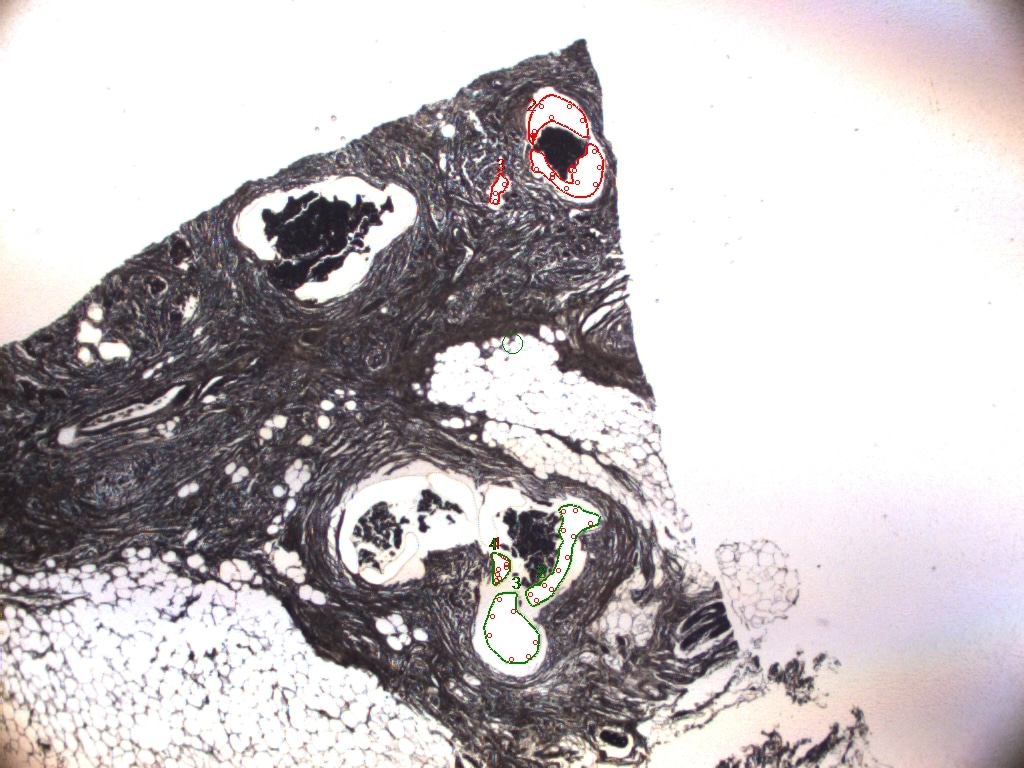

### DCIS-1.jpeg

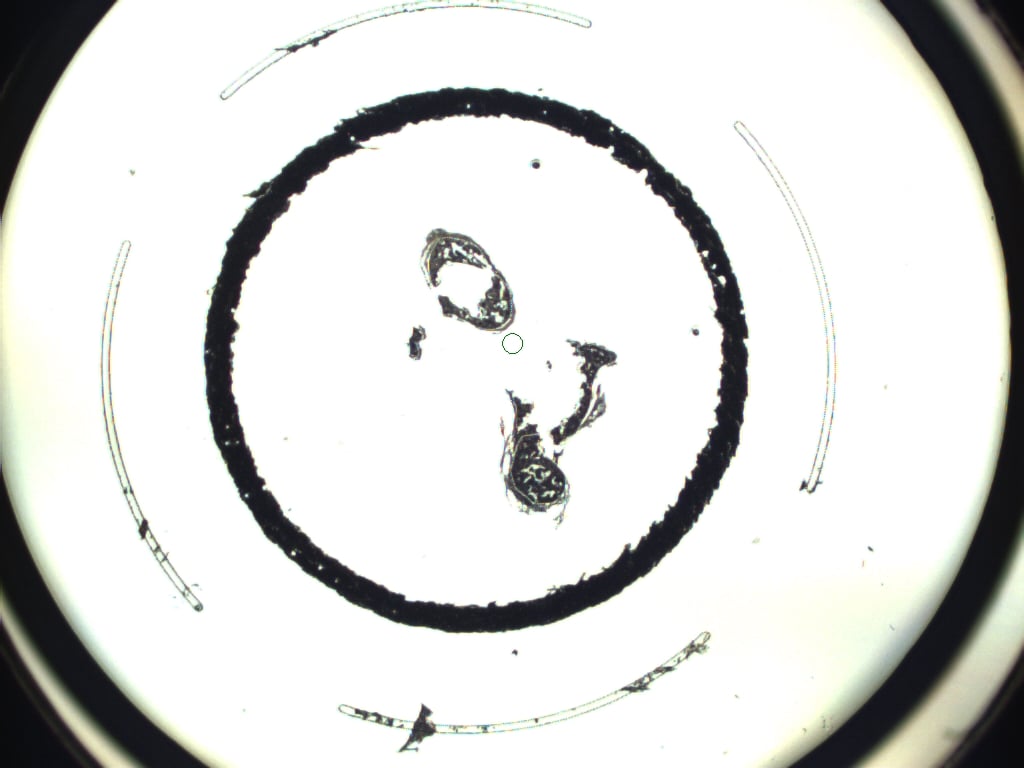

### DCIS-2 40x.jpeg

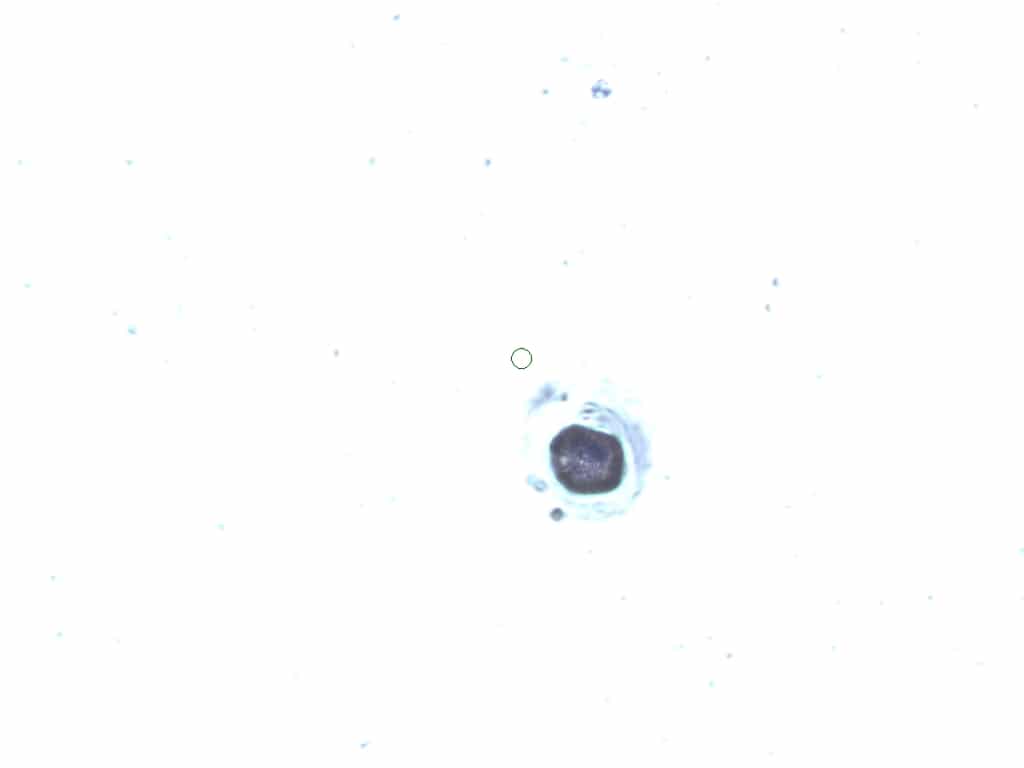

### DCIS-2 AFTER2X.jpeg

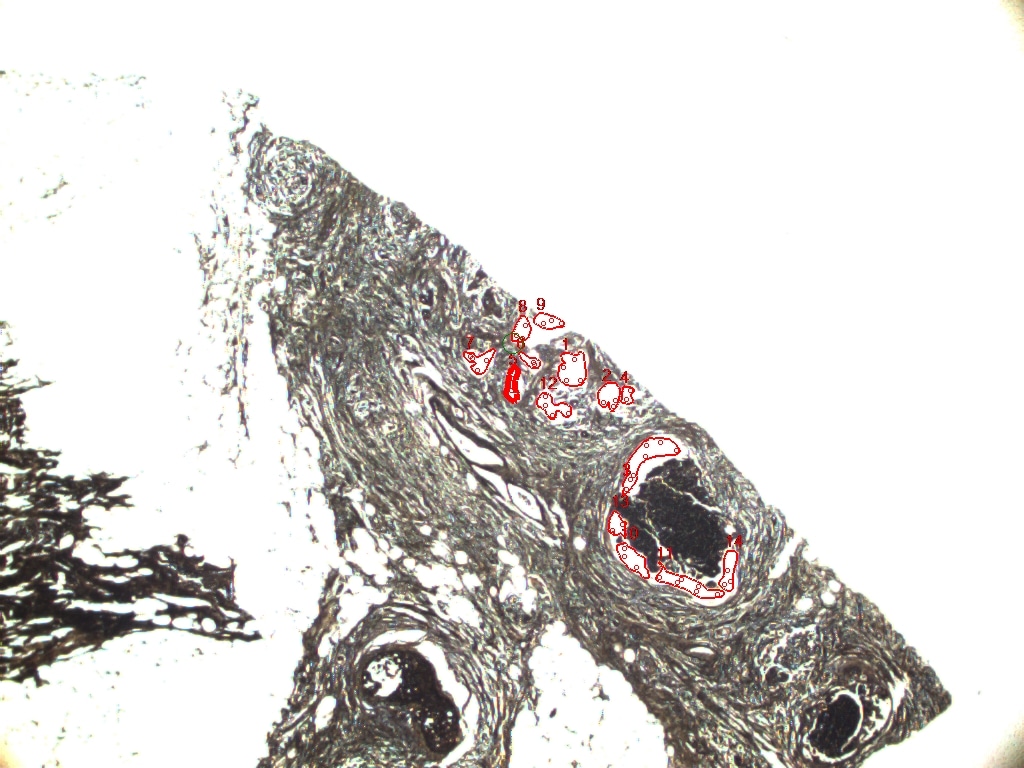

### DCIS-2 AFTER.jpeg

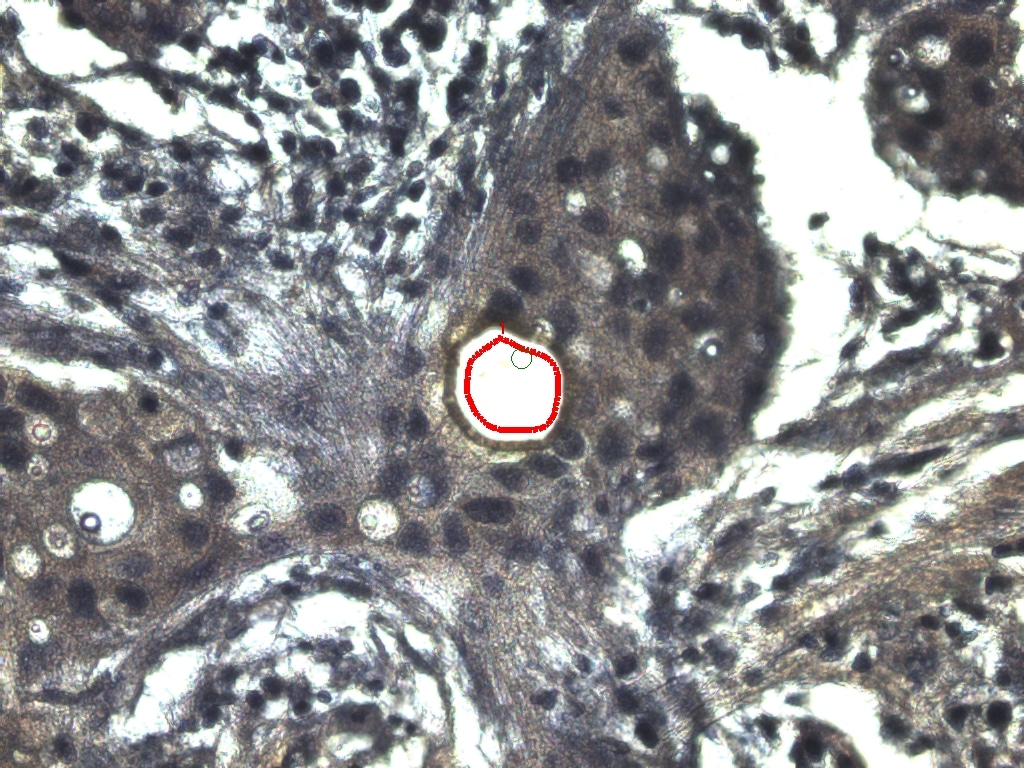

### DCIS-2 BEFORE2X.jpeg

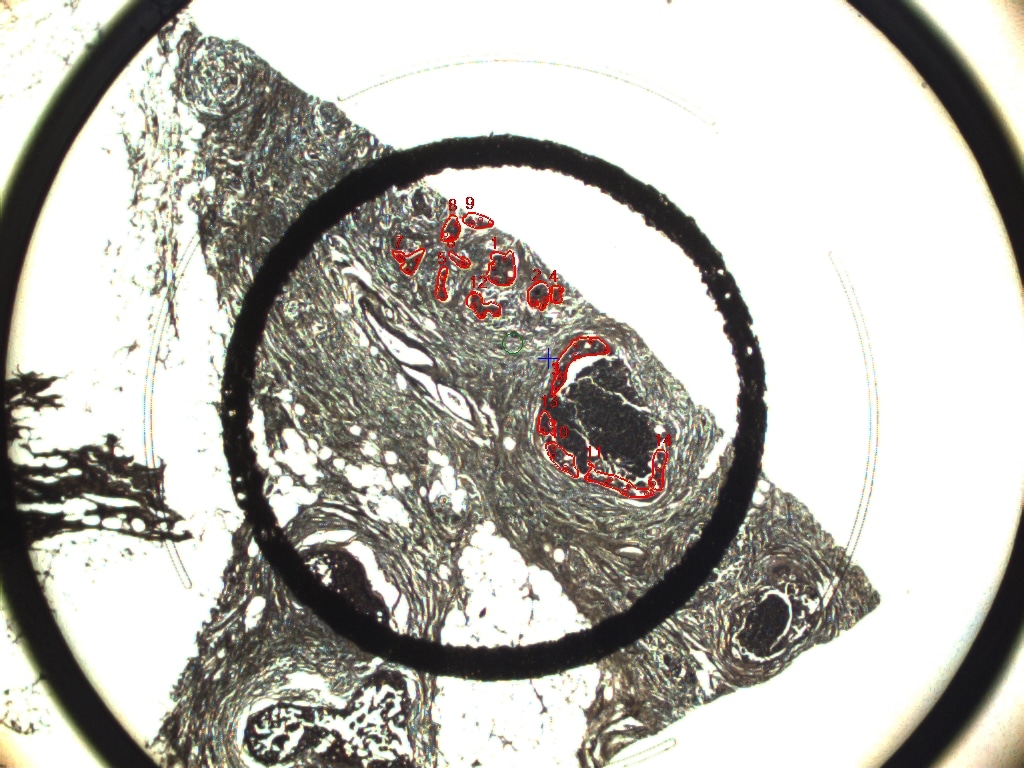

### DCIS-2 BEFORE.jpeg

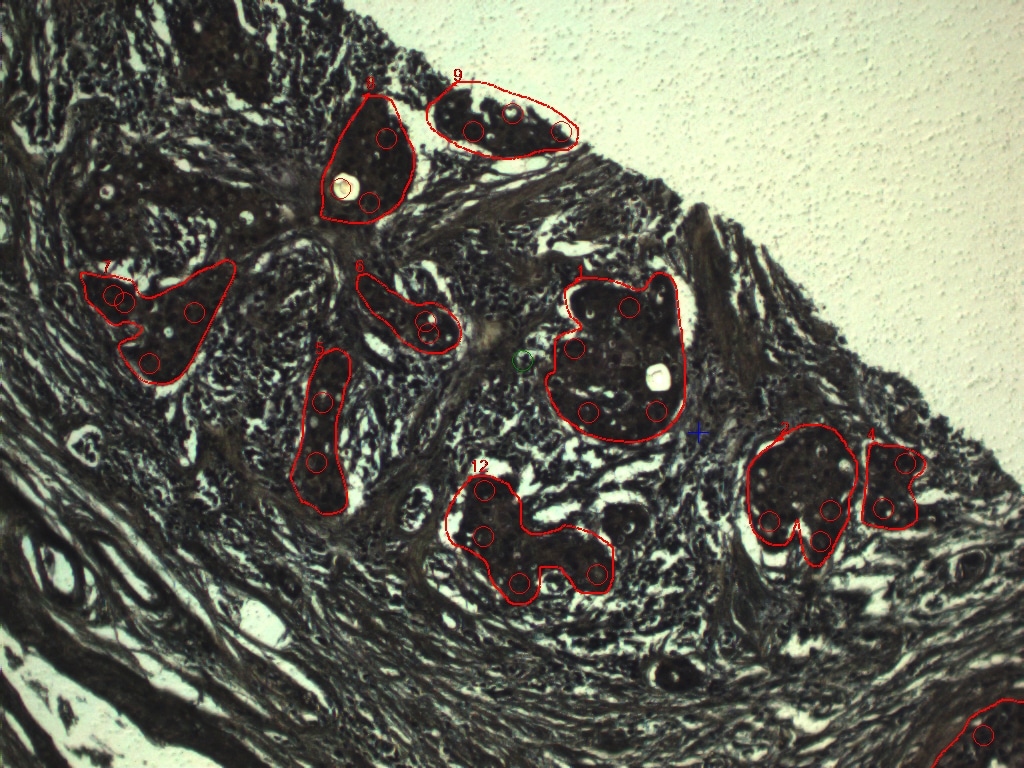

### DCIS-2.jpeg

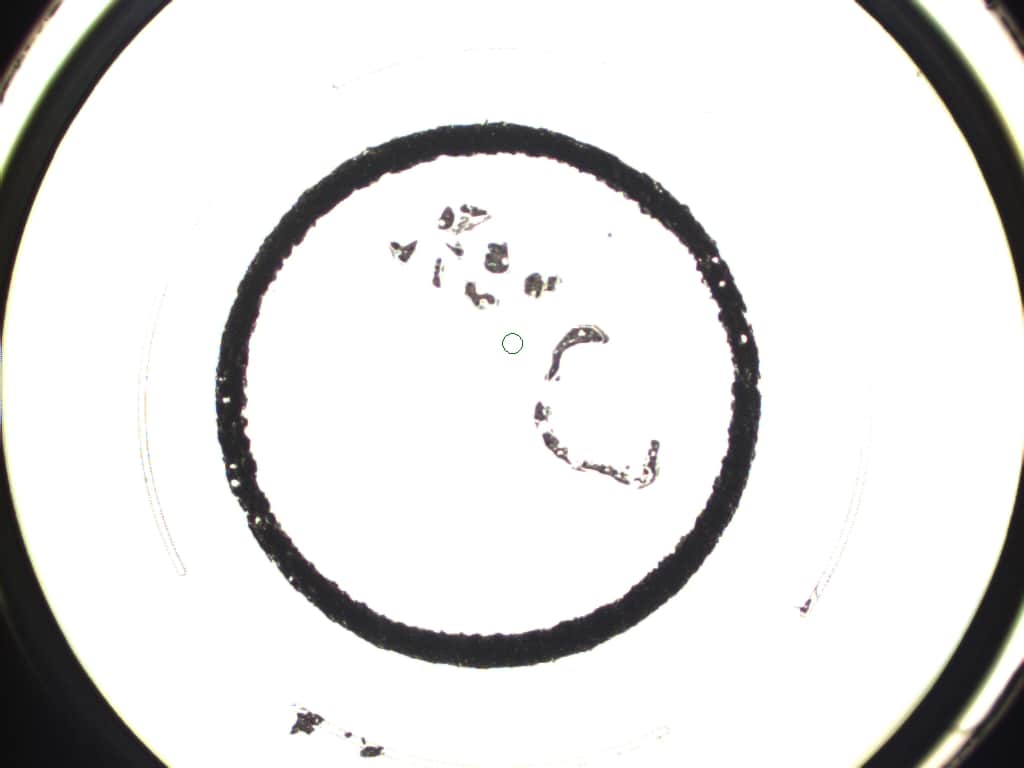

### DCIS-3 AFTER.jpeg

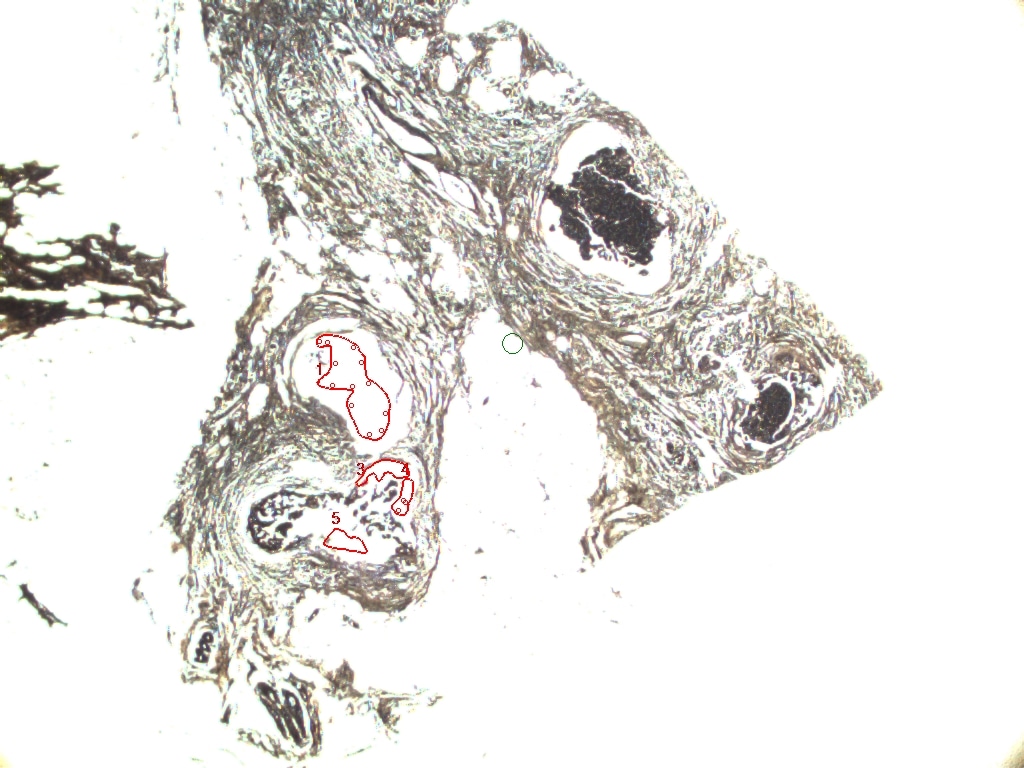

### DCIS-3 AFTER.jpeg

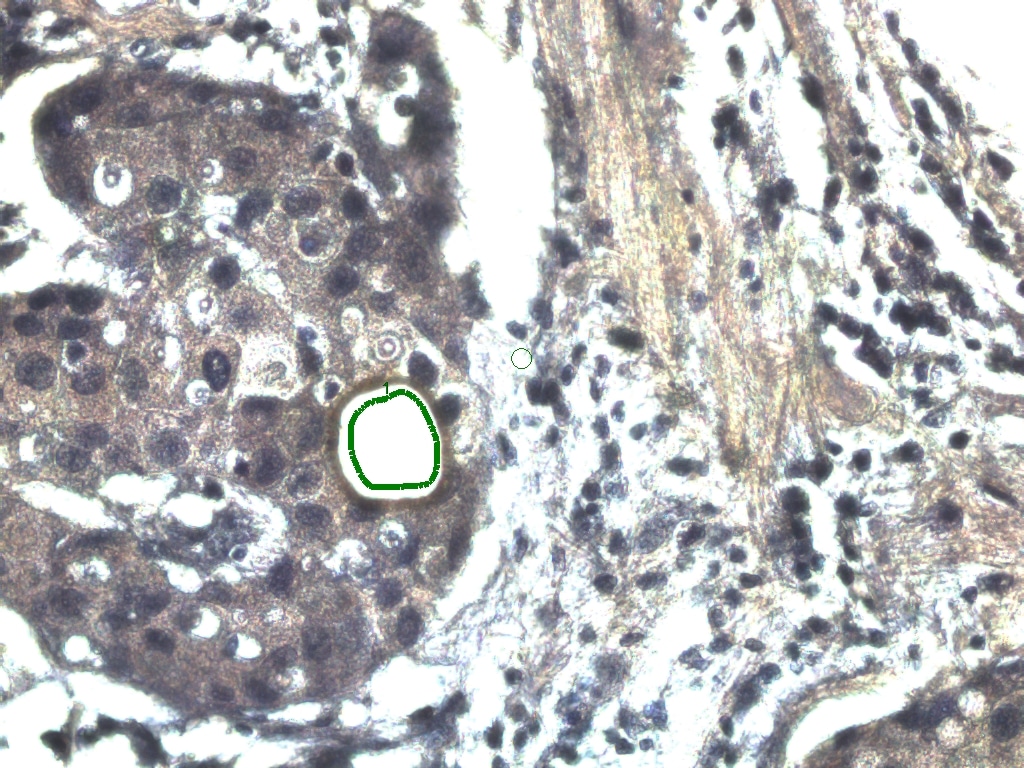

### DCIS-3 BEFORE.jpeg

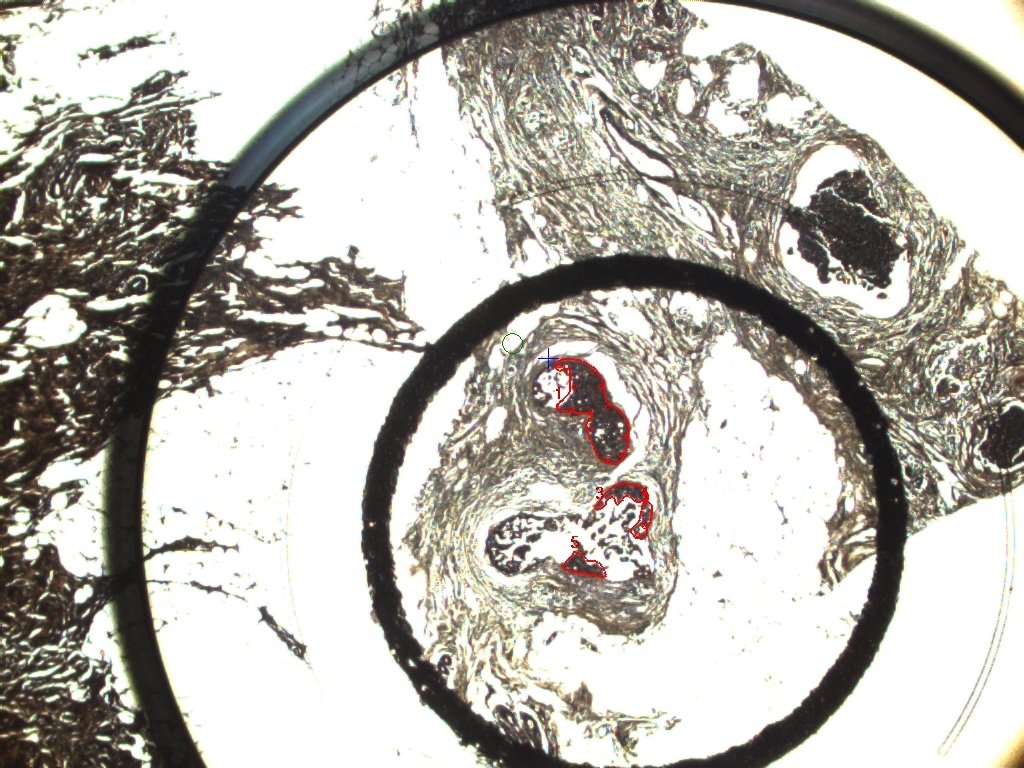

### DCIS-3 BEFORE.jpeg

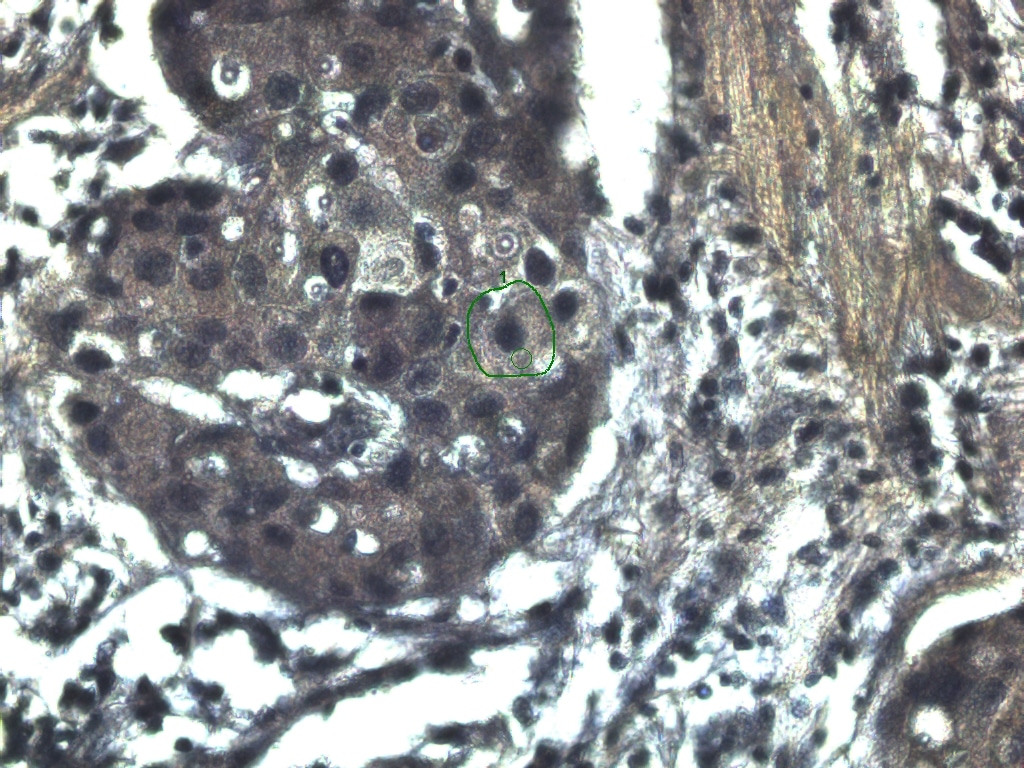

### DCIS-3.jpeg

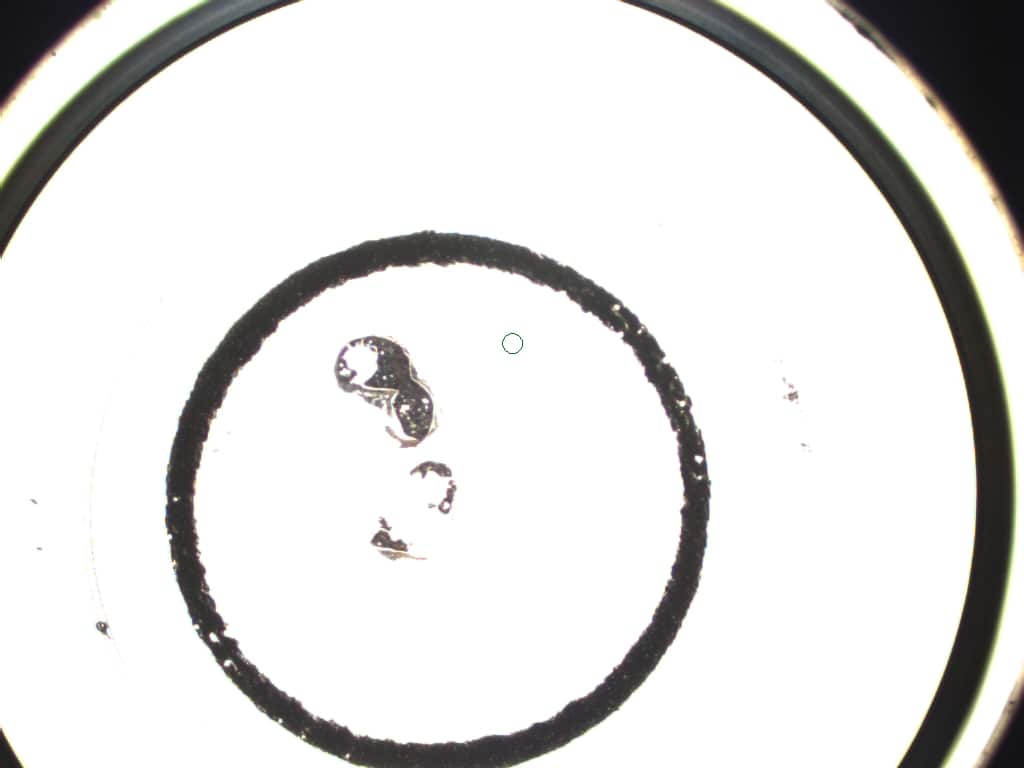

### DCIS-4 ABLATION.jpeg

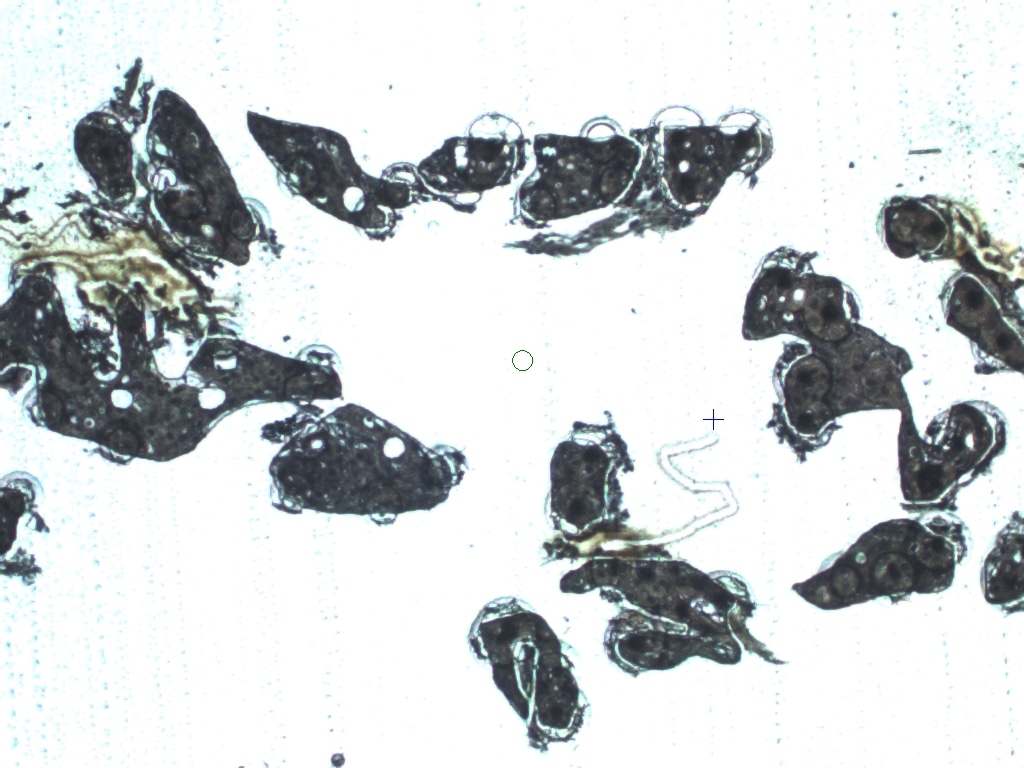

### DCIS-4 AFTER.jpeg

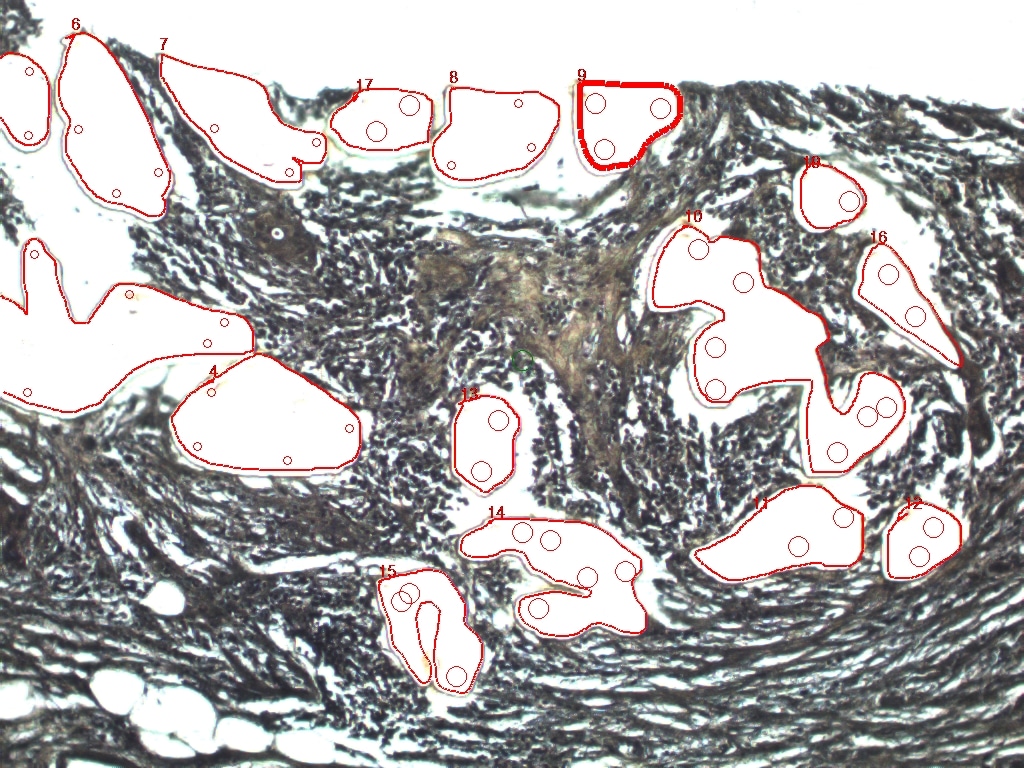

### DCIS-4 BEFORE .jpeg

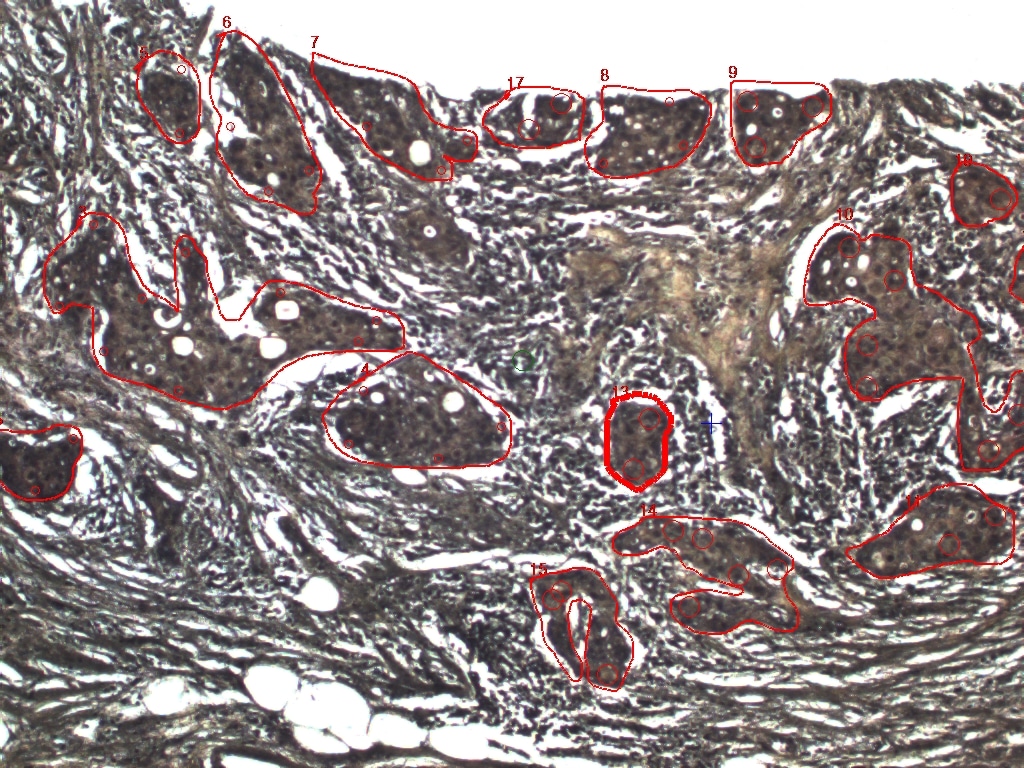

### DCIS-4 BEFORE.jpeg

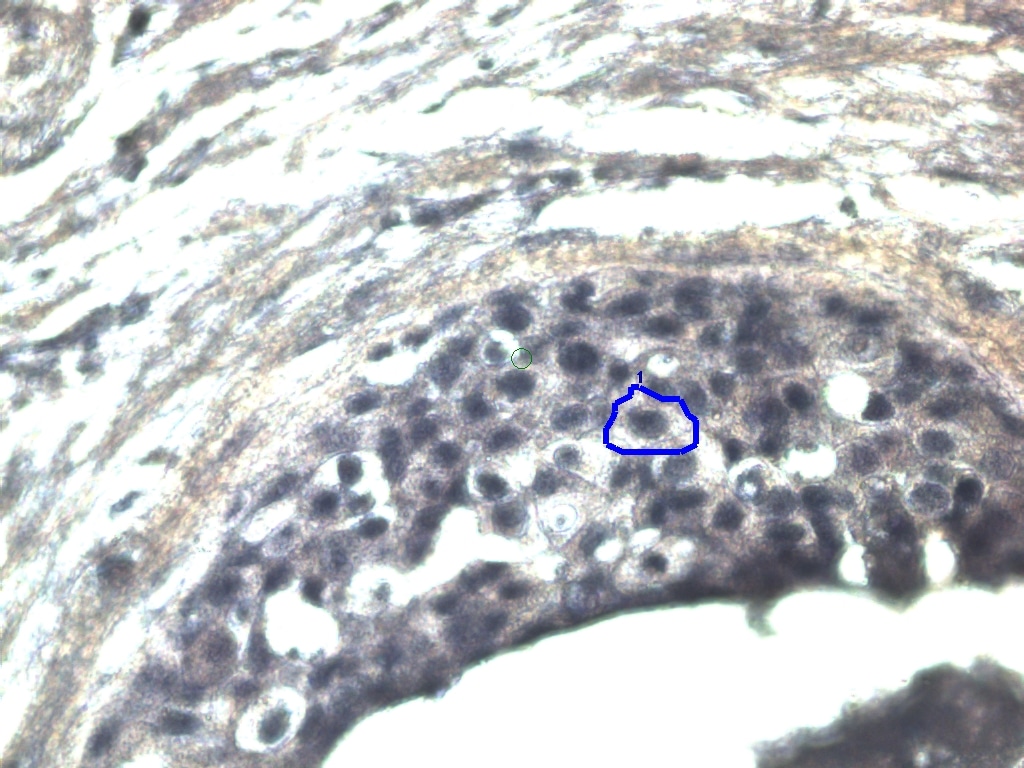

### DCIS-4.jpeg

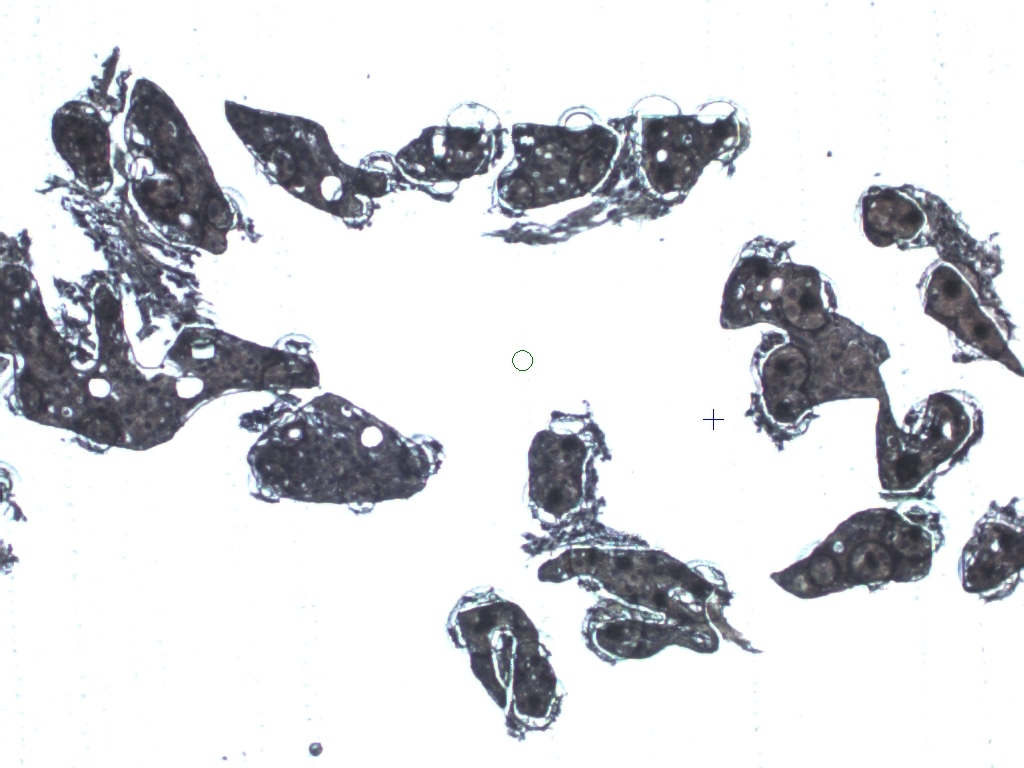

### DCIS-5 AFTER.jpeg

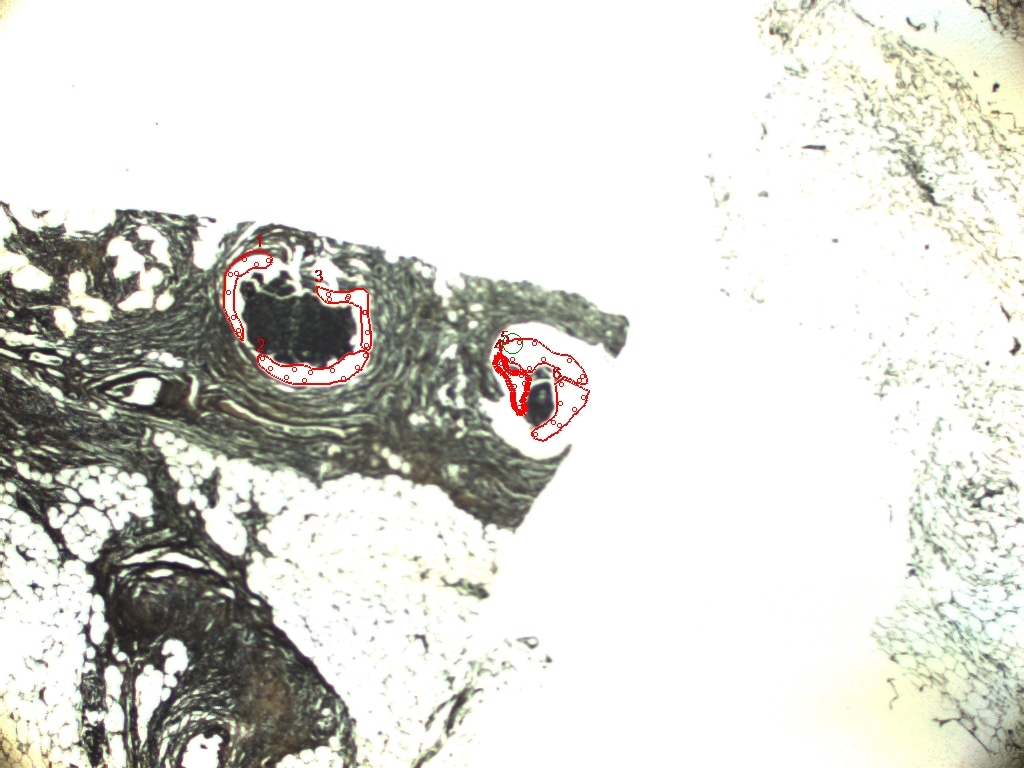

### DCIS-5 BEFORE2 .jpeg

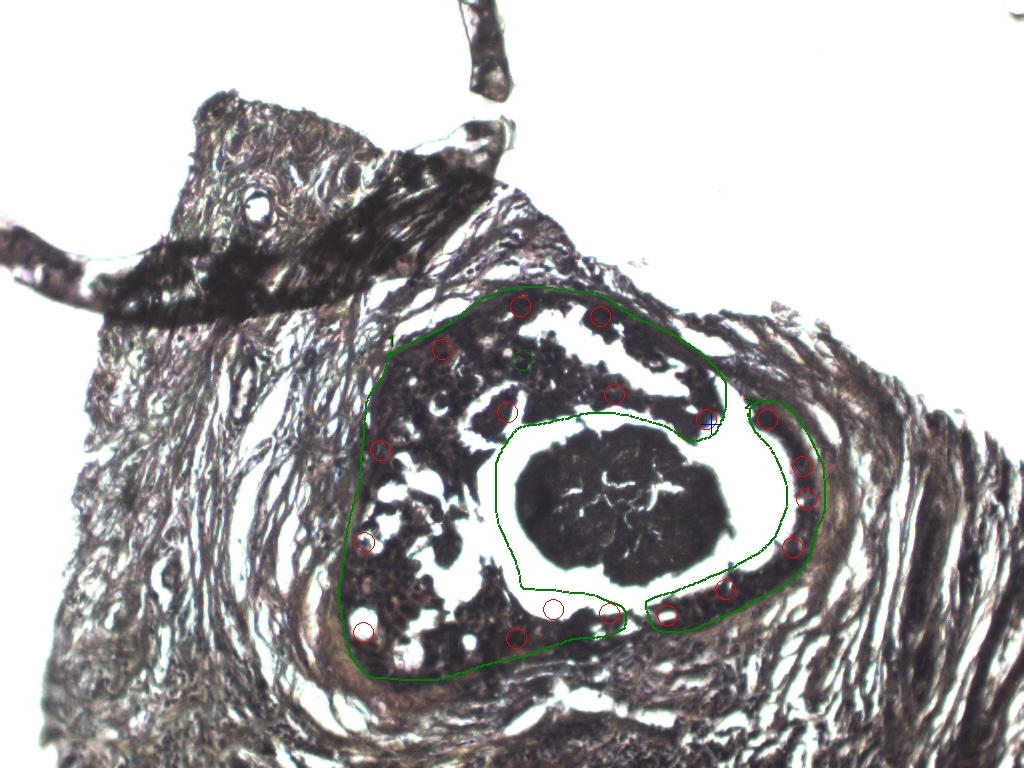

### DCIS-5 BEFORE .jpeg

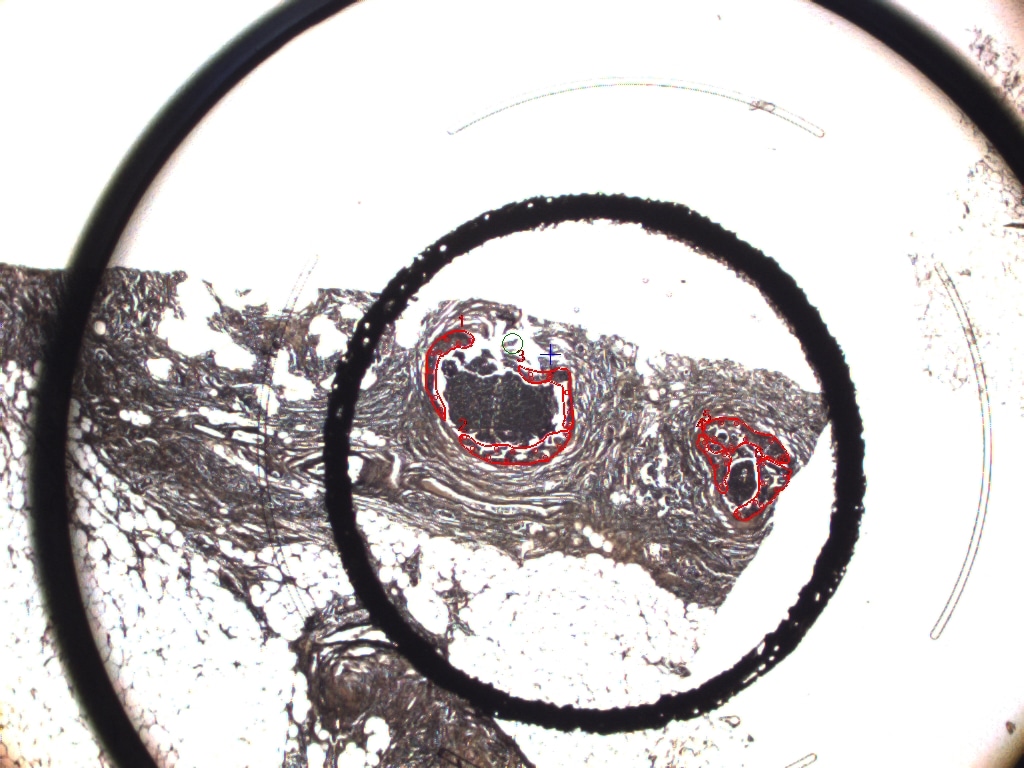

### DCIS-5.jpeg

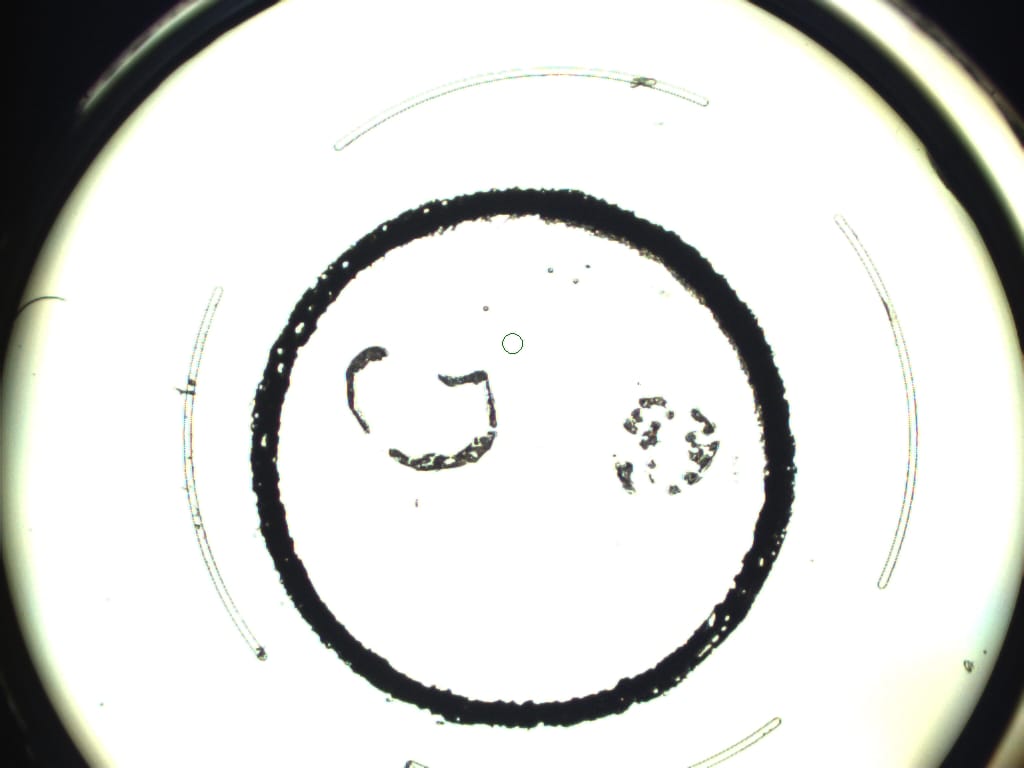

### DCIS-5FINAL.jpeg

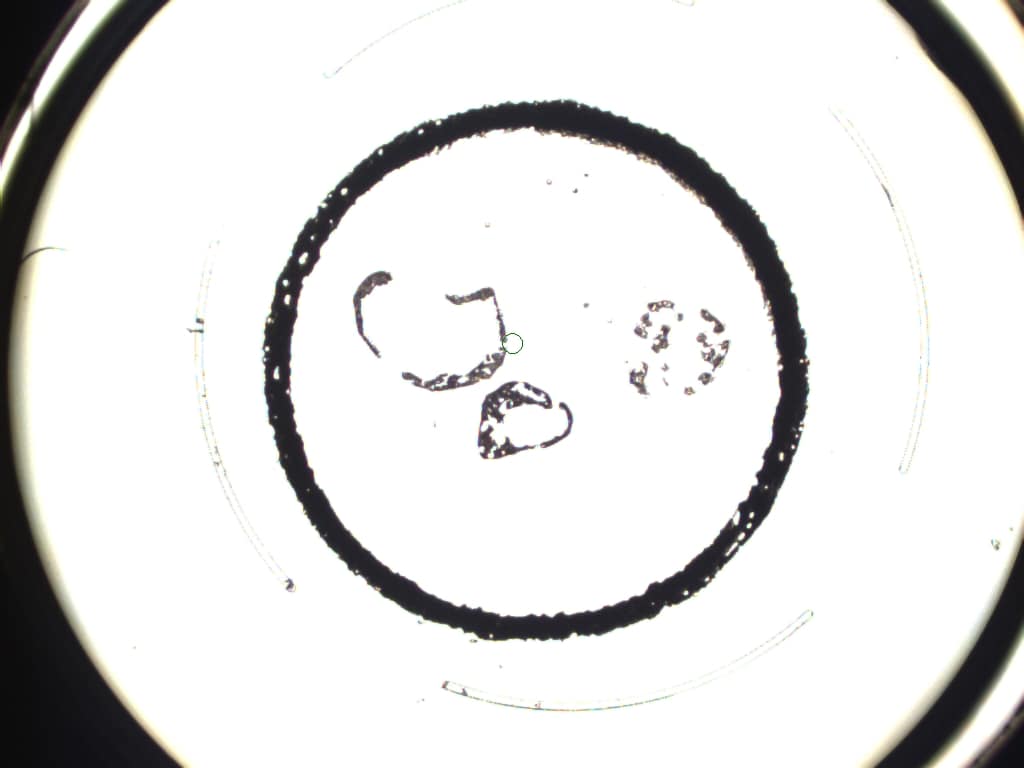

### DCIS-6 AFTER.jpeg

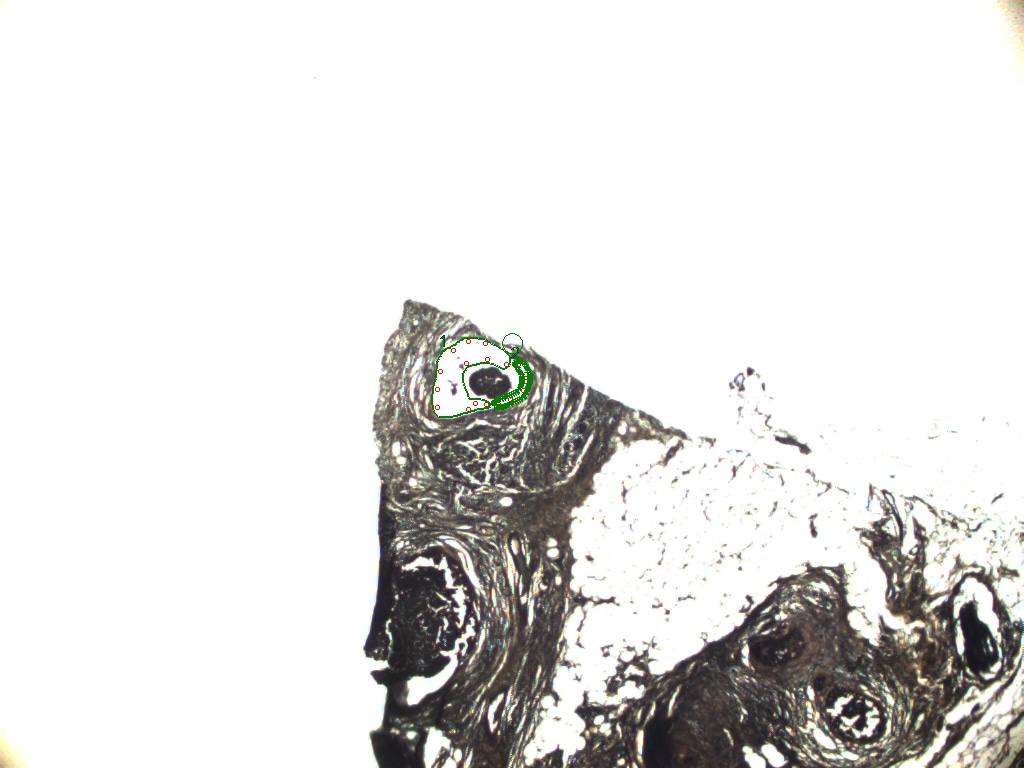

### DCIS-6 BEFORE2 .jpeg

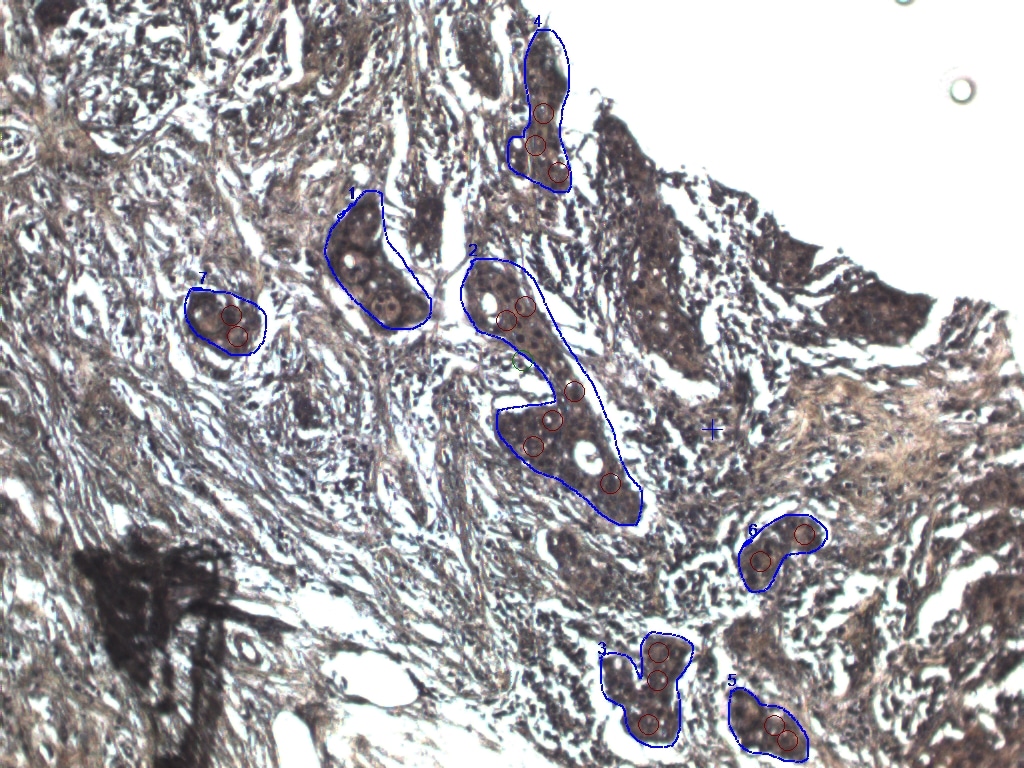

### DCIS-6 BEFORE .jpeg

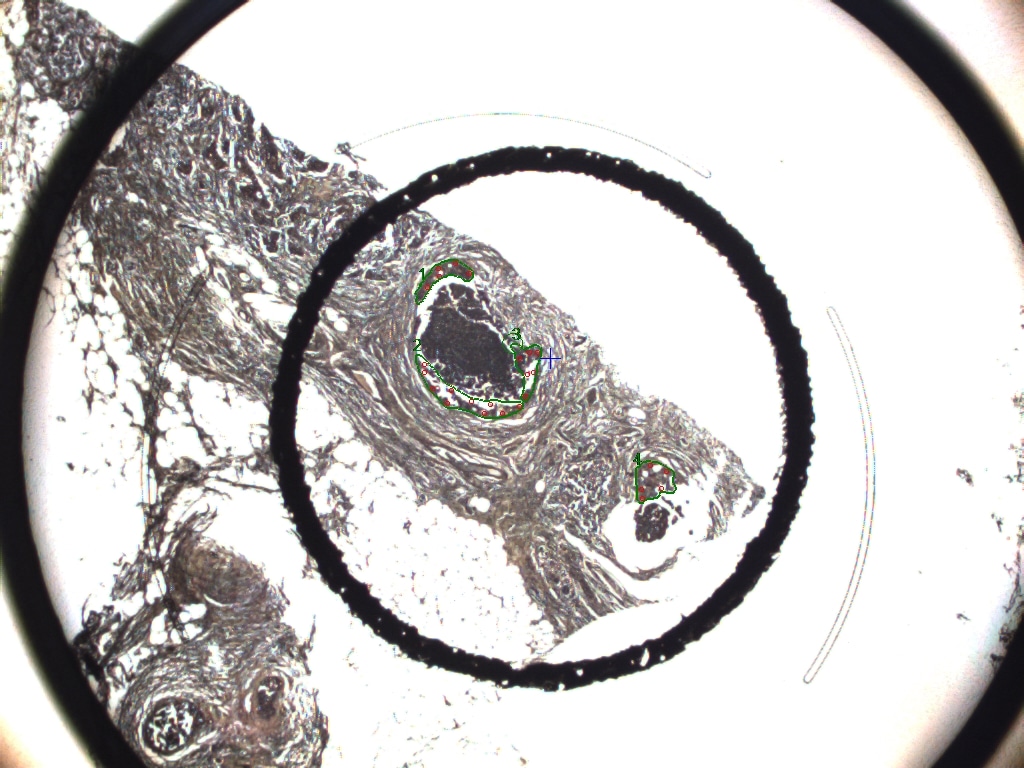

### DCIS-6 RIGHTAFTER2.jpeg

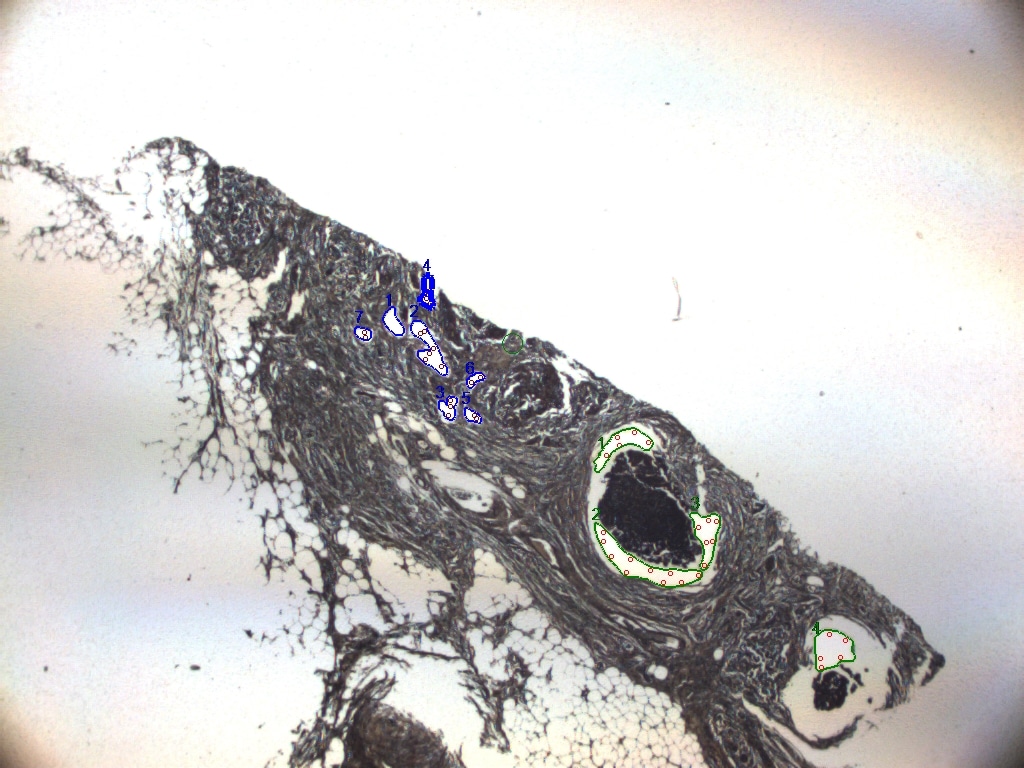

### DCIS-6 RIGHTAFTER.jpeg

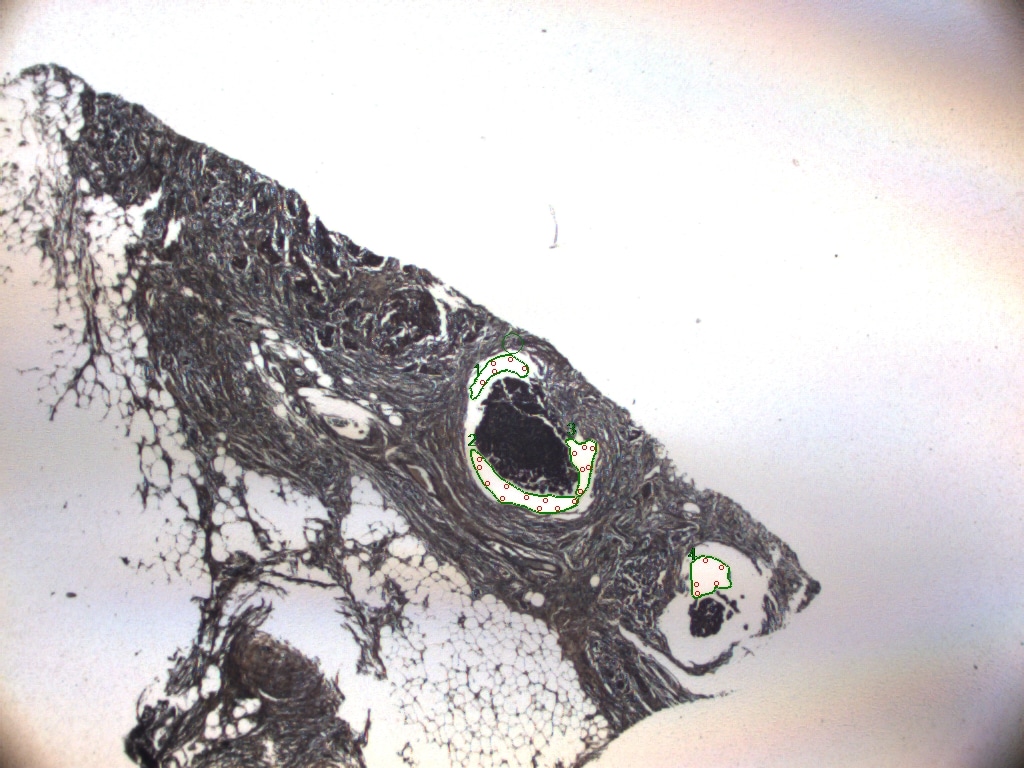

### DCIS-6.jpeg

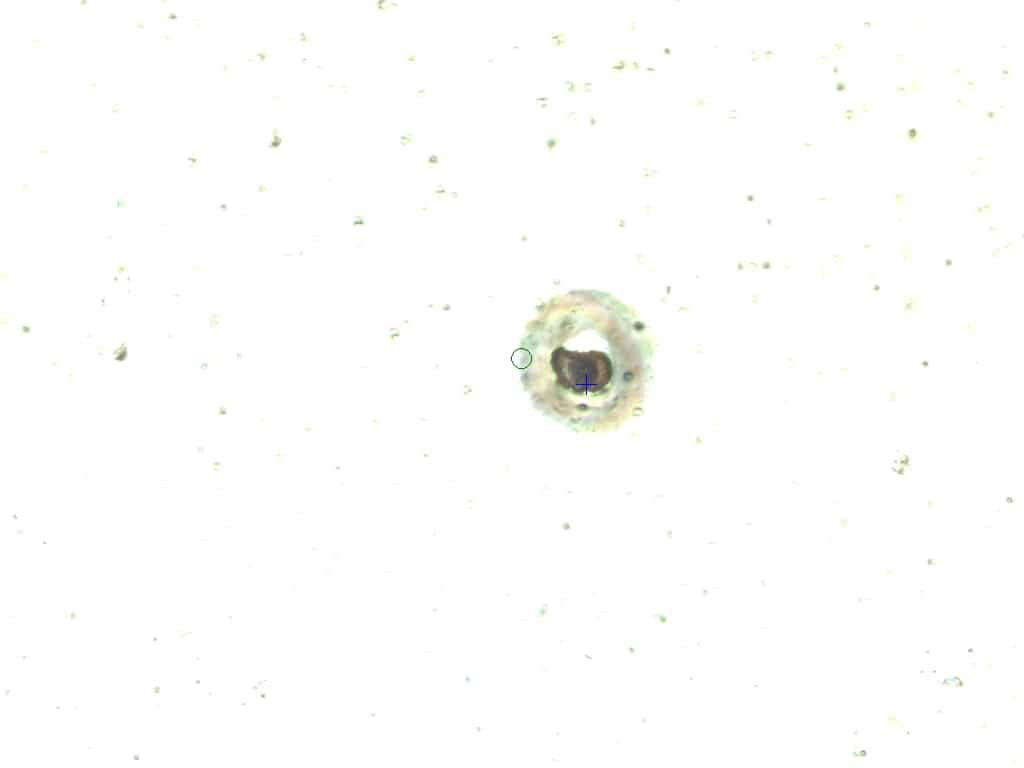

### DCIS-6FINAL.jpeg

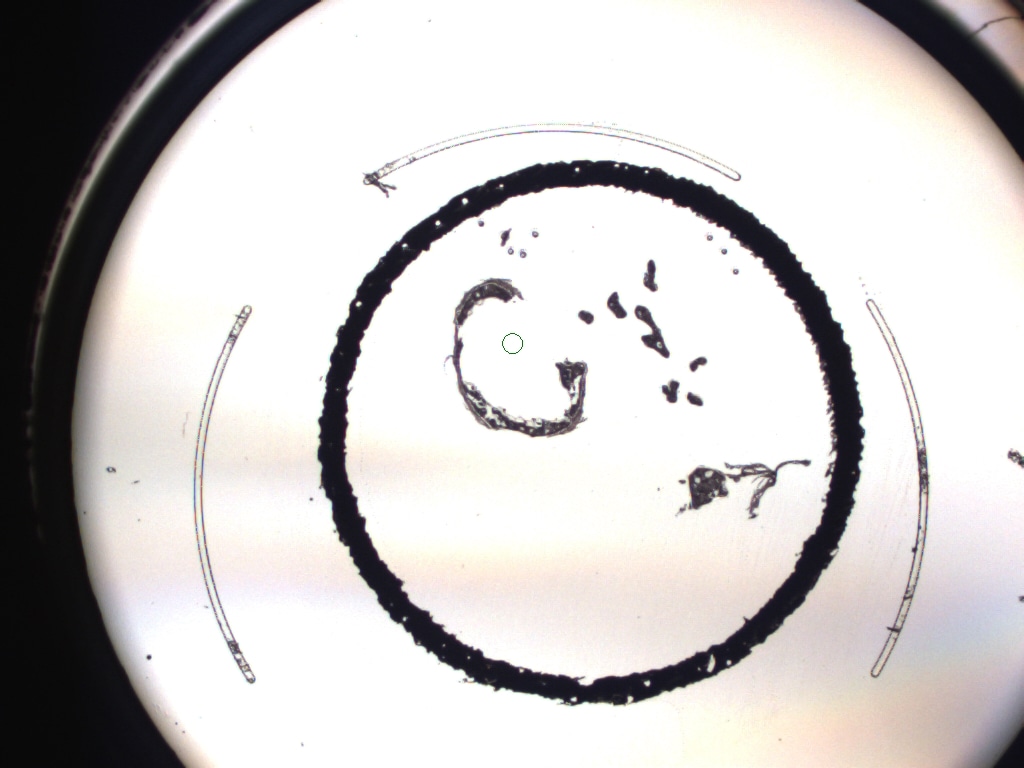

### DCIS-7 .jpeg

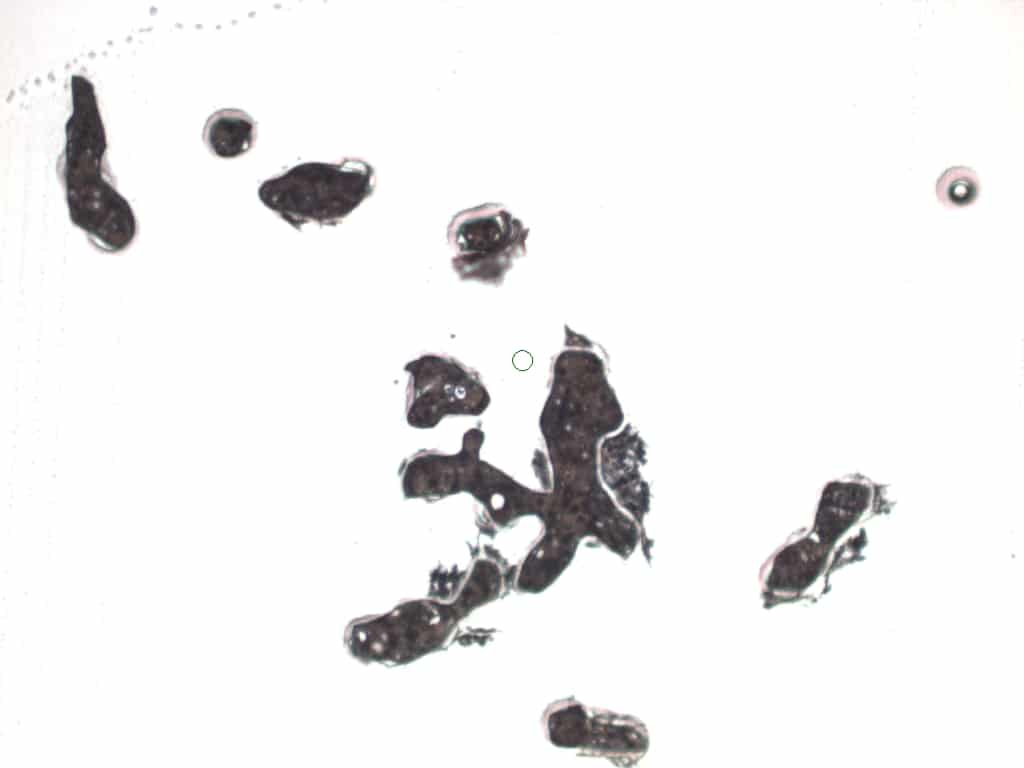

### DCIS-7 AFTER.jpeg

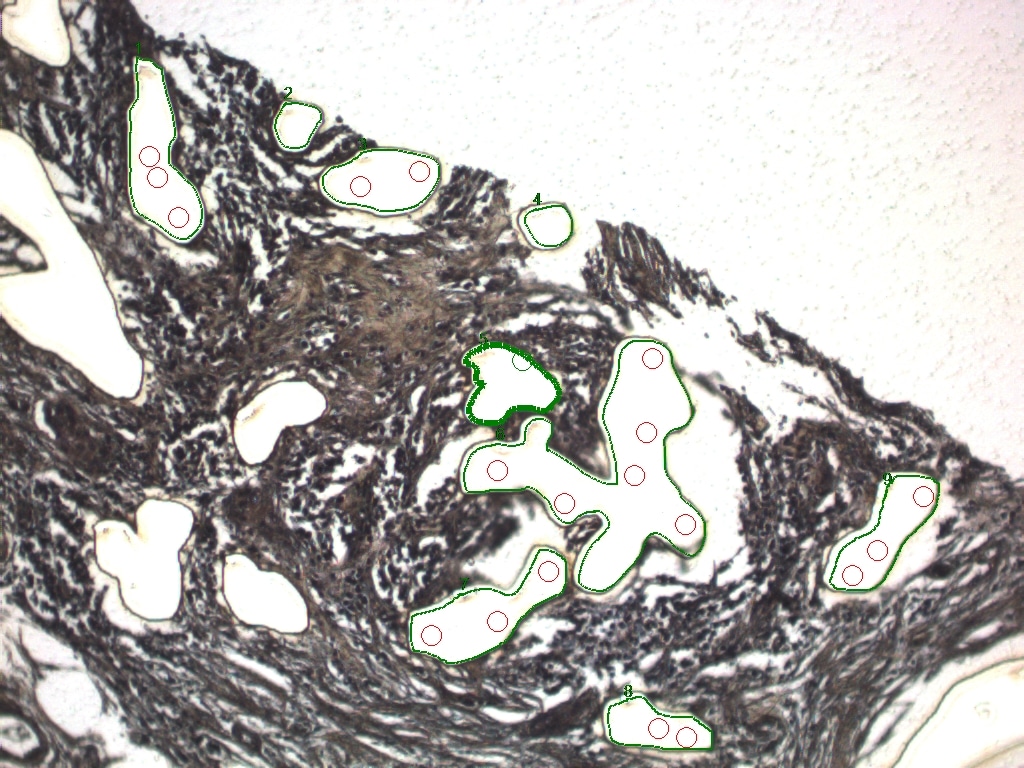

### DCIS-7 BEFORE .jpeg

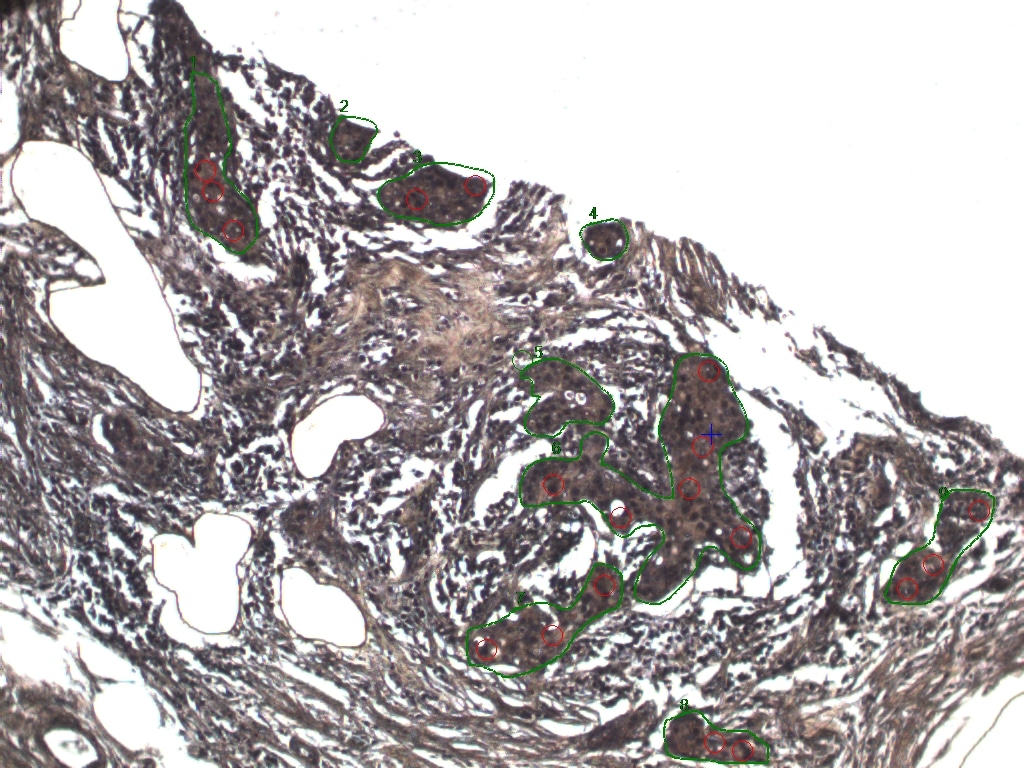

### DCIS-9 BEFORE.jpeg

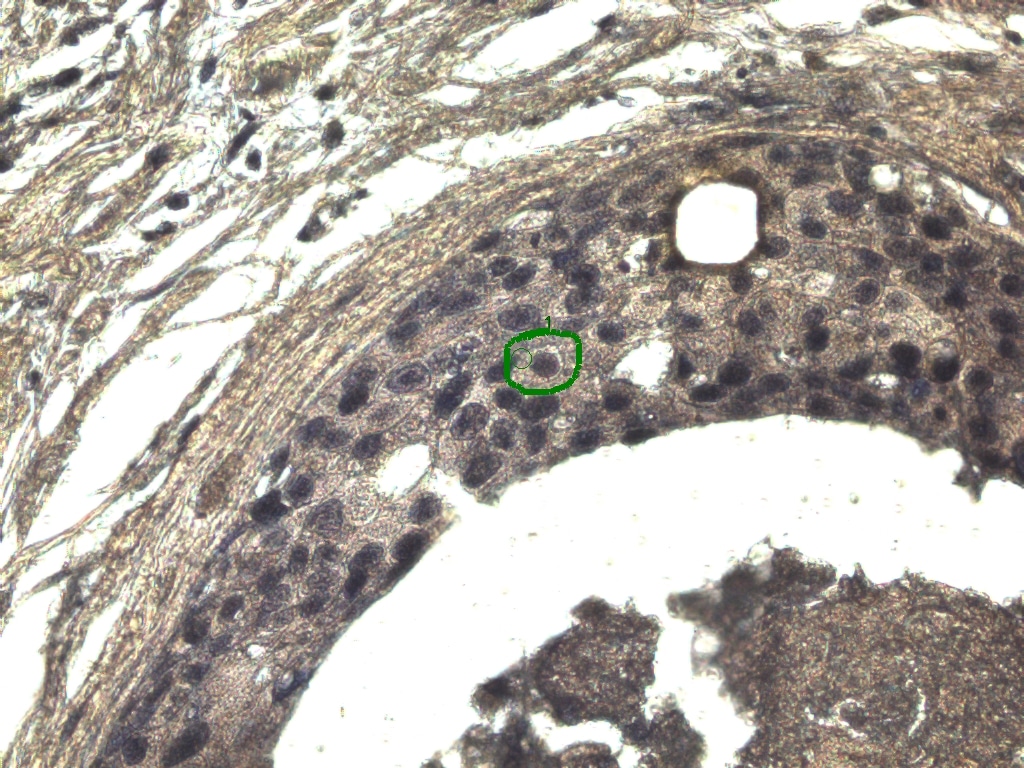

### IDC1 BEFORE.jpeg

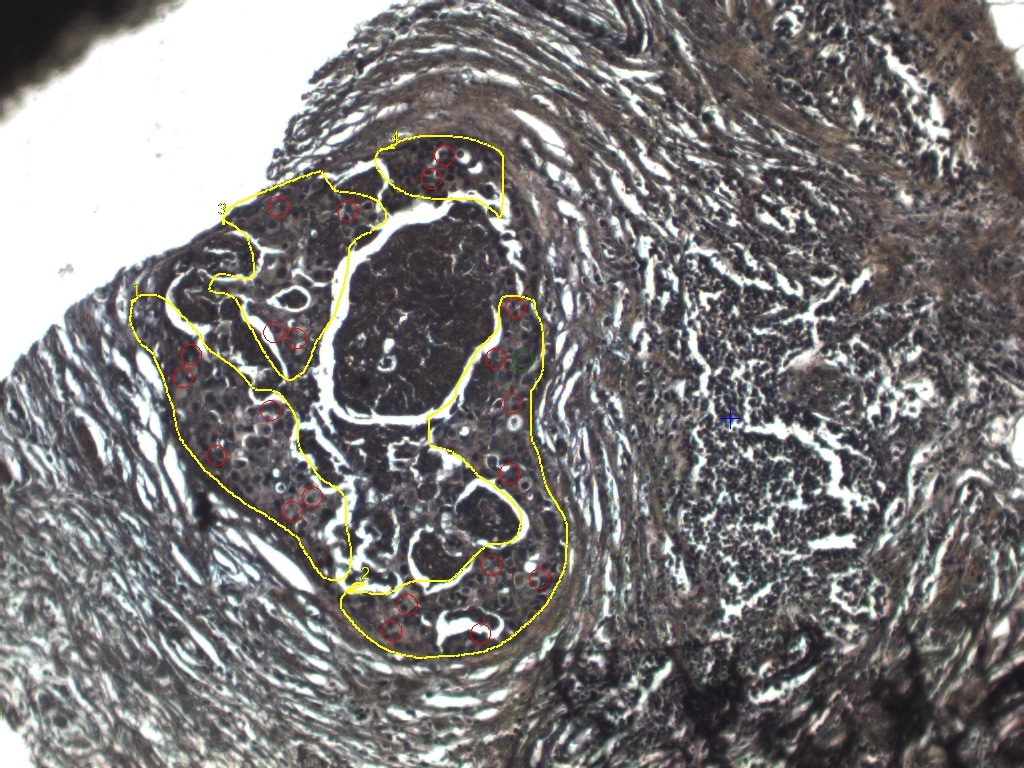

### IDC1-2 BEFORE.jpeg

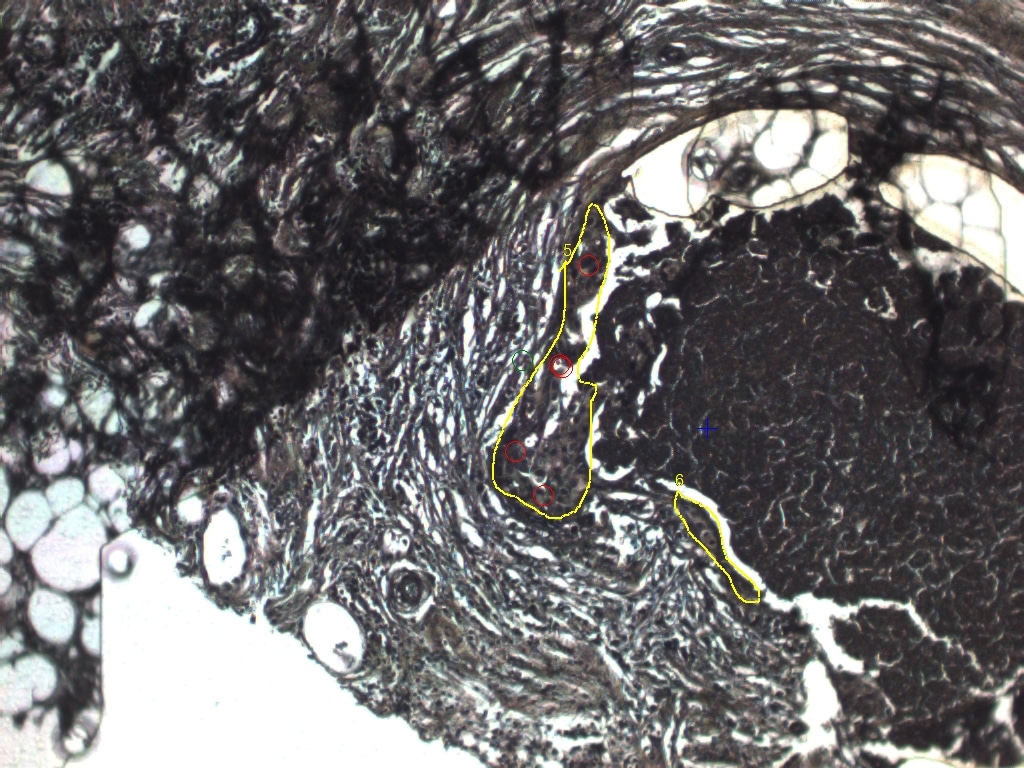

### IDC1AFTER.jpeg

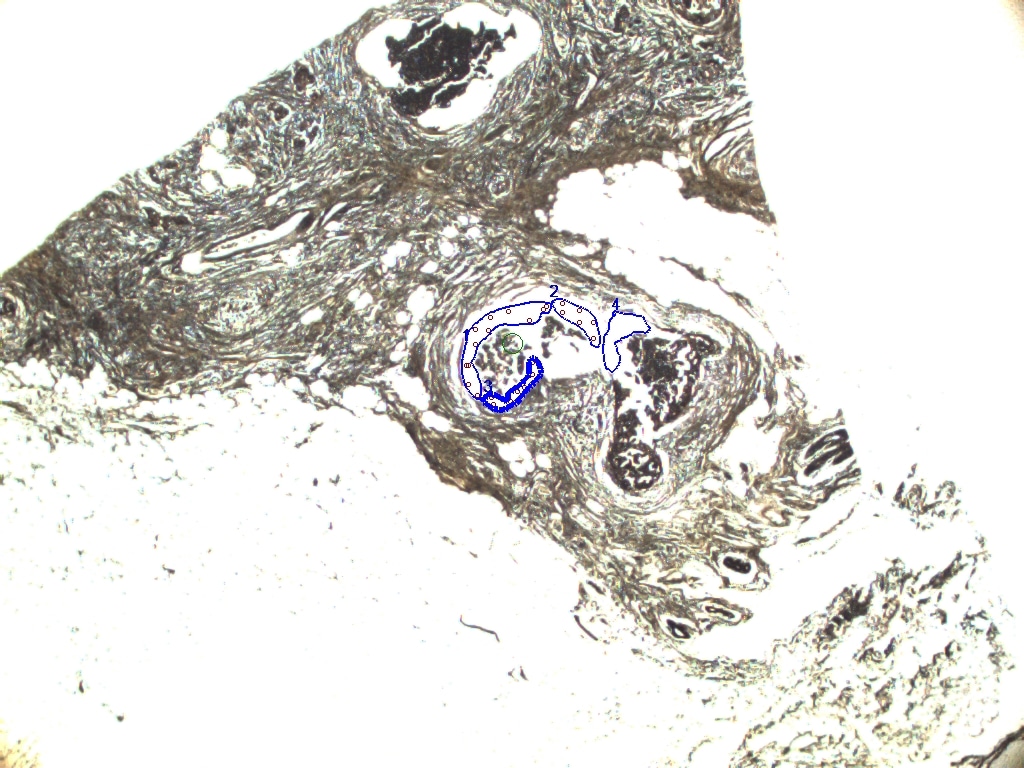

### IDC2 BEFORE.jpeg

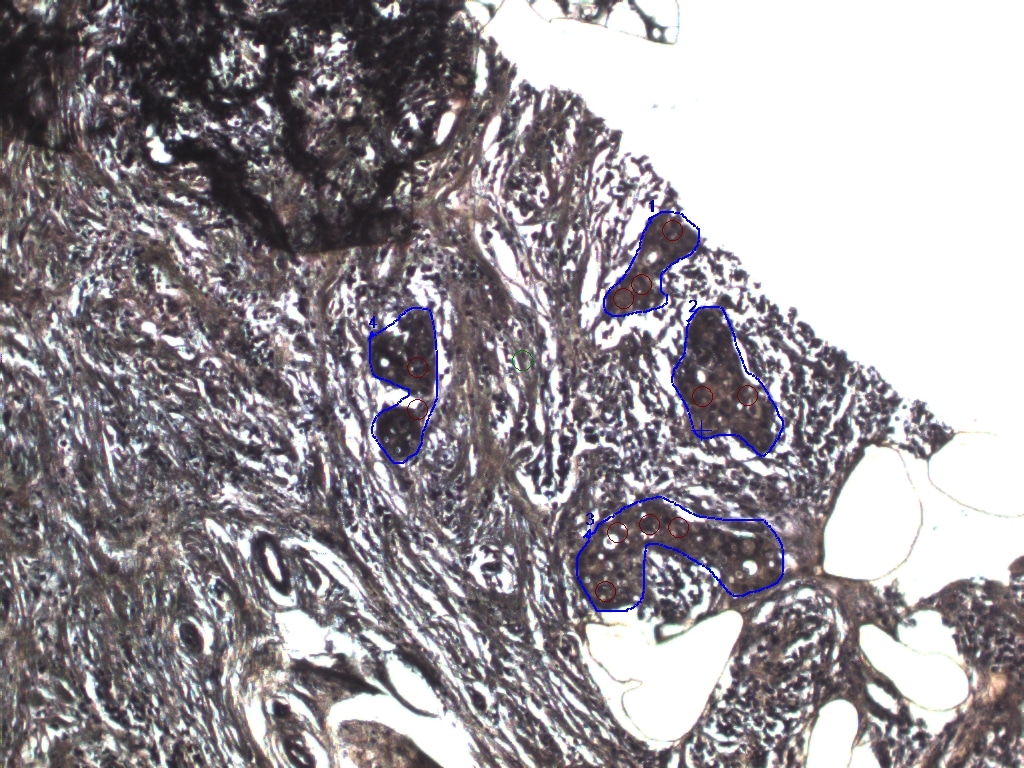

### IDC2-2 BEFORE.jpeg

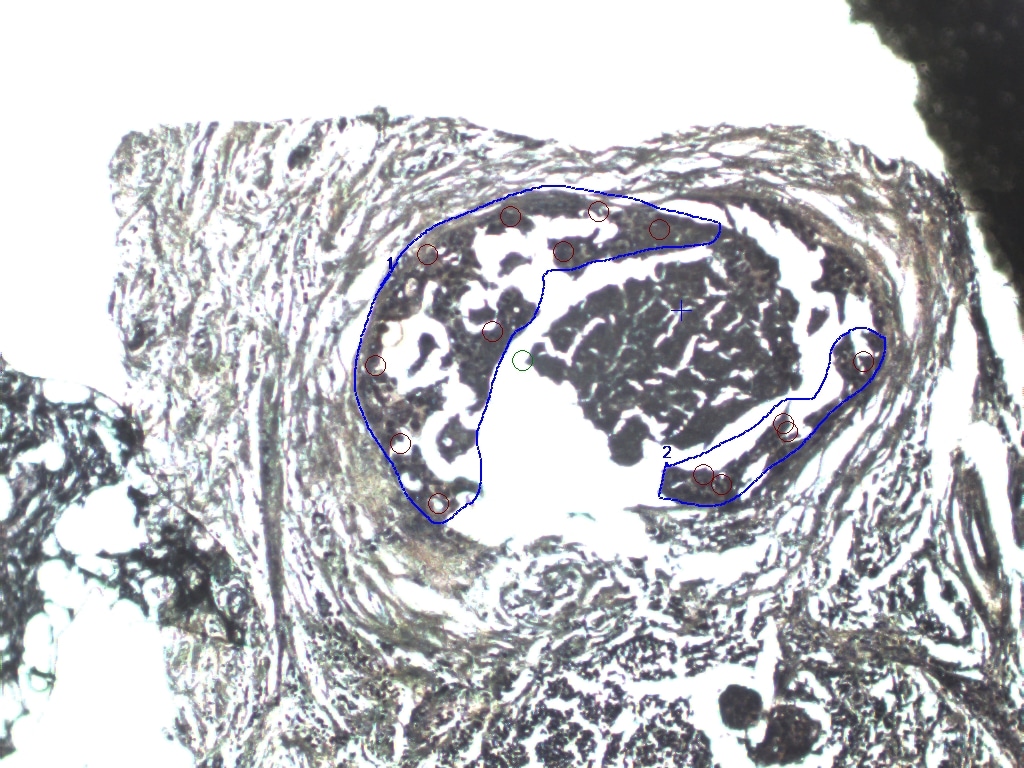

### MAC1 .jpeg

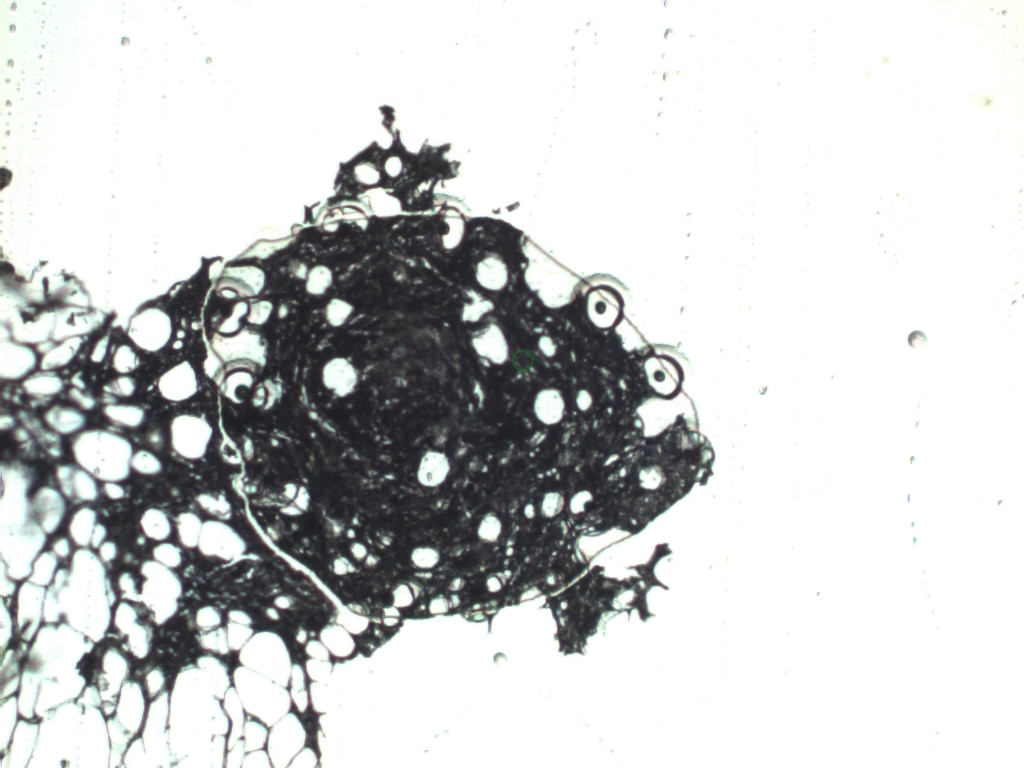

### MAC1 AFTER.jpeg

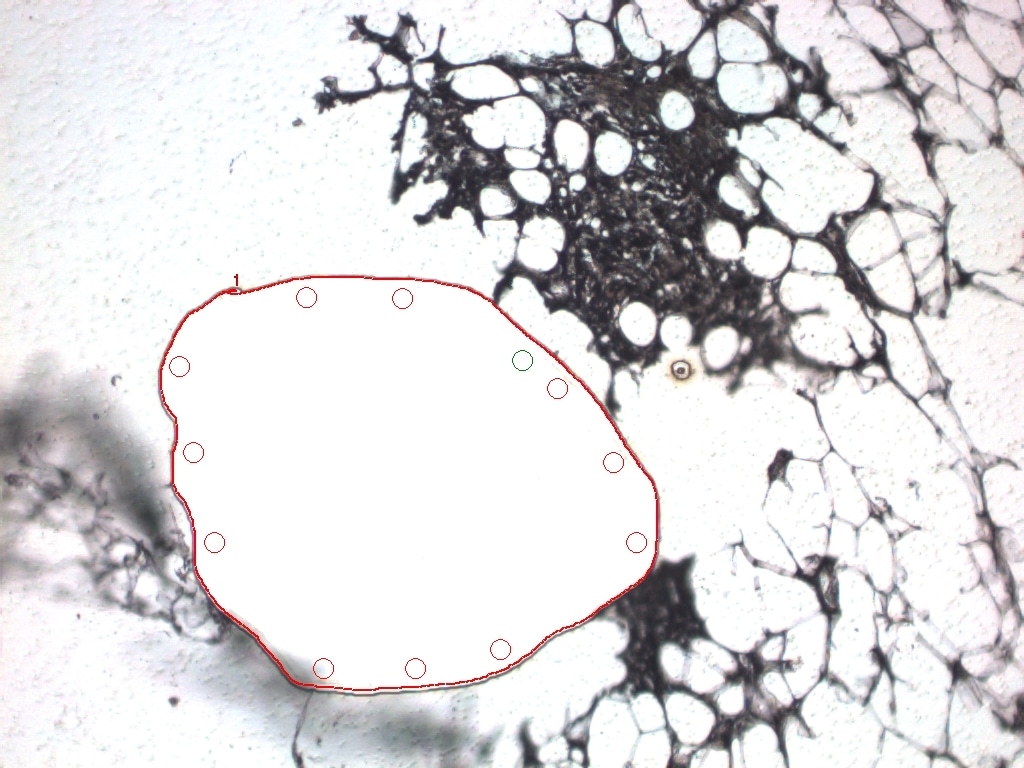

### MAC1 BEFORE.jpeg

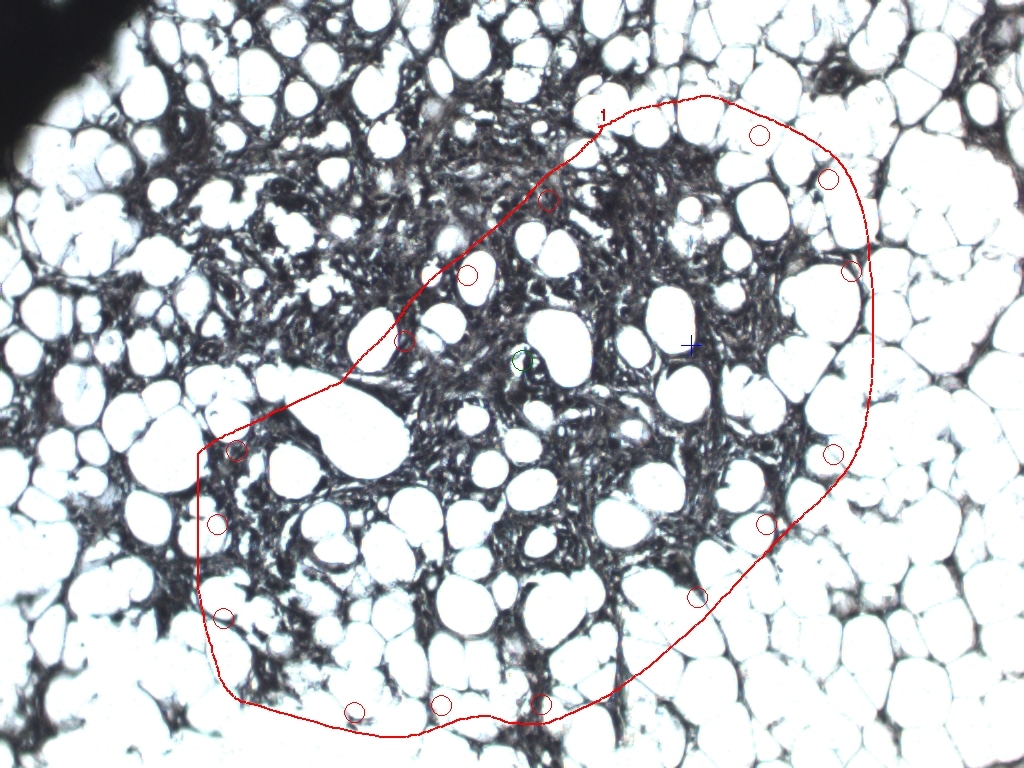

### MAC2 .jpeg

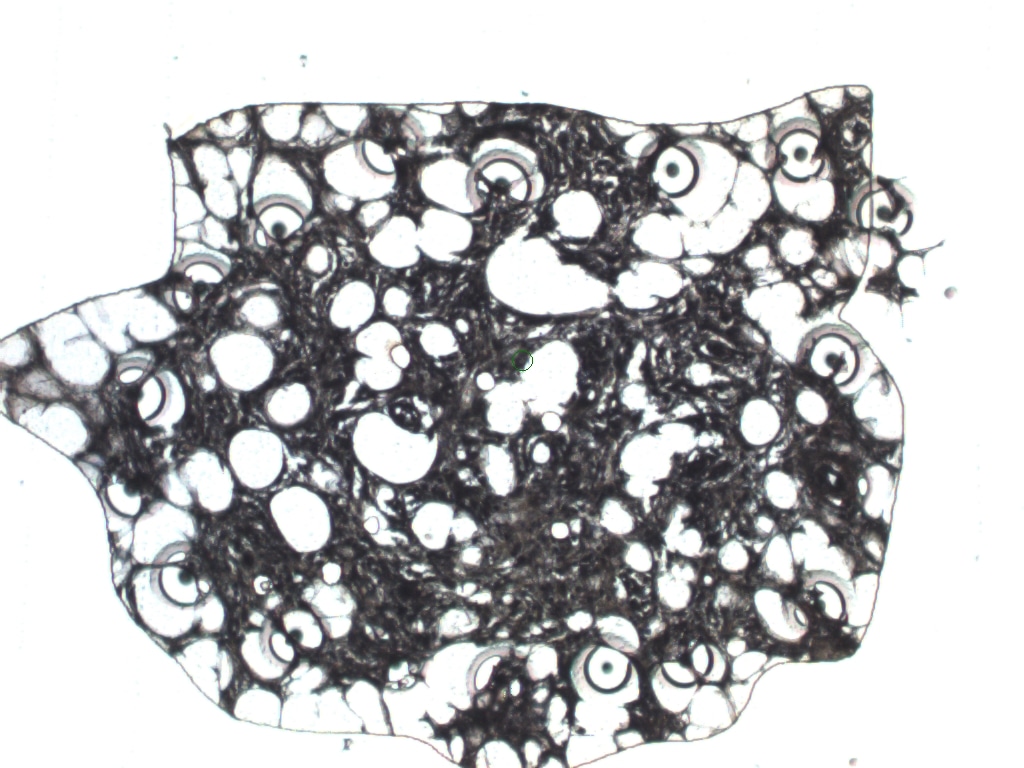

### MAC2 AFTER ABLATION10x.jpeg

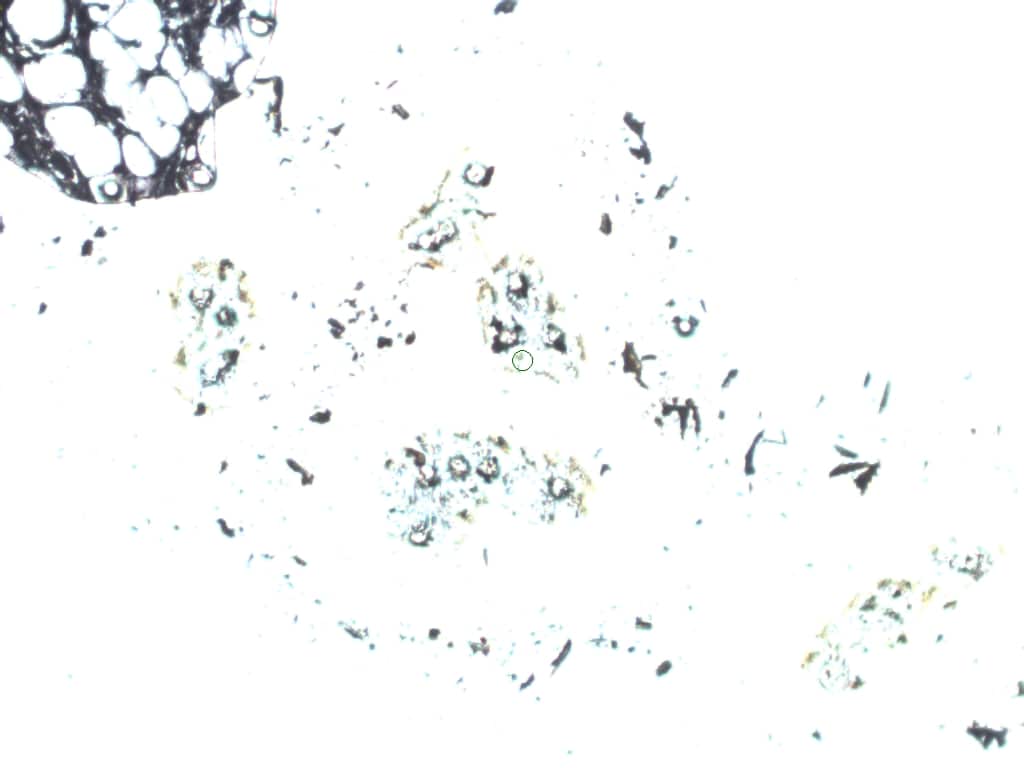

### MAC2 AFTER ABLATION .jpeg

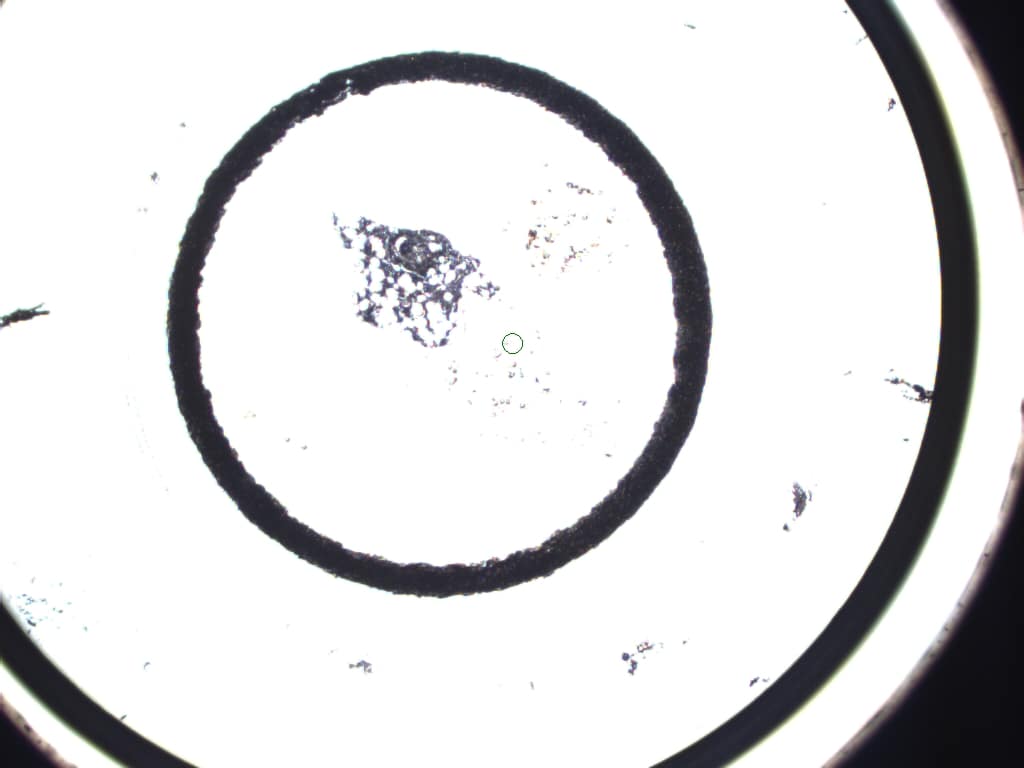

### MAC2 BEFORE ABLATION.jpeg

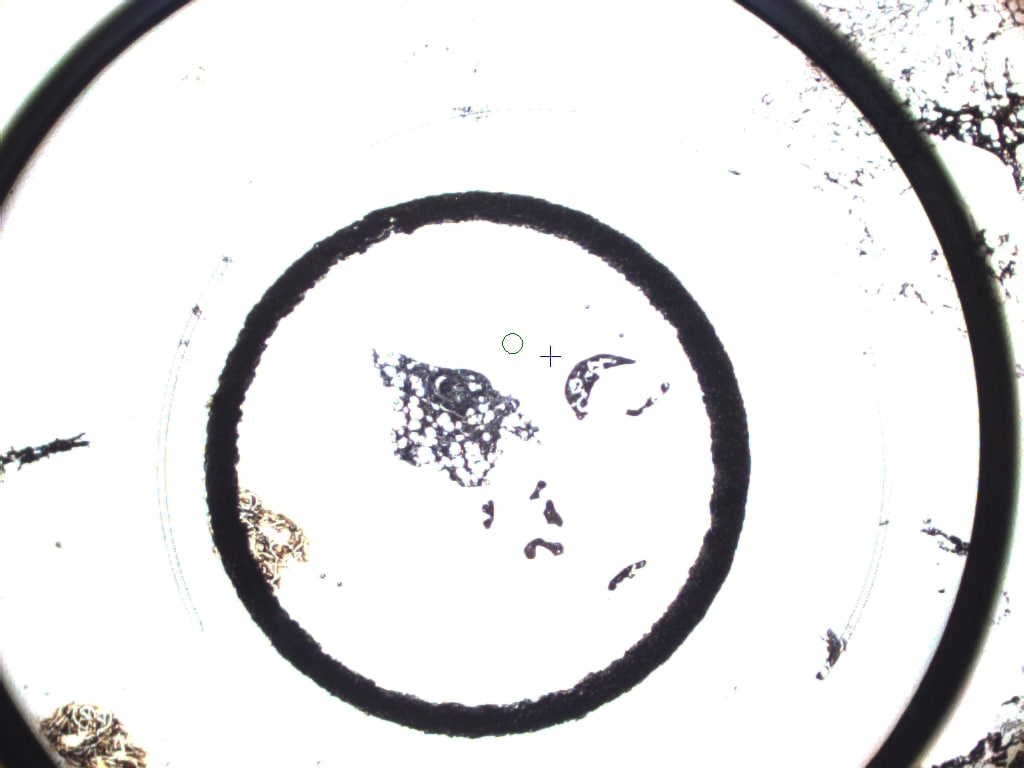

### MAC2 BEFORE.jpeg

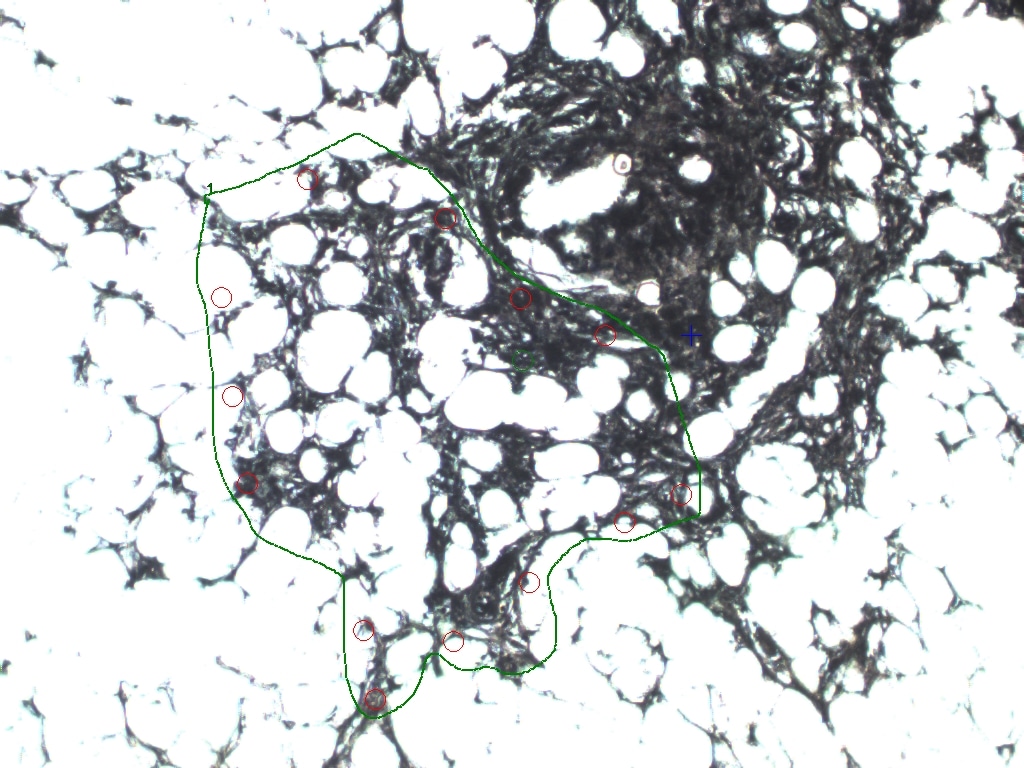

### MAC2AFTER.jpeg

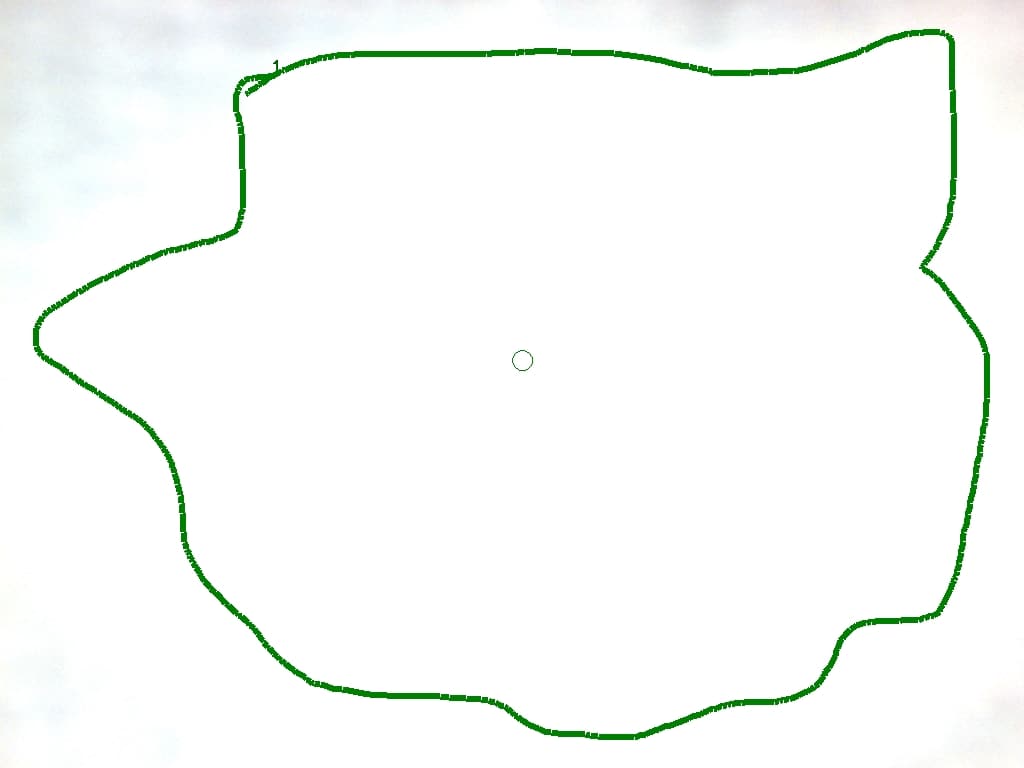

### MAC3 AFTER ABLATION10x.jpeg

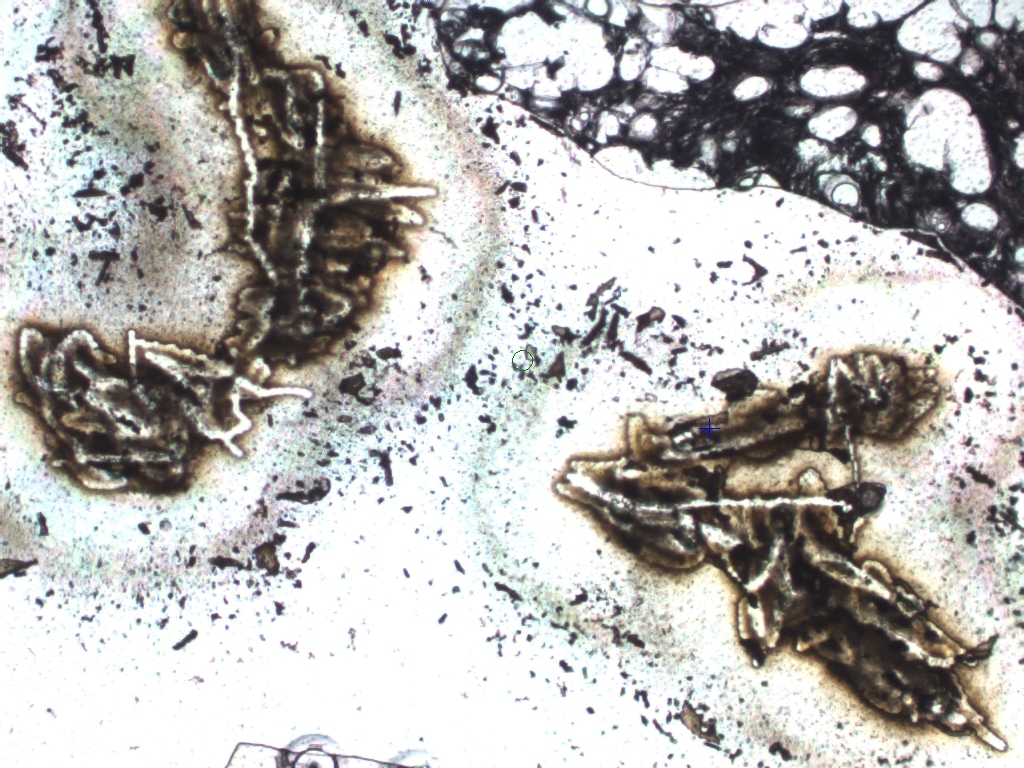

### MAC3 AFTER ABLATION .jpeg

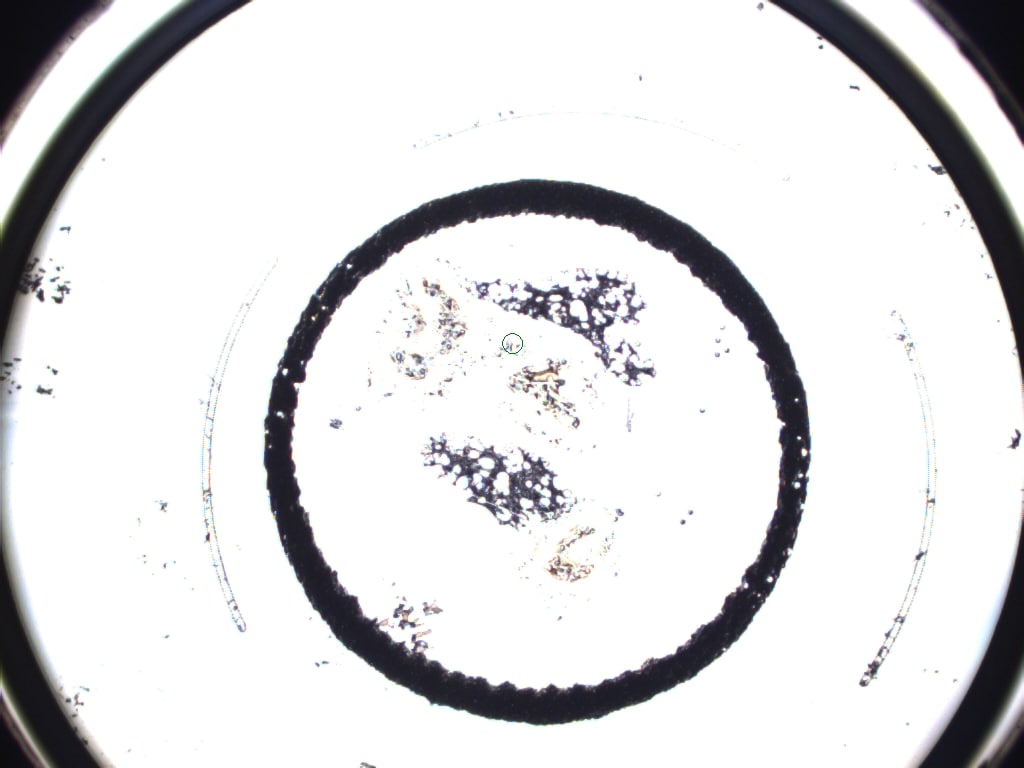

### MAC3 AFTER.jpeg

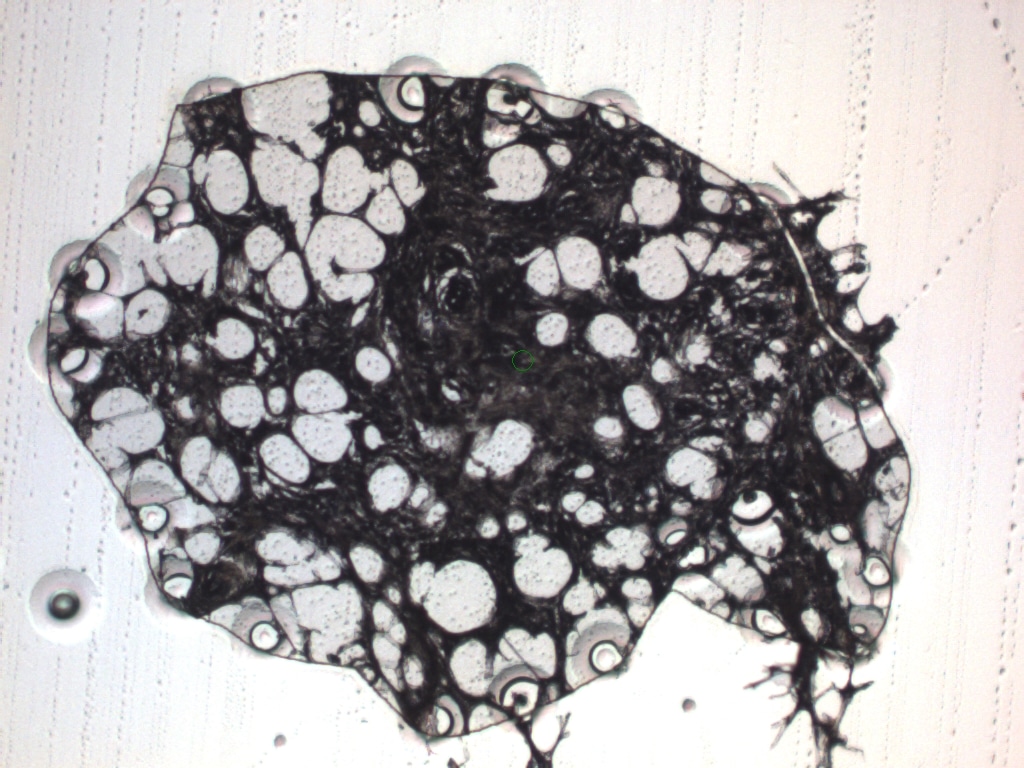

### MAC3 BEFORE ABLATION.jpeg

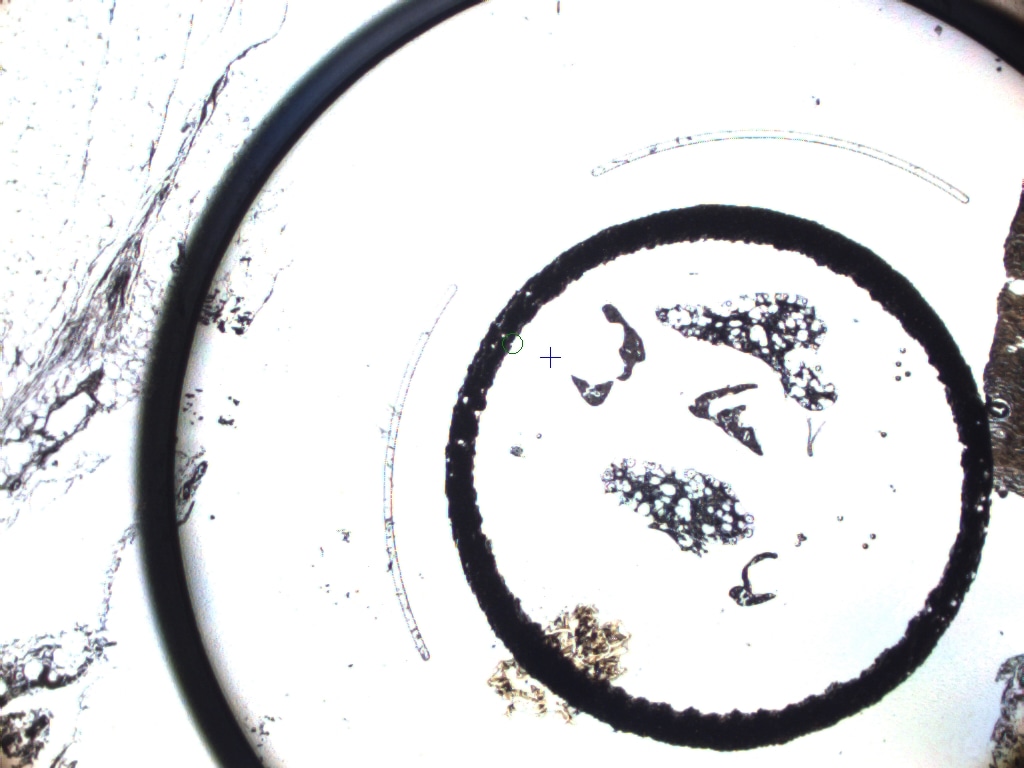

### MAC3 BEFORE.jpeg

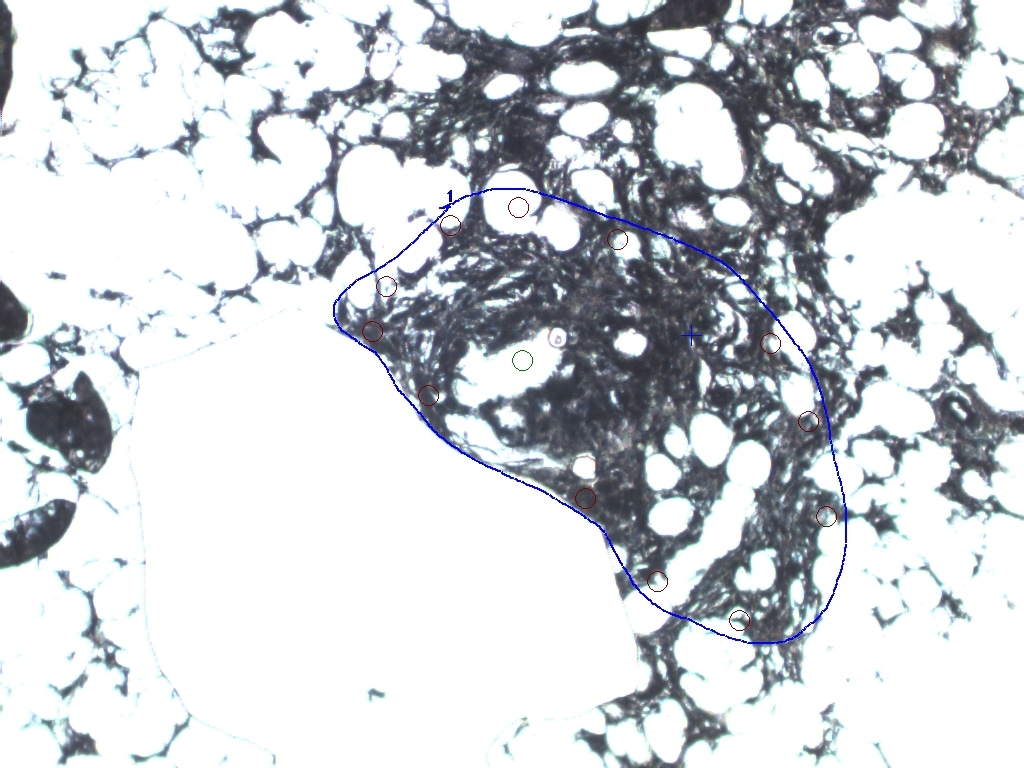

### MAC3AFTER RIGHT.jpeg

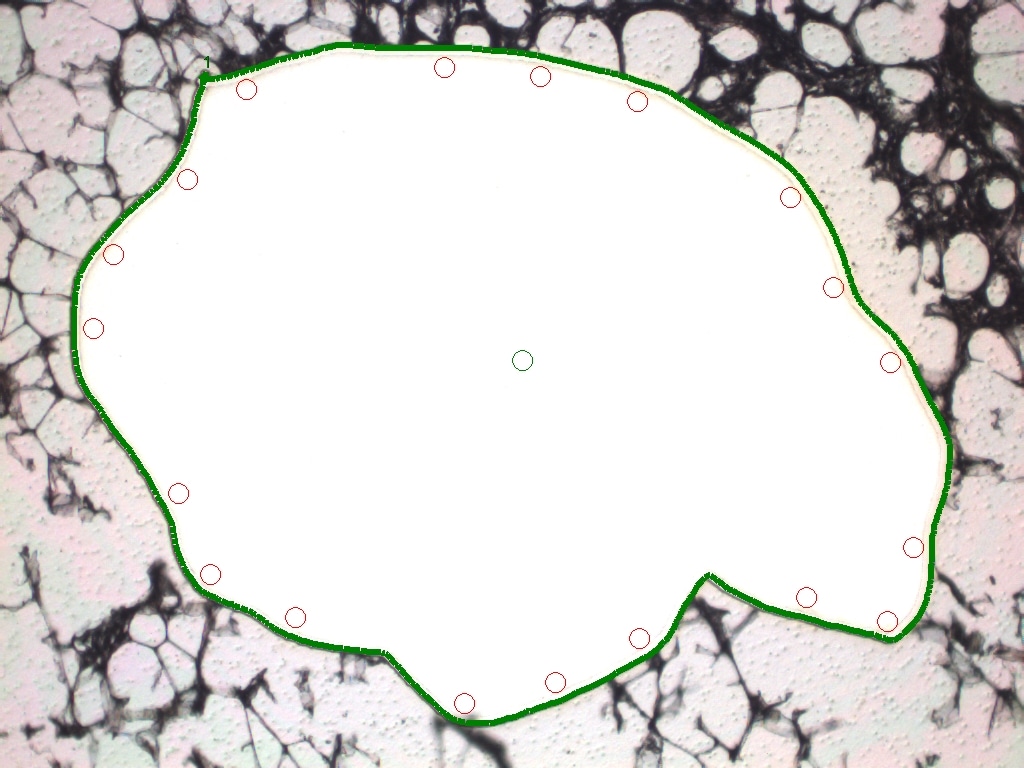

### MAC4 .jpeg

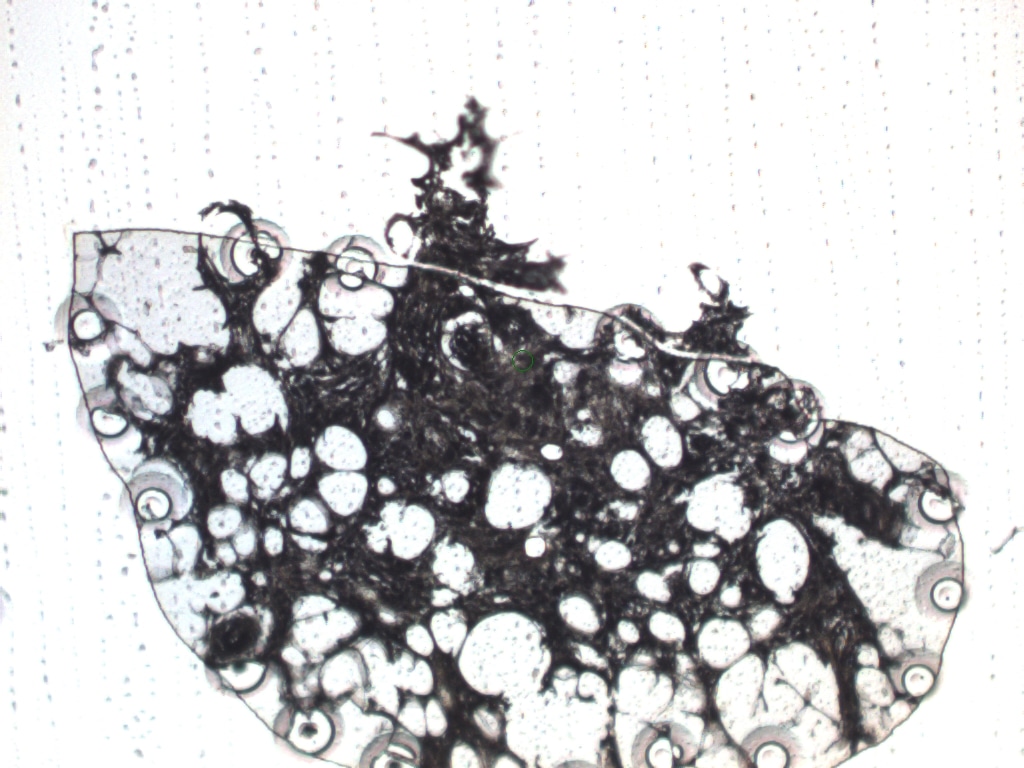

### MAC4 AFTER.jpeg

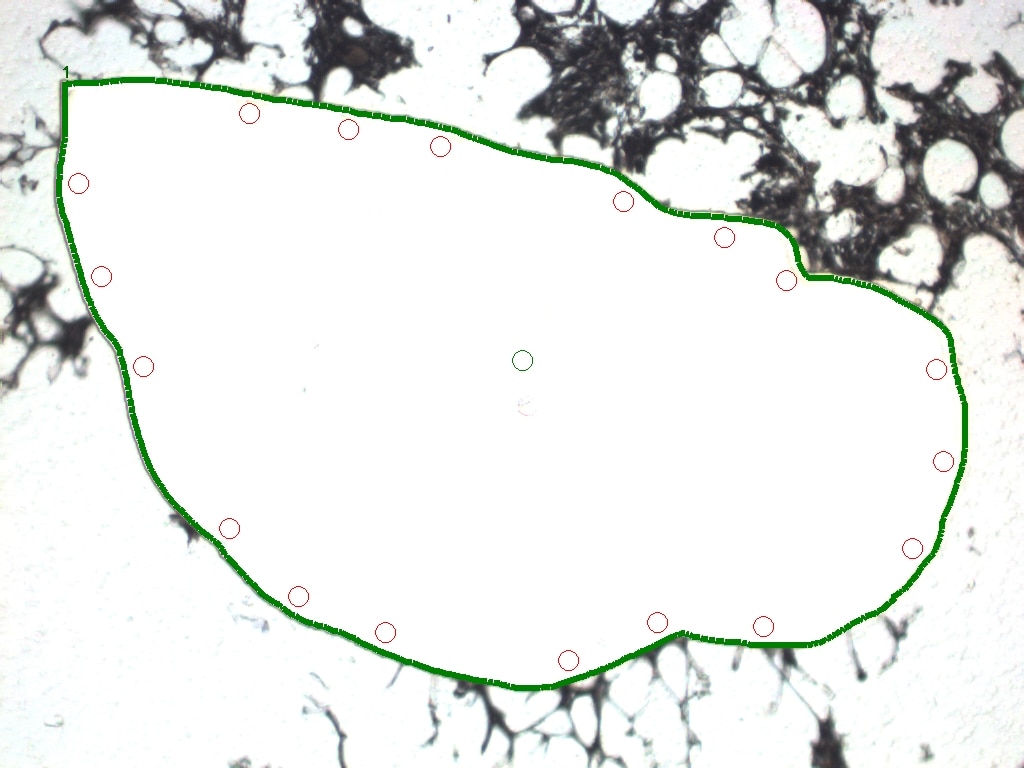

### MAC5 .jpeg

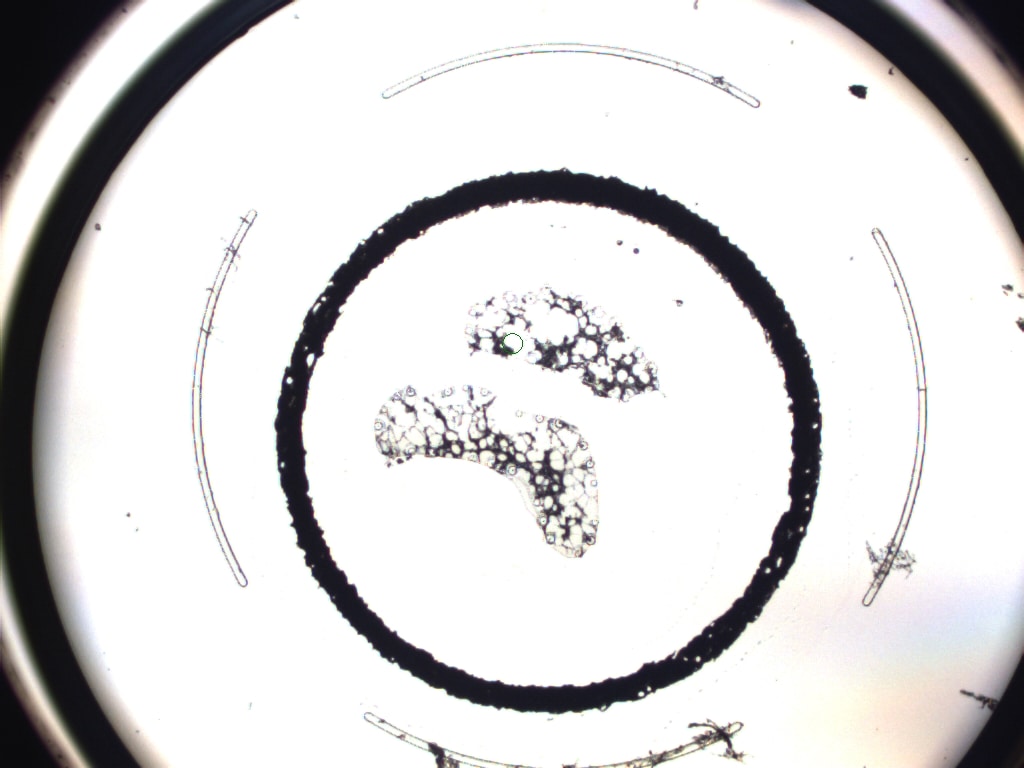

### MAC6 .jpeg

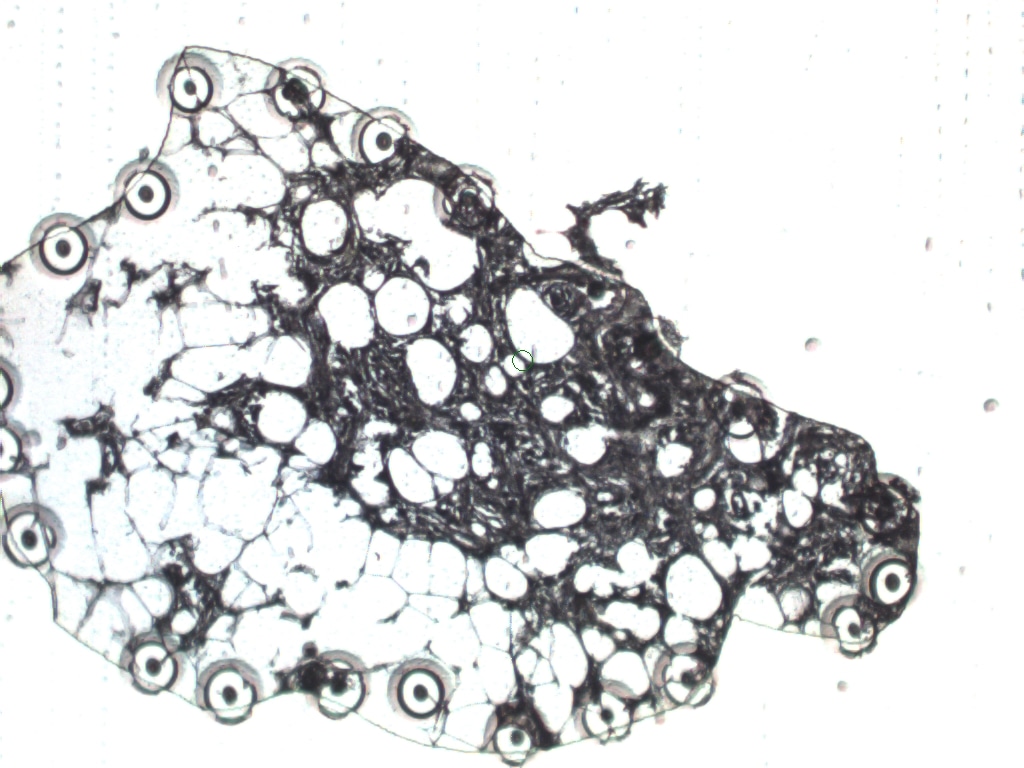

### MACROPHAGE1 BEFORE 10x.jpeg

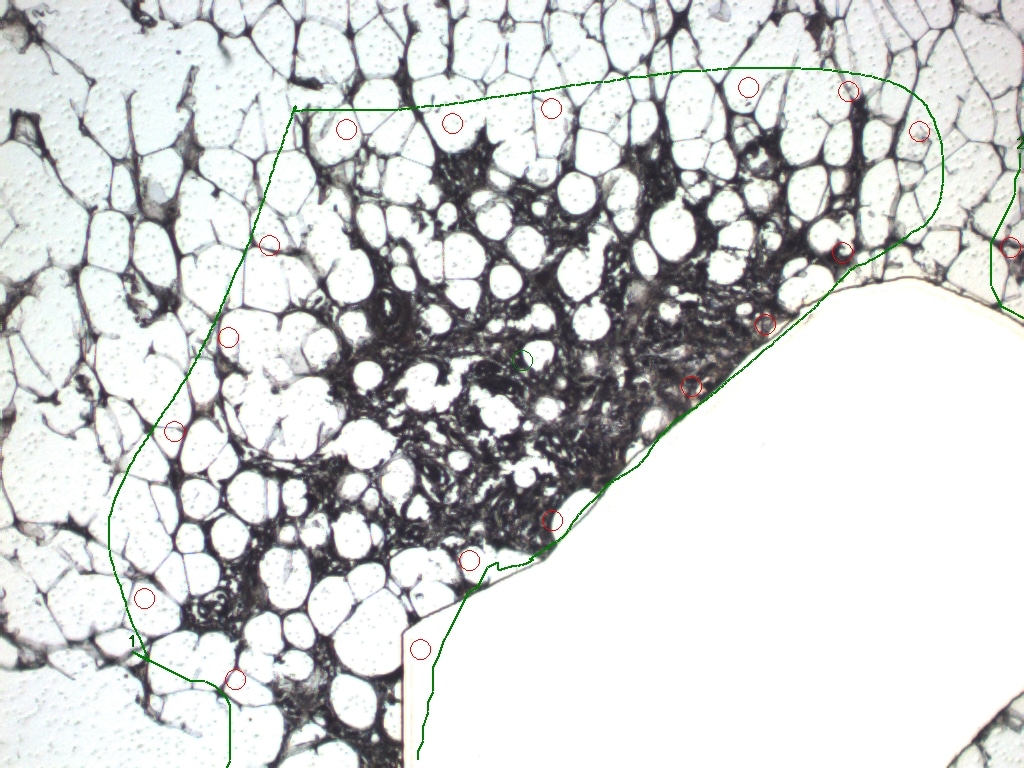

### MACROPHAGE2 BEFORE 10x.jpeg

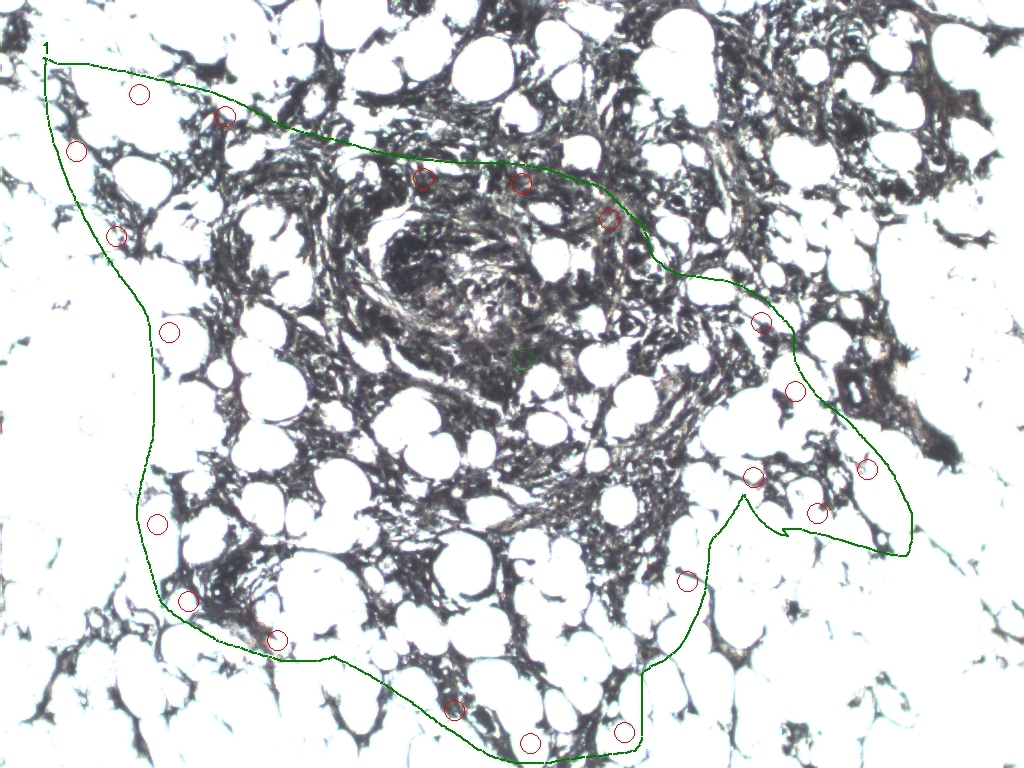

### MACROPHAGE2 BEFORE 10x.jpeg

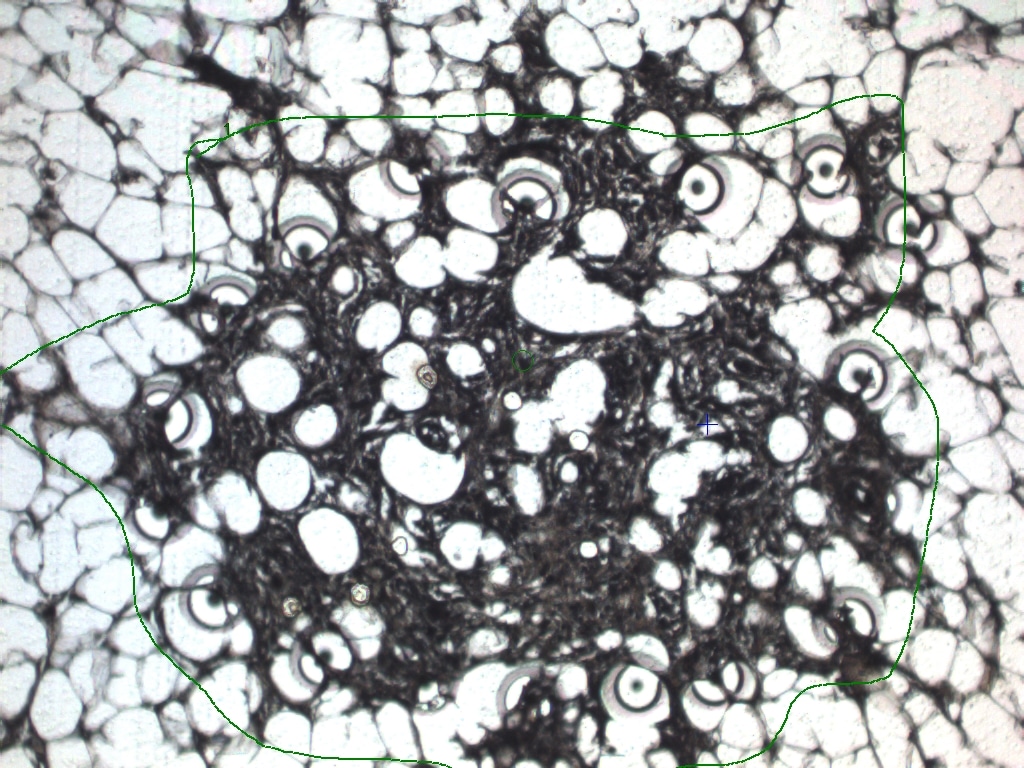

### MACROPHAGE2BEFORE 10x.jpeg

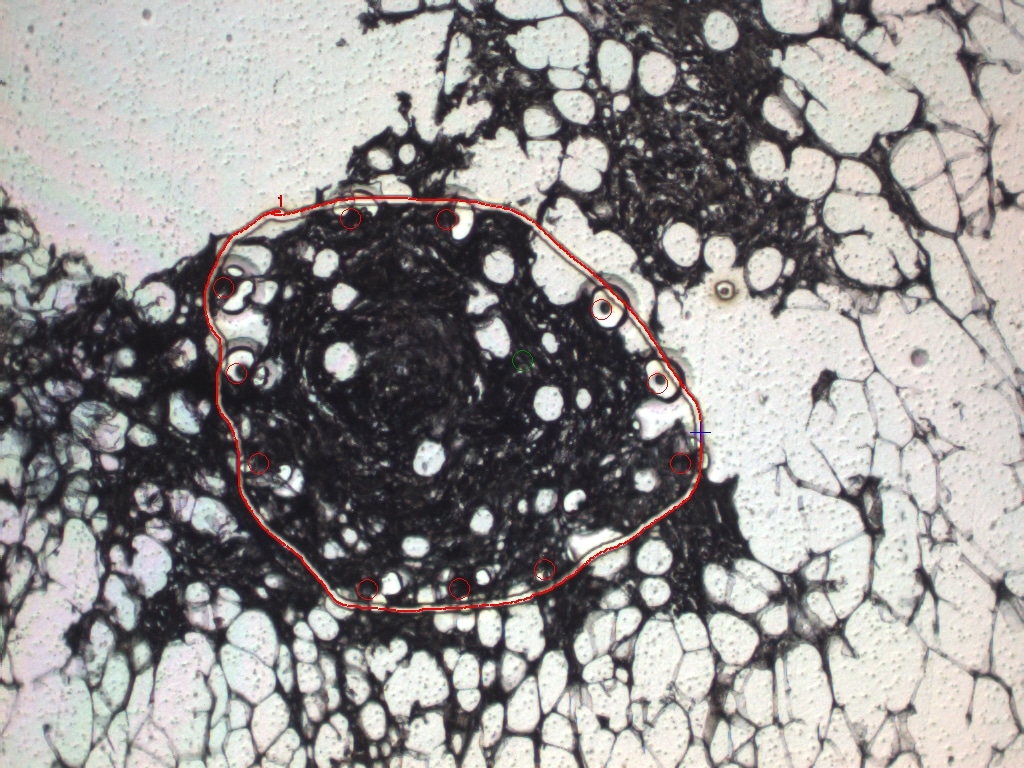

### MACROPHAGE3 BEFORE 10x.jpeg

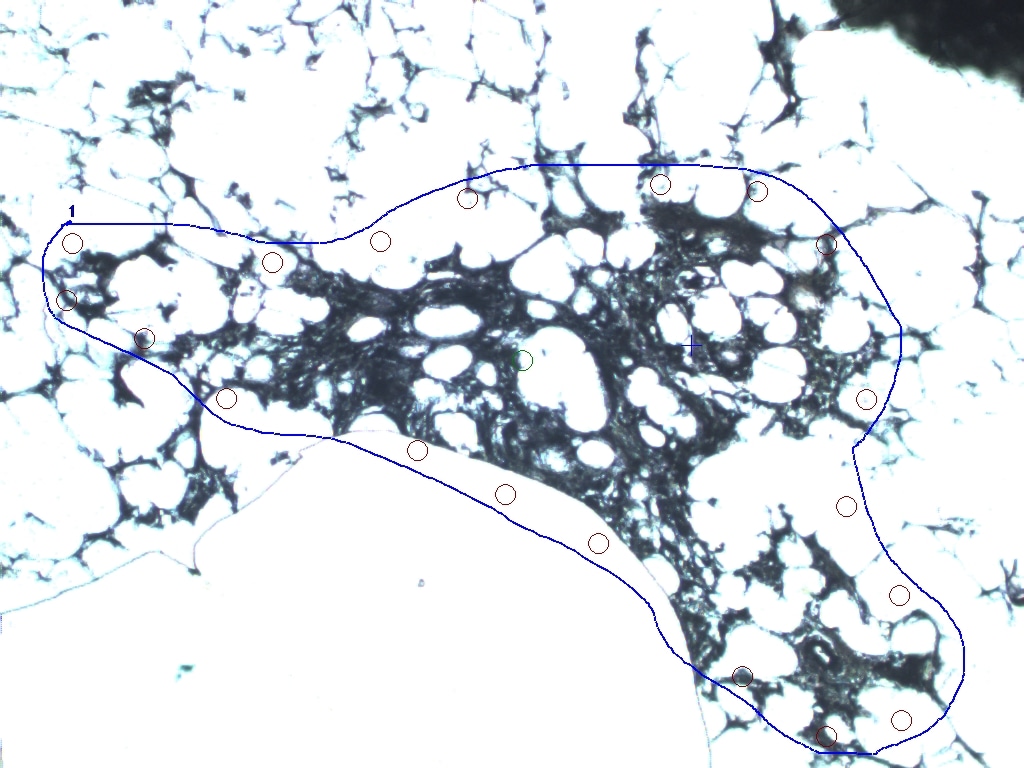

### MACROPHAGE3 BEFORE 10x.jpeg

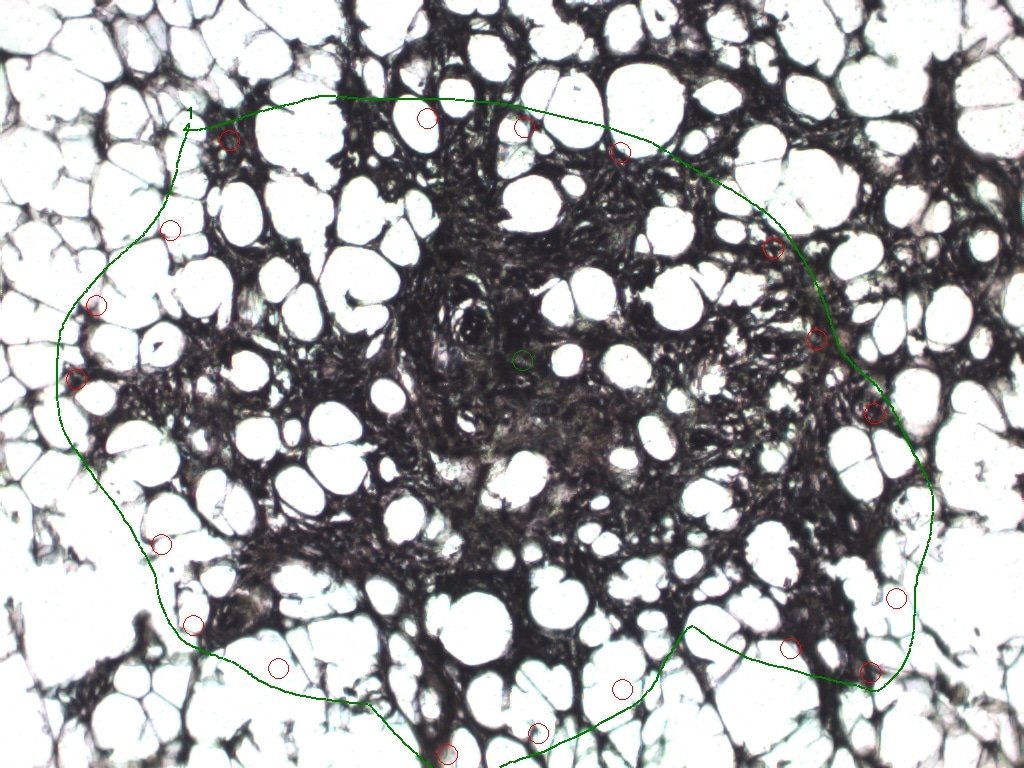

### MACROPHAGE3-2 BEFORE 10x.jpeg

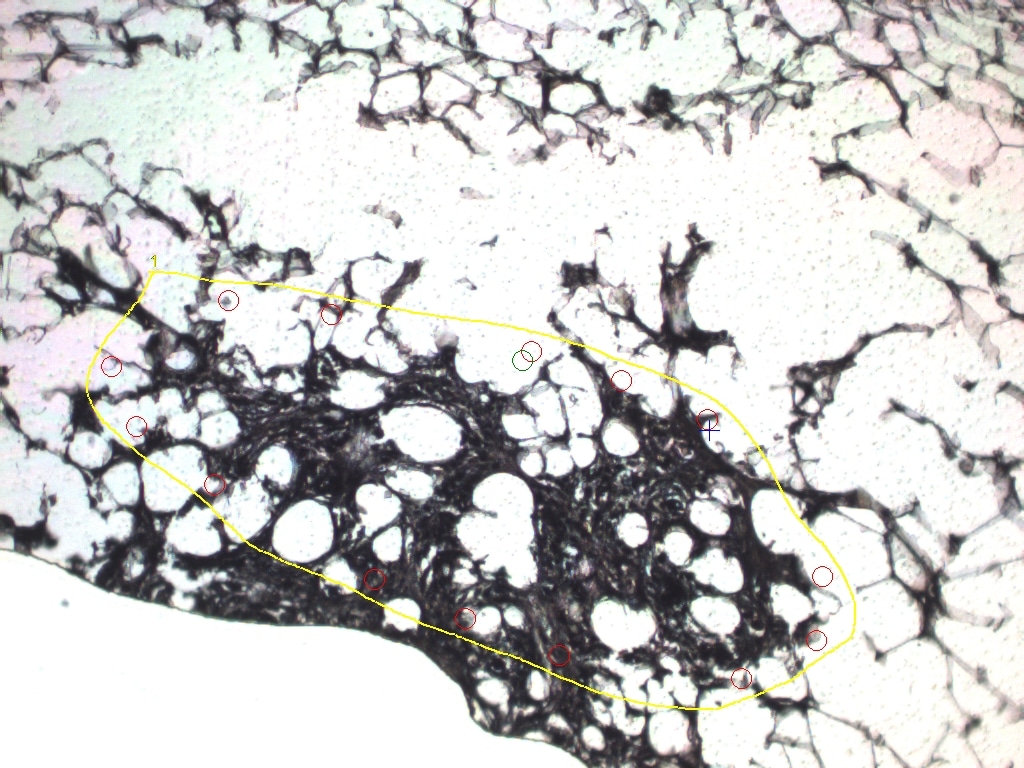

### MACROPHAGE4 BEFORE 10x.jpeg

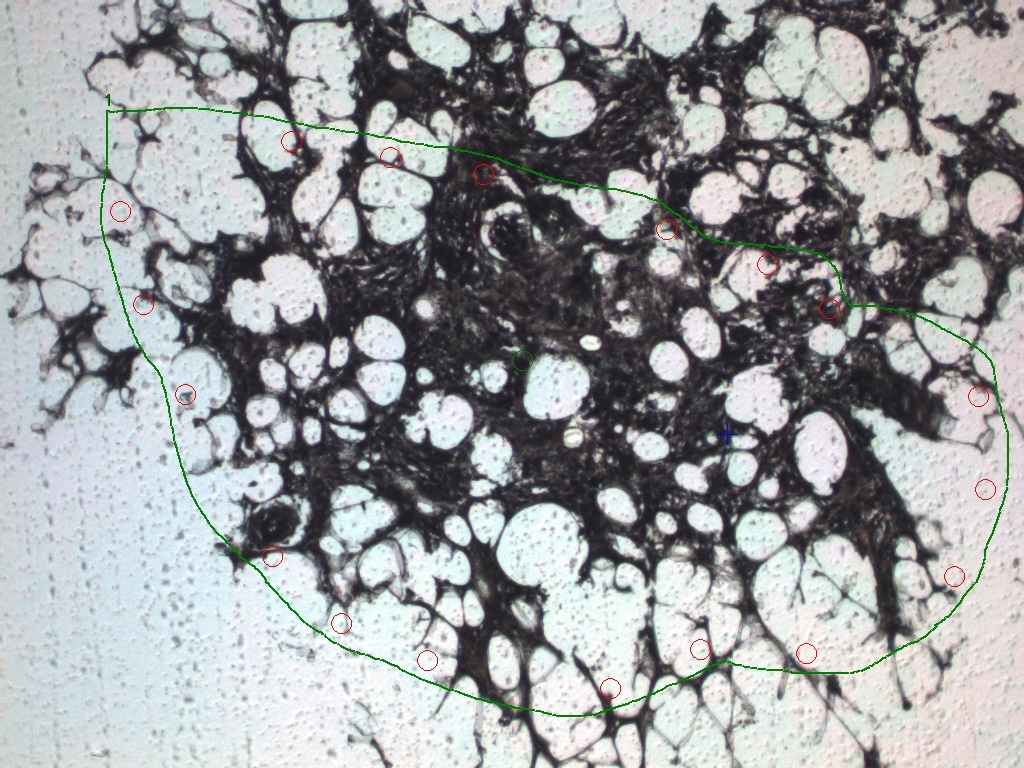

### MACROPHAGE6 AFTER.jpeg

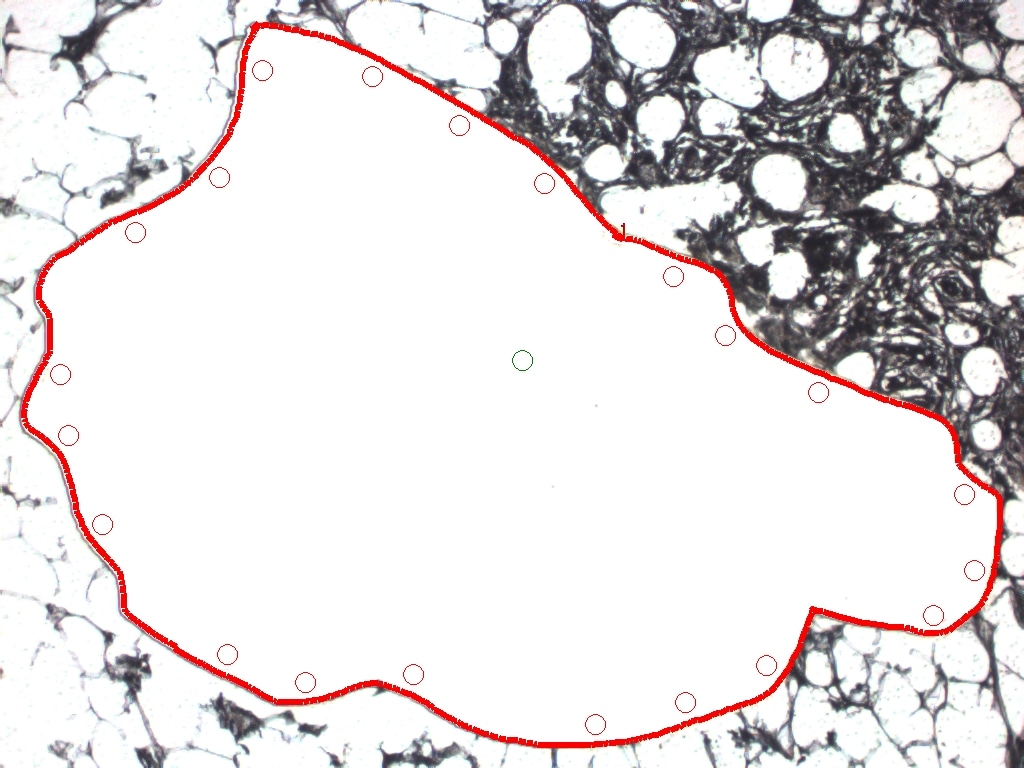

### MACROPHAGE6 BEFORE 2.jpeg

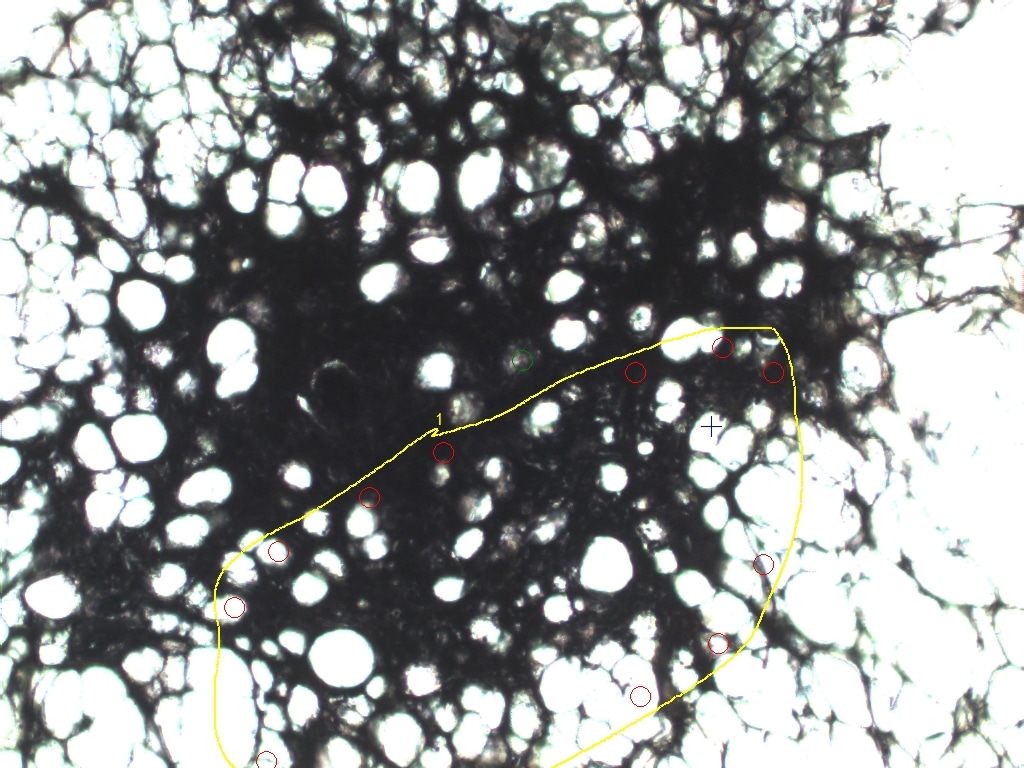

### MACROPHAGE6 BEFORE 10x.jpeg

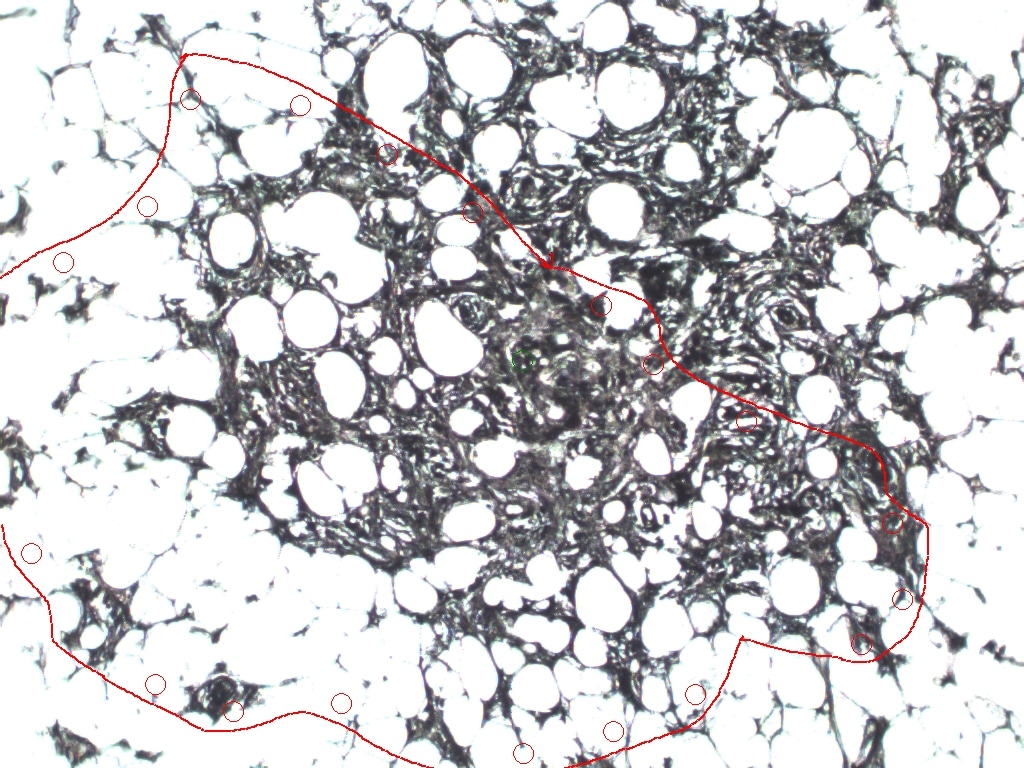

### MACROPHAGE 1 AFTER.jpeg

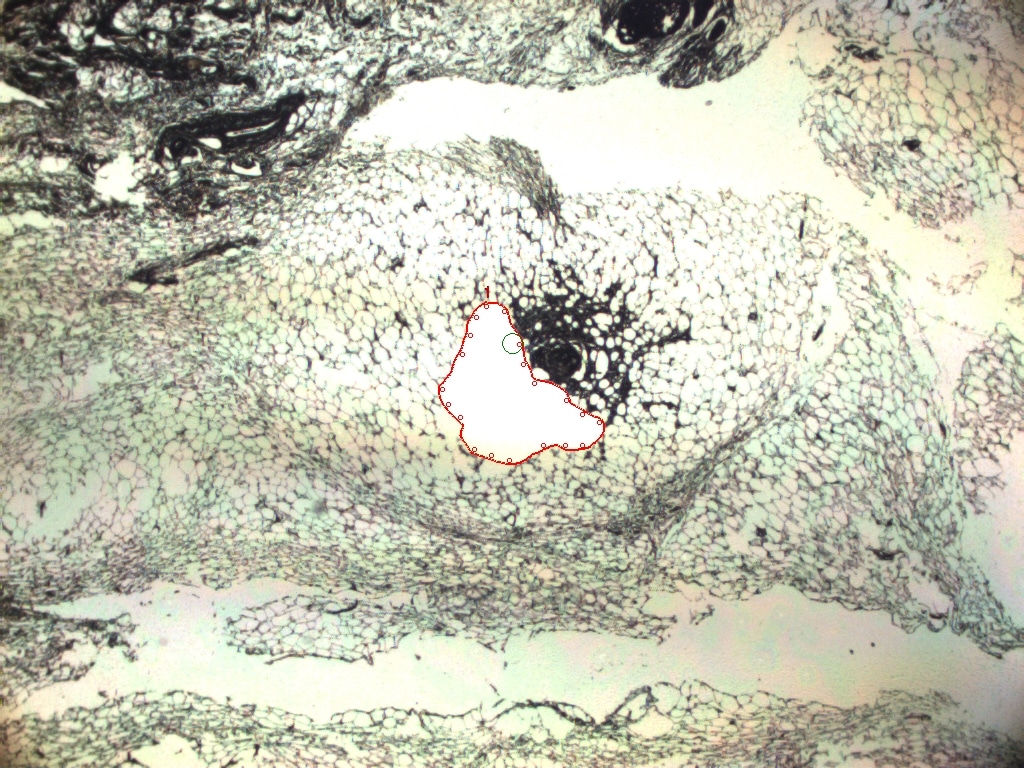

### MACROPHAGE 1 AFTER.jpeg

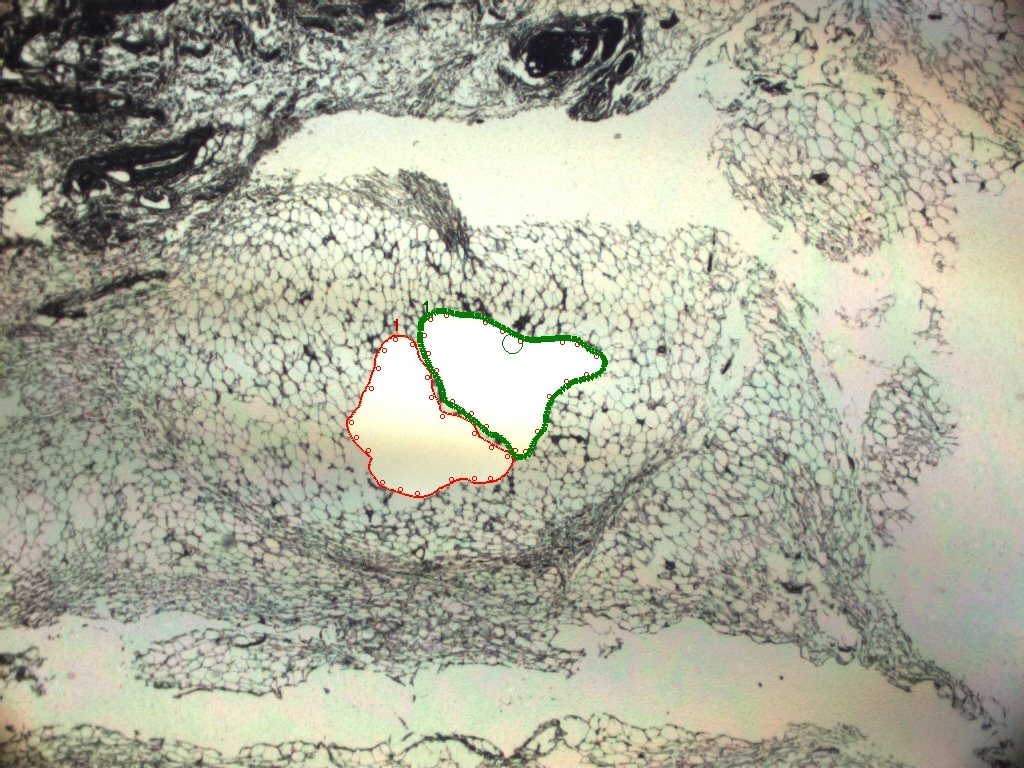

### MACROPHAGE 1 BEFORE 10x.jpeg

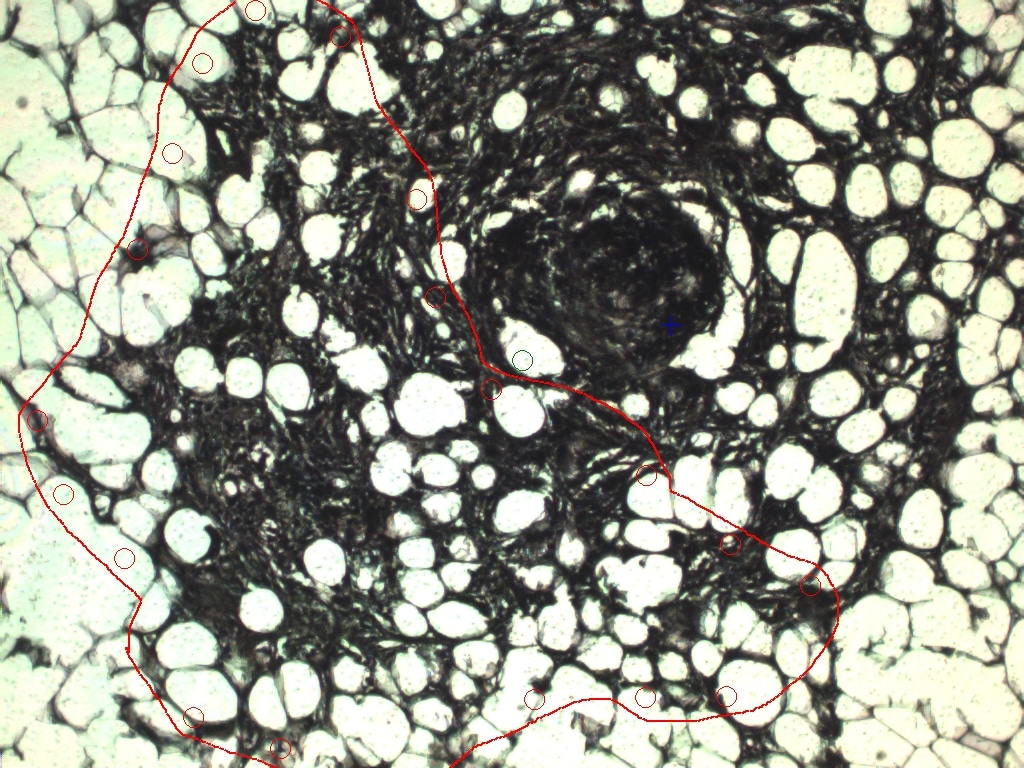

### MACROPHAGE 1 BEFORE.jpeg

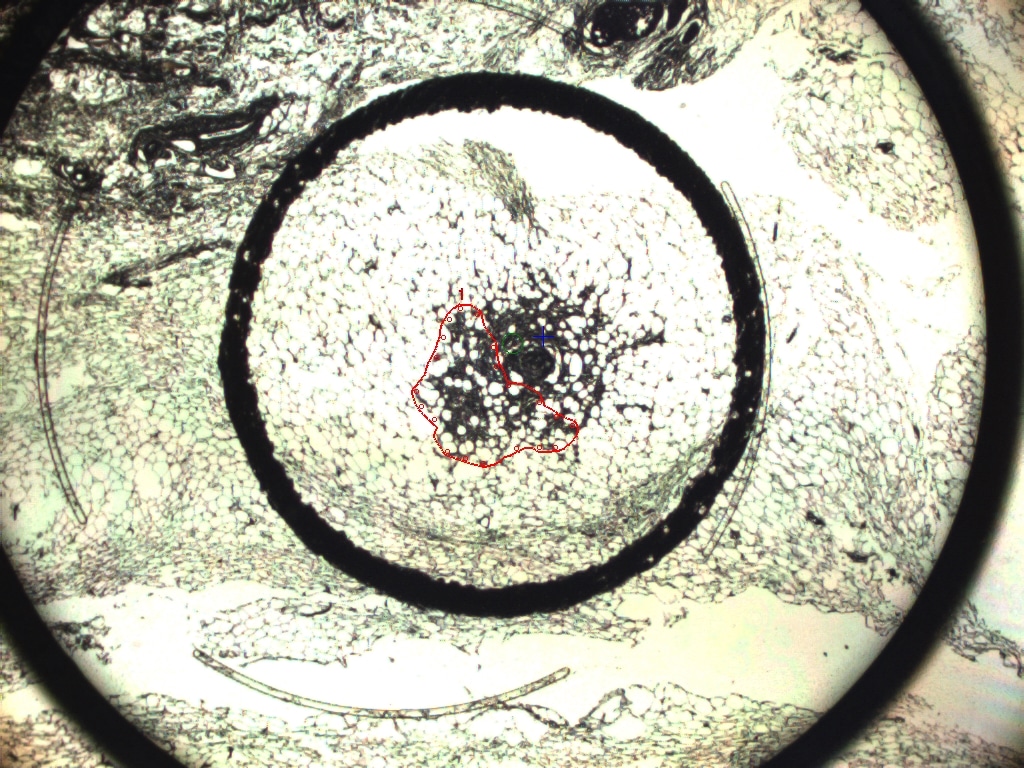

### MACROPHAGE 1 BEFORE.jpeg

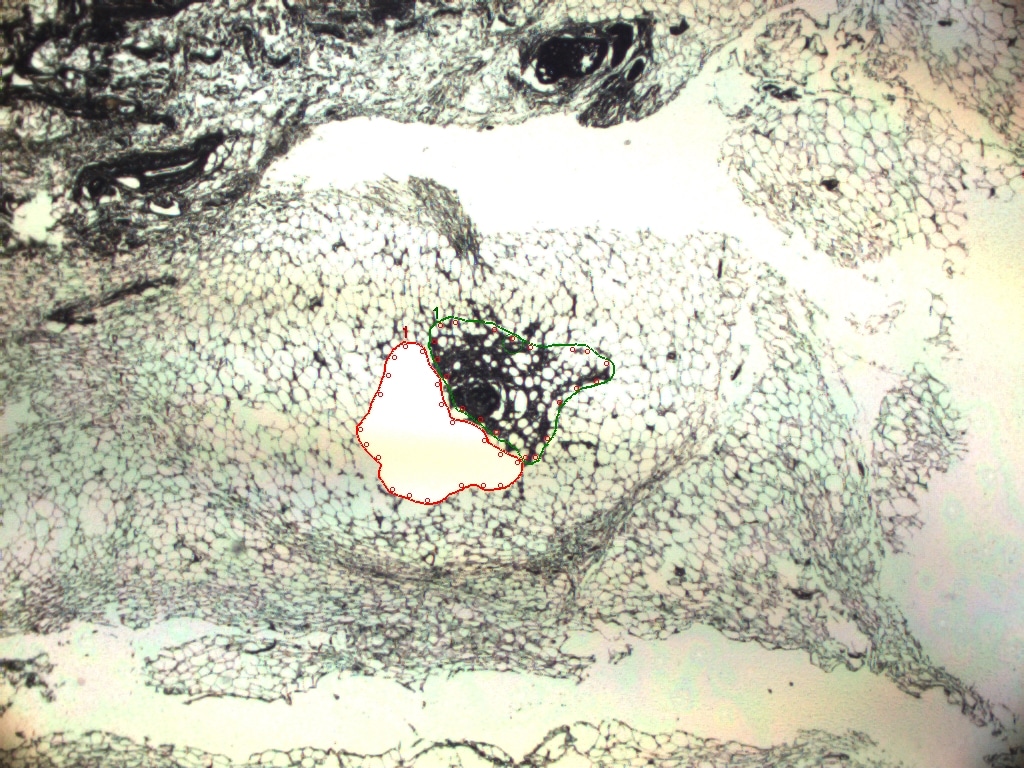

### MACROPHAGE 2 AFTER.jpeg

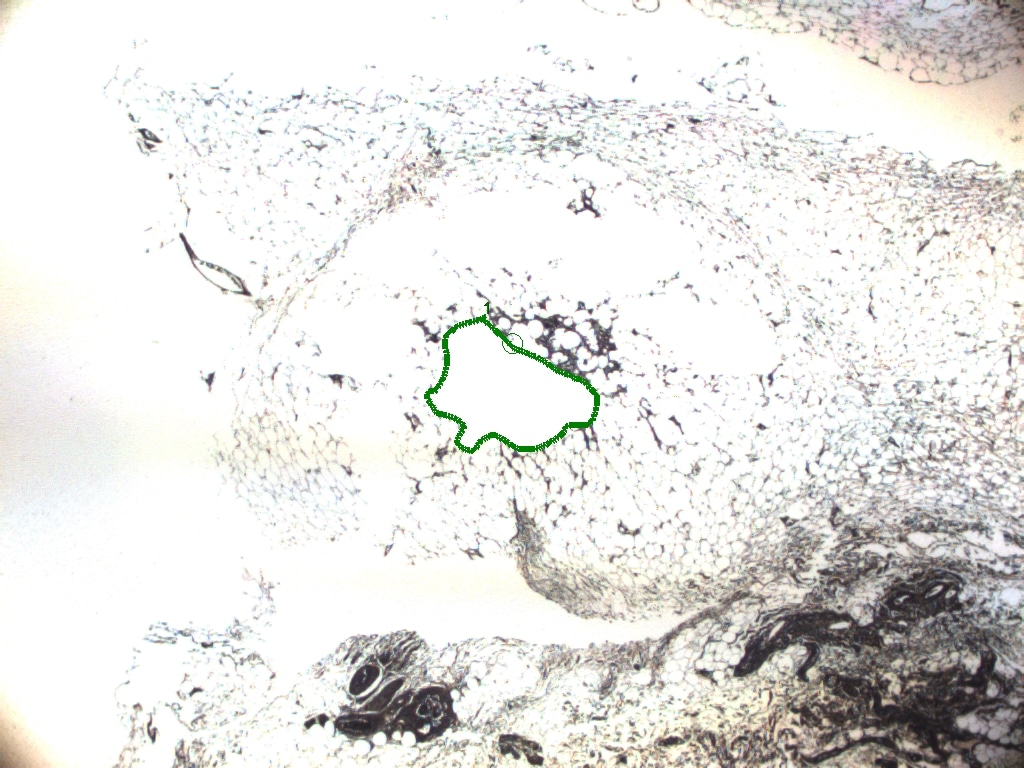

### NO BLATION CAP1.jpeg

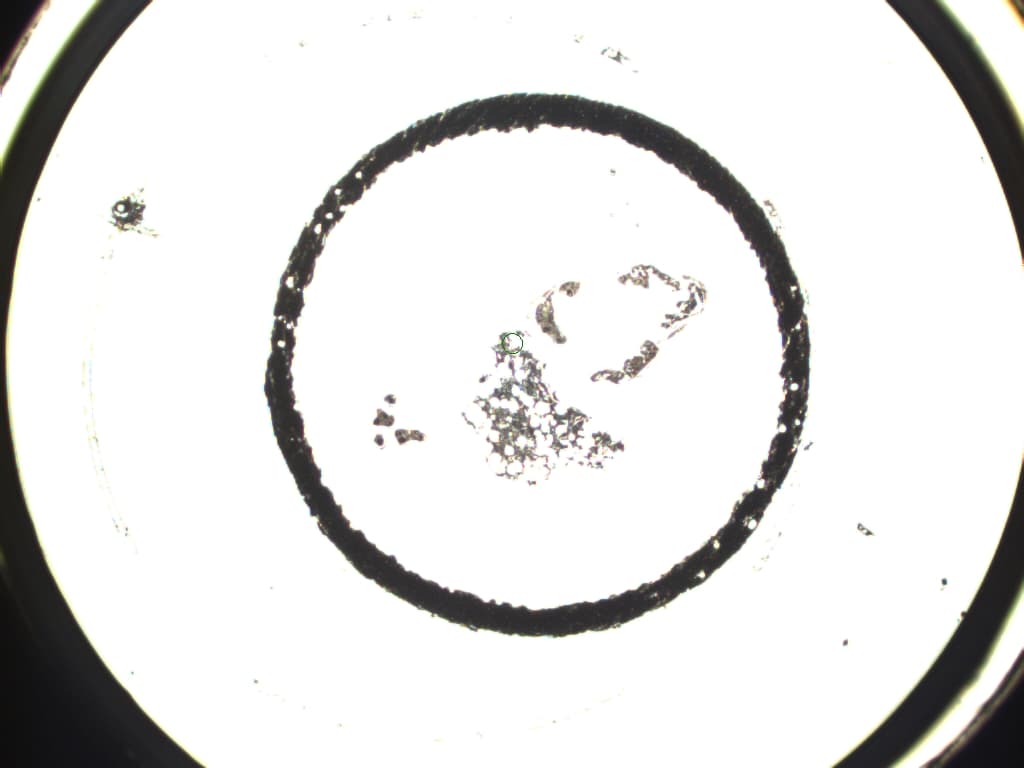

### NO BLATION CAP2 DCIS7+MAC.jpeg

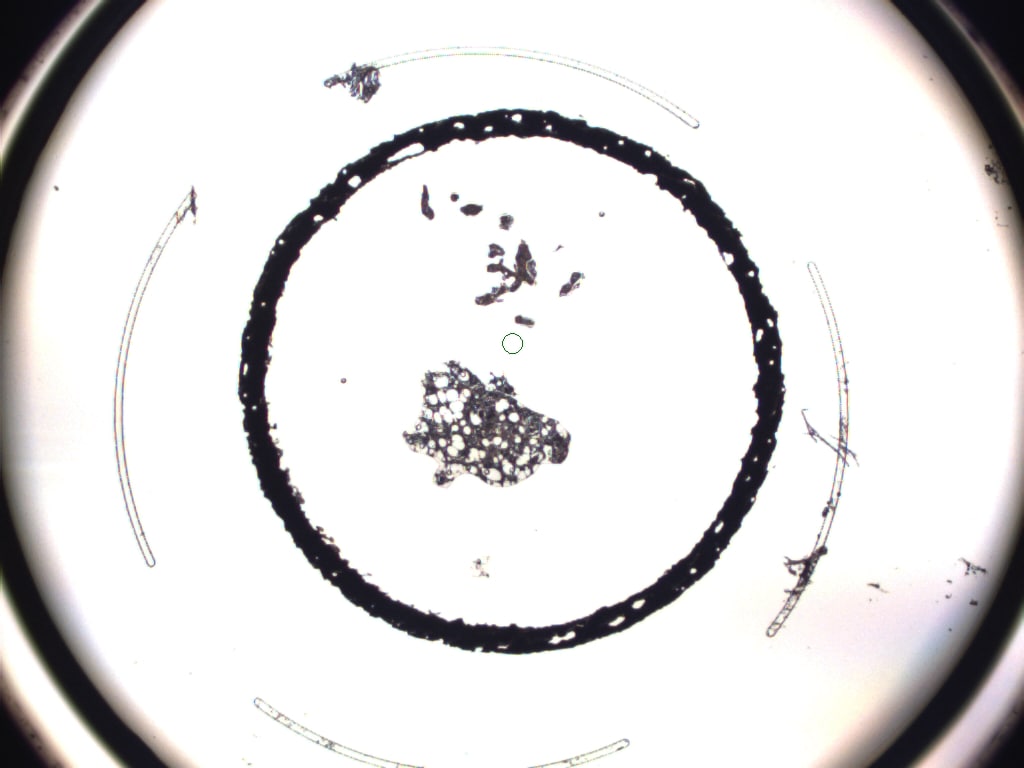
