## Supplemental Code for "Gene-expression profiling of single cells from archival tissue with laser-capture microdissection and Smart-3SEQ": heatmap.R

```
#! /usr/bin/Rscript

library(DESeq2)
library(reshape2)
library(ggplot2)

load("counts.RData")
load("conditions.RData")
load("rld.RData")

dds.bulk <- DESeq(DESeqDataSetFromMatrix(counts.filtered[,conditions$procedure == "bulk"], data.frame(tissue = conditions$tissue[conditions$procedure == "bulk"]), ~ tissue), parallel = T)
results.bulk <- results(dds.bulk, c("tissue", "DCIS", "macrophage"))
results.bulk.significant <- as.data.frame(results.bulk[! is.na(results.bulk$padj) & results.bulk$padj <= 0.01,])
results.bulk.significant <- results.bulk.significant[order(results.bulk.significant$log2FoldChange),]

bottom100 <- rownames(results.bulk.significant)[1:100]
top100 <- rownames(results.bulk.significant)[nrow(results.bulk.significant) - 1:100]

heatmap.frame <- melt(t(scale(t(rld[c(top100, rev(bottom100)),]))))
colnames(heatmap.frame) <- c("gene", "library", "expression")

heatmap.hundreds <- ggplot(heatmap.frame, aes(library, gene, fill = expression)) +
	geom_tile() +
	scale_fill_gradient(low = "white", high = "steelblue") +
	theme(
		legend.position = "none",
		panel.background = element_blank(),
		axis.title.x = element_blank(),
		axis.title.y = element_blank(),
		axis.text.y = element_blank(),
		axis.ticks.y = element_blank(),
		axis.text.x = element_text(angle = 90, hjust = 1, vjust = 0.5),
		axis.ticks.x = element_blank(),
		panel.grid.major.x = element_blank(),
		panel.grid.major.y = element_blank(),
		panel.grid.minor.x = element_blank(),
		panel.grid.minor.y = element_blank()
	)
ggsave("heatmap_hundreds.pdf", heatmap.hundreds, width = 6, height = 4.5)
```
