## Supplemental Code for "Gene-expression profiling of single cells from archival tissue with laser-capture microdissection and Smart-3SEQ": pc12.R

```
#! /usr/bin/Rscript

library(ggplot2)

load("conditions.RData")
load("rld.RData")

pca <- prcomp(t(rld))
prop.variance <- pca$sdev^2 / sum(pca$sdev^2)
frame.to.plot <- data.frame(conditions, pca$x)
pc12.plot <- ggplot(frame.to.plot, aes(PC1, PC2, color = tissue, shape = procedure)) +
	geom_point() +
	xlab(paste0("PC1: ", round(100 * prop.variance[1]), "% of variance")) +
	ylab(paste0("PC2: ", round(100 * prop.variance[2]), "% of variance")) +
	theme( # no axis labels because the units are arbitrary
		legend.position = c(0.85, 0.3), # fragile
		axis.text.y = element_blank(),
		axis.ticks.y = element_blank(),
		axis.text.x = element_blank(),
		axis.ticks.x = element_blank(),
			panel.grid.major = element_blank(),
			panel.grid.minor = element_blank()
	) +
	scale_color_manual(values = c(
		"DCIS" = hcl(15, 100, 60),
		"macrophage" = hcl(255, 100, 60),
		"mix" = hcl(315, 100, 60)
	))
ggsave("pc1_pc2.pdf", pc12.plot, width = 6, height = 4.5)
```
