## Supplemental Code for "Gene-expression profiling of single cells from archival tissue with laser-capture microdissection and Smart-3SEQ": correlation.R

```
#! /usr/bin/Rscript

library(parallel)
options(mc.cores = detectCores())

regularize <- function(c) log10(c + 1) # function to transform read counts so they're nice for correlations

load("counts.RData")
load("subsample.RData")
load("conditions.RData")

counts.regularized <- regularize(counts)

correlations <- sapply(subsampled, function(subsamples) {
	sapply(subsamples, function(these.counts) {
		result <- simplify2array(mclapply(colnames(these.counts), function(library) cor(regularize(these.counts[,library]), counts.regularized[,library])))
		names(result) <- colnames(these.counts)
		result
	})
})

save(correlations, file = "correlation.RData")
```
