## Supplemental Code for "Gene-expression profiling of single cells from archival tissue with laser-capture microdissection and Smart-3SEQ": genes_hit.R

```
#! /usr/bin/Rscript

gene.hit.threshold <- 10

library(parallel)
options(mc.cores = detectCores())

load("subsample.RData")
load("conditions.RData")

genes.hit <- sapply(subsampled, function(subsamples) {
	simplify2array(mclapply(subsamples, function(counts) colSums(counts >= gene.hit.threshold)))
})

save(genes.hit, gene.hit.threshold, file = "genes_hit.RData")
```
