## Supplemental Code for "Gene-expression profiling of single cells from archival tissue with laser-capture microdissection and Smart-3SEQ": graph.R

```
#! /usr/bin/Rscript

library(reshape2)
library(ggplot2)
library(scales)

load("conditions.RData")
load("genes_hit.RData")
load("correlation.RData")
load("significant_genes.RData")

result <- do.call(rbind, lapply(c("single", "bulk"), function(amount) cbind(
	rbind(
	
		cbind(
			melt(genes.hit[[amount]], varnames = c("library", "target")),
			what = "genes.hit"
		),
		
		cbind(
			melt(correlations[[amount]], varnames = c("library", "target")),
			what = "correlation"
		),
		
		cbind(
			library = amount,
			target = names(significant.genes[[amount]]),
			value = significant.genes[[amount]],
			what = "significant.genes"
		)
	),
	amount = amount
)))
result$target <- as.integer(result$target)
result$value <- as.numeric(unlist(result$value))
result$amount <- factor(as.character(result$amount))

combined.plot <- ggplot(result, aes(target / 1E6, value, color = amount, group = library)) +
	geom_line() +
	facet_wrap(~ what, scales = "free_y") +
	xlab("sequenced reads (millions)") +
	theme(axis.title.y = element_blank()) +
	scale_y_continuous(label = comma)
	
ggsave("depth_ffpe-lcm.pdf", combined.plot, "pdf", width = 12, height = 4)
```
