## Supplemental Code for "Gene-expression profiling of single cells from archival tissue with laser-capture microdissection and Smart-3SEQ": subsample.R

```
#! /usr/bin/Rscript

library(parallel)
options(mc.cores = detectCores())

increment <- 5E5

subsample <- function(counts, p) sapply(counts, function(count) rbinom(1, count, p))


load("counts.RData")
load("conditions.RData")
total.reads <- read.table("total_reads", col.names = c("library", "reads"))
total.reads.lookup <- total.reads$reads
names(total.reads.lookup) <- total.reads$library

conditions.selected <- conditions[conditions$amount != "control" & ! (conditions$tissue == "DCIS" & conditions$HER2.amp == "FALSE"),]

subsampled <- lapply(c("bulk", "single"), function(this.amount) {
	these.libraries <- rownames(subset(conditions.selected, amount == this.amount))
	counts.filtered <- counts[,these.libraries]
	targets <- increment * 1:floor(min(total.reads.lookup[these.libraries]) / increment)
	result <- mclapply(targets, function(target) {
		sapply(these.libraries, function(this.library) subsample(counts.filtered[,this.library], target / total.reads.lookup[this.library]))
	})
	names(result) <- sapply(targets, format, scientific = F)
	result
})
names(subsampled) <- c("bulk", "single")

save(subsampled, increment, file = "subsample.RData")
```
