## Supplemental Code for "Gene-expression profiling of single cells from archival tissue with laser-capture microdissection and Smart-3SEQ": significant_genes.R

```
#! /usr/bin/Rscript

library(DESeq2)
library(parallel)
options(mc.cores = detectCores())

load("subsample.RData")

significant.genes <- sapply(subsampled, function(subsamples) {
	unlist(mclapply(subsamples, function(counts) {
		sum(
			results(
				DESeq(
					DESeqDataSetFromMatrix(
						counts[rowSums(counts) > 0,],
						data.frame(template = conditions[colnames(counts),]$template),
						~ template
					),
					fitType = "local",
					quiet = T,
				)
			)$padj < 0.05,
			na.rm = T
		)
	}))
})

save(significant.genes, file = "significant_genes.RData")
```
