## Supplemental Code for "Gene-expression profiling of single cells from archival tissue with laser-capture microdissection and Smart-3SEQ": alignability.R

```
#! /usr/bin/Rscript

library(ggplot2)
library(reshape2)
library(scales)

category.colors <- c(
	"uniquely aligned" =  "green3",
	"multiply aligned" =  "darkgreen",
	"other" =             "yellow3",
	"too short" =         "yellow2",
	"RT dimer" =          "darkred",
	"PCR dimer" =         "red2"
)

read.category.counts <- read.delim("read_category_count.tsv", row.names = 1, check.names = F) # will have NAs for aligner output
read.category.counts <- read.category.counts[! (grepl("Undetermined", rownames(read.category.counts))),]
alignment.category.counts <- read.delim("alignment_category.tsv", row.names = 1)
read.category.counts[,"uniquely aligned"] <- alignment.category.counts[rownames(read.category.counts),"unique.alignment"]
read.category.counts[,"multiply aligned"] <- alignment.category.counts[rownames(read.category.counts),"multi.mapped"]
read.category.counts[,"too short"] <- alignment.category.counts[rownames(read.category.counts),"length"]
read.category.counts[,"other"] <- rowSums(alignment.category.counts[rownames(read.category.counts),c("no.mapping", "homopolymer")])

result.frame <- melt(as.matrix(read.category.counts[,-1]), varnames = c("library", "category"), value.name = "reads", as.is = T)
result.frame$category <- factor(result.frame$category, levels = c("PCR dimer", "RT dimer", "too short", "other", "multiply aligned", "uniquely aligned"))
result.frame$library <- factor(result.frame$library, levels = c(
	unlist(lapply(c("A", "B", "C", "D", "E"), function(dilution)
		lapply(c("L", "H"), function(pcr)
			lapply(1:2, function(mix) 
				lapply(1:2, function(rep) paste0(dilution, mix, pcr, "-rep", rep))
			)
		)
	)),
	unlist(lapply(c("L", "H"), function(pcr)
		lapply(1:2, function(rep) paste0("NTC", pcr, "-rep", rep))
	))
))

categories.plot <- ggplot(result.frame) +
	geom_col(aes(library, reads, fill = category), position = "fill", width = 1) +
	scale_y_continuous(label = percent, expand = c(0, 0)) +
	scale_fill_manual(values = category.colors) +
	theme(
		axis.text.x = element_blank(),
		axis.ticks.x = element_blank(),
		panel.grid.major.x = element_blank()
	)

ggsave("read_category_percent.pdf", categories.plot, "pdf", width = 10, height = 4)
```
