## Supplemental Code for "Gene-expression profiling of single cells from archival tissue with laser-capture microdissection and Smart-3SEQ": load_conc.R

```
#! /usr/bin/Rscript

ercc.lengths.table <- read.table("ercc_lengths.tsv", as.is = T)
ercc.lengths <- ercc.lengths.table[,2]
names(ercc.lengths) <- ercc.lengths.table[,1]

ercc.conc.raw <- read.table("ercc_exfold_conc.tsv", sep = "\t", header = T, as.is = T)
ercc.conc.raw <- ercc.conc.raw[order(ercc.conc.raw$ERCC.ID),]

ercc.amol.per.ul <- as.matrix(ercc.conc.raw[,grepl("^concentration", colnames(ercc.conc.raw))])
rownames(ercc.amol.per.ul) <- ercc.conc.raw$ERCC.ID
colnames(ercc.amol.per.ul) <- c("ERCC mix 1", "ERCC mix 2")

ercc.molecules.per.ul <- ercc.amol.per.ul * 6.022140857E23 * 1E-18

ercc.tpm <- 1E6 * sweep(ercc.molecules.per.ul, 2, colSums(ercc.molecules.per.ul), FUN = "/")

save(ercc.amol.per.ul, ercc.molecules.per.ul, ercc.tpm, ercc.lengths, file = "ercc_conc.RData")
```
