## Supplemental Code for "Gene-expression profiling of single cells from archival tissue with laser-capture microdissection and Smart-3SEQ": load_counts.R

```
#! /usr/bin/Rscript

# how much input? A = 1 uL @ 1/10 dilution, B @ 1/100, C @ 1/1000, etc.
# official table gives concentrations in amol/uL and they sum to 103,515
# I will round this off to 100 fmol
# so A contains 10 fmol, B 1 fmol, C 100 amol, D 10 amol, E 1 amol

dilution <- 10^-(1:5)
amount <- c("10 fmol", "1 fmol", "100 amol", "10 amol", "1 amol")
names(dilution) <- c("A", "B", "C", "D", "E")
names(amount) <- names(dilution)
pcr.cycles <- c("AH" = 11, "AL" = 7, "BH" = 15, "BL" = 11, "CH" = 18, "CL" = 14, "DH" = 22, "DL" = 17, "EH" = 25, "EL" = 21) # according to my lab notebook
pcr.category <- c("H" = "more PCR", "L" = "less PCR")

read.counts <- as.matrix(read.table("ercc_counts_withdup.tsv",
	header = T,
	sep = "\t",
	as.is = T,
	row.names = 1
))
colnames(read.counts) <- sub("\\.bam$", "", colnames(read.counts))
read.counts <- read.counts[,
	! grepl("^NTC", colnames(read.counts)) &
	! grepl("^Undetermined", colnames(read.counts))
]

conditions <- data.frame(
	row.names = colnames(read.counts),
	mix = factor(paste("ERCC mix", substr(colnames(read.counts), 2, 2))),
	dilution = dilution[substr(colnames(read.counts), 1, 1)],
	amount = factor(amount[substr(colnames(read.counts), 1, 1)], levels = amount),
	pcr.cyles = pcr.cycles[paste0(substr(colnames(read.counts), 1, 1), substr(colnames(read.counts), 3, 3))],
	pcr.category = factor(pcr.category[substr(colnames(read.counts), 3, 3)]),
	rep = sapply(colnames(read.counts), function(x) strsplit(x, "\\.")[[1]][2])
)

save(read.counts, file = "counts.RData")
save(conditions, file = "conditions.RData")
```
