## Supplemental Code for "Gene-expression profiling of single cells from archival tissue with laser-capture microdissection and Smart-3SEQ": load_seqc.R

```
#! /usr/bin/Rscript

seqc.name.to.ercc.mix <- c("E" = "ERCC mix 1", "F" = "ERCC mix 2")

load("ercc_conc.RData")

pair.counts.seqc <- as.matrix(read.table("bgi_ercc_counts_withdup.tsv",
	header = T,
	sep = "\t",
	as.is = T,
	row.names = 1
))
colnames(pair.counts.seqc) <- sub("\\.bam$", "", colnames(pair.counts.seqc))

count.per.base.seqc <- pair.counts.seqc / ercc.lengths[rownames(pair.counts.seqc)]
tpm.seqc <- 1E6 * sweep(count.per.base.seqc, 2, colSums(count.per.base.seqc), FUN = "/")

conditions.seqc <- data.frame(
	row.names = colnames(pair.counts.seqc),
	site = factor(sapply(colnames(pair.counts.seqc), function(x) strsplit(x, "_")[[1]][1])),
	mix = factor(seqc.name.to.ercc.mix[sapply(colnames(pair.counts.seqc), function(x) strsplit(x, "_")[[1]][2])]),
	rep = paste0("rep", sapply(colnames(pair.counts.seqc), function(x) strsplit(x, "_")[[1]][3]))
)

save(pair.counts.seqc, tpm.seqc, conditions.seqc, file = "tpm_seqc.RData")
```
