## Supplemental Code for "Gene-expression profiling of single cells from archival tissue with laser-capture microdissection and Smart-3SEQ": replicates.R

```
#! /usr/bin/Rscript

library(ggplot2)
library(scales)


load("counts.RData")
load("conditions.RData")

frame.to.plot <- data.frame(
	library = rep(rownames(conditions), each = nrow(read.counts)),
	mix = rep(conditions$mix[conditions$rep == "rep1"], each = nrow(read.counts)),
	amount = rep(conditions$amount[conditions$rep == "rep1"], each = nrow(read.counts)),
	pcr.category = rep(conditions$pcr.category[conditions$rep == "rep1"], each = nrow(read.counts)),
	reads.rep1 = as.vector(read.counts[,conditions$rep == "rep1"]),
	reads.rep2 = as.vector(read.counts[,conditions$rep == "rep2"])
)

correlations <- data.frame(
	mix = levels(frame.to.plot$mix),
	amount = rep(levels(frame.to.plot$amount), each = nlevels(frame.to.plot$mix)),
	pcr.category = rep(levels(frame.to.plot$pcr.category), each = nlevels(frame.to.plot$mix) * nlevels(frame.to.plot$amount))
)
correlations$corr <- apply(correlations, 1, function(row) {
	this.subset <- subset(frame.to.plot, mix == row[1] & amount == row[2] & pcr.category == row[3])
	cor(log10(this.subset$reads.rep1 + 1), log10(this.subset$reads.rep2 + 1), method = "pearson")
})

replicate.comparison <- ggplot(frame.to.plot, aes(reads.rep1, reads.rep2)) +
	geom_point(size = 1/2) +
	geom_text(data = correlations, aes(label = paste0("r*minute == ", round(corr, 3))), x = -Inf, y = Inf, hjust = -0.2, vjust = 1.2, parse = T) + 
	scale_x_log10(labels = comma) +
	scale_y_log10(labels = comma) +
	xlab("Smart-3SEQ reads in replicate 1") +
	ylab("Smart-3SEQ reads in replicate 2") +
	facet_grid(mix ~ amount * pcr.category) +
	theme(axis.text.x = element_text(angle = 45, hjust = 1))

ggsave("replicates.pdf", replicate.comparison, "pdf", width = 12, height = 9)
```
