## Supplemental Code for "Gene-expression profiling of single cells from archival tissue with laser-capture microdissection and Smart-3SEQ": sensitivity.R

```
#! /usr/bin/Rscript

library(ggplot2)
library(scales)

load("counts.RData")
load("conditions.RData")

frame.to.plot <- data.frame(
	library = rep(rownames(subset(conditions, amount != "10 fmol")), each = nrow(read.counts)),
	reads.this.dilution = as.vector(read.counts[,conditions$amount != "10 fmol"]),
	reads.first.dilution = as.vector(read.counts[,conditions$amount == "10 fmol"])
)
frame.to.plot <- cbind(frame.to.plot, conditions[frame.to.plot$library,])

correlations <- data.frame(library = levels(frame.to.plot$library))
correlations$corr <- sapply(correlations$library, function(this.library) cor(log10(subset(frame.to.plot, library == this.library)$reads.this.dilution + 1), log10(subset(frame.to.plot, library == this.library)$reads.first.dilution + 1)))
correlations <- cbind(correlations, conditions[correlations$library,])

sensitivity.plot <- ggplot(frame.to.plot, aes(reads.first.dilution, reads.this.dilution)) +
	geom_point(size = 1/2) +
	geom_text(data = correlations, aes(label = paste0("r*minute == ", round(corr, 3))), x = -Inf, y = Inf, hjust = -0.2, vjust = 1.2, parse = T) + 
	scale_x_log10(labels = comma) +
	scale_y_log10(labels = comma) +
	xlab("reads in first dilution") +
	ylab("reads in this dilution") +
	facet_grid(mix * rep ~ amount * pcr.category) +
	theme(axis.text.x = element_text(angle = 45, hjust = 1))

ggsave("sensitivity_3seq.pdf", sensitivity.plot, "pdf", width = 12, height = 9)
```
