## Supplemental Code for "Gene-expression profiling of single cells from archival tissue with laser-capture microdissection and Smart-3SEQ": stdcrv.R

```
#! /usr/bin/Rscript

library(ggplot2)
library(scales)
library(reshape2)

best.library <- "A2L.rep1" # the one with the lowest PCR cycles and highest count, for simplified figures

load("ercc_conc.RData")
load("conditions.RData")
load("counts.RData")
load("counts_3M.RData")

pdf("stdcrv_3seq_subsample_original.pdf", width = 12, height = 9) # note this will contain the subsampled one before the original one
for (read.counts.to.plot in list(read.counts.3M, read.counts)) {

	frame.to.plot <- melt(read.counts.to.plot, varnames = c("transcript", "library"), value.name = "reads")
	frame.to.plot <- cbind(frame.to.plot, conditions[frame.to.plot$library,])
	frame.to.plot$molecules <- frame.to.plot$dilution * sapply(1:nrow(frame.to.plot), function(i) ercc.molecules.per.ul[frame.to.plot$transcript[i], frame.to.plot$mix[i]])

	expected.lines <- data.frame(
		library = levels(frame.to.plot$library),
		slope = colSums(read.counts.to.plot) / sapply(levels(frame.to.plot$library), function(library) sum(frame.to.plot$molecules[frame.to.plot$library == library])),
		corr = sapply(levels(frame.to.plot$library), function(library) cor(log10(frame.to.plot[frame.to.plot$library == library, "molecules"]), log10(frame.to.plot[frame.to.plot$library == library, "reads"] + 1), method = "pearson")) # Pearson correlation of log10(molecules), log10(count + 1)
	)
	expected.lines <- cbind(expected.lines, conditions[expected.lines$library,])

	standard.curve.all <- ggplot(frame.to.plot, aes(molecules, reads)) +
		geom_point(size = 1/2) +
#		geom_abline(data = expected.lines, aes(slope = 1, intercept = log10(slope))) +
		geom_text(data = expected.lines, aes(label = paste0("r*minute == ", round(corr, 3))), x = -Inf, y = Inf, hjust = -0.2, vjust = 1.2, parse = T) + 
		scale_x_log10(labels = comma) +
		scale_y_log10(labels = comma) +
		xlab("expected copy number") +
		ylab("Smart-3SEQ reads") +
		facet_grid(mix * rep ~ amount * pcr.category, scales = "free_x") +
		theme(axis.text.x = element_text(angle = 45, hjust = 1))

	print(standard.curve.all)
}
dev.off()

# frame.to.plot etc. is still defined from the non-subsampled data
standard.curve.best <- ggplot(subset(frame.to.plot, library == best.library), aes(molecules, reads)) +
	geom_point() +
#	geom_abline(data = subset(expected.lines, library == best.library), aes(slope = 1, intercept = log10(slope))) + # intercept = log10(slope) because we're in a log-log plot, i.e. log10(y) = log10(bx) = log10(b) + log10(x)
	scale_x_log10(labels = comma) +
	scale_y_log10(labels = comma) +
	xlab("expected copy number") +
	ylab("Smart-3SEQ reads")
cat(best.library, "r′ =", expected.lines[best.library, "corr"], "\n")

ggsave("stdcrv_3seq_best.pdf", standard.curve.best, "pdf", width = 4, height = 4)
```
