## Supplemental Code for "Gene-expression profiling of single cells from archival tissue with laser-capture microdissection and Smart-3SEQ": stdcrv_seqc.R

```
#! /usr/bin/Rscript

library(ggplot2)
library(scales)
library(reshape2)

best.library <- "BGI_F_2" # the one with the highest count (and it's mix 2 to match Smart-3SEQ), for simplified figures

load("ercc_conc.RData")
load("tpm_seqc.RData")
load("tpm_seqc_3M.RData")

pdf("stdcrv_seqc.pdf", width = 8, height = 6) # note this will contain the subsampled one before the original one

for (read.counts.to.plot in list(tpm.seqc, tpm.seqc.3M)) {
	frame.to.plot <- melt(tpm.seqc, varnames = c("transcript", "library"), value.name = "tpm")
	frame.to.plot <- cbind(frame.to.plot, conditions.seqc[frame.to.plot$library,])
	frame.to.plot$true.tpm <- sapply(1:nrow(frame.to.plot), function(i) ercc.tpm[frame.to.plot$transcript[i], frame.to.plot$mix[i]])

	expected.lines.seqc <- data.frame(
		library = levels(frame.to.plot$library),
		corr = sapply(levels(frame.to.plot$library), function(library) cor(log10(frame.to.plot[frame.to.plot$library == library, c("true.tpm", "tpm")]), method = "pearson")[1,2]) # Pearson correlation of log10(TPM)
	)
	expected.lines.seqc <- cbind(expected.lines.seqc, conditions.seqc[expected.lines.seqc$library,])

	standard.curve.all <- ggplot(frame.to.plot, aes(true.tpm, tpm)) +
		geom_point() +
#		geom_abline(aes(slope = 1, intercept = 0)) +
		geom_text(data = expected.lines.seqc, aes(label = paste0("r == ", round(corr, 3))), x = -Inf, y = Inf, hjust = -0.2, vjust = 1.2, parse = T) + 
		scale_x_log10(labels = comma) +
		scale_y_log10(labels = comma) +
		xlab("true TPM") +
		ylab("RNA-seq TPM") +
		facet_wrap(~ mix * rep) +
		theme(axis.text.x = element_text(angle = 45, hjust = 1))
	
	print(standard.curve.all)
}
dev.off()

# frame.to.plot etc. is still defined from the non-subsampled data
standard.curve.best <- ggplot(subset(frame.to.plot, library == best.library), aes(true.tpm, tpm)) +
	geom_point() +
#	geom_abline(aes(slope = 1, intercept = 0)) +
	scale_x_log10(labels = comma) +
	scale_y_log10(labels = comma) +
	xlab("true TPM") +
	ylab("RNA-seq TPM")
cat(best.library, "r =", expected.lines.seqc[best.library, "corr"], "\n")

ggsave(paste0("stdcrv_seqc_best.pdf"), standard.curve.best, "pdf", width = 4, height = 4)
```
