## Supplemental Code for "Gene-expression profiling of single cells from archival tissue with laser-capture microdissection and Smart-3SEQ": subsample.R

```
#! /usr/bin/Rscript

target <- 3E6

source("subsample_function.R")
load("counts.RData")

total.reads.table <- read.table("total_reads.tsv", as.is = T)
total.reads <- total.reads.table[,2]
names(total.reads) <- make.names(total.reads.table[,1])

read.counts.3M <- subsample.counts(read.counts, target, total.reads)

save(read.counts.3M, file = "counts_3M.RData")
```
