## Supplemental Code for "Gene-expression profiling of single cells from archival tissue with laser-capture microdissection and Smart-3SEQ": subsample_function.R

```
subsample.counts <- function(read.counts, target, totals) {
	stopifnot(colnames(read.counts) %in% names(totals))
	p <- target / totals
	sapply(colnames(read.counts)[totals[colnames(read.counts)] >= target], function(library) sapply(read.counts[,library], function(count) rbinom(1, count, p[library])))
}
```
