## Supplemental Code for "Gene-expression profiling of single cells from archival tissue with laser-capture microdissection and Smart-3SEQ": subsample_seqc.R

```
#! /usr/bin/Rscript

target <- 3E6

source("subsample_function.R")
load("tpm_seqc.RData")
load("ercc_conc.RData")

total.pairs.table <- read.table("total_pairs_bgi.tsv", as.is = T)
total.pairs <- total.pairs.table[,2]
names(total.pairs) <- make.names(total.pairs.table[,1])

pair.counts.seqc.3M <- subsample.counts(pair.counts.seqc, target, total.pairs)

count.per.base.seqc.3M <- pair.counts.seqc.3M / ercc.lengths[rownames(pair.counts.seqc.3M)]
tpm.seqc.3M <- 1E6 * sweep(count.per.base.seqc.3M, 2, colSums(count.per.base.seqc.3M), FUN = "/")

save(pair.counts.seqc.3M, tpm.seqc.3M, file = "tpm_seqc_3M.RData")
```
