## Supplemental Code for "Gene-expression profiling of single cells from archival tissue with laser-capture microdissection and Smart-3SEQ": cnv.R

```
#! /usr/bin/Rscript

library(reshape2)
library(ggplot2)

bin.size <- 1E6 # 1 Mb
chr.to.exclude <- c("chrY", "chrM")
selected.chr <- "chr17"
selected.gene <- "ERBB2"

load("counts.RData")
load("conditions.RData")
load("rld.RData")
load("gene_id.RData")

libraries.to.exclude <- rownames(conditions)[conditions$tissue == "control"]

chr.lengths.table <- read.table("hg38_chrom_sizes")
chr.lengths <- chr.lengths.table[,2]
names(chr.lengths) <- chr.lengths.table[,1]

annotations.filtered <- counts.raw[rowSums(counts) > 0, 2:5]
rownames(annotations.filtered) <- rownames(counts.filtered)

annotations.filtered <- data.frame(
	chr = sapply(annotations.filtered$Chr, function(string) {
		result <- unique(strsplit(string, ";")[[1]])
		stopifnot(length(result) == 1)
		result
	}),
	strand = sapply(annotations.filtered$Strand, function(string) {
		result <- unique(strsplit(string, ";")[[1]])
		stopifnot(length(result) == 1)
		result
	}),
	left = as.integer(sapply(annotations.filtered$Start, function(string) strsplit(string, ";")[[1]][1])),
	right = as.integer(sapply(annotations.filtered$End, function(string) tail(strsplit(string, ";")[[1]], 1))),
	row.names = rownames(annotations.filtered),
	stringsAsFactors = FALSE
)
annotations.filtered$end <- as.integer(apply(annotations.filtered, 1, function(row) if (row["strand"] == "+") row["right"] else row["left"]))

selected.gene.end <- annotations.filtered$end[gene.name == selected.gene]
selected.gene.end <- selected.gene.end[! is.na(selected.gene.end)] # it might be listed more than once
stopifnot(length(selected.gene.end) == 1)

bins <- lapply(chr.lengths, function(chr.length) {
	n.bin <- ceiling(chr.length / bin.size)
	matrix(0, nrow = n.bin, ncol = ncol(counts.filtered), dimnames = list((1:n.bin - 1 / 2) * bin.size, colnames(counts.filtered))) # add 1/2 bin size so that each bin's rowname will be its *center*, since this is how it's graphed later with geom_raster
})

for (i in 1:nrow(rld)) {
	this.chr <- annotations.filtered$chr[i]
	this.bin <- ceiling(annotations.filtered$end[i] / bin.size)
	bins[[this.chr]][this.bin,] <- bins[[this.chr]][this.bin,] + rld[i,]
}

bins.scaled <- lapply(bins[! names(bins) %in% chr.to.exclude], function(bin) { # exclude the excluded ones at this step because otherwise chrM probably only has one row, so the subset doesn't return a matrix, so rowMeans fails
	result <- bin - rowMeans(bin[,conditions$amount == "bulk" & conditions$tissue == "macrophage"])
	result[rowMeans(bin) == 0,] <- NA
	result
})


heatmap.frame <- melt(bins.scaled)
colnames(heatmap.frame) <- c("bin.start", "library", "relative.expression", "chr")
heatmap.frame$chr <- factor(heatmap.frame$chr, levels = names(chr.lengths))
heatmap.frame$library <- factor(heatmap.frame$library, levels = colnames(counts.filtered)[rev(display.order)]) # reverse display order because geom_raster goes bottom to top and we want top to bottom

pdf("heatmap_by_chr.pdf", width = 10, height = 7.5)
	for (chr in names(bins.scaled)) {
		heatmap.chr <- ggplot(heatmap.frame[heatmap.frame$chr == chr & (! heatmap.frame$library %in% libraries.to.exclude),], aes(bin.start / bin.size, library, fill = relative.expression)) +
			geom_tile() +
			coord_cartesian(expand = FALSE) +
			scale_fill_gradient2(na.value = "lightgray") +
			theme(
				legend.position = "none",
				axis.title.y = element_blank(),
				axis.ticks.y = element_blank()
			) +
			xlab("position (Mb)") +
			ggtitle(chr)
		if (chr == selected.chr) print(heatmap.chr + geom_vline(xintercept = selected.gene.end / bin.size, col = "red")) else print(heatmap.chr)
	}
dev.off()
```
