## Supplemental Code for "Gene-expression profiling of single cells from archival tissue with laser-capture microdissection and Smart-3SEQ": comparison.R

```
#! /usr/bin/Rscript

library(DESeq2)

load("counts.RData")
load("conditions.RData")

# bulk DCIS vs. bulk macrophage
dds.bulk <- DESeq(DESeqDataSetFromMatrix(counts.filtered[,conditions$amount == "bulk"], data.frame(tissue = conditions$tissue[conditions$amount == "bulk"]), ~ tissue), parallel = T, fitType = "local")
results.bulk <- results(dds.bulk, c("tissue", "DCIS", "macrophage"))

# single DCIS (only with HER2-amp) vs. single macrophage
which.single.amp <- which(conditions$amount == "single" & (conditions$HER2.amp == TRUE | conditions$tissue == "macrophage"))
dds.single <- DESeq(DESeqDataSetFromMatrix(counts.filtered[,which.single.amp], data.frame(tissue = conditions$tissue[which.single.amp]), ~ tissue), parallel = T, fitType = "local")
results.single <- results(dds.single, c("tissue", "DCIS", "macrophage"))

# single DCIS HER2-amp vs. no amp
dds.amp <- DESeq(DESeqDataSetFromMatrix(counts.filtered[,conditions$amount == "single" & conditions$tissue == "DCIS"], data.frame(HER2.amp = conditions$HER2.amp[conditions$amount == "single" & conditions$tissue == "DCIS"]), ~ HER2.amp), parallel = T, fitType = "local")
results.amp <- results(dds.amp, c("HER2.amp", T, F))

# single DCIS no amp vs. single macrophage
which.single.noamp <- which(conditions$amount == "single" & (conditions$HER2.amp == FALSE | conditions$tissue == "macrophage"))
dds.site <- DESeq(DESeqDataSetFromMatrix(counts.filtered[,which.single.noamp], data.frame(tissue = conditions$tissue[which.single.noamp]), ~ tissue), parallel = T, fitType = "local")
results.site <- results(dds.site, c("tissue", "DCIS", "macrophage"))

# single vs. bulk, controlling for cell tissue
which.not.weird <- ! (conditions$amount == "control" | (conditions$amount == "single" & conditions$tissue == "DCIS" & conditions$HER2.amp == FALSE))
dds.single.vs.bulk <- DESeq(DESeqDataSetFromMatrix(counts.filtered[,which.not.weird], conditions[which.not.weird,], ~ amount + tissue), parallel = T, fitType = "local")
results.single.vs.bulk <- results(dds.single.vs.bulk, c("amount", "single", "bulk"))

# cell tissue, controlling for single vs. bulk
which.not.weird <- ! (conditions$amount == "control" | (conditions$amount == "single" & conditions$tissue == "DCIS" & conditions$HER2.amp == FALSE))
results.tissue <- results(dds.single.vs.bulk, c("tissue", "DCIS", "macrophage"))

save(dds.bulk, results.bulk, dds.single, results.single, dds.amp, results.amp, dds.site, results.site, dds.single.vs.bulk, results.single.vs.bulk, results.tissue, file = "comparison.RData")
```
