## Supplemental Code for "Gene-expression profiling of single cells from archival tissue with laser-capture microdissection and Smart-3SEQ": expression_table.R

```
#! /usr/bin/Rscript

library(DESeq2) # required to coerce "results" objects into data frames
library(WriteXLS)

load("counts.RData")
load("rld.RData")
load("comparison.RData")
load("gene_id.RData")

WriteXLS(
	lapply(c(
		list(
			counts.filtered,
			round(tpm, 1),
			round(rld, 3)
		), lapply(list(
			results.bulk,
			results.single,
			results.amp,
			results.site,
			results.single.vs.bulk,
			results.tissue
		), function(x) round(as.data.frame(x), 4))
	), function(frame) data.frame("Ensembl ID" = ensembl.gene.id, "gene name" = gene.name, frame, check.names = F)),
	ExcelFileName = "gene_expression.xlsx",
	SheetNames = c(
		"count",
		"TPM",
		"rlog",
		"bulk comparison",
		"single comparison",
		"amp comparison",
		"site comparison",
		"single vs. bulk",
		"DCIS vs. macrophage"
	),
	AdjWidth = T,
	BoldHeaderRow = T,
	FreezeRow = 1,
	FreezeCol = 2
)
```
