## Supplemental Code for "Gene-expression profiling of single cells from archival tissue with laser-capture microdissection and Smart-3SEQ": gene_id.R

```
#! /usr/bin/Rscript

library(biomaRt)

load("counts.RData")

ensembl = useMart("ENSEMBL_MART_ENSEMBL", dataset = "hsapiens_gene_ensembl", host = "uswest.ensembl.org")
ensembl.gene.id <- sub("\\..*", "", rownames(counts.filtered))
gene.name.table <- getBM(
	attributes =  c("ensembl_gene_id", "external_gene_name"),
	filters =     "ensembl_gene_id",
	values =      ensembl.gene.id,
	mart =        ensembl
)
rownames(gene.name.table) <- gene.name.table$ensembl_gene_id
gene.name = gene.name.table[ensembl.gene.id, "external_gene_name"]

save(ensembl.gene.id, gene.name, file = "gene_id.RData")
```
