## Supplemental Code for "Gene-expression profiling of single cells from archival tissue with laser-capture microdissection and Smart-3SEQ": load_conditions.R

```
#! /usr/bin/Rscript

load("counts.RData")

amount <- factor(sapply(colnames(counts), function(name) sub(".*_", "", sub("_[^_]+$", "", name))), levels = c("bulk", "single", "control"))
tissue <- factor(sapply(colnames(counts), function(library.name) if (grepl("DCIS", library.name)) "DCIS" else if (grepl("macrophage", library.name)) "macrophage" else if (grepl("control", library.name)) "control"), levels = c("macrophage", "DCIS", "control"))

conditions <- data.frame(amount, tissue)
# add annotation of HER2 amplification as detected from sliding-window analysis
conditions$HER2.amp <- F
conditions[scan("which_her2_amp.txt", character()), "HER2.amp"] <- T
conditions$HER2.amp <- factor(conditions$HER2.amp)

display.order <- c( # nice display order to group the DCIS & ductal macrophage separately
	which(conditions$amount == "bulk" & conditions$tissue == "macrophage"),
	which(conditions$amount == "bulk" & conditions$tissue == "DCIS"),
	which(conditions$amount == "single" & conditions$tissue == "macrophage"),
	which(conditions$amount == "single" & conditions$tissue == "DCIS" & conditions$HER2.amp == FALSE),
	which(conditions$amount == "single" & conditions$tissue == "DCIS" & conditions$HER2.amp == TRUE),
	which(conditions$amount == "control")
)
write(rownames(conditions)[display.order], file="display_order")

save(conditions, display.order, file = "conditions.RData")
```
