## Supplemental Code for "Gene-expression profiling of single cells from archival tissue with laser-capture microdissection and Smart-3SEQ": load_counts.R

```
#! /usr/bin/Rscript

counts.raw <- read.table("counts_withdup.tsv",
	sep = "\t",
	as.is = T,
	check.names = F,
	header = T
)
counts.raw <- counts.raw[! grepl("_PAR_Y$", counts.raw$Geneid),] # remove Y PAR genes since they have duplicate names
counts.raw <- counts.raw[,! grepl("Undetermined", colnames(counts.raw))]
#counts.raw <- counts.raw[! grepl("chr[MY]", counts.raw$Chr),] # not interested in mitochondrial genome and shouldn't see Y chromosome

counts <- as.matrix(counts.raw[,-c(1:6)])
rownames(counts) <- sub("\\..*", "", counts.raw$Geneid)
colnames(counts) <- sub("\\..*", "", sapply(colnames(counts), basename))
names(colnames(counts)) <- colnames(counts)

counts <- counts[,! (grepl("ablation", colnames(counts)) | grepl("mix", colnames(counts)))] # remove the ablation experiment from this analysis

counts.filtered <- counts[rowSums(counts) > 0,]

tpm <- 1E6 * sweep(counts.filtered, 2, colSums(counts.filtered), "/")

save(counts.raw, counts, counts.filtered, tpm, file = "counts.RData")
```
