## Supplemental Code for "Gene-expression profiling of single cells from archival tissue with laser-capture microdissection and Smart-3SEQ": marker_genes.R

```
#! /usr/bin/Rscript

marker.genes <- c("CD68", "CD163", "EPCAM", "KRT7", "KRT18", "ERBB2")

library(reshape2)
library(ggplot2)

load("counts.RData")
load("conditions.RData")
load("gene_id.RData")

which.marker.genes <- which(gene.name %in% marker.genes)
frame.marker.genes <- melt(tpm[which.marker.genes,], varnames = c("gene.id", "library"), value.name = "tpm")
frame.marker.genes$gene.symbol <- factor(gene.name[which.marker.genes], levels = marker.genes)
frame.marker.genes <- cbind(frame.marker.genes, conditions[frame.marker.genes$library,])
frame.marker.genes$category <- factor(paste(frame.marker.genes$tissue, frame.marker.genes$amount), levels = c("macrophage bulk", "macrophage single", "DCIS bulk", "DCIS single"))

plot.marker.genes <- ggplot(subset(frame.marker.genes, tissue != "control"), aes(category, tpm, color = tissue, shape = amount, solid = HER2.amp)) +
	geom_jitter(width = .1) +
	facet_grid(~ gene.symbol) +
	scale_y_log10() +
	scale_color_manual(values = c(
		"DCIS" = hcl(15, 100, 60),
		"macrophage" = hcl(255, 100, 60),
		"control" = hcl(315, 100, 60)
	)) +
	ylab("gene expression (TPM)") +
	theme(
		axis.text.x = element_blank(),
		axis.title.x = element_blank(),
		axis.ticks.x = element_blank(),
		panel.grid.major.x = element_blank()
	) +
	geom_point(data = subset(frame.marker.genes, amount == "single" & tissue == "DCIS" & HER2.amp == FALSE), col = "black", shape = 1, size = 2.5) # circle the HER2- DCIS single cells, but the circles will be horizontally off because of the jittering (fix manually)

ggsave("marker_genes.pdf", plot.marker.genes, width = 7.5, height = 3)
```
