## Supplemental Code for "Gene-expression profiling of single cells from archival tissue with laser-capture microdissection and Smart-3SEQ": rld.R

```
#! /usr/bin/Rscript

# this one takes a few minutes

library(DESeq2)

load("counts.RData")
load("conditions.RData")

dds <- DESeq(DESeqDataSetFromMatrix(counts.filtered, conditions, ~ 1), parallel = T, fitType = "local") # blind to conditions because we want to use this for an unbiased rlog (and nothing else)

rld <- assay(rlog(dds, blind = F)) # "blind = F" means it won't redo the mean-variance fitting, which was already blind, but was done with parallelization, which is impossible with rlog for some reason

save(dds, rld, file = "rld.RData")
```
