## Supplemental Code for "Gene-expression profiling of single cells from archival tissue with laser-capture microdissection and Smart-3SEQ": single_vs_bulk.R

```
#! /usr/bin/Rscript

library(ggplot2)

load("counts.RData")
load("conditions.RData")

mean.bulk.dcis <- rowMeans(tpm[,conditions$amount == "bulk" & conditions$tissue == "DCIS"])
mean.bulk.macrophage <- rowMeans(tpm[,conditions$amount == "bulk" & conditions$tissue == "macrophage"])
mean.single.dcis <- rowMeans(tpm[,conditions$amount =="single" & conditions$tissue == "DCIS" & conditions$HER2.amp == TRUE])
mean.single.macrophage <- rowMeans(tpm[,conditions$amount =="single" & conditions$tissue == "macrophage"])

pdf("single_vs_bulk.pdf", width = 10, height = 7)
	ggplot(data.frame(single = c(mean.single.dcis, mean.single.macrophage), bulk = c(mean.bulk.dcis, mean.bulk.macrophage), type = rep(c("DCIS", "macrophage"), each = nrow(counts.filtered))), aes(bulk, single)) +
		geom_hex() +
		scale_x_log10() +
		scale_y_log10() +
		facet_grid(~ type) +
		xlab("mean TPM in bulk") +
		ylab("mean TPM in single cells") +
		geom_abline(slope = 1, col = "red")
	ggplot(data.frame(dcis = c(mean.single.dcis, mean.bulk.dcis), macrophage = c(mean.single.macrophage, mean.bulk.macrophage), amount = rep(c("single", "bulk"), each = nrow(counts.filtered))), aes(macrophage, dcis)) +
		geom_hex() +
		scale_x_log10() +
		scale_y_log10() +
		facet_grid(~ amount) +
		xlab("mean TPM in macrophage") +
		ylab("mean TPM in DCIS") +
		geom_abline(slope = 1, col = "red")
dev.off()
```
