## Supplemental Code for "Gene-expression profiling of single cells from archival tissue with laser-capture microdissection and Smart-3SEQ": tsne.R

```
#! /usr/bin/Rscript

library(parallel)
library(ggplot2)
library(Rtsne)

options(mc.cores = detectCores())

load("conditions.RData")
load("rld.RData")

# remove controls
rld <- rld[,conditions$amount != "control"]
conditions <- subset(conditions, amount != "control")

# distance matrix (Euclidean)
dist.mat <- dist(t(rld))

# t-SNE
tsne.iter <- mclapply(1:floor((ncol(rld) - 1) / 3), function(p) Rtsne(dist.mat,
	perplexity = p,
	check_duplicates = F
)) # this takes a few seconds
tsne.frame.iter <- lapply(tsne.iter, function(tsne.result) {
	coords <- tsne.result$Y
	colnames(coords) <- c("x", "y")
	data.frame(coords, conditions)	
	}
)
pdf("tsne.pdf", width = 4, height = 3)
for (p in 1:length(tsne.iter)) {
	print(ggplot(tsne.frame.iter[[p]], aes(x, y, color = tissue, shape = amount)) +
		geom_point() +
		ggtitle(paste0("perplexity = ", p, ", cost =", sum(tsne.iter[[p]]$costs))) + 
		theme( # no axis labels because the units are arbitrary
			axis.title.y = element_blank(),
			axis.text.y = element_blank(),
			axis.ticks.y = element_blank(),
			axis.title.x = element_blank(),
			axis.text.x = element_blank(),
			axis.ticks.x = element_blank(),
			panel.grid.major = element_blank(),
			panel.grid.minor = element_blank()
		) +
		scale_color_manual(values = c(
			"DCIS" = hcl(15, 100, 60),
			"macrophage" = hcl(255, 100, 60)
		)) +
		geom_point(data = subset(tsne.frame.iter[[p]], amount == "single" & tissue == "DCIS" & HER2.amp == FALSE), col = "black", shape = 1, size = 2.5) # circle the HER2- DCIS single cells
	)
}
dev.off()
```
