## Supplemental Code for "Gene-expression profiling of single cells from archival tissue with laser-capture microdissection and Smart-3SEQ": ref-rna_read_length.R

```
#! /usr/bin/Rscript

library(ggplot2)
library(scales)

template <- c(H = "HBRR", U = "UHRR")
amount <- c(A = "100 ng", B = "10 ng", C = "1 ng", D = "100 pg", E = "10 pg")

total.reads <- read.table("ref-rna_total_reads", as.is = T)
total.reads.lookup <- total.reads[,2]
names(total.reads.lookup) <- total.reads[,1]
star.logs <- read.table("ref-rna_aggregate_logs.tsv", sep = "\t", header = T, as.is = T)

star.results <- data.frame(
	run = sub("\\.align\\.log$", "", star.logs$library),
	library = sub("\\..*", "", star.logs$library),
	uniquely.aligned = star.logs$Uniquely.mapped.reads.number,
	template = factor(template[substr(star.logs$library, 1, 1)]),
	amount = factor(amount[substr(star.logs$library, 3, 3)], levels = amount),
	length = as.integer(sub("^trunc", "", sapply(star.logs$library, function(x) strsplit(x, "\\.")[[1]][2])))
)

star.results$percent.uniquely.aligned <- star.results$uniquely.aligned / total.reads.lookup[as.character(star.results$library)]


result.plot <- ggplot(star.results, aes(length, percent.uniquely.aligned, group = library, color = amount)) +
	geom_line() +
	geom_vline(xintercept = c(51, 76, 101, 151), col = "darkgray", linetype = "dashed") +
	scale_x_continuous(breaks = c(50, 100, 150)) + # if I let it go automatically it uses increments of 30, which are hard to read
	scale_y_continuous(label = percent) +
	xlab("read length (nt)") +
	ylab("uniquely aligned reads")

ggsave("read_length_ref-rna.pdf", result.plot, width = 8, height = 4)
```
