## Supplemental Code for "Gene-expression profiling of single cells from archival tissue with laser-capture microdissection and Smart-3SEQ": compare_3seq_seqc.R

```
#! /usr/bin/Rscript

fudge.factor.3seq <- 0.1
fudge.factor.seqc <- 0.0001

library(ggplot2)
library(scales)

load("tpm_seqc.RData")
load("counts.RData")

stopifnot(nrow(read.counts) == nrow(tpm.seqc))

tpm.3seq <- 1E6 * sweep(read.counts, 2, colSums(read.counts), FUN = "/")

frame.to.plot <- data.frame(
	sample = rep(c("HBRR", "UHRR"), each = nrow(read.counts)),
	tpm.seqc = c(rowMeans(tpm.seqc[,conditions.seqc$template == "HBRR"]), rowMeans(tpm.seqc[,conditions.seqc$template == "UHRR"])),
	tpm.3seq = c(rowMeans(tpm.3seq[,conditions$template == "HBRR" & conditions$amount == "100 ng"]), rowMeans(tpm.3seq[,conditions$template == "UHRR" & conditions$amount == "100 ng"]))
)

correlations <- data.frame(
	sample = levels(frame.to.plot$sample),
  corr = sapply(levels(frame.to.plot$sample), function(this.sample) cor(subset(frame.to.plot, sample == this.sample)$tpm.seqc, subset(frame.to.plot, sample == this.sample)$tpm.3seq, method = "spearman"))
)

seq.comparison <- ggplot(frame.to.plot, aes(tpm.seqc + fudge.factor.seqc, tpm.3seq + fudge.factor.3seq)) +
	geom_hex() +
  geom_text(data = correlations, aes(label = paste0("rho == ", round(corr, 3))), x = -Inf, y = Inf, hjust = -0.2, vjust = 1.2, parse = T) +  
	scale_x_log10(label = comma) +
	scale_y_log10(label = comma) +
	facet_wrap(~ sample) +
	xlab(paste("RNA-seq mean TPM +", format(fudge.factor.seqc, scientific = F))) +
	ylab(paste("Smart-3SEQ mean TPM +", format(fudge.factor.3seq, scientific = F))) +
	scale_fill_continuous(trans = "log10", name = "genes")

ggsave("compare_3seq_seqc.pdf", seq.comparison, "pdf", width = 8, height = 4)
```
