## Supplemental Code for "Gene-expression profiling of single cells from archival tissue with laser-capture microdissection and Smart-3SEQ": compare_3seq_sybr.R

```
#! /usr/bin/Rscript

library(ggplot2)
library(scales)

best.library <- "U1A" # the one with the highest input and highest count, for simplified figures

load("counts.RData")
load("sybr.RData")

sybr.cq.common <- sybr.cq[rownames(sybr.cq) %in% rownames(read.counts),]


frame.to.plot <- data.frame(
	gene.id = rownames(sybr.cq.common),
	cq = as.vector(sybr.cq.common[,as.character(conditions$template)]),
	reads = as.vector(read.counts[rownames(sybr.cq.common),]),
	library = rep(colnames(read.counts), each = nrow(sybr.cq.common)),
	template = rep(conditions$template, each = nrow(sybr.cq.common)),
	replicate = sub("mix", "rep", rep(conditions$ercc.mix, each = nrow(sybr.cq.common))),
	amount = rep(conditions$amount, each = nrow(sybr.cq.common))
)

expected.lines <- data.frame(
	library = levels(frame.to.plot$library),
  corr = sapply(levels(frame.to.plot$library), function(this.library) cor(subset(frame.to.plot, library == this.library)$cq, subset(frame.to.plot, library == this.library)$reads, method = "spearman", use = "complete.obs"))
)
expected.lines <- cbind(expected.lines, conditions[expected.lines$library,])

result <- ggplot(frame.to.plot, aes(cq, reads + 1)) +
	geom_hex() +
  geom_text(data = expected.lines, aes(label = paste0("rho == ", round(corr, 3))), x = -Inf, y = Inf, hjust = -0.2, vjust = 1.2, parse = T) +  
	scale_x_reverse() +
	scale_y_log10(label = comma) +
	xlab("SYBR qPCR measurement (Cq)") +
	ylab("Smart-3SEQ measurement (read count + 1)") +
	facet_grid(template * replicate ~ amount) +
	scale_fill_continuous(trans = "log10", name = "genes")

result.best <- ggplot(subset(frame.to.plot, library == best.library), aes(cq, reads + 1)) +
	geom_hex() +
	scale_x_reverse() +
	scale_y_log10(label = comma) +
	xlab("SYBR qPCR measurement (Cq)") +
	ylab("Smart-3SEQ measurement (read count + 1)") +
	scale_fill_continuous(trans = "log10", name = "genes")

cat(best.library, "rho =", expected.lines[best.library, "corr"], "\n")

ggsave("compare_3seq_sybr.pdf", result, "pdf", width = 8, height = 6)
ggsave("compare_3seq_sybr_best.pdf", result.best, "pdf", width = 8, height = 6)
```
