## Supplemental Code for "Gene-expression profiling of single cells from archival tissue with laser-capture microdissection and Smart-3SEQ": compare_3seq_taqman.R

```
#! /usr/bin/Rscript

library(ggplot2)
library(scales)

best.library <- "U1A" # the one with the highest input and highest count, for simplified figures

load("counts.RData")
load("taqman.RData")

taqman.expr.common <- taqman.expr[rownames(taqman.expr) %in% rownames(read.counts),]


frame.to.plot <- data.frame(
	gene.id = rownames(taqman.expr.common),
	expr = as.vector(taqman.expr.common[,as.character(conditions$template)]),
	reads = as.vector(read.counts[rownames(taqman.expr.common),]),
	library = rep(colnames(read.counts), each = nrow(taqman.expr.common)),
	template = rep(conditions$template, each = nrow(taqman.expr.common)),
	replicate = rep(conditions$replicate, each = nrow(taqman.expr.common)),
	amount = rep(conditions$amount, each = nrow(taqman.expr.common))
)

correlations <- data.frame(
	library = levels(frame.to.plot$library),
  corr = sapply(levels(frame.to.plot$library), function(this.library) cor(subset(frame.to.plot, library == this.library)$expr, subset(frame.to.plot, library == this.library)$reads, method = "spearman"))
)
correlations <- cbind(correlations, conditions[correlations$library,])

result <- ggplot(frame.to.plot, aes(expr, reads)) +
	geom_point(size = 1/10) +
  geom_text(data = correlations, aes(label = paste0("rho == ", round(corr, 3))), x = -Inf, y = Inf, hjust = -0.2, vjust = 1.2, parse = T) +  
	scale_x_log10() +
	scale_y_log10(label = comma) +
	xlab("qPCR measurement (relative expression)") +
	ylab("Smart-3SEQ reads") +
	facet_grid(template * replicate ~ amount)

result.best <- ggplot(subset(frame.to.plot, library == best.library), aes(expr, reads)) +
	geom_point() +
	scale_x_log10() +
	scale_y_log10(label = comma) +
	xlab("qPCR measurement (relative expression)") +
	ylab("Smart-3SEQ reads")
cat(best.library, "rho =", correlations[best.library, "corr"], "\n")

ggsave("compare_3seq_taqman.pdf", result, "pdf", width = 8, height = 6)
ggsave("compare_3seq_taqman_best.pdf", result.best, "pdf", width = 4, height = 4)
```
