## Supplemental Code for "Gene-expression profiling of single cells from archival tissue with laser-capture microdissection and Smart-3SEQ": compare_seqc_sybr.R

```
#! /usr/bin/Rscript

fudge.factor.seqc <- 0.001

library(ggplot2)
library(scales)

load("tpm_seqc.RData")
load("sybr.RData")

sybr.cq.common <- sybr.cq[rownames(sybr.cq) %in% rownames(tpm.seqc),]


frame.to.plot <- data.frame(
	gene.id = rownames(sybr.cq.common),
	cq = as.vector(sybr.cq.common[,as.character(conditions.seqc$template)]),
	tpm = as.vector(tpm.seqc[rownames(sybr.cq.common),]),
	library = rep(colnames(tpm.seqc), each = nrow(sybr.cq.common)),
	template = rep(conditions.seqc$template, each = nrow(sybr.cq.common)),
	replicate = sub("mix", "rep", rep(conditions.seqc$replicate, each = nrow(sybr.cq.common)))
)

correlations <- data.frame(
	library = levels(frame.to.plot$library),
  corr = sapply(levels(frame.to.plot$library), function(this.library) cor(subset(frame.to.plot, library == this.library)$cq, subset(frame.to.plot, library == this.library)$tpm, method = "spearman", use = "complete.obs"))
)
correlations <- cbind(correlations, conditions.seqc[correlations$library,])


result.all <- ggplot(frame.to.plot, aes(cq, tpm + fudge.factor.seqc)) +
	geom_hex() +
  geom_text(data = correlations, aes(label = paste0("rho == ", round(corr, 3))), x = -Inf, y = Inf, hjust = -0.2, vjust = 1.2, parse = T) +  
	scale_x_reverse() +
	scale_y_log10(label = comma) +
	xlab("SYBR qPCR measurement (Cq)") +
	ylab(paste("RNA-seq TPM +", format(fudge.factor.seqc, scientific = F))) +
	facet_grid(template ~ replicate) +
	scale_fill_continuous(trans = "log10", name = "genes")

ggsave("compare_seqc_sybr.pdf", result.all, "pdf", width = 8, height = 6)
```
