## Supplemental Code for "Gene-expression profiling of single cells from archival tissue with laser-capture microdissection and Smart-3SEQ": compare_sybr_taqman.R

```
#! /usr/bin/Rscript

library(ggplot2)
library(scales)
library(reshape2)

load("taqman.RData")
load("sybr.RData")

common.genes <- intersect(rownames(taqman.expr), rownames(sybr.cq))

frame.to.plot <- data.frame(
	gene.id = rep(common.genes, 2),
	sample = rep(c("HBRR", "UHRR"), each = length(common.genes)),
	taqman.expr = c(taqman.expr[common.genes, "HBRR"], taqman.expr[common.genes, "UHRR"]),
	primepcr.cq = c(sybr.cq[common.genes, "HBRR"], sybr.cq[common.genes, "UHRR"])
)

correlations <- data.frame(
	sample = levels(frame.to.plot$sample),
  corr = sapply(levels(frame.to.plot$sample), function(this.sample) cor(subset(frame.to.plot, sample == this.sample)$taqman.expr, subset(frame.to.plot, sample == this.sample)$primepcr.cq, method = "spearman", use = "complete.obs"))
)

sybr.comparison <- ggplot(frame.to.plot, aes(taqman.expr, primepcr.cq)) +
	geom_point(size = 1/10) +
  geom_text(data = correlations, aes(label = paste0("rho == ", round(corr, 3))), x = -Inf, y = Inf, hjust = -0.2, vjust = 1.2, parse = T) +  
	scale_x_log10() +
	scale_y_reverse() +
	facet_wrap(~ sample) +
	xlab("TaqMan measurement (relative expression)") +
	ylab("SYBR measurement (Cq)")

ggsave("compare_sybr_taqman.pdf", sybr.comparison, "pdf", width = 8, height = 4)
```
