## Supplemental Code for "Gene-expression profiling of single cells from archival tissue with laser-capture microdissection and Smart-3SEQ": load_counts.R

```
#! /usr/bin/Rscript

template <- c("H" = "HBRR", "U" = "UHRR")
amount <- c("A" = "100 ng", "B" = "10 ng", "C" = "1 ng", "D" = "100 pg", "E" = "10 pg")

counts.raw <- read.table("counts_withdup.tsv",
	header = T,
	sep = "\t",
	as.is = T,
	skip = 1
)

read.counts <- as.matrix(counts.raw[,
	grepl("_run2\\.bam$", colnames(counts.raw)) &
	! grepl("^Undetermined_run", colnames(counts.raw)) &
	! grepl("^NTC", colnames(counts.raw))
])
colnames(read.counts) <- sub("_run2\\.bam$", "", colnames(read.counts))
rownames(read.counts) <- sub("\\..*", "", counts.raw$Geneid)

conditions <- data.frame(
	row.names = colnames(read.counts),
	template = factor(template[substr(colnames(read.counts), 1, 1)]),
	ercc.mix = factor(paste0("mix", substr(colnames(read.counts), 2, 2))),
	amount = factor(amount[substr(colnames(read.counts), 3, 3)], levels = amount)
)
conditions$replicate <- sub("mix", "rep", conditions$ercc.mix)

save(read.counts, conditions, file = "counts.RData")
```
