## Supplemental Code for "Gene-expression profiling of single cells from archival tissue with laser-capture microdissection and Smart-3SEQ": load_seqc.R

```
#! /usr/bin/Rscript

seqc.name.to.template <- c("A" = "UHRR", "B" = "HBRR")


counts.raw <- read.table("bgi_hg38_counts_withdup.tsv",
	header = T,
	sep = "\t",
	as.is = T,
	skip = 1
)

fragment.counts.seqc <- as.matrix(counts.raw[,grepl("\\.bam$", colnames(counts.raw))])
colnames(fragment.counts.seqc) <- sub("\\.bam$", "", colnames(fragment.counts.seqc))
rownames(fragment.counts.seqc) <- sub("\\..*", "", counts.raw$Geneid)

gene.lengths <- counts.raw$Length
names(gene.lengths) <- rownames(fragment.counts.seqc)

count.per.base.seqc <- fragment.counts.seqc / gene.lengths[rownames(fragment.counts.seqc)]
tpm.seqc <- 1E6 * sweep(count.per.base.seqc, 2, colSums(count.per.base.seqc), FUN = "/")

conditions.seqc <- data.frame(
	row.names = colnames(fragment.counts.seqc),
	template = factor(seqc.name.to.template[sapply(colnames(fragment.counts.seqc), function(x) strsplit(x, "_")[[1]][2])]),
	replicate = paste0("rep", sapply(colnames(fragment.counts.seqc), function(x) strsplit(x, "_")[[1]][3]))
)

save(fragment.counts.seqc, tpm.seqc, conditions.seqc, file = "tpm_seqc.RData")
```
