## Supplemental Code for "Gene-expression profiling of single cells from archival tissue with laser-capture microdissection and Smart-3SEQ": load_sybr.R

```
#! /usr/bin/Rscript

sybr.raw <- read.table("GSE56457_SEQC_qPCR_GEOSub.txt",
	header = T,
	sep = "\t",
	comment.char = "",
	quote = "\"",
	as.is = T
)

sybr.cq <- as.matrix(sybr.raw[,c("SEQC_RTPCR_B", "SEQC_RTPCR_A")])
rownames(sybr.cq) <- sybr.raw$ensembl_gene
colnames(sybr.cq) <- c("HBRR", "UHRR")
sybr.cq[sybr.cq == 0] <- NA # they put 0 for missing values, I guess

save(sybr.cq, file = "sybr.RData")
```
