## Supplemental Code for "Gene-expression profiling of single cells from archival tissue with laser-capture microdissection and Smart-3SEQ": load_taqman.R

```
#! /usr/bin/Rscript

gene.convert.table <- read.table("MAQC_TaqMan_GPL4097/refseq_to_ensgene.tsv", sep = "\t", fill = T, as.is = T, header = T, quote = "")
gene.lookup <- gene.convert.table$To
names(gene.lookup) <- gene.convert.table$From

assay.desc <- read.table("MAQC_TaqMan_GPL4097/assay_description_GPL4097.tsv", sep = "\t", header = T, as.is = T)
refseq.id <- lapply(assay.desc$GB_LIST, function(x) grep("^NM_", strsplit(x, ",")[[1]], value = T))
ensembl.id <- lapply(refseq.id, function(refseq.ids) {
	result <- gene.lookup[refseq.ids]
	names(result) <- NULL
	result
})
ensembl.id.clean <- sapply(ensembl.id, function(ensembl.ids) {
	if (length(ensembl.ids) == 0) {
		return(NA)
	} else if (length(ensembl.ids) == 1) {
		return(ensembl.ids[1])
	} else if (length(unique(ensembl.ids) == 1)) {
		return(ensembl.ids[1])
	} else {
		return(NA)
	}
})

taqman.expr.raw <- sapply(Sys.glob("MAQC_TaqMan_GPL4097/*_GSM*.tsv"), function(data.file) read.table(data.file, sep = "\t", header = T, strip.white = T)[,2])
colnames(taqman.expr.raw) <- sub("\\.tsv$", "", basename(colnames(taqman.expr.raw)))
template <- sub("_.*", "", colnames(taqman.expr.raw))

taqman.expr <- t(sapply(sort(unique(ensembl.id.clean[! is.na(ensembl.id.clean)])), function(gene) sapply(unique(template), function(this.template) mean(taqman.expr.raw[which(ensembl.id.clean == gene), which(template == this.template)]))))

save(taqman.expr, file = "taqman.RData")
```
